# Models and Algorithms for Equilibrium Analysis of Mixed-Material Nucleic Acid Systems

**DOI:** 10.1101/2025.06.30.662484

**Authors:** Avinash Nanjundiah, Mark E. Fornace, Samuel J. Schulte, Niles A. Pierce

## Abstract

Dynamic programming algorithms within the NUPACK software suite enable analysis of equilibrium base-pairing properties for complex and test tube ensembles containing arbitrary numbers of interacting nucleic acid strands. Currently, calculations are limited to single-material systems that are either all-RNA or all-DNA. Here, to enable analysis of mixed-material systems that are critical for modern applications in vitro, in situ, and in vivo, we develop physical models and dynamic programming algorithms that allow the material of the system to be specified at nucleotide resolution. Free energy parameter sets are constructed for both RNA/DNA and RNA/2^′^OMe-RNA mixed-material systems by combining available empirical mixed-material parameters with single-material parameter sets to enable treatment of the full complex and test tube ensembles. New dynamic programming recursions account for the material of each nucleotide throughout the recursive process. For a complex with *N* nucleotides, the mixed-material dynamic programming algorithms maintain the *O*(*N* ^3^) time complexity of the single-material algorithms, enabling efficient calculation of diverse physical quantities over complex and test tube ensembles (e.g., complex partition function, equilibrium complex concentrations, equilibrium base-pairing probabilities, minimum free energy secondary structure(s), and Boltzmann-sampled secondary structures) at a cost increase of roughly 2.0–3.5*×* . The results of existing single-material algorithms are exactly reproduced when applying the new mixed-material algorithms to single-material systems. Accuracy is significantly enhanced using mixed-material models and algorithms to predict RNA/DNA and RNA/2^′^OMe-RNA duplex melting temperatures from the experimental literature as well as RNA/DNA melt profiles from new experiments. Mixed-material analyses can be performed online using the NUPACK web app (www.nupack.org) or locally using the NUPACK Python module.^¶^

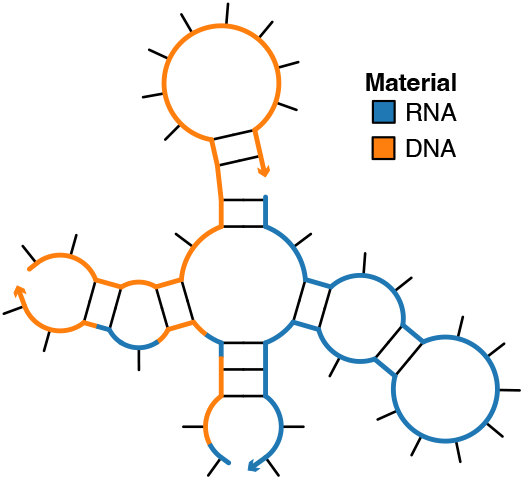

## Introduction

Mixed-material nucleic acid interactions are common in life sciences applications. For example, hybridization of a DNA probe to an RNA target in an in situ imaging experiment^1, 2^ illustrates the *hybrid case*, in which a nucleic acid strand of one material hybridizes to a nucleic acid strand of another material. Or, for example, hybridization of a probe containing a mixture of RNA and 2^′^OMe-RNA to an RNA target in an in vivo regulation experiment^3, 4^ illustrates the *chimeric case*, in which a nucleic acid strand contains more than one material. However, equilibrium analysis algorithms usually require that only one material is treated in a given calculation (e.g., all-RNA or all-DNA).^5–10^ An algorithm has been developed for equilibrium analysis of RNA/DNA dimers for the hybrid case where one strand is all-RNA and one strand is all-DNA.^11^ Currently, no models or algorithms exist for analyzing mixed-material systems over complex and test tube ensembles containing arbitrary numbers of interacting strand species for either the hybrid or chimeric cases. Here, we develop these models and algorithms, allowing the material composition to be specified at nucleotide resolution.

Using a nucleic acid secondary structure model, a given secondary structure is decomposed into its constituent loops (Figure 1a) and the free energy of the secondary structure is estimated via a sum of empirical loop free energies. One challenge to the formulation of mixed-material physical models is the scarcity of empirical loop free energies for mixed-material loops. To enable modeling of mixed-material duplexes, empirical free energy parameters have been measured for stack loops for a variety of mixed materials including RNA/DNA duplexes,^12–16^ RNA/2^′^OMe-RNA duplexes,^17^ a mixture of 2^′^OMe-RNA and LNA duplexed with RNA,^18^ and a mixture of LNA and DNA duplexed with DNA.^19^ A partial set of internal mismatches have also been measured for RNA/DNA duplexes.^20^ To construct mixed-material parameter sets encompassing all loop types, we combine available stack loop and internal mismatch free energies (which enable calculation of the free energy only for duplex structures with or without internal mismatches) with single-material parameter sets for other loop types (to enable modeling of the full complex and test tube ensembles). For a mixed-material secondary structure, the parameters used for each loop depend on the material composition of the loop. We define mixed-material free energy parameter sets for both RNA/DNA systems and RNA/2^′^OMe-RNA systems.

**Figure 1:**
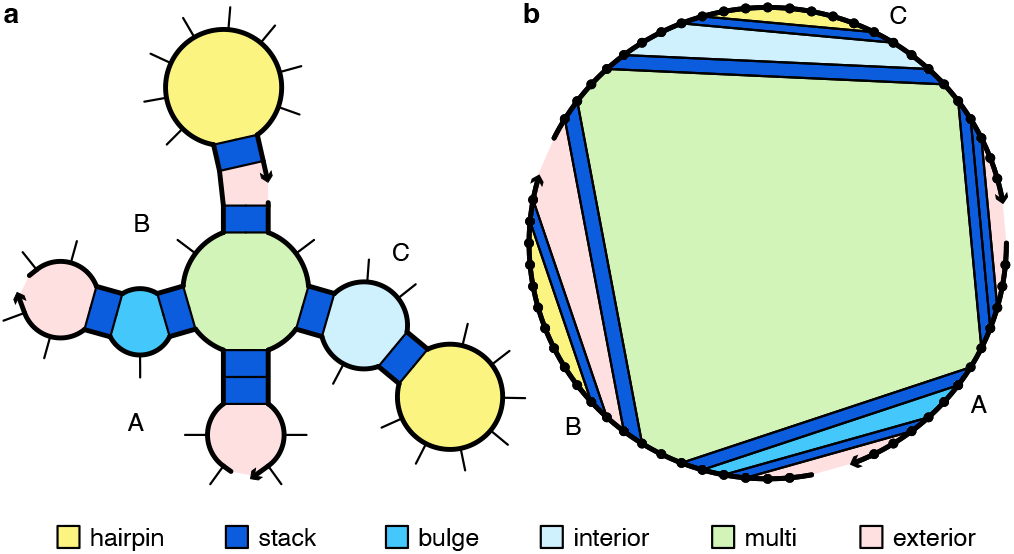
Loop-based free energy model for a complex. (a) Canonical loop types for complex with strand ordering *π* = ABC. (b) Equivalent polymer graph representation. An arrowhead denotes the 3^′^ end of each strand. Adapted with permission from Fornace et al., *ACS Synth Biol*, 9, 2665-2678, 2020. Copyright 2020 American Chemical Society.

Dynamic programming algorithms enable efficient calculation of nucleic acid base-pairing properties over complex and test tube ensembles by building up information about the full sequence recursively from calculations on subsequences of increasing length. Within the new mixed-material dynamic programming algorithms, the appropriate single- or mixed-material free energy parameters (e.g., RNA, DNA, or RNA/DNA) must be used for each loop depending on the material composition of the loop. For some loop types, contributions are built up recursively for multiple loops simultaneously without knowing the full composition of each loop at any single recursive level, necessitating the definition of separate single- and mixed-material recursions that apply the correct parameters to each loop encountered in the recursive process. We describe mixed-material dynamic programming algorithms that apply the appropriate free energy model to each loop in a mixed-material system while exactly reproducing single-material results when applied to single-material systems.

To assess the utility of the new mixed-material models and algorithms, we compare computed and empirical duplex melting temperatures for several literature data sets. As a further reality check, we performed new experimental melt studies to compare computed and empirical melt curves and melting temperatures for hybrid duplexes synthesized with each of four different strand compositions: DNA:DNA, DNA:RNA, RNA:DNA, and RNA:RNA.

## Mixed-Material Free Energy Models

### Complex and Test Tube Ensembles

NUPACK algorithms operate over two fundamental ensembles:^10, 21–23^

- *Complex ensemble:* The ensemble of all (unpseudoknotted connected) secondary structures for an arbitrary number of interacting nucleic acid strands.
- *Test tube ensemble:* The ensemble of a dilute solution containing an arbitrary number of nucleic acid strand species (introduced at user-specified concentrations) interacting to form an arbitrary number of complex species.

Furthermore, to enable reaction pathway engineering of dynamic hybridization cascades (e.g., shape and sequence transduction using small conditional RNAs^24^) or large-scale structural engineering including pseudoknots (e.g., RNA origamis^25^), NUPACK algorithms operate on multi-tube ensembles.^10, 26, 27^

The sequence, *ϕ*, of one or more interacting strands is specified as a list of nucleotides *ϕ*^*a*^ for *a* = 1, …, |*ϕ*|, where *ϕ*^*a*^ specifies both the material and base. For RNA nucleotides, *ϕ*^*a*^ ∈ {rA, rC, rG, rU}. For DNA nucleotides, *ϕ*^*a*^ ∈ {dA, dC, dG, dT} . For 2^′^OMe-RNA nucleotides, *ϕ*^*a*^ ∈ {mA, mC, mG, mU} . Hence, for a mixed-material system (RNA/DNA or RNA/2^′^OMe-RNA), the sequence alphabet contains eight letters rather than four. By convention, sequences are specified 5^′^ to 3^′^ and the material need only be specified when there is a change in material (e.g., rACGdAT denotes a strand with 3 RNA nucleotides followed by 2 DNA nucleotides).

A secondary structure, *s*, of one or more interacting RNA strands is defined by a set of base pairs, each a Watson–Crick pair or a wobble pair. An example secondary structure is displayed in Figure 1a. For RNA base pairs, Watson–Crick pairs are rA·rU and rC·rG and wobble pairs are rG·rU. For DNA base pairs, Watson– Crick pairs are dA·dT and dC·dG and there are no wobble pairs (dG×dT is considered to form a mismatch).^28^ For 2^′^OMe-RNA base pairs, Watson–Crick pairs are mA·mU and mC·mG and wobble pairs are mG·mU. For RNA/DNA mixed-material base pairs, Watson– Crick pairs are rG·dC, dG·rC, rA·dT, dA·rU and wobble pairs are rG·dT; dG and rU are considered to form mismatch dG×rU and not a wobble pair.^29^ For RNA/2^′^OMe-RNA mixed-material base pairs, Watson–Crick pairs are rG·mC, mG·rC, rA·mU, and mA·rU and wobble pairs are rG·mU and mG·rU.^17^

For algorithmic purposes, it is convenient to describe secondary structures using a *polymer graph* representation, constructed by ordering the strands around a circle, drawing the backbones in succession from 5^′^ to 3^′^ around the circumference with a *nick* between each strand, and drawing straight lines connecting paired bases (e.g., Figure 1b). A secondary structure is *unpseudoknotted* if there exists a strand ordering for which the polymer graph has no crossing lines, or *pseudoknotted* if all strand orderings contain crossing lines. A secondary structure is *connected* if no subset of the strands is free of the others.

Consider a *complex* of *L* distinct strands (e.g., each with a unique identifier in {1, …, *L*}) corresponding to strand ordering *π*. The complex ensemble 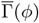 contains all connected polymer graphs with no crossing lines for sequence *ϕ*and strand ordering *π* (i.e., all unpseudoknotted secondary structures).^21^ Out of algorithmic necessity, dynamic programs operate on complex ensemble 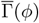 treating all strands as distinct.^10, 21^ However, in the laboratory, strands with the same sequence are typically indistinguishable with respect to experimental observables. For comparison to experimental data, physical quantities calculated over ensemble 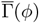 can be post-processed to obtain the corresponding quantities calculated over ensemble Γ(*ϕ*) in which strands with the same sequence are treated as indistinguishable.^10^

A *test tube* ensemble is a dilute solution containing a set of strand species, Ψ^0^, introduced at user-specified concentrations, that interact to form a set of complex species, Ψ, each corresponding to a different strand ordering treating strands with the same sequence as indistinguishable.

### Loop-based free energy model

For each (unpseudoknotted connected) secondary structure 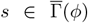, the free energy, 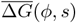, is estimated as the sum of the empirically determined free energies of the constituent loops^5,30–34^ plus a strand association penalty,^35^ *G*^assoc^, applied *L −* 1 times for a complex of *L* strands:

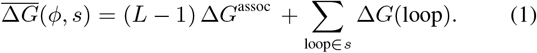

The secondary structure and polymer graph of Figure 1 illustrate the different loop types, with free energies modeled as follows:^5,30–34^

- ·A *hairpin loop* is closed by a single base-pair *i·j*. The loop free energy, 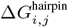, depends on sequence and loop size.
- An *interior loop* is closed by two base pairs (*i·j* and *d·e* with *i < d < e < j*). The loop free energy, 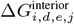depends on sequence, loop size, and loop asymmetry. *Bulge loops* (where either *d* = *i* + 1 or *e* = *j*−1) and *stack loops* (where both *d* = *i* + 1 and *e* = *j* − 1) are treated as special cases of interior loops.
- A *multiloop* is closed by three or more base pairs. The loop free energy is modeled as the sum of three sequence-independent penalties: (1) 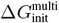 for formation of a multiloop, (2) 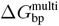 for each closing base pair, (3) 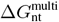 for each unpaired nucleotide inside the multiloop; plus two sequence-dependent terms: (4) 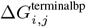, a penalty for each closing base pair *i*·*j*, (5) 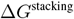, an optional coaxial and dangle stacking bonus.
- An *exterior loop* contains a nick between strands and any number of closing base pairs. The exterior loop free energy is the sum of 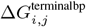 over all closing base pairs *i*·*j* and an optional coaxial and dangle stacking bonus, Δ*G*^stacking^. Hence, an unpaired strand has a free energy of zero, corresponding to the reference state.^21^

Within a multiloop or an exterior loop, a closing base pair can form one *coaxial stack* with an adjacent closing base pair or can form a *dangle stack* with at most two adjacent unpaired bases; unpaired bases can form a dangle stack with at most one adjacent closing base pair (see Section S2.2 for details). NUPACK dynamic programming algorithms provide the option to incorporate all possible coaxial and dangle stacking states in the complex ensemble, with each secondary structure incorporating Boltzmann-weighted contributions from its subensemble of stacking states.^10^

For a secondary structure *s ∈* Γ(*ϕ*) with an *R*-fold rotational symmetry there is an *R*-fold reduction in distinguishable conformational space, so the free energy (1) must be adjusted by a symmetry correction:^21^

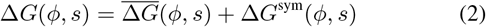

where

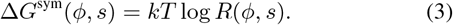

Because the symmetry factor *R*(*ϕ, s*) is a global property of each secondary structure *s* Γ(*ϕ*), it is not suitable for use with dynamic programs that treat multiple subproblems simultaneously without access to global structural information. As a result, dynamic programs operate on ensemble 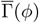 using physical model (1), after which the Distinguishability Correction Theorem of Dirks et al.^21^ enables exact conversion of physical quantities to ensemble Γ(*ϕ*) using physical model (2). Interestingly, ensembles 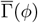 and Γ(*ϕ*) both have utility when examining the physical properties of a complex as they provide related but different perspectives, akin to complementary thought experiments (see Section S2.7).

Single-material loop-based free energy models are defined using empirical temperature-dependent parameter sets that provide loop free energies for either RNA^5, 31^–^34, 36, 37^ or DNA.^5, 28, 30, 38, 39^ Salt corrections enable DNA calculations in user-specified concentrations of 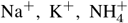, and Mg^++^.^28, 30, 38, 40^ Based on empirical data,^14, 41, 42^ we further apply the salt correction to enable RNA calculations in user-specified concentrations of Na^+^ (see Section S2.4). Additionally, we construct a single-material parameter set for 2^′^OMe-RNA in 0.12 M Na^+^ for use in the context of mixed-material RNA/2^′^OMe-RNA calculations.^17^ See Section S3 for details on RNA, DNA, and 2^′^OMe-RNA single-material parameter sets.

### Mixed-material free energy parameter sets

By contrast, empirical free energies have been measured for relatively few mixed-material loops (RNA/DNA or RNA/2^′^OMe-RNA). Here, we construct mixed-material parameter sets by leveraging empirical mixed-material loop free energies when possible and drawing on existing single-material parameter sets when necessary. Figure 2 illustrates a mixed-material secondary structure for a complex containing both mixed-material RNA/DNA loops and single-material RNA loops and DNA loops. With the new mixed-material physical model, the parameters used for each loop depend on the material composition of the loop.

**Figure 2:**
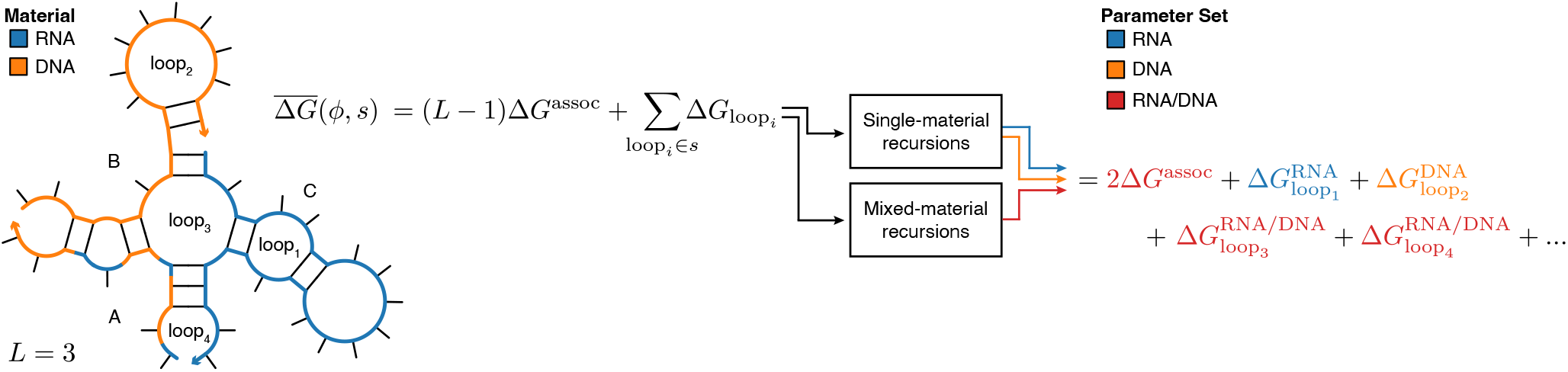
Overview of the mixed-material model and algorithm. Complex ABC consists of a chimeric RNA/DNA strand (A), a DNA strand (B), and an RNA strand (C). Each loop in a given secondary structure, 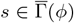, is either all-RNA (e.g., loop_1_), all-DNA (e.g., loop_2_), or a mixture of RNA and DNA (e.g., loop_3_ and loop_4_). Whereas current single-material algorithms use parameters for one material (either RNA or DNA) for all loops regardless of loop composition, the new mixed-material algorithms combine single- and mixed-material recursions to use the appropriate parameter set based on the composition of each loop (e.g., RNA parameters for loop_1_, DNA parameters for loop_2_, and mixed-material RNA/DNA parameters for loop_3_ and loop_4_).

The new mixed-material temperature-dependent parameter set for RNA/DNA (see Section S4 for details) leverages empirical free energies for the full set of 16 hybrid stack loops (rr/dd) for Watson–Crick base pairing,^13^ a partial set of hybrid internal mismatches (rA×dA, rC×dC, rG×dG, rU×dT, rG×dT, rU×dG, rA×dG, rG×dA, rA×dC, rU×dC, rC×dA, rC×dT),^20, 29^ a partial set of chimeric stack loops,^15^ and empirical trends in chimeric stack loop free energies.^15^ Other parameters for which mixed-material RNA/DNA experiments were not available were derived by averaging RNA-only and DNA-only single-material parameters. A weighted average was used for sequence-dependent parameters (depending on the number or RNA and DNA nucleotides in a given loop), and a simple average was used for size-dependent parameters (as for these parameters, the information need to calculate a weighted average is not available during the dynamic programming recursions). Based on empirical data,^14,16^ we further apply the DNA salt correction to enable mixed-material RNA/DNA calculations in user-specified concentrations of Na^+^ (see Section S2.4).

The new mixed-material temperature-dependent parameter set for RNA/2^′^OMe-RNA (see Section S5 for details) leverages empirical free energies for the full set of 16 hybrid stack loops (rr/mm) for Watson–Crick base pairing,^17^ the hybrid strand association penalty,^17^ and the terminal base pair penalty, 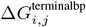.^17^ Other parameters for which mixed-material RNA/2^′^OMe-RNA experiments were not available use single-material RNA parameters. Mixed-material RNA/2^′^OMe-RNA calculations are supported in 0.12 M Na^+^ to match the experimental conditions that were used for parameterization studies.^17^

As new empirical parameters become available, whether for different experimental conditions, additional loop types, or new materials, single- and mixed-material parameter sets can be updated and used to improve the accuracy and relevance of future calculations.

## Mixed-Material Dynamic Programming Algorithms

We previously described a unified dynamic programming framework that enables calculation of diverse physical quantities for interacting nucleic acid strands using three ingredients:^10^ 1) *recursions* that specify the dependencies between subproblems and incorporate the details of the structural ensemble and the free energy model, 2) *evaluation algebras* that define the mathematical form of each subproblem, 3) *operation orders* that specify the computational trajectory through the dependency graph of subproblems. The recursions are coded generically and then compiled with a quantity-specific evaluation algebra and operation order to generate an executable for calculation of each physical quantity: partition function, equilibrium base-pairing probabilities, MFE energy and proxy structure, suboptimal structures, and Boltzmann-sampled structures. Here, to convey the essence of the new mixed-material algorithms, we focus on describing recursions for calculation of the partition function without coaxial and dangle stacking. These ideas are then generalized to dynamic programs for all of the above physical quantities using the unified dynamic programming framework.^10^

### Single-material algorithm recursion overview

For a complex with sequence *ϕ*, the partition function,

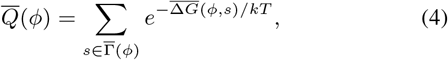

over ensemble 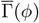 can be used to calculate the equilibrium probability of any secondary structure 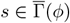:

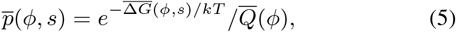

where *k* is the Boltzmann constant and *T* is temperature.

Before describing the new mixed-material partition function recursions, it is helpful to briefly summarize the recursions for the single-material algorithm that operate on complex ensemble 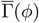.^10, 21^ The complex ensemble size, 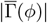, grows exponentially with the number of nucleotides, *N ≡ ∣ϕ ∣*, but the partition function can be calculated in *O*(*N* ^3^) time and *O*(*N* ^2^) space using a dynamic programming algorithm.^10, 21, 43^ The algorithm calculates the subsequence partition function *Q*_*i,j*_ for each subsequence [*i, j*] via a forward sweep from short subsequences to the full sequence (Figure 3), finally yielding the partition function of the full sequence, *Q*_1,*N*_ . The recursions used to calculate *Q*_*i,j*_ from previously calculated subsequence partition functions can be depicted as recursion diagrams (Figure 4 left; with free energy contributions colored to match the loop types of Figure 1) or equivalently using recursion equations (Figure 4 right). The *Q* recursion relies on additional restricted partition functions *Q*^*s*^ and *Q*^*b*^, and *Q*^*b*^ in turn relies on restricted partition functions *Q*^*m*^ and *Q*^*ms*^, all of which are calculated recursively. Collectively, the *Q, Q*^*s*^, *Q*^*b*^, *Q*^*m*^, and *Q*^*ms*^ recursions yield 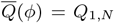, incorporating the partition function contributions of every secondary structure, 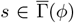, based on free energy model (1) treating all strands as distinct.

**Figure 3:**
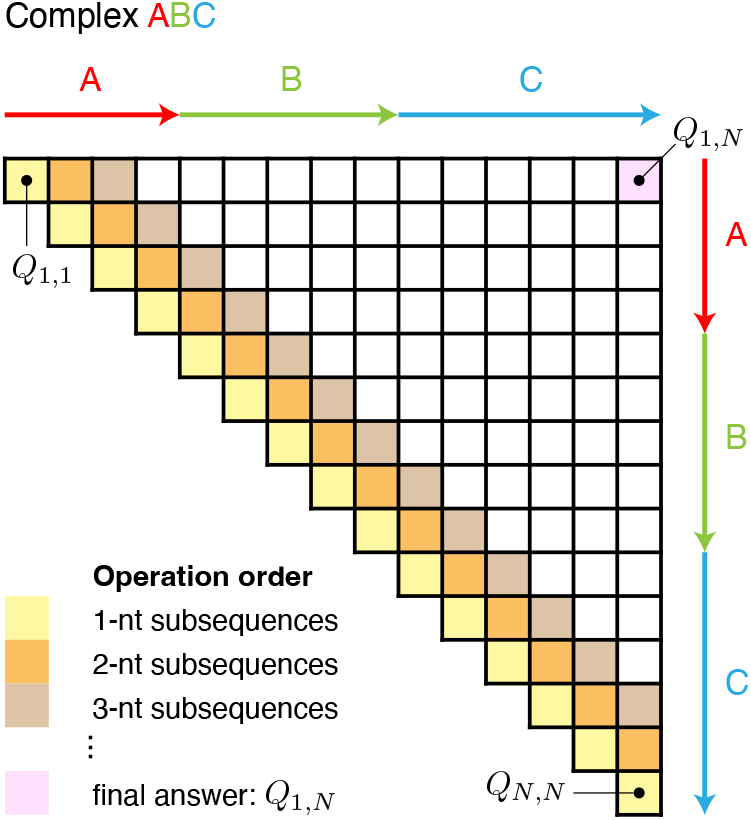
Operation order for partition function dynamic program over a complex ensemble with *N* nucleotides. Used with permission from Fornace et al., *ACS Synth Biol*, 9, 2665-2678, 2020. Copyright 2020 American Chemical Society.

**Figure 4:**
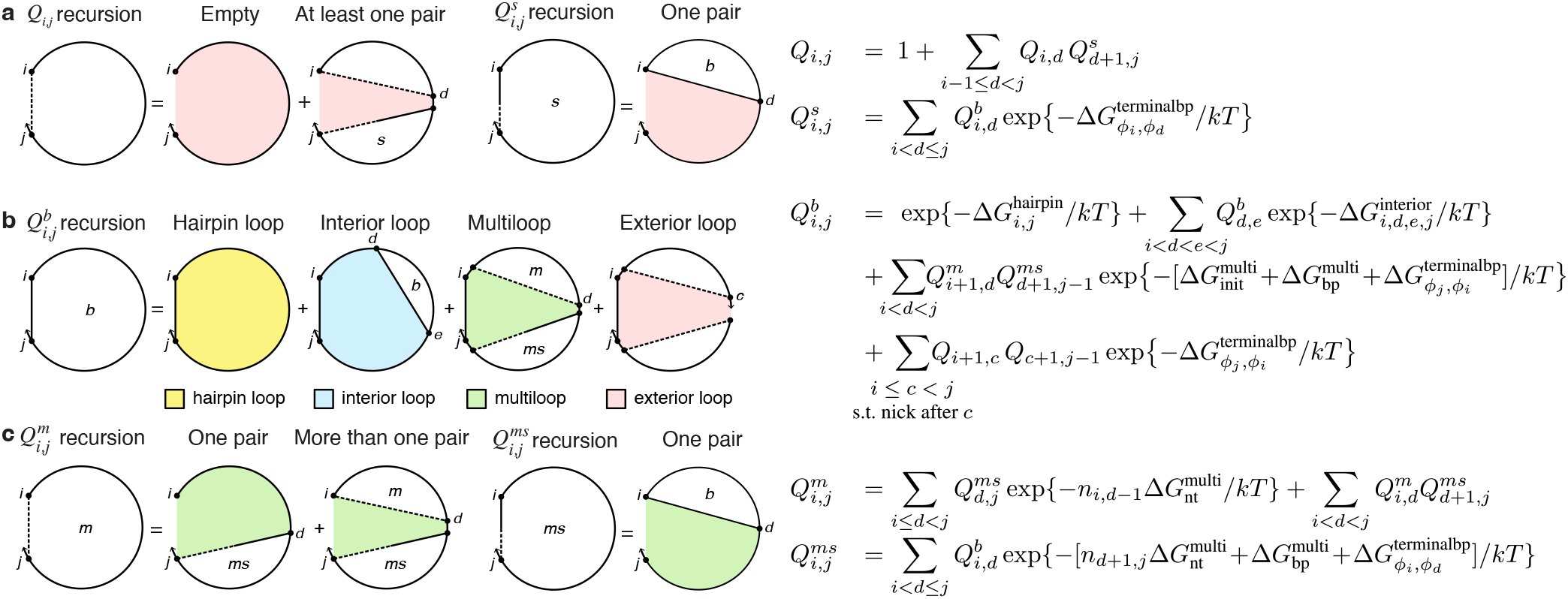
Partition function dynamic program recursion diagrams (left) and recursion equations (right) without coaxial and dangle stacking.^21^ A solid straight line indicates a base pair and a dashed line demarcates a region without implying that the connected bases are paired. A half-solid/half-dashed straight line indicates that the nucleotide connected on the solid side is base-paired to a nucleotide within the demarcated region. Shaded regions correspond to loop free energies that are explicitly incorporated at the current level of recursion (colors correspond to the loop types of Figure 1). (a) *Q*_*i,j*_ represents the partition function for subsequence [*i, j*]. There are two cases: either there are no base pairs (corresponding to the reference free energy 0 and partition function contribution 1) or there is a 3^′^-most base pair (between *d* + 1 and a nucleotide ≤ *j*). In the latter case, determination of the partition function contribution makes use of previously computed subsequence partition functions 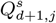 and *Q*_*i,d*_. 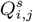 represents the partition function for subsequence [*i, j*] in an exterior loop where *i* is base-paired to a nucleotide ≤ *j*. By the distributive law, multiplication of these subsequence partition functions (each representing a sum over substructures) implicitly sums over all pairwise combinations of substructures. The independence of the loop contributions in the energy model implies that these products appropriately add the free energies in the exponents. (b) 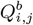 is the partition function for subsequence [*i, j*] with the restriction that bases *i* and *j* are paired. There are four cases: either there are no additional base pairs (corresponding to a hairpin loop), there is exactly one additional base pair *d*·*e* (corresponding to an interior loop), there is more than one additional base pair (corresponding to a multiloop) with 3^′^-most pair (between *d* + 1 and a nucleotide *< j*) and at least one additional pair specified in a previously computed subsequence partition function 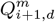, or there is an exterior loop containing a nick after nucleotide *c*. (c) 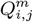 is the partition function for subsequence [*i, j*] with the restrictions that the subsequence is inside a multiloop and contains at least one base pair. There are two cases: either there is exactly one additional base pair defining the multiloop (between *d* and a nucleotide *j*), or there is more than one additional base pair defining the multiloop (with 3^′^-most pair between *d* + 1 and a nucleotide≤ *j*). In either case, the 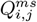 recursion represents the partition function for subsequence [*i, j*] in a multiloop with *i* base-paired to a nucleotide≤ *j. n*_*i,j*_ *j i* + 1 denotes the number of nucleotides between *i* and *j* inclusive. The time complexity of recursions (a)-(c) is *O*(*N* ^3^) (indices *i, d, j* or *i, c, j* on the right hand side) and the space complexity is *O*(*N* ^2^) (indices *i, j* on the left hand side); interior loop contributions have four indices (*i, d, e, j*) on the right hand side but are calculated with *O*(*N* ^3^) time complexity as described in Section S6.4.

### Mixed-material recursion overview

Recursions for the single-material algorithm use free energy parameters for only one material (either RNA or DNA) for all loops independent of the material of each nucleotide in a given loop. Our mixed-material algorithm switches parameter sets based on the material composition of each loop, with mixed-material loops using RNA/DNA parameters, RNA loops using RNA parameters, and DNA loops using DNA parameters (Figure 2). As a result, a given single-material loop is treated identically by single- and mixed-material algorithms, and in the special case where a system contains only one material, mixed-material algorithms return the same results as single-material algorithms.

Parameter switching is trivial for loop types whose contributions to the partition function are calculated within a single recursion level since the material for each nucleotide can be checked and the appropriate free energy parameters applied to the loop as a whole. However, the situation is more challenging for loop types whose contributions to the partition function are calculated across multiple recursion levels, as these loops are never considered in their entirety, necessitating the use of both single- and mixed-material recursions to keep track of the material properties of each loop, employing the appropriate single- or mixed-material free energy parameters on the fly. Here, we outline the mixed-material recursion concepts for each loop type. Full details are provided in Section S6, including recursions with coaxial and dangle stacking.

### Hairpin loop treatment

The partition function contribution of a hairpin loop is incorporated within a single recursion level. Thus, the material of the entire loop can be checked and either single- or mixed-material parameters used as appropriate.

### Interior loop treatment

Consider an interior loop closed by base pairs *i*·*j* and *d*·*e* with *i < d < e < j* and let *n*_1_ ≡*d− i−* 1 and *n*_2_≡ *j −e−* 1 denote the number of unpaired bases for the two sides of the interior loop. Interior loops with *n*_1_ *<* 4 or *n*_2_ *<* 4 are explicitly enumerated within a single recursion level, so the material can be checked for the entire loop and single- or mixed-material parameters used as appropriate.

For interior loops with *n*_1_ ≥ 4 and *n*_2_ ≥ 4, the free energy is given by:

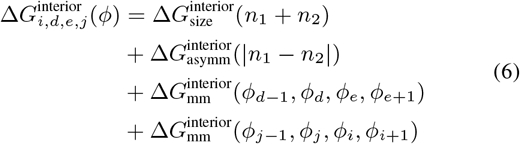

corresponding to a size term (dependent on the sum of the lengths of the sides, *n*_1_ + *n*_2_), an asymmetry term (dependent on the difference in the length of the sides, |*n*_1_−*n*_2_|), and sequence-dependent mismatch terms (reflecting stacking of bases *i* + 1 and *j*−1 on closing base pair *i*·*j* and bases *d*−1 and *e*+1 on closing base pair *d*·*e*). These loops are termed *extensible interior loops* because the free energy of smaller loops can be used to calculate the free energy of larger loops in order to reduce the time complexity of the algorithm. Note that extending both the 5^′^ and 3^′^ ends of the loop by 1 nucleotide so that the loop is closed by base pair *i*−1·*j* + 1 instead of *i*·*j* does not change the asymmetry and 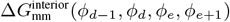 mismatch terms. Hence, the free energy of extensible interior loops of size *l ≡ n*_1_ + *n*_2_ can be calculated from extensible loops of size *l*−2 by swapping out the size parameter 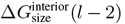 for 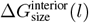 and replacing the mismatch term 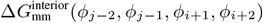 with 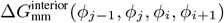. For example, in calculating the partition function this corresponds to:

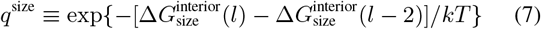

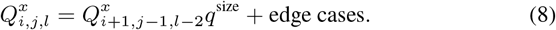

Here 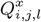 is the partition function contribution for extensible interior loops of size *l* bounded by indices *i* and *j*. This operation in effect replaces 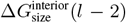 with 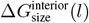 for each loop in 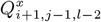, obtaining the partition function contribution for loops extended by one nucleotide on each side. Note that *Q*^*x*^ does not include the 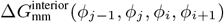 mismatch term, which is incorporated at a higher recursion level for each closing base pair *I*·*j*, making it unnecessary to swap out this mismatch contribution as the loop is extended. *Q*^*x*^ also includes edge cases corresponding to contributions from extensible loops with *n*_1_ = 4 or *n*_2_ = 4 that cannot be extended from smaller extensible loops. The process of loop extension reduces the time complexity of calculating interior loop contributions from *O*(*N* ^4^) to *O*(*N* ^3^), using three loop indices (*i, j, l*) instead of four (*i, j, d, e*).

To accommodate mixed materials, the *Q*^*x*^ recursion must be split into separate recursions that track extensible interior loops that are either single- or mixed-material, using either single- or mixed-material values of the 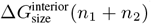 and 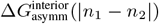 parameters according to the overall material properties of each loop. Let *µ*_*i*_ denote the material of nucleotide *i* (e.g., either RNA or DNA) and let the Kronecker delta function

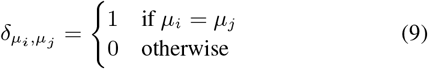

indicate whether *i* and *j* are the same or different materials.

To extend single-material loops, the two nucleotides that are added (*i* and *j*) must both match the material of the other nucleotides in the loop, which can be verified by comparing to *i* + 1 (Figure 5a):

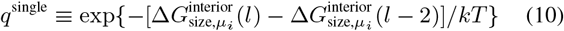

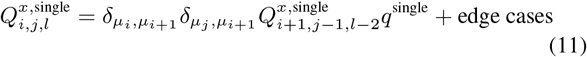

**Figure 5:**
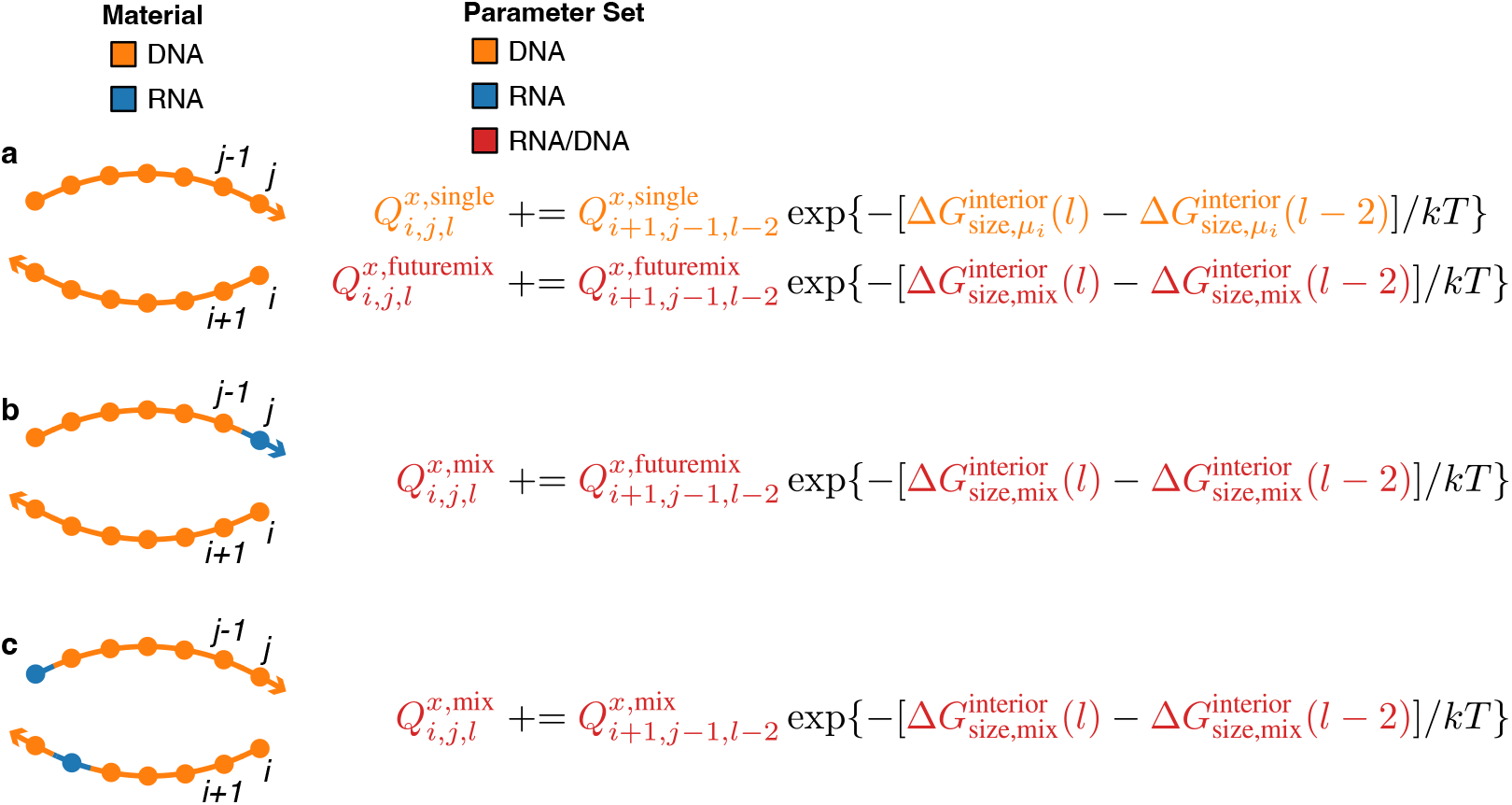
Treatment of extensible interior loops in the mixed-material algorithm. (a) Extension to single-material loop. If the material of the extension (nucleotides *i* and *j*) matches the material of the previous nucleotides (checked by comparing to nucleotide *i* + 1), single-material loop contributions cached in 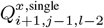 are extended as single-material contributions in 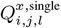. In preparation for a potential mixed-material extension to previously single-material loops, the recursion 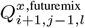 caches contributions of the same loops using the mixed-material parameter set instead of the single-material parameter set. (b) Mixed-material extension to previously single-material loops. If there is a mismatch between the material of the extension (nucleotides *i* and *j*) and the material of the previous nucleotides (checked by comparing to nucleotide *i* + 1), mixed-material contributions previously cached in 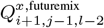 are extended as mixed-material contributions in 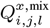. (c) Extension to mixed-material loops. 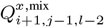 is always extended in 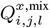, since mixed-material loops remain mixed-material no matter which materials are introduced by the extension.

If the material of *i* or *j* does not match *i* + 1 and *j*−1 (Figure 5b), then single material loops of size *l*−2 cannot be extended as single-material loops to size *l* and must instead be extended as mixed-material loops. This requires a related recursion, *Q*^*x*,futuremix^, that tracks the same single-material loops as *Q*^*x*,single^, but uses the mixed-material parameter set:

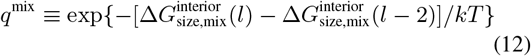

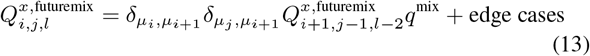

so that single-material loops can be extended as mixed-material loops if *i* or *j* introduce a new material:

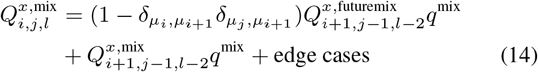

Here, the *Q*^*x*,mix^ recursion tracks mixed-material loops, which are always extended as mixed-material since they remain mixed no matter what material *i* and *j* are (Figure 5c). See Section S6.4 for full details on interior loop recursions for the mixed-material algorithm.

### Multiloop treatment

A multiloop is closed by 3 or more terminal base pairs and may be defined by the series of bounding subsequences [*ϕ*], the ends of which form the terminal base pairs. The free energy for a multiloop (without coaxial and dangle stacking) is modeled as:

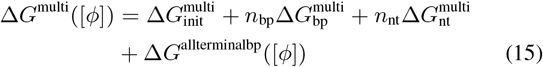

where 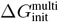denotes the penalty for formation of a multiloop, 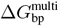 denotes the sequence-independent penalty for a terminal base pair in a multiloop, *n*_bp_ denotes the number of terminal base pairs, 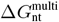 denotes the penalty per unpaired nucleotide in a multiloop, and *n*_nt_ denotes the number of unpaired nucleotides. Δ*G*^allterminalbp^([*ϕ*]) is the sum of the sequence-dependent penalty 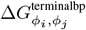for each terminal base pair *i*·*j* closing the multiloop. We define 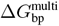 and 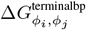 based on the material(s) of a given terminal base pair whether the overall loop is single- or mixed-material. For example, these parameters are the same for an RNA terminal base pair whether the multiloop is all-RNA or a mixture of RNA and DNA. Likewise, we define 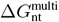 based on the material of each unpaired nucleotide within the multiloop whether the overall loop is single- or mixed-material. By contrast, 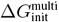applies to the multiloop as a whole and has different values for single- and mixed-material multiloops. For efficiency reasons, multiloop partition function contributions are calculated using multiple levels of recursions without ever knowing the full multiloop structure at any single recursion level. As a result, it is necessary to define single- and mixed-material recursions so that once the partition function contributions for a given multiloop have been calculated through the recursive process, the correct value of 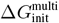can be incorporated depending on whether the multiloop turns out to be single- or mixed-material.

To illustrate this concept, consider one of the contributions to the *Q*^*m*^ recursion of Figure 4c,

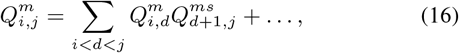

corresponding to the partition function contribution for subsequence [*i, j*] containing more than one base pair inside a multiloop (with 3^′^-most pair between *d* + 1 and a nucleotide ≤ *j*). Figure 6a illustrates different single-material multiloops and the single-material recursions used to generate them for an RNA sequence.

**Figure 6:**
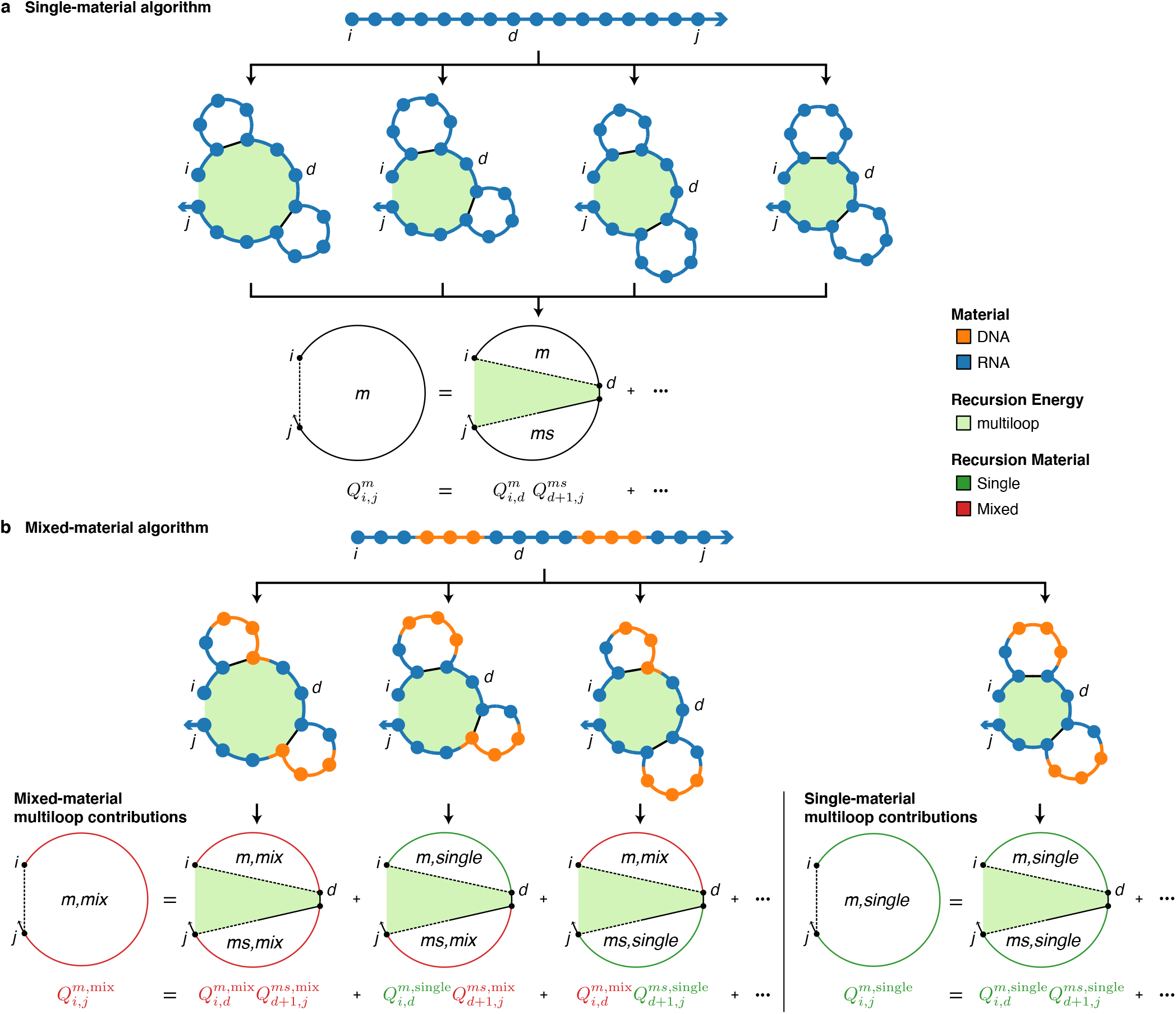
Comparison of multiloop treatment for single- and mixed-material algorithms. (a) Single-material algorithm. (b) Mixed-material algorithm. For each algorithm, recursions are depicted with recursion diagrams (above) and recursion equations (below). In recursion diagrams, a dashed line demarcates a region without implying that the connected bases are paired; a half-solid/half-dashed straight line indicates that the nucleotide connected on the solid side is base-paired to a nucleotide within the demarcated region; shaded regions correspond to loop free energies that are explicitly incorporated at the current level of recursion. Examples of partial multiloops (each corresponding to a shaded region closed by two terminal base pairs in subsequence [*i, j*]) are depicted indicating the recursion that incorporates their contributions to the partition function.

If we split *Q*^*m*^ and *Q*^*ms*^ into single- and mixed-material recursions, we have the following:

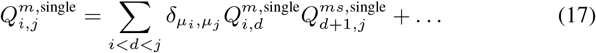

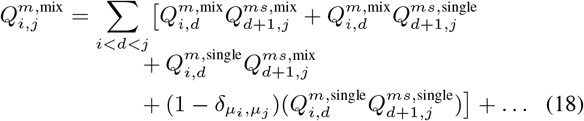

Unlike the single-material algorithm, which contains the 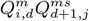 contribution (16), the mixed-material algorithm treats four combinations of these terms that are assigned to a higher level 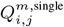 recursion (17) or 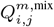 recursion (18) depending on whether the combination represents an overall single- or mixed-material multiloop contribution. Any combination involving at least one mixed-material recursion element 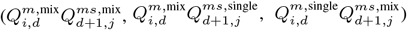 represents an overall mixed-material contribution that is included in 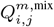. The remaining combination, 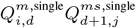, represents a single-material contribution if the materials for the two single-material subsequences [*i, d*] and [*d* + 1, *j*] match (added to (17) using the conditional 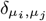), and a mixed-material contribution if they do not (added to (18) using the conditional 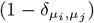). Figure 6b illustrates different mixed-material and single-material multiloops and the recursions used to generate them for a chimeric RNA/DNA sequence.

For the single-material algorithm, the *Q*^*b*^ recursion (Figure 4b) recursively calls *Q*^*m*^ and *Q*^*ms*^ to complete the calculation of the partition function contributions for each multiloop, incorporating the final terminal base pair, *j·i*, to close the loop. For the mixed- the *Q*^*m*^ and *Q*^*ms*^ material algorithm, 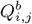 is evaluated by comparing the materials of the 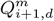 and 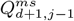 recursions as well as the material of the terminal base pair *j·i* and using either a single- or mixed-material value for 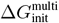, the penalty for formation of a multiloop, depending on whether the overall multiloop turns out to be single- or mixed-material:

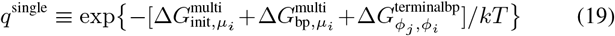

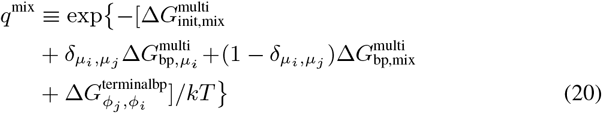

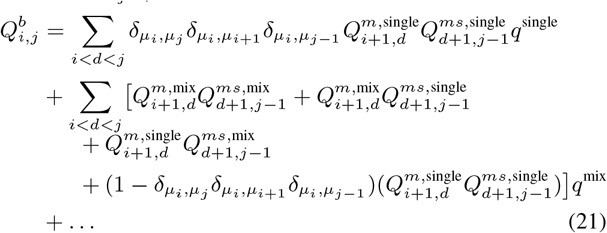

See Sections S6.3 and S6.5 for full details on multiloop recursions without/with coaxial and dangle stacking.

### Exterior loop treatment

An exterior loop contains a nick between strands, is closed by zero or more terminal base pairs, and may be defined by the series of bounding subsequences [*ϕ*], two ends of which border the nick and the other ends of which form the terminal base pairs. An unpaired strand is an exterior loop with a free energy of zero, corresponding to the reference state.^21^ The free energy for an exterior loop without coaxial and dangle stacking is modeled as:

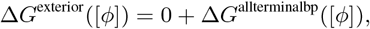

where Δ*G*^allterminalbp^([*ϕ*]) is a sum of the sequence-dependent penalty 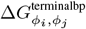 for each terminal base pair *i·j* closing the exterior loop. As in the multiloop setting, we define 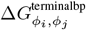 based on the material( s) of a given terminal base pair whether the overall exterior loop is single- or mixed-material. For example, the parameter is the same for an RNA terminal base pair whether the exterior loop is all-RNA or a mixture of RNA and DNA. As a result, when a terminal base pair is encountered in an exterior loop context within a recursion (e.g., the terminal base pair *j·i* in the exterior loop recursion of Figure 4b), the material of the terminal base pair is checked and either a single- or mixed-material parameter value is used as appropriate.

## Results and Validation

### Validation with empirical RNA/DNA melting temperature data from the literature

The new mixed-material RNA/DNA parameter set incorporates available stack loop and internal mismatch free energies derived in the literature from duplex melt experiments.^13, 15, 20^ For each duplex used in the parameterization experiments, we used the new mixed-material algorithms to calculate the melting temperature (*T*_*m*_) for comparison to the experimental melting temperature. Figure 7 displays the computed and empirical *T*_*m*_ for the duplexes used in four studies: RNA/DNA hybrid duplexes with no mismatches^13^ (panel a), RNA/DNA chimeric duplexes with no mismatches^15^ (panel b), RNA/DNA hybrid duplexes with or without a mismatch^20^ (panel c), RNA/DNA hybrid duplexes with or without a single hybrid or chimeric mismatch (panel d). In each plot, ideal data would fall on the diagonal. Using mixed-material RNA/DNA simulations, the mean absolute error (MAE) in *T*_*m*_ was *{*2.1, 0.58, 2.1, 2.5*}* ^*°*^C for these four data sets. In the absence of mixed-material parameter sets and algorithms, current standard practice is to simulate RNA/DNA systems as either all-RNA or all-DNA. Using all-DNA simulations for these four data sets, the MAE increased to *{*4.9, 5.2, 4.3, 4.7*}* ^*°*^C with the *T*_*m*_ typically over-estimated for two data sets and typically under-estimated for two others. Using all-RNA simulations for these four data sets, the MAE increased further to *{*15, 12, 18, 16*}* ^*°*^C and the *T*_*m*_ was typically overestimated for all four data sets. These results are consistent with previous observations that mixed RNA/DNA base-pairing is energetically more similar to DNA-only base-pairing than to RNA-only base-pairing.^13^ The chimeric duplexes of Figure 7b have no mismatches and either of two duplex sequences, but with different RNA/DNA compositions; mixed-material simulations capture the empirical trend in *T*_*m*_, while single-material RNA-only or DNA-only simulations cannot, predicting the same *T*_*m*_ for all duplexes with the same sequence (depicted as horizontal lines).

**Figure 7:**
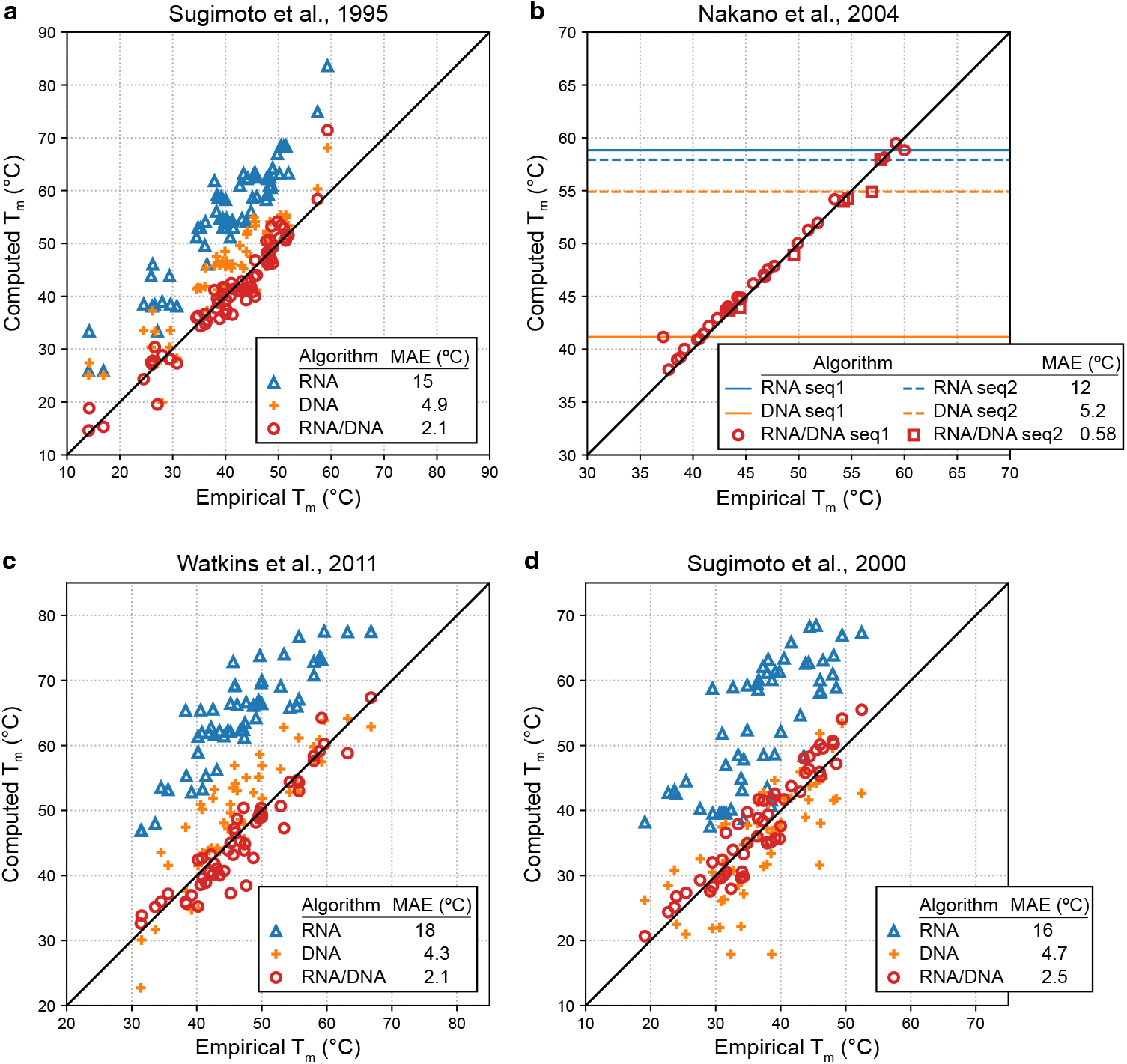
Improved prediction of duplex melting temperature using a mixed-material RNA/DNA algorithm compared to single-material RNA-only or DNA-only algorithms. Computed vs empirical melting temperatures (*T*_*m*_) for four different experimental studies. Mean absolute error (MAE). (a) Sugimoto et al, 1995.^13^ 67 RNA/DNA hybrid duplexes with no mismatches; one duplex was omitted that did not melt between 0 ^*°*^C and 100 ^*°*^C using RNA parameters. (b) Nakano et al., 2004.^15^ 37 chimeric duplexes with no mismatches (30 duplexes with the same non-self-complementary sequence (termed seq1) with varying RNA and DNA composition e.g., rGACUAdGGT/rACCUAGUC, dGACTrAGGU/dACCTAGTC, dGACTrAGGU/rACCUdAGTC; 7 duplexes with the same self-complementary sequence (termed seq2) with varying RNA and DNA composition e.g., rCGCGCdG/rCGCGCdG, rCGCdGCG/rCGCdGCG; the single-material predictions are the same for all duplexes with the same sequence (depicted as horizontal lines). (c) Watkins et al., 2011.^20^ 54 RNA/DNA hybrid duplexes: 43 contain a single internal rA *×*dA, rC *×*dC, rG*×* dG, or rU*×* dT mismatch, 11 contain no mismatch; duplexes that appear in Sugimoto et al., 2000^29^ were omitted. (d) Sugimoto et al., 2000.^29^ 57 RNA/DNA hybrid duplexes: 39 contain a single mismatch (rA*×* dA, rA *×*dC, rA *×*dG, rC *×*dA, rC *×*dT, rG*×* dA, rG *×*dG, rU *×*dC, rU *×*dG, or rU *×*dT), 9 contain a wobble pair (rG *·* dT), 4 contain a chimeric mismatch (dT *×*dG), 5 contain no mismatch; eight sequences containing dU were omitted since there are no DNA parameters for U. Each strand: 50 µM. Salt conditions:1.0 M Na^+^.

### Validation with empirical RNA/2′OMe-RNA melting temperature data from the literature

The new mixed-material algorithms can also be used with the new mixed-material RNA/2^′^OMe-RNA parameter set to model systems that contain a mixture of RNA and 2^′^OMe-RNA nucleotides. Figure 8 displays the computed and empirical *T*_*m*_ for RNA/2^′^OMe-RNA hybrid duplexes without mismatches that were used to parameterize the model. Mixed-material RNA/2^′^OMe-RNA simulations produce an MAE of 0.97 ^*°*^C compared to 5.1 ^*°*^C for RNA-only simulations, which tend to underestimate the *T*_*m*_.

**Figure 8:**
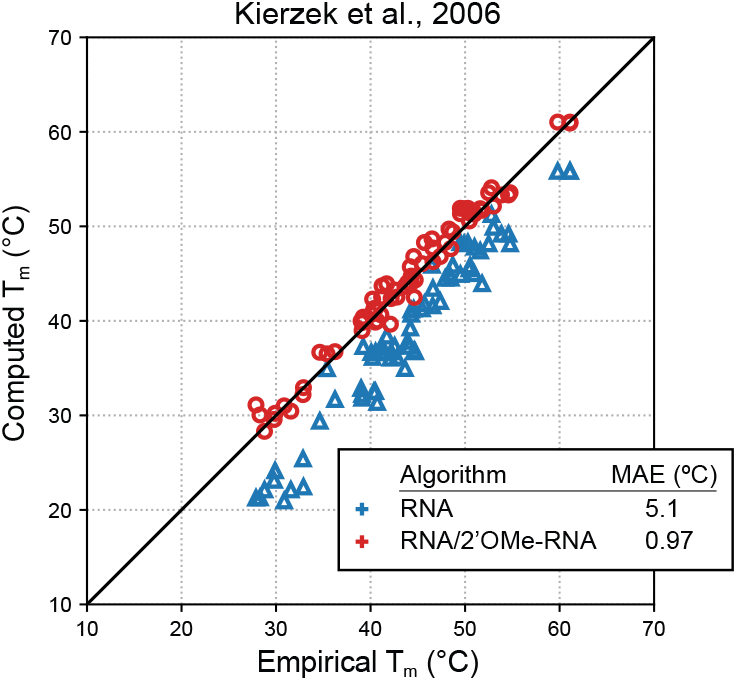
Improved prediction of duplex melting temperature using a mixed-material RNA/2^′^OMe-RNA algorithm compared to a single-material RNA-only algorithm. Computed vs empirical melting temperature (*T*_*m*_) for the experimental study of Kierzek et al., 2006.^17^ Mean absolute error (MAE). 68 RNA/2^′^OMe-RNA hybrid duplexes without mismatches.^17^ Each strand: 50 µM. Salt conditions: 0.12 M Na^+^.

### Validation with melt experiments on different duplex material compositions

As a further test of the utility of the mixed-material model and algorithms, we synthesized duplexes with each of four different strand compositions (DNA:DNA, DNA:RNA, RNA:DNA, and RNA:RNA) and performed melt experiments to enable comparison of computed and experimental melt curves for a given duplex sequence with these different material compositions. Figure 9 displays the computed and experimental melt curves and melting temperatures for three duplex sequences. The mixed-material simulations correctly predict the stability order between the different material compositions. This is particularly significant because while RNA:RNA duplexes are generally more stable than DNA:DNA duplexes, whether DNA:DNA is more or less stable than DNA:RNA and/or RNA:DNA hybrid duplexes is sequence-dependent.^13^ Indeed, the order of increasing stability for Duplex 1 is DNA:RNA *<* RNA:DNA *<* DNA:DNA *<* RNA:RNA, with the all-DNA duplex more stable than either of the hybrid duplexes, but for Duplexes 2 and 3, the order of increasing stability switches to be DNA:RNA *<* DNA:DNA *<* RNA:DNA *<* RNA:RNA, with the all-DNA duplex of intermediate stability between the two hybrid duplexes. The MAE in *T*_*m*_ across the three duplex sequences was 1.7°C for DNA:DNA duplexes, 1.7°C for DNA:RNA duplexes, 2.6°C for RNA:DNA duplexes, and 1.1°C for RNA:RNA duplexes.

### Relative efficiency of single- and mixed-material dynamic programs

Both the single- and mixed-material algorithms have time complexity *O*(*N* ^3^) using dynamic programming recursions that operate on a maximum of three nucleotide indices in each recursion. However, the mixed-material algorithms include new conditional statements to check the material composition of each nucleotide, as well as duplicate recursions to appropriately handle single- and mixed-material loop contributions within the dynamic program. Figure 10 compares the speed of single- and mixed-material partition function algorithms for 3-stranded complexes. Across a wide range of complex sizes, the mixed-material algorithm is *≈*2–3.5 *×* slower than the single-material algorithm. Hence, the significant accuracy benefits achieved by modeling mixed-material systems at nucleotide resolution come at the cost of only a modest increase in simulation time.

**Figure 9:**
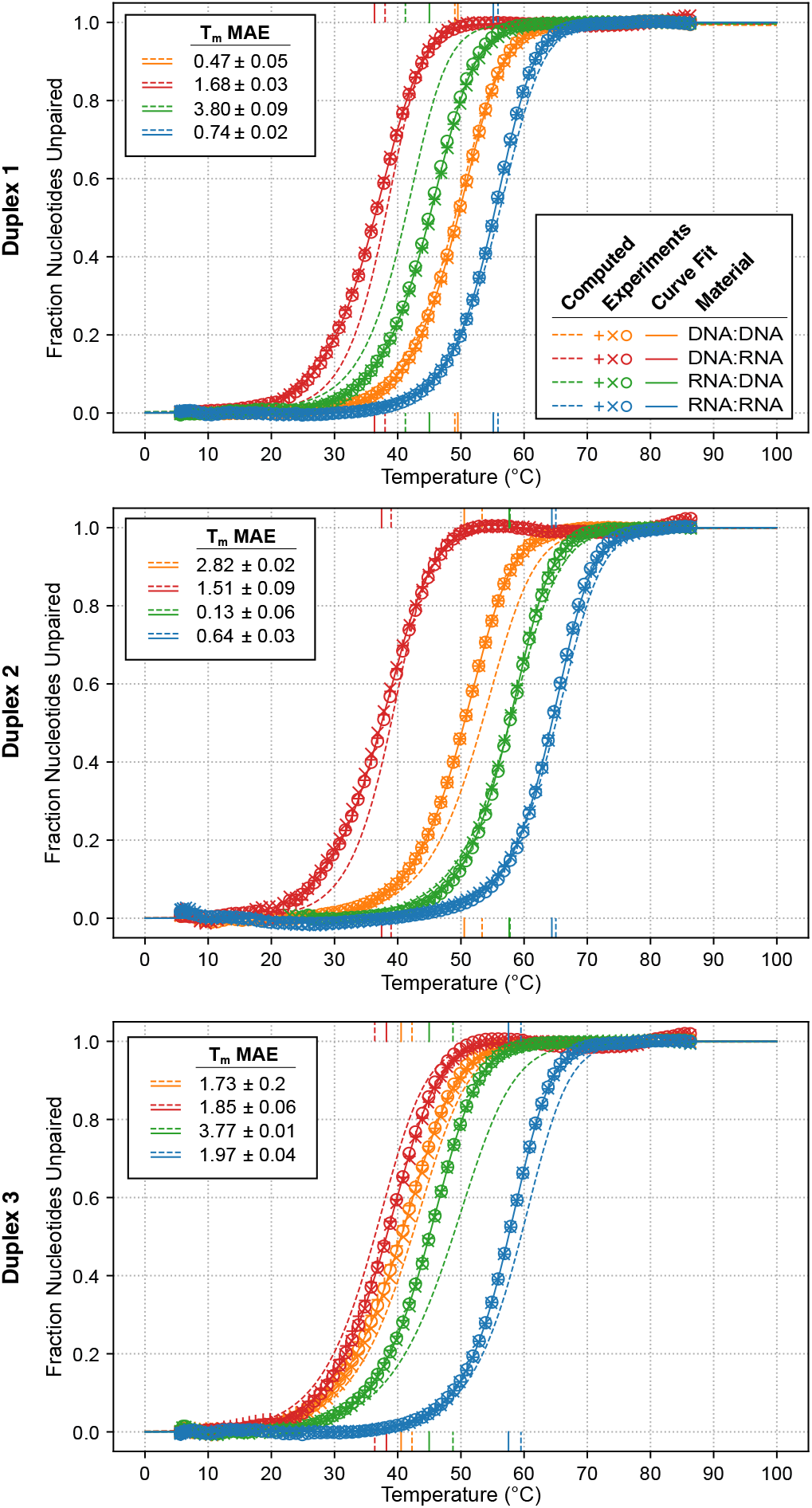
Comparison of computed vs experimental melt curves and melting temperatures for three duplex sequences each synthesized with four different strand compositions (DNA:DNA, DNA:RNA, RNA:DNA, and RNA:RNA). Computed melt: fraction of nucleotides unpaired. Experimental melt: fraction of nucleotides unpaired based on curve fit with 2-state model. Symbols denote *N* = 3 replicate experiments; each experimental curve fit represents the mean of the curve fits for the replicates. Computed and experimental melting temperatures (*T*_*m*_) depicted as vertical lines. Mean absolute error (MAE) *±* SEM for *N* = 3 replicate experiments. Each strand: 5 µM. Salt conditions: 1.0 M Na^+^.

**Figure 10:**
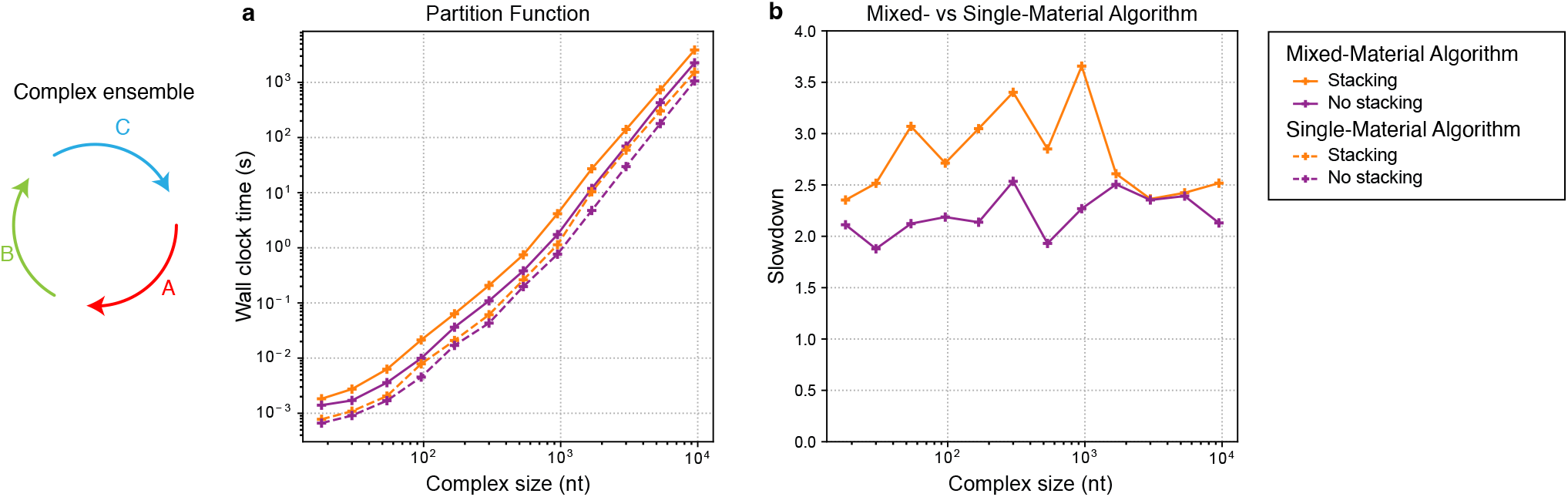
Relative speed of single- and mixed-material partition function algorithms. Calculation of the partition function with or without coaxial and dangle stacking for a complex of 3 strands, each with a different random sequence of uniform length. For the mixed-material tests, each strand contained a random mix of RNA and DNA nucleotides. For single-material tests, the same strand sequences were used with only DNA nucleotides. (a) Computational cost. Mean wall clock time over 75 sets of random sequences per complex size *<* 300 nt, and 5 sets of random sequences per complex size *≥* 300 nt. (b) Computational slowdown (ratio of mean wall clock times). Conditions: 37 ^*°*^C, 1.0 M Na^+^. See Section S7.1 for additional timing data.

## Conclusions

We formulated free energy parameter sets for RNA/DNA and RNA/2^′^OMe-RNA mixed-material systems to enable analysis of complex and test tube ensembles with the material composition specified at nucleotide resolution. We developed new single- and mixed-material dynamic programming recursions that apply the appropriate single- or mixed-material free energy parameters to each loop within a secondary structure depending on the material properties of a given loop. The mixed-material algorithm exactly reproduces the results of the single-material algorithm when applied to a single-material system. In the context of the unified dynamic programming framework underlying NUPACK,^10^ the new mixed-material recursions enable calculation of diverse physical quantities for mixed-material systems (complex partition function, equilibrium complex concentrations, equilibrium base-pairing probabilities, minimum free energy secondary structure(s), and Boltzmann-sampled secondary structures; see Section S2.9 for details). The mixed-material algorithms retain the *O*(*N* ^3^) time complexity of the single-material algorithms with a *≈*2–3.5*×* increase in run time.

We validated the mixed-material models and algorithms by predicting duplex melting temperatures from literature experiments for both RNA/DNA and RNA/2^′^OMe-RNA data sets. For mixed-material systems, the mixed-material models and algorithms are significantly more accurate in predicting melting temperatures than single-material models and algorithms. We further validated the mixed-material simulations by comparing to new duplex melt curves for three hybrid duplex sequences synthesized with four combinations of strand compositions (DNA:DNA, DNA:RNA, RNA:DNA, and RNA:RNA). Mixed-material simulations correctly predicted the sequence-dependent order of duplex stability for the different material compositions.

The new mixed-material parameter sets incorporate the limited supply of empirical mixed-material loop free energies that have been regressed in the literature, filling in missing mixed-material values with blended single-material values. These new mixed-material algorithms significantly increase the motivation to perform additional mixed-material empirical parameterization studies as updates to the parameters files can be directly leveraged in future mixed-material simulations of complex and test tube ensembles, including for new combinations of materials.

## Materials and Methods

### Materials

Sequences for the three duplexes used for UV melt experiments are shown in Table 1. The sequences were selected for experimental study based on predictions using the mixed-material algorithm that they would fully melt within the range 0 °C to 100 °C and have different *T*_*m*_ values for each of the four material combinations. Each strand was synthesized and HPLC-purified as an all-RNA oligo and as an all-DNA oligo by Integrated DNA Technologies.

**Table 1:**
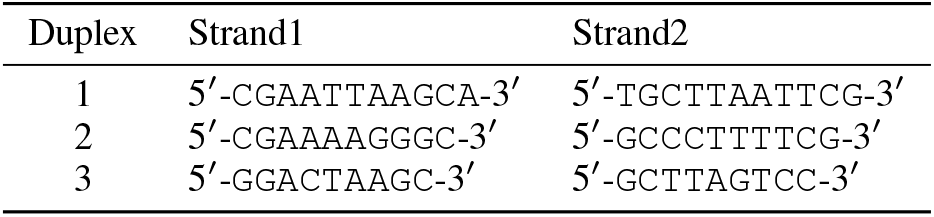
Three duplex sequences each synthesized with four different strand compositions (DNA:DNA, DNA:RNA, RNA:DNA, RNA:RNA). `U` replaces `T` for RNA strands.

### Duplex melt experiments

Each strand was resuspended in a solution of 1 M NaCl, 0.5 mM Tris, 5 µM EDTA. Concentrations were determined by measuring absorbance at 260 nm. For each duplex, two complementary strands were mixed at a 1:1 ratio with each strand at 5 µM. Melts were performed using the Agilent Cary 3500 Multicell Peltier UV-Vis Double Beam Spectrophotometer. Absorbance vs temperature curves were collected at 260 nm from 5 to 90 °C at a heating rate of 2 °C/min. Below 25 °C, N_2_ gas was streamed through the instrument to prevent condensation on the cuvettes. *N* = 3 replicate melt experiments were performed for each material composition (DNA:DNA, DNA:RNA, RNA:DNA, and RNA:RNA) for each of the three duplex sequences in Table 1.

### Curve fitting for duplex melt experiments

For each replicate melt experiment, UV absorbance data were fit with a two-state model where each strand is assumed to be either in a single-stranded (ss) unpaired state (strands A and B) with no base pairs formed or in a double-stranded (ds) duplex state (duplex AB) with all base pairs formed. Absorbance curve fits with sloping baselines for the ss and ds states were performed using the Levenberg–Marquardt (LM) algorithm, yielding the temperature-dependent fraction of strands in the single-stranded state, *f*_ss_(*T*) (see Section S2.8 for details).^44^ The empirical melting temperature, *T*_*m*_, is defined as the temperature at which *f*_ss_(*T*) = 0.5. Figure 9 displays the mean *f*_ss_(*T*) curve fit and mean *T*_*m*_ over *N* = 3 replicate experiments. Additionally, replicate absorbance measurements are displayed as equivalent *f*_ss_(*T*) values calculated as a fraction of the absorbance difference between the regressed single-stranded and double-stranded sloping baselines at a given temperature. Note that for a two-state model applied to strands that form a fully complementary duplex (as is the case for Figure 9), *f*_ss_(*T*) is equivalent to the fraction of nucleotides that are unpaired in solution.

### Computational melt studies

Computational melt studies for Figures 7–9 were performed using mixed-material temperature-dependent free energy parameter sets (`rna_dna06` for RNA/DNA and `rna_merna06` for RNA/2^*/*^OMe-RNA) using the `nostacking` complex ensemble (no coaxial or dangle stacking^10, 27^) for a test tube ensemble that includes all monomers and dimers formed from strands A and B for a given duplex (i.e., A, B, AA, BB, AB). Single-material calculations were performed using single-material parameter sets `rna06` for RNA and `dna04.2` for DNA. Strand concentrations and salt conditions were set as follows to match experimental conditions: each strand at 50 µM in 1.0 M Na^+^ for Figure 7; each strand at 50 µM in 0.12 M Na^+^ for Figure 8; each strand at 5 µM in 1.0 M Na^+^ for Figure 9. For 1 °C increments from 5 to 90 °C we calculated the partition function and equilibrium base-pairing probability matrix for each complex. The partition functions for the complexes in the test tube were then used to calculate the equilibrium complex concentration for each complex species. The equilibrium complex concentrations and equilibrium base-pairing probability matrices were then used to calculate the fraction of unpaired nucleotides, *f*_u_(*T*) for the test tube ensemble,^27^ which is displayed as a computational melt curve in Figure 9. For fully complementary duplexes, *f*_u_(*T*) falls in the interval [0, 1] and computational melting temperature, *T*_*m*_, is defined as the temperature at which *f*_u_(*T*) = 0.5. For duplexes with an internal mismatch (1*×* 1 interior loop), *f*_u_(*T*) falls in the interval [1*/n*, 1], where *n* is the number of nucleotides in each strand, and the computational *T*_*m*_ is defined as the temperature at which *f*_u_(*T*) = (*n* + 1)*/*(2*n*), corresponding to the midpoint of the interval. For the timing comparisons of Figure 10, jobs were run in serial on dual-processor 8-core 3.30 GHz Intel E5-2667v2 Xeon nodes with 64 GB of memory.

Literature free energy parameters were regressed from experimental studies using a two-state model that models each duplex as either fully associated or fully disassociated. On the other hand, the partition function algorithm operates on a complex ensemble containing all possible (unpseudoknotted connected) secondary structures, with or without coaxial and dangle stacking. If coaxial and dangle stacking states are included in the ensemble, each secondary structure incorporates Boltzmann-weighted contributions from a subensemble of stacking states.^10^ We found that including coaxial and dangle stacking states using literature parameters derived using a two-state model tended to overstabilize the duplex and overestimate the melting temperature compared to experimental results (see Section S7.2). This is not to say that the `stacking` ensemble is less accurate than the `nostacking` ensemble, but rather that it is less compatible with how the literature free energy parameters were regressed from experimental data. Efforts are underway to regress free energy parameters from scratch for use with the full complex ensemble including coaxial and dangle stacking states.

## RESOURCES

### NUPACK web app and Python module

Mixed-material jobs can be run interactively via the NUPACK web app (nupack.org) or scripted locally using the NUPACK Python module, subject to the NUPACK software license agreement. Please direct questions, comments, feature requests, and bug reports to.

## ASSOCIATED CONTENT

### Supporting Information

Strand association penalty for a complex, coaxial and dangle stacking subensembles within multiloops or exterior loops, complex ensembles with and without coaxial and/or dangle stacking, validation and implementation of salt corrections, temperature dependence of the free energy model, treatment of constant free energy terms for complex ensembles, distinguishability issues, curve fitting for duplex melt experiments, calculation of diverse physical quantities over complex and test tube ensembles using a unified dynamic programming framework, single-material free energy models for RNA, DNA, and 2^*/*^OMe-RNA, mixed-material free energy model for RNA/DNA and RNA/2^*/*^OMe-RNA, mixed-material recursions without or with coaxial and dangle stacking, additional studies, validation studies.

## AUTHOR INFORMATION

### Notes

The authors declare no competing financial interests.

## ACKNOWLEDGMENTS

This work was funded by the National Science Foundation (NSF-CHE-2317395 and NSF-OAC-1835414), by the Programmable Molecular Technology Center (PMTC) within the Beckman Institute at Caltech, by the AWS/IST Cloud Credit Program at Caltech. S.J.S was supported in part by National Institutes of Health NIGMS training grant GM008042. M.E.F. was supported in part by the U.S. Department of Energy, Office of Science, Office of Advanced Scientific Computing Research’s Applied Mathematics Competitive Portfolios program under Contract No. AC02-05CH11231.

## Footnotes

¶ NUPACK software for performing mixed-material calculations will be made available in conjunction with publication in a peer-reviewed journal.

