## Supplementary Information for "Models and Algorithms for Equilibrium Analysis of Mixed-Material Nucleic Acid Systems"

### List of Tables

### List of Algorithms

S1 Generalized dot product over multiple vectors of equal length . . . 31
S2 Enumerate valid positions for vectorization in an interstrand block . . . 32

### S1 Presentation of mixed-material models and algorithms

The supplementary information of Fornace et al.[1] provides a detailed description of a unified dynamic programming framework for calculating diverse physical quantities over single-material (RNA-only or DNA-only) complex and test tube ensembles. To facilitate comparison to the single-material case, here we follow the same presentation as closely as possible, modifying text, equations, algorithms, and figures as necessary to describe the models and algorithms for mixed-material ensembles.

### S2 Additional details

#### S2.1 Strand association penalty for a complex

Based on dimensional analysis, we define the complex concentrations $x_\Psi$ for a test tube containing the set of complexes $\Psi$ as mole fractions rather than molarities (see equation (S68)).[1,2] Therefore, we adjust the strand association penalty

$$\Delta G^{\text{assoc}} = \Delta G^{\text{assoc}}_{\text{pub}} - kT \log \frac{\rho_{\text{H}_2\text{O}}}{1\text{M}} \tag{S1}$$

where $\Delta G^{\text{assoc}}_{\text{pub}}$ is the published value for two strands associating[3] and $\rho_{\text{H}_2\text{O}}$ is the molarity of water (e.g., $\rho_{\text{H}_2\text{O}} = 55.14$ M at 37 °C).[2] The strand association penalty for a complex of $L$ strands (see equation (1)) is then

$$(L-1)\Delta G^{\text{assoc}}. \tag{S2}$$

To see how this adjustment arises, consider reaction

$$\text{A} + \text{B} \leftrightarrow \text{AB}. \tag{S3}$$

At equilibrium

$$\exp\left\{-\frac{\Delta G_{\text{AB}} - \Delta G_{\text{A}} - \Delta G_{\text{B}}}{kT}\right\} = \frac{x_{\text{AB}}}{x_{\text{A}} x_{\text{B}}} \tag{S4}$$

$$= \frac{[\text{AB}]/\rho_{\text{H}_2\text{O}}}{([\text{A}]/\rho_{\text{H}_2\text{O}})([\text{B}]/\rho_{\text{H}_2\text{O}})} \tag{S5}$$

$$= \frac{[\text{AB}]\rho_{\text{H}_2\text{O}}}{[\text{A}][\text{B}]} \tag{S6}$$

or equivalently

$$\Delta G_{\text{AB}} - \Delta G_{\text{A}} - \Delta G_{\text{B}} = -kT \log \frac{x_{\text{AB}}}{x_{\text{A}} x_{\text{B}}} \tag{S7}$$

$$= -kT \log \frac{[\text{AB}]\rho_{\text{H}_2\text{O}}}{[\text{A}][\text{B}]} \tag{S8}$$

Note that ratio of mole fractions is dimensionless as it must be in order to take the logarithm, or to be the result of exponentiation of a dimensionless quantity. A common practice is to write

$$\exp\left\{-\frac{\Delta G'_{\text{AB}} - \Delta G_{\text{A}} - \Delta G_{\text{B}}}{kT}\right\} = \frac{[\text{AB}]}{[\text{A}][\text{B}]} \tag{S9}$$

or equivalently

$$\Delta G'_{\text{AB}} - \Delta G_{\text{A}} - \Delta G_{\text{B}} = -kT \log \frac{[\text{AB}]}{[\text{A}][\text{B}]} \tag{S10}$$

which is formally problematic since the ratio of concentrations is dimensional so it is not proper to take the logarithm of this ratio, or to state that this ratio is the result of exponentiation of a dimensionless quantity. The situation can be rescued by non-dimensionalizing each concentration with a reference concentration

$$\exp\left\{-\frac{\Delta G'_{\text{AB}} - \Delta G_{\text{A}} - \Delta G_{\text{B}}}{kT}\right\} = \frac{[\text{AB}]/(1\text{ M})}{([\text{A}]/(1\text{ M}))([\text{B}]/(1\text{ M}))} \tag{S11}$$

or equivalently

$$\Delta G'_{\rm AB} - \Delta G_{\rm A} - \Delta G_{\rm B} = -kT \log \frac{[{\rm AB}](1\,{\rm M})}{[{\rm A}][{\rm B}]} \tag{S12}$$

so that the numerical values do not change but the quantities are now dimensionless to ensure compatibility of logarithms and exponentials. Subtracting equation (S12) from (S8) we then have

$$\Delta G_{\rm AB} - \Delta G'_{\rm AB} = -kT \log \frac{\rho_{\rm H_2O}}{1\,{\rm M}} \tag{S13}$$

Historically, the strand association penalty $\Delta G^{\rm assoc}_{\rm pub}$ that contributes to the free energy of dimer AB was parameterized using the formulation of (S9). We prefer the formulation (S4) rather than (S9) so we use the strand association penalty

$$\Delta G^{\rm assoc} = \Delta G^{\rm assoc}_{\rm pub} - kT \log \frac{\rho_{\rm H_2O}}{1\,{\rm M}} \tag{S14}$$

#### S2.2 Coaxial and dangle stacking subensembles within multiloops and exterior loops

Within a multiloop or exterior loop, a base pair can form one *coaxial stack* with an adjacent base pair,[4–6] or can form a *dangle stack* with at most two adjacent unpaired bases;[6–9] unpaired bases can either form no stack, or can form a dangle stack with at most one adjacent base pair. See Figure S1 for an illustration of the valid stacking states for a multiloop (panel a) or two exterior loops (panels b and c). For a given multiloop or exterior loop, the energetic contributions of all possible coaxial and dangle stacking states are enumerated so as to calculate the free energy:

$$\Delta G^{\rm stacking} = -kT \log \sum_{\omega \in {\rm loop}} \prod_{{\rm x} \in \omega} e^{-\Delta G_x / kT} \tag{S15}$$

where $\omega$ indexes the possible stacking states within the loop and $x$ indexes the individual stacks (coaxial or dangle) within a stacking state. The free energy of a multiloop or exterior loop is augmented by the corresponding $\Delta G^{\rm stacking}$ bonus. Hence, the stacking states within a given multiloop or exterior loop are treated as a structural subensemble that contributes in a Boltzmann-weighted fashion to the free energy model for the loop. Let $s'' \in s$ denote a stacking state of the paired and unpaired bases in secondary structure $s$. We may equivalently define the free energy of secondary structure $s$ in terms of the free energies for all stacking states $s'' \in s$:

$$\overline{\Delta G}(\phi, s) = -kT \log \sum_{s'' \in s} e^{-\overline{\Delta G}(\phi, s'')/kT} \tag{S16}$$

Let $\overline{\Gamma}''(\phi)$ denote the ensemble of stacking states corresponding to the complex ensemble of secondary structures $\overline{\Gamma}(\phi)$.

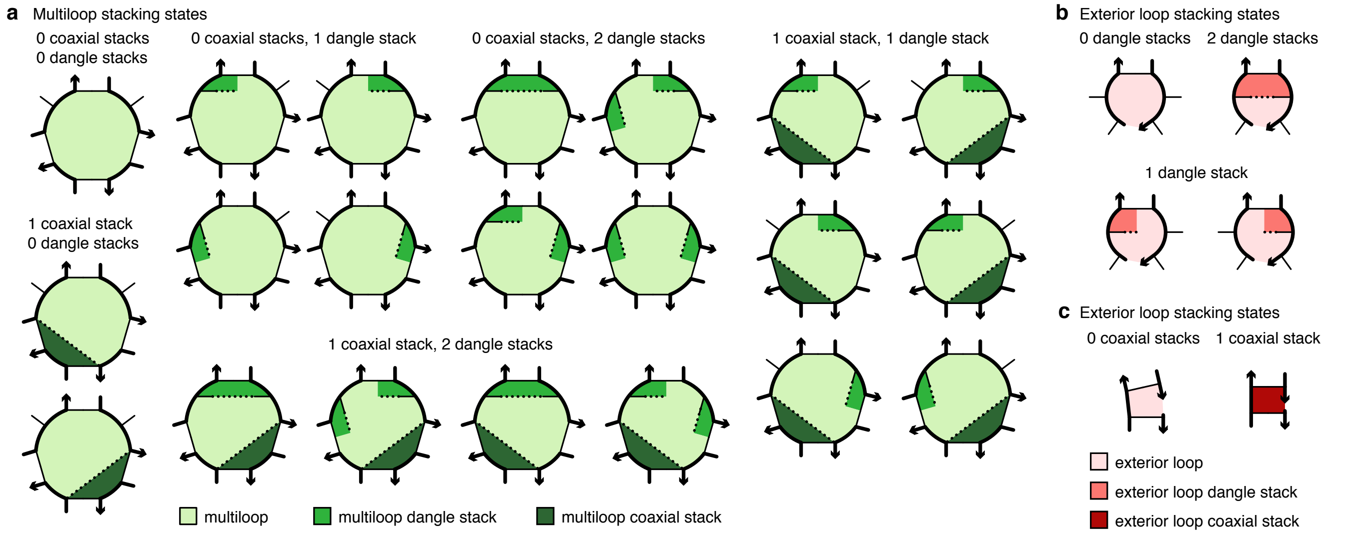

Figure S1: **Coaxial and dangle stacking states for multiloops and exterior loops.** (a) Stacking subensemble for the multiloop of Figure 1a. (b,c) Stacking subensembes for two exterior loops from Figure 1a. Used with permission from Fornace et al., *ACS Synth Biol*, 9, 2665-2678, 2020. Copyright 2020 American Chemical Society.

#### S2.3 Complex ensembles with or without coaxial and/or dangle stacking

NUPACK 4 algorithms perform calculations on the following complex ensembles:[1,10]

- `stacking`: complex ensemble with coaxial and dangle stacking in multiloops and exterior loops (ensemble $\overline{\Gamma}^{\shortparallel}(\phi)$).
- `dangle-stacking`: complex ensemble with dangle stacking in multiloops and exterior loops.
- `coaxial-stacking`: complex ensemble with coaxial stacking in multiloops and exterior loops.
- `nostacking`: complex ensemble without coaxial and dangle stacking in multiloops and exterior loops (ensemble $\overline{\Gamma}(\phi)$).

These ensembles can be used for calculations in combination with either single- or mixed-material parameter sets.

#### S2.4 Validation and implementation of salt corrections

##### S2.4.1 Literature DNA salt corrections for [Na$^+$], [K$^+$], [NH$_4^+$], [Mg$^{++}$]

The default salt condition for DNA[5,8,9,11] parameter sets is [Na$^+$] = 1.0 M. Salt corrections are available for DNA parameters[5,11–13] to permit calculations in user-specified sodium, potassium, ammonium, and magnesium ion concentrations. Following SantaLucia and co-workers, the free energy of a DNA duplex at 37°C is augmented by

$$-0.114\,\frac{N}{2}\log[\mathrm{Na}^+], \tag{S17}$$

for user-specified $0.05\,\mathrm{M} \le [\mathrm{Na}^+] \le 1.0\,\mathrm{M}$, where $N$ is the number of phosphates in the duplex and it is assumed that $\Delta H$ is independent of [Na$^+$], which is valid for this salt regime.[11,12] This salt correction was derived using duplexes with 16 bp or less and the accuracy decreases as duplex length increases further.[11,12] The expression can be generalized to monovalent potassium and ammonium ions[12] as well as to divalent magnesium cations:[5,13]

$$-0.114\,\frac{N}{2}\log\left([\mathrm{Na}^+] + [\mathrm{K}^+] + [\mathrm{NH}_4^+] + 3.3\,[\mathrm{Mg}^{++}]^{1/2}\right), \tag{S18}$$

for user-specified $0.05\,\mathrm{M} \le [\mathrm{Na}^+] + [\mathrm{K}^+] + [\mathrm{NH}_4^+] \le 1.1\,\mathrm{M}$ and $0.0\,\mathrm{M} \le [\mathrm{Mg}^{++}] \le 0.2\,\mathrm{M}$.

The DNA salt correction for a duplex with $N$ phosphates is then defined as:[11,12]

$$\frac{N}{2}\Delta G^{\mathrm{salt}}. \tag{S19}$$

Augmenting equation (2), the salt-corrected secondary structure free energy is then:

$$\Delta G(\phi, s)^{\text{corrected}} \equiv \Delta G(\phi, s) + \frac{N}{2}\Delta G^{\mathrm{salt}}, \tag{S20}$$

where

$$\Delta G^{\mathrm{salt}} \equiv \alpha^{\mathrm{DNA}}_{\mathrm{Na}^+}\,\log\left([\mathrm{Na}^+] + [\mathrm{K}^+] + [\mathrm{NH}_4^+] + 3.3\,[\mathrm{Mg}^{++}]^{1/2}\right)\frac{T}{T_{37}} \tag{S21}$$

for user-specified

$$0.05\,\mathrm{M} \le [\mathrm{Na}^+] + [\mathrm{K}^+] + [\mathrm{NH}_4^+] \le 1.1\,\mathrm{M}, \tag{S22}$$

$$0.0\,\mathrm{M} \le [\mathrm{Mg}^{++}] \le 0.2\,\mathrm{M}, \tag{S23}$$

with $\alpha^{\mathrm{DNA}}_{\mathrm{Na}^+} = -0.114$ kcal/mol and $T_{37} \equiv 310.15$ K.

##### S2.4.2 Regression of an RNA salt correction for [Na$^+$]

To define an RNA salt correction, we regress a value for $\alpha^{\mathrm{RNA}}_{\mathrm{Na}^+}$ using literature melt data for 29 RNA duplexes[14–16] measured at varying Na$^+$ concentrations in the range 0.012 M $\le$ [Na$^+$] $\le$ 0.621 M. Some sequences were measured at multiple Na$^+$ concentrations in that range. Many of the 29 RNA duplexes were also measured at the [Na$^+$] = 1.0 M reference conditions.[14–17] Therefore, $\alpha_{\mathrm{Na}^+}$ was regressed to fit $\Delta G^{\mathrm{salt}}$ (S19) at $T = T_{37}$ to the difference between $\Delta G_{\mathrm{rxn}}$ of the sequence measured at $0.012\,\mathrm{M} \le [\mathrm{Na}^+] \le 0.621$ M and $\Delta G_{\mathrm{rxn}}$ of the sequence measured at [Na$^+$] = 1.0 M ($\Delta G_{\mathrm{rxn}}^{0.012\mathrm{M}\le[\mathrm{Na}^+]\le 0.621\,\mathrm{M}} - \Delta G_{\mathrm{rxn}}^{[\mathrm{Na}^+]=1.0\,\mathrm{M}}$).

For those sequences for which $\Delta G_{\text{rxn}}^{[\text{Na}^+]=1.0\text{ M}}$ was not available in the literature, we used $\Delta G_{\text{rxn}}^{[\text{Na}^+]=1.0\text{ M}} \approx \Delta G(\phi, s)$ computed using equation (2) with parameter set `rna06`. This regression yielded

$$\alpha_{\text{Na}^+}^{\text{RNA}} = -0.114 \text{ kcal/mol}, \tag{S24}$$

so we define the RNA salt correction to be identical to the literature DNA salt correction of SantaLucia and co-workers for $[\text{Na}^+]$.[11,12]

##### S2.4.3 Unified $[\text{Na}^+]$ salt correction for DNA, RNA, and RNA/DNA

Based on our implementation of salt corrections in a dynamic programming context (see Section S2.4.4), if salt-corrections are to be supported for mixed-material calculations, it is necessary to use a unified salt correction for all single-material and mixed-material loops in a secondary structure. Because the same salt correction parameter is derived for single-material DNA and RNA duplexes, we further extend this parameter to mixed RNA/DNA duplexes, $\alpha_{\text{Na}^+}^{\text{RNA/DNA}} = -0.114$ kcal/mol.

Figure S2 displays the computed vs experimental melting temperatures for DNA duplexes (panel a), RNA duplexes (panel b), and RNA/DNA duplexes (panel c) in a range of $\text{Na}^+$ concentrations. DNA results are shown for 159 DNA duplexes[15,18] studied at one or more $\text{Na}^+$ concentrations in the range 0.011 M $\leq [\text{Na}^+] \leq$ 0.810 M (46 of these sequences were compiled by Owczarzy et al.[18] from 20 published sources). RNA results are shown for the same 29 RNA duplexes[14–16] used for the regression above. RNA/DNA results are shown for 40 RNA/DNA hybrid duplexes[15,19] measured at $[\text{Na}^+]$ = 0.122 M.

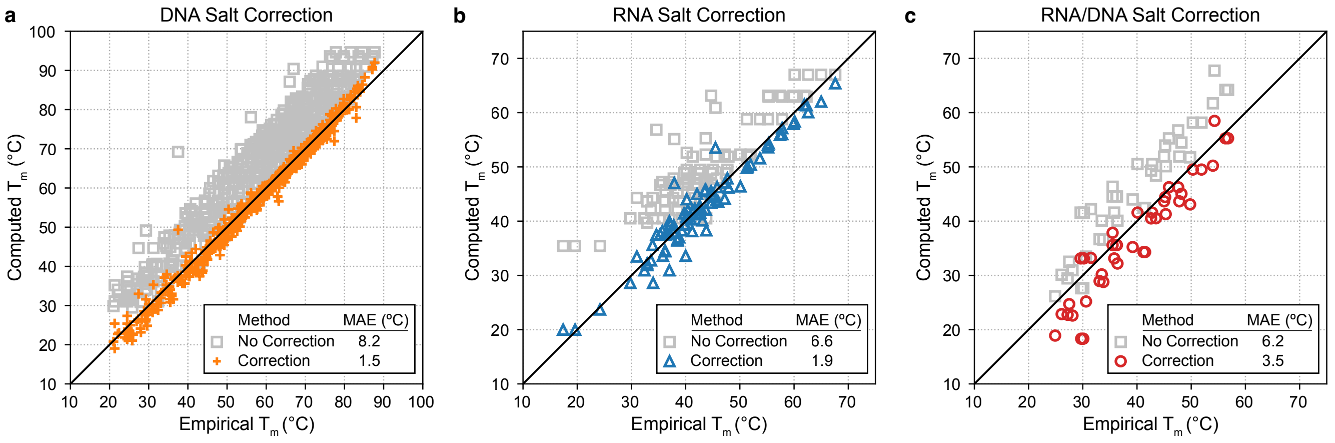

Figure S2: Computational vs experimental melting temperature ($T_m$) with and without $\text{Na}^+$ salt correction for DNA, RNA, and RNA/DNA duplex melt studies. Mean absolute error (MAE). The same SantaLucia salt correction[11,12] was used for RNA, DNA, and RNA/DNA with $\alpha_{\text{Na}^+}^{\text{DNA}} = \alpha_{\text{Na}^+}^{\text{RNA}} = \alpha_{\text{Na}^+}^{\text{RNA/DNA}} = -0.114$ kcal/mol. (a) 159 DNA duplexes[15,18] measured at $\text{Na}^+$ concentrations in the range 0.011 M $\leq [\text{Na}^+] \leq$ 0.810 M. (b) 29 RNA duplexes[14–16] measured at $\text{Na}^+$ concentrations in the range 0.012 M $\leq [\text{Na}^+] \leq$ 0.621 M. (c) 40 RNA/DNA hybrid duplexes[15,19] measured at $[\text{Na}^+]$ = 0.122 M. For each duplex, melting temperature was computed at the strand concentrations used in the original studies.

The $T_m$ for each duplex was calculated using a two-state model where each strand is assumed to be either in a single-stranded unpaired state with no base pairs formed or in a double-stranded duplex state with all base pairs formed. For non-self-complementary sequences (A + B $\leftrightarrow$ AB), this yields:

$$T_m = \frac{\Delta H(\phi, s)}{k \log\left(\frac{c}{4\rho_{\text{H}_2\text{O}}}\right) + \Delta S} \tag{S25}$$

where $c$ is the total strand concentration and $\rho_{\text{H}_2\text{O}}$ is the molarity of water, which follows from setting $f_{\text{ss}}(T) = 0.5$ and $T = T_m$ in equation (S57). For self-complementary sequences (2A $\leftrightarrow$ AA), we have analogously:

$$T_m = \frac{\Delta H(\phi, s)}{k \log\left(\frac{c}{\rho_{\text{H}_2\text{O}}}\right) + \Delta S} \tag{S26}$$

which follows from setting $f_{\text{ss}}(T) = 0.5$ and $T = T_m$ in the self-complementary analog to (S57):

$$\exp\left(-\frac{\Delta H_{\text{rxn}}}{kT} + \frac{\Delta S_{\text{rxn}}}{k}\right) = \frac{(1 - f_{\text{ss}}(T))\rho_{\text{H}_2\text{O}}}{2 f_{\text{ss}}(T)^2 c}. \tag{S27}$$

For $T_m$ computed without a salt correction, let

$$\Delta S \equiv \Delta S(\phi, s) = \frac{\Delta H(\phi, s) - \Delta G(\phi, s)}{310.15\,\text{K}} \tag{S28}$$

and for $T_m$ computed with a salt correction, let

$$\Delta S \equiv \Delta S(\phi, s)^{\text{corrected}} = \frac{\Delta H(\phi, s) - \Delta G(\phi, s)^{\text{corrected}}}{310.15\,\text{K}} \tag{S29}$$

where $\Delta G(\phi, s)$ and $\Delta H(\phi, s)$ are computed using `dna04.2` for DNA, `rna06` for RNA, and `rna_dna06` for RNA/DNA. The $T_m$ predictions of Figures S2a and S2b indicate that the unified salt correction of equation (S30) is comparably effective for both RNA and DNA. For DNA, the mean absolute error (MAE) is 8.2 °C without salt correction and 1.5 °C with salt correction. For RNA, the MAE is 6.6 °C without salt correction and 1.9 °C with salt correction. On the other hand, the $T_m$ predictions of Figure S2c suggest that the unified salt correction is not as well-suited for RNA/DNA duplexes, although it still provides a significant improvement over not using a salt correction. For RNA/DNA, the MAE is 6.2 °C without salt correction and 3.5 °C with salt correction.

The unified [$Na^+$] salt correction for DNA, RNA, and RNA/DNA is then

$$\Delta G^{\text{salt}} \equiv \alpha_{\text{Na}^+} \log[\text{Na}^+] \frac{T}{T_{37}} \tag{S30}$$

with

$$\alpha_{\text{Na}^+} = -0.114\ \text{kcal/mol}. \tag{S31}$$

The literature DNA salt correction has been validated for the range $0.05\,\text{M} \le [\text{Na}^+] \le 1.1\,\text{M}$.[11,12] The RNA regression and validation were based on melt studies performed in the range $0.012\,\text{M} \le [\text{Na}^+] \le 1.0\,\text{M}$. To be conservative, we define the RNA salt correction in the range

$$0.05\,\text{M} \le [\text{Na}^+] \le 1.0\,\text{M}, \tag{S32}$$

adopting the lower bound from the DNA studies and the upper bound from the RNA studies. Validation of the RNA/DNA salt correction was based on melt studies performed at [$Na^+$] = 0.12 M so we define the RNA/DNA salt correction in the range

$$0.12\,\text{M} \le [\text{Na}^+] \le 1.0\,\text{M}. \tag{S33}$$

##### S2.4.4 Implementation of salt corrections in dynamic programs

The default salt conditions for RNA,[4,6,7,9,20,21] DNA,[5,8,9,11] and the new RNA/DNA parameter sets are all [$Na^+$] = 1.0 M. To apply the salt correction of equation (S19) to a complex of $L$ strands at temperature $T$, consider a secondary structure $s$ containing one or more duplexes. We assume that strands are synthesized with one phosphate per base so that $N/2 = n_{\text{bp}}(s)$ where $N$ is the total number of phosphates in duplexes and $n_{\text{bp}}(s)$ is the total number of base pairs in $s$. (If strands are synthesized without a $5'$ terminal phosphate, then $2n_{\text{bp}}(s)$ approximates the total number of phosphates in duplexes.) We further assume that $\Delta H$ is independent of cation concentration in the regime of the salt correction. To facilitate incorporation into dynamic programs, the salt correction of equation (S19) is implemented in the form

$$n_{\text{bp}}(s)\Delta G^{\text{salt}}. \tag{S34}$$

To incorporate this salt correction in dynamic programs without explicitly calculating $n_{\text{bp}}(s)$,[1] note that for a complex of $L$ strands, the total number of loops in each secondary structure is

$$n_{\text{loop}}(s) = n_{\text{bp}}(s) + 1. \tag{S35}$$

This may be seen, for example, by starting with a single strand with no base pairs (corresponding to a single exterior loop). Each addition of a base pair adds one loop. Once all base pairs in $s$ have been added, each addition of a nick increases the number of strands by one without changing the number of loops (all secondary structures in the complex ensemble are connected so introduction of each nick converts a loop from another type to an exterior loop). Let $n_{\text{loop}}^{\text{other}}$ denote the total number of non-exterior loops and $n_{\text{loop}}^{\text{exterior}}$ denote the total number of exterior loops, so we have

$$n_{\text{loop}}(s) = n_{\text{loop}}^{\text{exterior}}(s) + n_{\text{loop}}^{\text{other}}(s). \tag{S36}$$

For a complex of $L$ strands, $n_{\text{loop}}^{\text{exterior}}(s) = L$. Thus, the salt correction (S34) becomes

$$n_{\text{bp}}(s)\Delta G^{\text{salt}} = (L-1)\Delta G^{\text{salt}} + n_{\text{loop}}^{\text{other}}(s)\Delta G^{\text{salt}}. \tag{S37}$$

Hence, the salt correction can be implemented by adding

$$\Delta G^{\text{salt}} \tag{S38}$$

to every $\Delta G(\text{loop})$ except for exterior loops as a pre-processing step, using our suite of dynamic programs without modification, and then treating the constant term $(L-1)\Delta G^{\text{salt}}$ in a post-processing step (see Section S2.6).

#### S2.5 Temperature dependence of the free energy model

The loop-based free energy model (1) is temperature dependent. Each loop free energy is calculated using

$$\Delta G(\text{loop}) = \Delta H(\text{loop}) - T\Delta S(\text{loop}) \tag{S39}$$

where $T$ is in Kelvin and $\Delta H(\text{loop})$ and $\Delta S(\text{loop})$ are assumed to be temperature independent.[12] Model parameters are provided for RNA[4,6,7,9,20,21] and DNA[5,8,9,11] in the form of $\Delta G_{37}(\text{loop})$ and $\Delta H(\text{loop})$ which can be used to calculate

$$\Delta S(\text{loop}) = \frac{1}{T_{37}}[\Delta H(\text{loop}) - \Delta G_{37}(\text{loop})] \tag{S40}$$

with $T_{37} = 310.15$ K, so (S39) becomes

$$\Delta G(\text{loop}) = \Delta H(\text{loop}) - \frac{T}{T_{37}}[\Delta H(\text{loop}) - \Delta G_{37}(\text{loop})]. \tag{S41}$$

Similarly, for the strand association penalty (S1):

$$\Delta G_{\text{pub}}^{\text{assoc}} = \Delta H_{\text{pub}}^{\text{assoc}} - T\Delta S_{\text{pub}}^{\text{assoc}}. \tag{S42}$$

and the provided parameters $\Delta G_{37,\text{pub}}^{\text{assoc}}$ and $\Delta H_{\text{pub}}^{\text{assoc}}$ yield

$$\Delta G_{\text{pub}}^{\text{assoc}} = \Delta H_{\text{pub}}^{\text{assoc}} - \frac{T}{T_{37}}[\Delta H_{\text{pub}}^{\text{assoc}} - \Delta G_{37,\text{pub}}^{\text{assoc}}]. \tag{S43}$$

The temperature dependence is explicit in the form of the symmetry correction (3) and salt correction (S34).

#### S2.6 Treatment of constant free energy terms for complex ensembles

Consider a complex of $L$ strands containing a total of $N$ nucleotides. Suppose that free energy model terms have been pre-processed as described above for units [see (S1)], salt corrections [see (S38)], and temperature corrections [see (S41 and S42)] prior to calculating any physical quantities. The secondary structure free energy (1) then becomes

$$\overline{\Delta G}(\phi, s) = (L-1)[\Delta G^{\text{assoc}} + \Delta G^{\text{salt}}] + \sum_{\text{loop}\in s} \Delta G(\text{loop}). \tag{S44}$$

After running the partition function dynamic program to calculate $Q_{1,N}$, the partition function is then

$$\overline{Q}(\phi) = \exp\{-(L-1)[\Delta G^{\text{assoc}} + \Delta G^{\text{salt}}]/kT\}Q_{1,N}. \tag{S45}$$

where this post-processing step accounts for the constant terms $\Delta G^{\text{assoc}}$ and $\Delta G^{\text{salt}}$ that affect all secondary structures in the complex ensemble. Likewise, after running the MFE dynamic program to calculate $F_{1,N}$, the free energy of the MFE stacking state is then

$$\overline{\Delta G}(\phi, s^{\shortparallel}_{\text{MFE}}) = (L-1)[\Delta G^{\text{salt}} + \Delta G^{\text{assoc}}] + F_{1,N}. \tag{S46}$$

The equilibrium base-pairing probability $\overline{P}_{i,j}(\phi)$ is calculated via equation (17) of Reference 1 using the values of $Q^b_{i,j}(\phi)$, $Q^b_{j,N+i}(\phi')$ and $Q_{1,N}(\phi)$ returned by the dynamic program; the constant terms $\Delta G^{\text{assoc}}$ and $\Delta G^{\text{salt}}$ do not affect the calculation as they are omitted in both the numerator and the denominator. The dynamic programs for calculating the MFE proxy structure, suboptimal structures, or sampled structures are unaffected by the constant terms $\Delta G^{\text{assoc}}$ and $\Delta G^{\text{salt}}$ so no post-processing is required for those quantities.

#### S2.7 Distinguishability issues

For a complex of $L$ strands, the ensemble $\overline{\Gamma}$ treats each strand as distinct while the ensemble $\Gamma$ treats strands with the same sequence as indistinguishable.[1,2] Both ensembles have conceptual utility as they provide different perspectives when examining the physical properties of a complex. In laboratory experiments, strands with the same sequence are typically indistinguishable, so calculations over ensemble $\Gamma$ are crucial for comparison to experimental data (e.g., equilibrium secondary structure probabilities and equilibrium complex concentrations). On the other hand, calculations over ensemble $\overline{\Gamma}$ can sometimes provide information that is valuable precisely because it cannot be measured experimentally (e.g., equilibrium base-pairing probability matrix).

All of the dynamic programs described in the present work operate on ensemble $\overline{\Gamma}$ using free energy model (1) where each strand is treated as distinct. This is a matter of algorithmic necessity, as the free energy model (2) used for ensemble $\Gamma$ contains a symmetry correction that depends on the global rotational symmetry $R$ of each secondary structure $s \in \Gamma$. For efficiency reasons, the dynamic programs avoid explicitly enumerating each structure, instead operating on local loop free energies to incorporate information for multiple structures simultaneously while operating only on local loop free energies. As a result, the dynamic programs cannot incorporate a different global rotational symmetry correction for each structure because they never have access to global structural information. However, to facilitate comparisons to experimental data, physical quantities calculated using a dynamic program over ensemble $\overline{\Gamma}$ using physical model (1) can be post-processed to obtain the corresponding physical quantities over ensemble $\Gamma$ using physical model (2). See Section S5 of Reference 1 for details.

#### S2.8 Curve fitting for duplex melt experiments

For each replicate melt experiment, UV absorbance data were fit with a two-state model as described previously,[22] where each strand is assumed to be either in a single-stranded (ss) unpaired state with no base pairs formed (A and B) or in a double-stranded (ds) duplex state AB with all base pairs formed:

$$\underbrace{\mathrm{A}+\mathrm{B}}_{\text{ss state}} \leftrightarrow \underbrace{\mathrm{AB}}_{\text{ds state}} . \tag{S47}$$

At equilibrium (see equation (S4)):

$$\exp\left(-\frac{\Delta G_{\mathrm{AB}} - \Delta G_{\mathrm{A}} - \Delta G_{\mathrm{B}}}{kT}\right) = \frac{x_{\mathrm{AB}}}{x_{\mathrm{A}} x_{\mathrm{B}}} \tag{S48}$$

subject to conservation of mass

$$x_{\mathrm{A}} + x_{\mathrm{AB}} = x^0_{\mathrm{A}}, \tag{S49a}$$

$$x_{\mathrm{B}} + x_{\mathrm{AB}} = x^0_{\mathrm{B}}, \tag{S49b}$$

where $x_{\mathrm{A}}$, $x_{\mathrm{B}}$, and $x_{\mathrm{AB}}$ are mole fractions (see Section S2.1), and $x^0_{\mathrm{A}}$ and $x^0_{\mathrm{B}}$ represent the total concentration of strands A and B present in the experiment. For our melt experiments, the two strands are introduced at equal concentration ($x^0_{\mathrm{A}} = x^0_{\mathrm{B}}$) and are also at equal concentration at equilibrium ($x_{\mathrm{A}} = x_{\mathrm{B}}$), so the right hand side of (S48) can be restated as:

$$\frac{x_{\mathrm{AB}}}{x_{\mathrm{A}} x_{\mathrm{B}}} = \frac{[\mathrm{AB}]\rho_{\mathrm{H_2O}}}{[\mathrm{A}][\mathrm{B}]} \tag{S50}$$

$$= \frac{([\mathrm{A}]_0 - [\mathrm{A}])\rho_{\mathrm{H_2O}}}{[\mathrm{A}]^2} \tag{S51}$$

$$= \frac{2(1 - f_{\mathrm{ss}}(T))\rho_{\mathrm{H_2O}}}{f_{\mathrm{ss}}(T)^2 c} \tag{S52}$$

where $\rho_{\mathrm{H_2O}}$ is the molarity of water, $f_{\mathrm{ss}}(T)$ is the temperature-dependent fraction of strands in the single-stranded state, and

$$c \equiv [A]_0 + [B]_0 = 2[A]_0 \tag{S53}$$

is the total concentration of strands in the experiment (determined by measuring absorbance at 260 nm). The left hand side of (S48) can be restated as:

$$\exp\left(-\frac{\Delta G_{\mathrm{AB}} - \Delta G_{\mathrm{A}} - \Delta G_{\mathrm{B}}}{kT}\right) = \exp\left(-\frac{\Delta H_{\mathrm{rxn}}}{kT} + \frac{\Delta S_{\mathrm{rxn}}}{k}\right) \tag{S54}$$

in terms of the reaction enthalpy and entropy (assumed to be temperature-independent):

$$\Delta H_{\text{rxn}} \equiv \Delta H_{\text{AB}} - \Delta H_{\text{A}} - \Delta H_{\text{B}}, \tag{S55}$$
$$\Delta S_{\text{rxn}} \equiv \Delta S_{\text{AB}} - \Delta S_{\text{A}} - \Delta S_{\text{B}}. \tag{S56}$$

Hence, we may restate equation (S48) in the form

$$\exp\left(-\frac{\Delta H_{\text{rxn}}}{kT} + \frac{\Delta S_{\text{rxn}}}{k}\right) = \frac{2(1 - f_{\text{ss}}(T))\rho_{\text{H}_2\text{O}}}{f_{\text{ss}}(T)^2 c}. \tag{S57}$$

The temperature-dependent absorbance of the solution, $A(T)$, is modeled using the two-state model as

$$A(T) = f_{\text{ss}} A_{\text{ss}}(T) + (1 - f_{\text{ss}}(T)) A_{\text{ds}}(T), \tag{S58}$$

where $A_{\text{ss}}(T)$ and $A_{\text{ds}}(T)$ are the absorbances of the the single- and double-stranded states, respectively, which are assumed to be linear functions of temperature:

$$A_{\text{ss}}(T) \equiv m_{\text{ss}} T + b_{\text{ss}}, \tag{S59a}$$
$$A_{\text{ds}}(T) \equiv m_{\text{ds}} T + b_{\text{ds}}. \tag{S59b}$$

Absorbance curve fits were performed using the Levenberg–Marquardt (LM) algorithm for nonlinear least squares optimization.[22] For each curve fit, the modeled absorbance, $A(T)$, governed by (S57) and (S58) is fit to a set of experimental absorbance values $A_i$ measured at temperatures $T_i \in \{5, 6, \ldots, 90\}$°C. The LM algorithm returns a set of temperature-dependent $f_{\text{ss}}(T_i)$ values as well as values for the six unknowns $m_{\text{ss}}$, $b_{\text{ss}}$, $m_{\text{ds}}$, $b_{\text{ds}}$, $\Delta H_{\text{rxn}}$, and $\Delta S_{\text{rxn}}$. The experimental melting temperature, $T_m$, is defined as the temperature at which $f_{\text{ss}}(T) = 0.5$.

#### S2.9 Calculation of diverse physical quantities

##### S2.9.1 Quantities defined over a complex ensemble

Consider a complex with sequence $\phi$. NUPACK 4 enables calculation of the diverse physical quantities listed below for single-material complexes.[1] Here, we provide dynamic programs to calculate these quantities for mixed-material complexes:

- the partition function,

$$\overline{Q}(\phi) = \sum_{s \in \overline{\Gamma}(\phi)} e^{-\overline{\Delta G}(\phi,s)/kT}, \tag{S60}$$

  over ensemble $\overline{\Gamma}(\phi)$ treating all strands as distinct. The equilibrium probability of any secondary structure $s \in \overline{\Gamma}(\phi)$ is then

$$\overline{p}(\phi, s) = e^{-\overline{\Delta G}(\phi,s)/kT} / \overline{Q}(\phi). \tag{S61}$$

  Post-processing $\overline{Q}(\phi)$ yields the partition function $Q(\phi)$ over ensemble $\Gamma(\phi)$ treating strands with the same sequence as indistinguishable.[1,2]

- the base-pairing probability matrix $\overline{P}(\phi)$ with entries $\overline{P}^{i,j}(\phi) \in [0, 1]$ corresponding to the probability

$$\overline{P}^{i,j}(\phi) = \sum_{s \in \overline{\Gamma}(\phi)} \overline{p}(\phi, s) S^{i,j}(s) \tag{S62}$$

  that base pair $i \cdot j$ forms at equilibrium within ensemble $\overline{\Gamma}(\phi)$, treating all strands as distinct. Here, $S(s)$ is a *structure matrix* with entries $S^{i,j}(s) = 1$ if structure $s$ contains base pair $i \cdot j$ and $S^{i,j}(s) = 0$ otherwise. Abusing notation, the entry $S^{i,i}(s)$ is 1 if base $i$ is unpaired in structure $s$ and 0 otherwise; the entry $P^{i,i}(\phi) \in [0, 1]$ denotes the equilibrium probability that base $i$ is unpaired over ensemble $\overline{\Gamma}(\phi)$. Hence $S(s)$ and $\overline{P}(\phi)$ are symmetric matrices with row and column sums of 1.

- the free energy of the minimum free energy (MFE) stacking state $s''_{\text{MFE}}(\phi) \in \overline{\Gamma}''(\phi)$ treating all strands as distinct:

$$\overline{\Delta G}(\phi, s''_{\text{MFE}}) = \min_{s'' \in \overline{\Gamma}''(\phi)} \overline{\Delta G}(\phi, s''). \tag{S63}$$

- the MFE proxy structure

$$s_{\text{MFE}'} = \{s \in \overline{\Gamma}(\phi) | s^{\shortparallel}_{\text{MFE}} \in s, s^{\shortparallel}_{\text{MFE}}(\phi) = \arg\min_{s^{\shortparallel} \in \overline{\Gamma}^{\shortparallel}(\phi)} \overline{\Delta G}(\phi, s^{\shortparallel})\}. \tag{S64}$$

defined as the secondary structure containing the MFE stacking state within its subensemble. If there is more than one MFE stacking state, the algorithm returns all corresponding MFE proxy structures.

- the set of suboptimal secondary structures

$$\overline{\Gamma}_{\text{subopt}}(\phi, \Delta G_{\text{gap}}) = \{s \in \overline{\Gamma}(\phi) | s^{\shortparallel} \in s, \overline{\Delta G}(\phi, s^{\shortparallel}) \leq \overline{\Delta G}(\phi, s^{\shortparallel}_{\text{MFE}}) + \Delta G_{\text{gap}}\} \tag{S65}$$

with stacking states within a specified $\Delta G_{\text{gap}} \geq 0$ of the MFE stacking state.

- a set of $J$ secondary structures Boltzmann sampled from ensemble $\overline{\Gamma}(\phi)$ treating all strands as distinct:

$$\overline{\Gamma}_{\text{sample}}(\phi, J) \in \overline{\Gamma}(\phi). \tag{S66}$$

Post-processing then yields the set of $J$ secondary structures Boltzmann sampled from ensemble $\Gamma(\phi)$ treating strands with the same sequence as indistinguishable:

$$\Gamma_{\text{sample}}(\phi, J) \in \Gamma(\phi). \tag{S67}$$

##### S2.9.2 Quantities defined over a test tube ensemble

Now consider a test tube ensemble containing an arbitrary set of strand species $\Psi^0$ interacting to form an arbitrary set of complex species $\Psi$. NUPACK 4 enables calculation of the physical quantities listed below for single-material test tubes.[1] We provide algorithms to calculate these quantities for mixed-material test tubes:

- the set of equilibrium concentrations $x_\Psi \equiv x_c \; \forall c \in \Psi$, (specified as mole fractions) that are the unique solution to the strictly convex optimization problem:[2]

$$\min_{x_\Psi} \sum_{c \in \Psi} x_c (\log x_c - \log Q_c - 1) \tag{S68a}$$

$$\text{subject to} \quad \sum_{c \in \Psi} A_{i,c} x_c = x_i^0 \quad \forall i \in \Psi^0, \tag{S68b}$$

expressed in terms of the previously calculated set of partition functions $Q_\Psi$. Here, the constraints impose conservation of mass: $A$ is the stoichiometry matrix such that $A_{i,c}$ is the number of strands of type $i$ in complex $c$, and $x_i^0$ is the total concentration of strand $i$ present in the test tube. Based on dimensional analysis,[2] the convex optimization algorithm operates on mole fractions, but for convenience, accepts molar strand concentrations $[i]^0 = x_i^0 \rho_{\text{H}_2\text{O}}$ as inputs and returns molar complex concentrations $[c] = x_c \rho_{\text{H}_2\text{O}}$ as outputs, where $\rho_{\text{H}_2\text{O}}$ is the molarity of water.

- the ensemble pair fractions for the test tube ensemble, for example

$$f_A(i_A \cdot j_B) \tag{S69}$$

denotes the fraction of A strands that form base pair $i_A \cdot j_B$ (correspondingly $f_B(i_A \cdot j_B)$ denotes the fraction of B strands that form base pair $i_A \cdot j_B$). In order to calculate these base-pairing observables, it is first necessary to calculate the set of equilibrium concentrations $x_\Psi$ and the set of base-pairing probability matrices $\overline{P}_\Psi$.

##### S2.9.3 Unified dynamic programming framework

With the unified dynamic programming framework introduced in Reference 1, each dynamic program combines three ingredients (Figure S3): (1) a set of model-specific recursions that specify the dependencies of each subproblem and incorporate the structural details of the complex ensemble and the energetic details of the loop-based free energy model, (2) quantity-specific evaluation algebras that define the mathematical form of each subproblem for a given physical quantity, (3) quantity-specific operation orders that define the computational trajectory through the dependency graph of subproblems for a given physical quantity. Hence, to implement mixed-material dynamic programs, here we define new mixed-material recursions (see Section S6) and leverage the previously defined evaluation algebras (see Section S3 of Reference 1) and operation orders (see Section S4 of Reference 1) without modification. The algorithmic ingredients used to calculate each physical quantity are summarized in Table 1 of Reference 1.

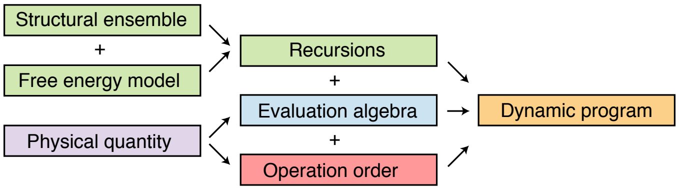

Figure S3: Unified dynamic programming framework. To calculate a physical quantity of interest based on a physical model comprising a structural ensemble and a free energy model, each dynamic program combines three ingredients: model-specific recursions, a quantity-specific evaluation algebra, and a quantity-specific operation order. Used with permission from Fornace et al., *ACS Synth Biol*, 9, 2665-2678, 2020. Copyright 2020 American Chemical Society.

### S3 Single-material free energy model details: RNA, DNA, 2′OMe-RNA

#### S3.1 Single-material parameter sets

A single-material parameter set (either RNA, DNA, or 2′OMe-RNA) contains parameters for single-material loops. The following temperature-dependent parameter sets are supported:

- `rna95`: RNA parameters based on (Serra & Turner, 1995)[7] with additional parameters[9] including coaxial stacking[4,6] and dangle stacking[6,7,9] in a user-specified concentration of $Na^+$.
- `dna04.1`: DNA parameters based on (SantaLucia, 1998)[11] and (SantaLucia & Hicks, 2004)[12] with additional parameters[9] including coaxial stacking[5] and dangle stacking[8,9] in user-specified concentrations of $Na^+$, $K^+$, $NH_4^+$, and $Mg^{++}$.[5,11–13]
- `dna04.2`: DNA parameters based on `dna04.1` with updated GT internal mismatch values,[12,23–27] internal asymmetry values,[12] and terminal mismatch values[6,28] in user-specified concentrations of $Na^+$, $K^+$, $NH_4^+$, and $Mg^{++}$.[5,11–13]
- `rna06`: RNA parameters based on (Mathews et al., 1999),[4] (Mathews et al., 2004),[29] and (Lu et al., 2006)[21] with additional parameters[9,20] including coaxial stacking[4,6] and dangle stacking[6,7,9] in a user-specified concentration of $Na^+$.
- `merna06`: 2′OMe-RNA parameters for calculations in 1.2 M $Na^+$ based on (Kierzek et al., 2006)[30] using stack loop, coaxial stacking, terminal base pair, and strand association values from the RNA/2′OMe-RNA mixed-material parameter set `rna_merna06` and all other values from the RNA single-material parameter set `rna06`. Note that due to the unavailability of single-material 2′OMe-RNA melt experiments, this parameter set is based on values regressed from mixed-material RNA/2′OMe-RNA melt experiments and single-material RNA melt experiments. This single-material parameter set is needed for use in conjunction with that the mixed-material parameter set `rna_merna06` which requires both `rna06` and `merna06`.
- `custom`: Custom parameters based on user-specified parameters representing nucleic acids or synthetic nucleic acid analogs in experimental conditions of choice.

#### S3.2 Definition of single-material Watson–Crick and wobble pairs

Single-material base pairs are either Watson–Crick pairs:

- RNA: rG·rC and rA·rU,
- DNA: dG·dC and dA·dT,
- 2′OMe-RNA: mG·mC and mA·mU,

or wobble pairs:

- RNA: rG·rU,
- 2′OMe-RNA: mG·mU.

Note that for DNA, dG and dT are considered to form a mismatch dG×dT and not a wobble pair.[12]

#### S3.3 Historical single-material ensembles and parameter sets (for backwards compatibility with NUPACK 3)

For backwards compatibility, the following historical single-material complex ensembles without coaxial stacking and with approximate dangle stacking are supported (see Section S2.5 in Reference 1 for details):

- `none-nupack3`: no dangle stacking and no coaxial stacking (dangles "`none`" option for NUPACK 3)
- `some-nupack3`: some dangle stacking and no coaxial stacking (dangles "`some`" option for NUPACK 3)
- `all-nupack3`: all dangle stacking and no coaxial stacking (dangles "`all`" option for NUPACK 3)

For these historical ensembles, base pairs are either Watson–Crick pairs (G·C and A·U for RNA; G·C and A·T for DNA) or wobble pairs (G·U for RNA; G·T for DNA). Note that for the historical ensembles, G·T is classified as a DNA wobble pair and not as a mismatch. The historical ensembles prohibit a wobble pair (G·U or G·T) as a terminal base pair in an exterior loop or a multiloop. As a result, an attempt to evaluate a free energy for a sequence $\phi$ and secondary structure $s$ that place a wobble pair as a terminal base pair in an exterior loop or multiloop will return $\overline{\Delta G}(\phi, s) = \Delta G(\phi, s) = \infty$. These historical ensembles can be used for calculations in combination with the following historical DNA and RNA parameter sets:

- `rna95-nupack3` is the same as `rna95` except that terminal mismatch free energies in exterior loops and multiloops are replaced by two dangle stacking free energies (see equation (S55) of Reference 1).
- `dna04-nupack3` is the same as `dna04.1` except that G·T was treated as a wobble pair (analogous to a G·U RNA wobble pair) instead of being classified as a G×T mismatch. Note that while terminal mismatch free energies in exterior loops and multiloops are replaced by two dangle stacking free energies (see equation (S55) of Reference 1), this is the same treatment as in `dna04.1`, as terminal mismatch parameters are not public for DNA.[12]
- `rna99-nupack3` based on (Mathews et al., 1999)[4] with additional parameters[9,20] including dangle stacking[6,7,9] in 1M $Na^+$. Terminal mismatch free energies in exterior loops and multiloops are replaced by two dangle stacking free energies (see equation (S55) of Reference 1). Parameters are provided only for 37 °C.

#### S3.4 Single-material salt conditions

##### S3.4.1 DNA salt conditions

The default salt condition for DNA[5,8,9,11] parameter sets is [$Na^+$] = 1.0 M. For a DNA secondary structure $s$ containing $n_{\rm bp}$ base pairs, the free energy, $\Delta G(\phi, s)$, is augmented by (see Section S2.4):

$$n_{\rm bp}(s)\Delta G^{\rm salt} \tag{S70}$$

with

$$\Delta G^{\rm salt} = -0.114\,\log\left([{\rm Na}^+] + [{\rm K}^+] + [{\rm NH}_4^+] + 3.3\,[{\rm Mg}^{++}]^{1/2}\right)\frac{T}{T_{37}} \tag{S71}$$

for user-specified

$$0.05\,{\rm M} \le [{\rm Na}^+] + [{\rm K}^+] + [{\rm NH}_4^+] \le 1.1\,{\rm M}, \tag{S72}$$

$$0.0\,{\rm M} \le [{\rm Mg}^{++}] \le 0.2\,{\rm M}, \tag{S73}$$

with $T_{37} = 310.15$ K.

##### S3.4.2 RNA salt conditions

The default salt condition for RNA[4,6,7,9,20,21] parameter sets is [$Na^+$] = 1.0 M. For an RNA secondary structure $s$ containing $n_{\rm bp}$ base pairs, the free energy, $\Delta G(\phi, s)$, is augmented by (see Section S2.4):

$$n_{\rm bp}(s)\Delta G^{\rm salt} \tag{S74}$$

with

$$\Delta G^{\rm salt} = -0.114\,\log\left([{\rm Na}^+]\right)\frac{T}{T_{37}} \tag{S75}$$

for user-specified

$$0.05\,{\rm M} \le [{\rm Na}^+] \le 1.0\,{\rm M}, \tag{S76}$$

with $T_{37} = 310.15$ K.

##### S3.4.3 2′OMe-RNA salt conditions

The default salt condition for 2′OMe-RNA[30] parameter sets is [$\text{Na}^+$] = 0.12 M based on the experimental conditions used for RNA/2′OMe-RNA melt studies. Due to the unavailability of empirical melt data for other salt conditions, no salt correction is supported to enable 2′OMe-RNA calculations at other $\text{Na}^+$ concentrations.

#### S3.5 Single-material strand association penalty

The strand association penalty, $\Delta G^{\text{assoc}}_{\text{pub}}$, uses empirical values for `rna95`,[20] `rna06`,[20] and `dna04.1/dna04.2`.[12] In the absence of an empirical 2′OMe-RNA-only value, `merna06` uses the empirical value derived for RNA/2′OMe-RNA hybrid duplexes.[30]

#### S3.6 Free energy model for single-material hairpin loops

A hairpin loop is defined for a subsequence $\phi_{[i:j]}$ by the single base pair $i \cdot j$ such that there are no nicks or additional base pairs in the range $[i : j]$. Let $n \equiv j - i - 1$ denote the number of unpaired nucleotides in the hairpin loop. Steric effects are assumed to prevent hairpin loops with $n < 3$ for both RNA[6,7] and DNA.[12] The functional form of the hairpin free energy is as follows:

$$\Delta G^{\text{hairpin}}(\phi_{[i:j]}) = \Delta G^{\text{hairpin}}_{\text{size}}(n) + \Delta G^{\text{hairpin}}_{\text{seq}}(\phi_{[i:j]}) \tag{S77}$$

For the size-dependent term:[4,7,12,21]

$$\Delta G^{\text{hairpin}}_{\text{size}}(n) = \begin{cases} \infty, & n < 3 \\ \Delta G^{\text{hairpinsize}}_{n}, & 3 \le n \le 30 \\ \Delta G^{\text{hairpinsize}}_{30} + \log\left(\frac{n}{30}\right) \Delta G^{\text{polymer}}_{\text{entropy}}, & n > 30 \end{cases} \tag{S78}$$

- $\Delta G^{\text{hairpinsize}}_{n}$: a lookup table up to $n = 30$. `rna95` and `rna06` populate the lookup table using empirical values of $\Delta G^{\text{hairpinsize}}_{n}$ up to $n = 9$ and logarithmic extrapolation for larger $n$.[4,7,21] `dna04.1/dna04.2` populate the lookup table using empirical values of $\Delta G^{\text{hairpinsize}}_{n}$ for a subset of $3 \le n \le 30$ and logarithmic interpolation for the other values.[12]
- $\Delta G^{\text{polymer}}_{\text{entropy}}$: a logarithmic extrapolation parameter based on Jacobson-Stockmayer polymer theory for $n > 30$. `rna95`, `dna04.1/dna04.2`, and `rna06` use previously published values.[7,12]

For the sequence-dependent term:[4,12,21]

$$\Delta G^{\text{hairpin}}_{\text{seq}}(\phi_{[i:j]}) = \begin{cases} \Delta G^{\text{triloop}}_{\phi_{[i:j]}} + \Delta G^{\text{terminalbp}}_{\phi_j,\phi_i} & n = 3 \\ \Delta G^{\text{tetraloop}}_{\phi_{[i:j]}} & n = 4 \\ \Delta G^{\text{hairpinmm}}_{\phi_{j-1},\phi_j,\phi_i,\phi_{i+1}} & n \ge 5 \end{cases} \tag{S79}$$

- $\Delta G^{\text{triloop}}_{\phi_{[i:j]}}$: sequence-dependent penalty for hairpin loop of length $n = 3$. 0 kcal/mol for `rna95`.[7] Empirical values for `dna04.1/dna04.2`[12] and `rna06`.[29] `merna06` uses `rna06` values treating RNA nucleotides as 2′OMe-RNA nucleotides.
- $\Delta G^{\text{terminalbp}}_{\phi_j,\phi_i}$: sequence-dependent penalty for non-GC terminal base pair at the end of a duplex. Empirical values for `rna95`,[7] `dna04.1/dna04.2`,[12] and `rna06`.[29] `merna06` uses the `rna_merna06` values (defined in Section S5.5) treating RNA nucleotides as 2′OMe-RNA nucleotides.
- $\Delta G^{\text{tetraloop}}_{\phi_{[i:j]}}$: sequence-dependent penalty for hairpin loop of length $n = 4$. Empirical values for `rna95`,[7] `dna04.1/dna04.2`,[12] and `rna06`.[29] `merna06` uses `rna06` values treating RNA nucleotides as 2′OMe-RNA nucleotides.
- $\Delta G^{\text{hairpinmm}}_{\phi_{j-1},\phi_j,\phi_i,\phi_{i+1}}$: sequence-dependent term for mismatched bases adjacent to base pair $i \cdot j$. Empirical values set equal to $\Delta G^{\text{terminalmm}}_{\phi_{j-1},\phi_j,\phi_i,\phi_{i+1}}$ (defined in Section S3.8) plus sequence-dependent modifications for `rna95`[7] and `rna06`.[4,21] `dna04.1` uses unpublished values for $\Delta G^{\text{hairpinmm}}_{\phi_{j-1},\phi_j,\phi_i,\phi_{i+1}}$ made available in the Mfold software.[9] `dna04.2` sets $\Delta G^{\text{hairpinmm}}_{\phi_{j-1},\phi_j,\phi_i,\phi_{i+1}}$ equal to empirical values for $\Delta G^{\text{terminalmm}}_{\phi_{j-1},\phi_j,\phi_i,\phi_{i+1}}$ (defined in Section S3.8). `merna06` uses `rna06` values treating RNA nucleotides as 2′OMe-RNA nucleotides.

#### S3.7 Free energy model for single-material interior loops

An interior loop may be defined via a pair of subsequences $\phi_{[i:d]}$ and $\phi_{[e:j]}$ such that $i < d < e < j$ with base pairs $i \cdot j$ and $d \cdot e$, with no additional paired bases or nicks within the two subsequences.

**Stack loops.** Stack loops are the special case where $d = i + 1$ and $j = e + 1$.

$$\Delta G^{\text{stackedpair}}(\phi_{[i:i+1]}, \phi_{[j-1:j]}) = \Delta G^{\text{stack}}_{\phi_j,\phi_i,\phi_{i+1},\phi_{j-1}} \tag{S80}$$

- $\Delta G^{\text{stack}}_{\phi_j,\phi_i,\phi_{i+1},\phi_{j-1}}$: the stack free energy has been determined for all allowable base pair combinations from experimental results for `rna95`,[7] `rna06`,[4,20,21] and `dna04.1/dna04.2`.[12] `merna06` uses the `rna_merna06` values (defined in Section S5.6) treating RNA nucleotides as 2′OMe-RNA nucleotides.

**Bulge loops.** A bulge loop is the special case with either $d = i + 1$ or $j = e + 1$ but not both. Here, we will outline the functional form when $d = i + 1$. Let $n \equiv j - e - 1$ denote the number of unpaired nucleotides in the bulge loop.

$$\Delta G^{\text{bulge}}(\phi_{[i:i+1]}, \phi_{[e:j]}) = \Delta G^{\text{bulge}}_{\text{size}}(n) + \Delta G^{\text{bulge}}_{\text{seq}}(\phi_{[i:i+1]}, \phi_{[e:j]}) \tag{S81}$$

For the size-dependent term:

$$\Delta G^{\text{bulge}}_{\text{size}}(n) = \begin{cases} \Delta G^{\text{bulgesize}}_{n}, & n \leq 30 \\ \Delta G^{\text{bulgesize}}_{30} + \log\left(\frac{n}{30}\right) \Delta G^{\text{polymer}}_{\text{entropy}}, & n > 30 \end{cases} \tag{S82}$$

- $\Delta G^{\text{bulge size}}_{n}$: `rna95` uses empirical values for $1 \leq n \leq 5$.[7] `rna06` uses empirical values for $1 \leq n \leq 6$.[4,21] `dna04.1/dna04.2` use empirical values for a subset of $1 \leq n \leq 30$.[12] Each parameter set uses a logarithmic approximation for all other values of $n$. `merna06` uses `rna06` values.

For the sequence-dependent term:

$$\Delta G^{\text{bulge}}_{\text{seq}}(\phi_{[i:i+1]}, \phi_{[e:j]}) = \begin{cases} \Delta G^{\text{stack}}_{\phi_j,\phi_i,\phi_{i+1},\phi_e}, & e + 2 = j \\ \Delta G^{\text{terminalbp}}_{\phi_j,\phi_i} + \Delta G^{\text{terminalbp}}_{\phi_{i+1},\phi_e}, & \text{otherwise} \end{cases} \tag{S83}$$

**Other small interior loops.** The free energies for interior loops with $2 \leq d - i \leq 3$ and $2 \leq j - e \leq 3$ are kept in a lookup table.

- **1 × 1 interior loop.** Corresponds to $d - i = 2$ and $j - e = 2$. `rna95` assigns a sequence-independent $\Delta G$.[7] `rna06` uses unpublished parameters made available in the Mfold software.[9] `dna04.1` models these loops using (S84) below.[129] `dna04.2` models these loops using empirical values[12,23–27] with a positive constant free energy (10 kcal/mol) assigned for mismatches where the unpaired nucleotides are Watson–Crick complements. `merna06` uses `rna06` values treating RNA nucleotides as 2′OMe-RNA nucleotides.

- **1 × 2 interior loop.** Corresponds to $d-i = 2$ and $j-e = 3$, or $d-i = 3$ and $j-e = 2$. `rna95` and `dna04.1/dna04.2` model these loops using (S84) below.[7,12] For `dna04.1/dna04.2`, a positive constant free energy is assigned for mismatches where the unpaired nucleotides are Watson–Crick complements.[9] `rna06` models these loops using a combination of tabulated data and averaging.[4,21] `merna06` uses `rna06` values treating RNA nucleotides as 2′OMe-RNA nucleotides.

- **2 × 2 interior loop.** Corresponds to $d - i = 3$ and $j - e = 3$. `rna95` and `dna04.1/dna04.2` model these loops using (S84) below.[7,12] For `dna04.1/dna04.2`, a positive constant free energy is assigned for mismatches where the unpaired nucleotides are Watson–Crick complements.[9] `rna06` models these loops using tabulated symmetric tandem interior mismatches and averaging for asymmetric tandem interior mismatches.[4,21] `merna06` uses `rna06` values treating RNA nucleotides as 2′OMe-RNA nucleotides.

**Other interior loops.** Let $n_1 \equiv d-i-1$ and $n_2 \equiv j-e-1$ denote the number of unpaired nucleotides for the two sides of the interior loop. For the general case of interior loops not handled via special cases above, the following formula is used:

$$\Delta G^{\text{interior}}(\phi_{[i:d]}, \phi_{[e:j]}) = \Delta G^{\text{interior}}_{\text{size}}(n_1+n_2) + \Delta G^{\text{interior}}_{\text{asymm}}(n_1, n_2) + \Delta G^{\text{interior}}_{\text{mm}}(\phi_{[i:d]}, \phi_{[e:j]}) \tag{S84}$$

For the size-dependent term:

$$\Delta G^{\text{interior}}_{\text{size}}(n) = \begin{cases} \Delta G^{\text{interiorsize}}_{n}, & n \leq 30 \\ \Delta G^{\text{interiorsize}}_{30} + \log\left(\frac{n}{30}\right) \Delta G^{\text{polymer}}_{\text{entropy}}, & n > 30 \end{cases} \tag{S85}$$

- $\Delta G^{\text{interiorsize}}_{n}$: `rna95` uses empirical values for $2 \leq n \leq 6$.[7] `rna06` uses empirical values for $4 \leq n \leq 6$.[4,21] `dna04.1`/`dna04.2` use empirical values for a subset of values in $3 \leq n \leq 30$.[12] Each parameter set uses a logarithmic approximation for all other values of $n$. `merna06` uses `rna06` values.

For the asymmetry-based term:

$$\Delta G^{\text{interior}}_{\text{asymm}}(n_1, n_2) = \min(\Delta G^{\text{interiorasymm}}_{4}, |n_1 - n_2| \Delta G^{\text{interiorasymm}}_{\min(4, n_2, n_1)}) \tag{S86}$$

- $\Delta G^{\text{interiorasymm}}_{n}$: `rna95`,[7] `rna06`,[4,21] and `dna04.2`[12] use values regressed from empirical data. `dna04.1` uses values made available from the Mfold software.[9] `merna06` uses `rna06` values.

For the mismatch-based term:

$$\Delta G^{\text{interior}}_{\text{mm}}(\phi_{[i:d]}, \phi_{[e:j]}) = \begin{cases} \Delta G^{\text{interiormm}'}_{\phi_{j-1},\phi_j,\phi_i,\phi_{i+1}} + \Delta G^{\text{interiormm}'}_{\phi_{d-1},\phi_d,\phi_e,\phi_{e+1}} & i+2=d \text{ or } e+2=j \\ \Delta G^{\text{interiormm}}_{\phi_{j-1},\phi_j,\phi_i,\phi_{i+1}} + \Delta G^{\text{interiormm}}_{\phi_{d-1},\phi_d,\phi_e,\phi_{e+1}} & \text{otherwise} \end{cases} \tag{S87}$$

- $\Delta G^{\text{interiormm}}_{\phi_{j-1},\phi_j,\phi_i,\phi_{i+1}}$: `rna95`[7] and `rna06`[4,21] use independently determined values for loops without complementary unpaired bases. `dna04.1`/`dna04.2` equate $\Delta G^{\text{interiormm}}_{\phi_{j-1},\phi_j,\phi_i,\phi_{i+1}}$ with $\Delta G^{\text{terminalmm}}_{\phi_{j-1},\phi_j,\phi_i,\phi_{i+1}}$,[12] which are assigned unpublished values made available in the Mfold software.[9] `merna06` uses `rna06` values.

- $\Delta G^{\text{interiormm}'}_{\phi_{j-1},\phi_j,\phi_i,\phi_{i+1}}$: `rna95`,[7] `rna06`,[4,21] `dna04.1`/`dna04.2`[12] use different parameters for the case when one side of the interior loop has only one unpaired nucleotide.[29] `merna06` uses `rna06` values.

#### S3.8 Free energy model for single-material multiloops

A multiloop contains 3 or more terminal base pairs and no nicks. It may be defined as a series of bounding subsequences $[\phi]$. If the number of terminal base pairs is $n_{\text{bp}}$ and the number of unpaired nucleotides is $n_{\text{nt}}$, the free energy for a multiloop in a specified stacking state, $\omega$, is modeled as follows:

$$\begin{aligned} \Delta G^{\text{multi}}([\phi], \omega) = {} & \Delta G^{\text{multi}}_{\text{init}} + n_{\text{bp}} \Delta G^{\text{multi}}_{\text{bp}} + n_{\text{nt}} \Delta G^{\text{multi}}_{\text{nt}} \\ & + \Delta G^{\text{allterminalbp}}([\phi]) + \Delta G^{\text{allcoax}}([\phi], \omega) + \Delta G^{\text{alldangle}}([\phi], \omega) \end{aligned} \tag{S88}$$

where $\Delta G^{\text{multi}}_{\text{init}}$ denotes the penalty for formation of a multiloop, $\Delta G^{\text{multi}}_{\text{bp}}$ denotes the sequence-independent penalty for a terminal base pair in a multiloop, and $\Delta G^{\text{multi}}_{\text{nt}}$ denotes the penalty per unpaired nucleotide in a multiloop. Note that in contrast to interior loops and hairpin loops, the free energy of a multiloop is assumed to scale linearly, not logarithmically, with the number of unpaired nucleotides; the linear simplification facilitates the derivation of $O(N^3)$ multiloop recursions.

- $\Delta G^{\text{multi}}_{\text{init}}$: empirical values for `rna95`;[7] recently regressed empirical values for `rna06`;[1,31] unpublished values for `dna04.1`/`dna04.2`.[9] `merna06` uses `rna06` values.

- $\Delta G^{\text{multi}}_{\text{bp}}$: empirical values for `rna95`;[7] recently regressed empirical values for `rna06`;[1,31] unpublished values for `dna04.1`/`dna04.2`.[9] `merna06` uses `rna06` values.

- $\Delta G^{\text{multi}}_{\text{nt}}$: empirical values for `rna95`;[7] recently regressed empirical values for `rna06`;[1,31] unpublished values for `dna04.1`/`dna04.2`.[9] `merna06` uses `rna06` values.

The parameters $\Delta G^{\text{multi}}_{\text{init}}$, $\Delta G^{\text{multi}}_{\text{bp}}$, $\Delta G^{\text{multi}}_{\text{nt}}$ were regressed[1] from literature source data for multiloops.[31]

$\Delta G^{\text{allterminalbp}}([\phi])$ is a sum of the sequence-dependent free energy $\Delta G^{\text{terminalbp}}_{\phi_i,\phi_j}$ (defined in Section S4.5) for each terminal base pair $i \cdot j$ in the multiloop.

$\Delta G^{\text{allcoax}}([\phi],\omega)$ is a sum over each coaxial stack present in the multiloop stacking state $\omega$. Owing to a lack of parameters, only coaxial stacks between adjacent terminal base pairs (with no intervening unpaired bases) are considered. Each coaxial stack between base pairs $i \cdot d$ and $d+1 \cdot j$ contributes a free energy of $\Delta G^{\text{coax}}_{\phi_i,\phi_d,\phi_{d+1},\phi_j}$. For the recursions with coaxial stacking (Section S6.5), we use $\Delta G^{\text{coax}}_{i,d,j}(\phi)$ to denote $\Delta G^{\text{coax}}_{\phi_i,\phi_d,\phi_{d+1},\phi_j}$ since only three indices may vary freely. For the recursions without coaxial stacking (Section S6.3), the term $\Delta G^{\text{allcoax}}([\phi],\omega)$ is neglected.

- $\Delta G^{\text{coax}}_{\phi_i,\phi_d,\phi_{d+1},\phi_j}$: `rna95`[7] and `rna06`[4,21] set $\Delta G^{\text{coax}}_{\phi_i,\phi_d,\phi_{d+1},\phi_j}$ equal to $\Delta G^{\text{stack}}_{\phi_i,\phi_d,\phi_{d+1},\phi_j}$. `dna04.1/dna04.2` use independently estimated values.[5] `merna06` sets $\Delta G^{\text{coax}}_{\phi_i,\phi_d,\phi_{d+1},\phi_j}$ equal to the `rna_merna06` values (defined in Section S5.7) treating any RNA nucleotides as 2′OMe-RNA nucleotides.

$\Delta G^{\text{alldangle}}([\phi],\omega)$ is a sum of the sequence-dependent free energy, $\Delta G^{\text{dangle}}_{i,j}(\phi)$, for each terminal base pair $i \cdot j$ that is not in a coaxial stack in stacking state $\omega$. For a given terminal base pair $i \cdot j$, $\Delta G^{\text{dangle}}_{i,j}(\phi)$ takes one of four values to match the dangle stacking state for a given $\omega$:

$$\Delta G^{\text{dangle}}_{i,j}(\phi) = \begin{cases} 0 & \text{no dangles} \\ \Delta G^{5'\text{dangle}}_{\phi_i,\phi_{i+1},\phi_j} & 5' \text{ dangle} \\ \Delta G^{3'\text{dangle}}_{\phi_i,\phi_{j-1},\phi_j} & 3' \text{ dangle} \\ \Delta G^{\text{terminalmm}}_{\phi_i,\phi_{i+1},\phi_{j-1},\phi_j} & \text{terminal mismatch} \end{cases} \tag{S89}$$

Note that the state where both 5′ and 3′ dangles stack on terminal base pair $i \cdot j$ is classified as a terminal mismatch. For the recursions without dangle stacking (Section S6.3), the term $\Delta G^{\text{alldangle}}([\phi],\omega)$ is neglected.

- $\Delta G^{5'\text{dangle}}_{\phi_i,\phi_j,\phi_k}$: 5′ dangle free energy parametrized for `rna95`,[7] `rna06`,[4,21] and `dna04.1/dna04.2`.[12] `merna06` uses `rna06` values treating RNA nucleotides as 2′OMe-RNA nucleotides.
- $\Delta G^{3'\text{dangle}}_{\phi_i,\phi_j,\phi_k}$: 3′ dangle free energy parametrized for `rna95`,[7] `rna06`,[4,21] and `dna04.1/dna04.2`.[12] `merna06` uses `rna06` values treating RNA nucleotides as 2′OMe-RNA nucleotides.
- $\Delta G^{\text{terminalmm}}_{\phi_i,\phi_{i+1},\phi_{j-1},\phi_j}$: `rna95`,[7] `rna06`,[4,21] and `dna04.2`[6,28] use empirical parameters for $\Delta G^{\text{terminalmm}}$. `dna04.1` assigns $\Delta G^{\text{terminalmm}}_{\phi_i,\phi_{i+1},\phi_{j-1},\phi_j}$ to be the sum of $\Delta G^{5'\text{dangle}}_{\phi_i,\phi_{i+1},\phi_j}$ and $\Delta G^{3'\text{dangle}}_{\phi_i,\phi_{j-1},\phi_j}$. `merna06` uses `rna06` values treating RNA nucleotides as 2′OMe-RNA nucleotides.

##### S3.9 Free energy model for single-material exterior loops

An exterior loop is a loop containing one nick and zero or more terminal base pairs. An exterior loop may be defined as a series of bounding subsequences $[\phi]$ with a given nick location. An unpaired strand is an exterior loop with a free energy of zero, corresponding to the reference state.[2] The free energy of an exterior loop in a specified stacking state $\omega$ is modeled as follows:

$$\Delta G^{\text{exterior}}([\phi],\omega) = 0 + \Delta G^{\text{allterminalbp}}([\phi]) + \Delta G^{\text{allcoax}}([\phi],\omega) + \Delta G^{\text{alldangle}}([\phi],\omega). \tag{S90}$$

The functions $\Delta G^{\text{allterminalbp}}([\phi])$, $\Delta G^{\text{allcoax}}([\phi],\omega)$, and $\Delta G^{\text{alldangle}}([\phi],\omega)$ are defined in Section S3.8 as for multiloops. For the recursions without coaxial and dangle stacking (Section S6.3), the terms $\Delta G^{\text{allcoax}}([\phi],\omega)$ and $\Delta G^{\text{alldangle}}([\phi],\omega)$ are neglected.

#### S4 Mixed-material free energy model details: RNA/DNA

The functional form of the free energy model for mixed-material systems is the same as for single-material systems (see Section S3). Here, we describe mixed-material RNA/DNA parameters for each term in the model, taking advantage of empirically derived parameters where available and averaging single-material parameters where they are not.

#### S4.1 Mixed-material parameter set: RNA/DNA

A mixed-material parameter set contains parameters for mixed-material loops and also specifies the names of two single-material parameter sets with which it is used in conjunction. The following temperature-dependent parameter set is supported:

- `rna_dna06`: RNA/DNA parameters for hybrid stack loops,[32] hybrid internal mismatches,[33,34] and chimeric stack loops[35] with additional parameters for mixed-material loops as defined below; for use in conjunction with single-material parameter sets `rna06` and `dna04.2`.

#### S4.2 Definition of mixed-material Watson–Crick and wobble pairs: RNA/DNA

Mixed-material base pairs are either Watson–Crick pairs:

- RNA/DNA: rG·dC, dG·rC, rA·dT, dA·rU,

or wobble pairs:

- RNA/DNA: rG·dT,

Note that for RNA/DNA, dG and rU are considered to form a mismatch dG×rU and not a wobble pair.[34]

#### S4.3 Mixed-material salt conditions: RNA/DNA

The default salt condition for our RNA/DNA[32–35] parameter set is [$Na^+$] = 1.0 M. For a mixed-material RNA/DNA secondary structure $s$ containing $n_{\text{bp}}$ base pairs, the free energy, $\Delta G(\phi, s)$, is augmented by (see Section S2.4):

$$n_{\text{bp}}(s)\Delta G^{\text{salt}} \tag{S91}$$

with

$$\Delta G^{\text{salt}} = -0.114 \log\left([\text{Na}^+]\right)\frac{T}{T_{37}} \tag{S92}$$

for user-specified

$$0.12\,\text{M} \leq [\text{Na}^+] \leq 1.0\,\text{M}, \tag{S93}$$

with $T_{37} = 310.15$ K.

#### S4.4 Mixed-material strand association penalty: RNA/DNA

A $\Delta G^{\text{assoc}}_{\text{pub, hybrid(RNA:DNA)}}$ association penalty derived from RNA/DNA hybrid duplexes is available in the literature.[32] Because `rna06` and `dna04.2` parameters were derived with different association penalties ($\Delta G^{\text{assoc}}_{\text{pub, RNA}}$ and $\Delta G^{\text{assoc}}_{\text{pub, DNA}}$ ), a weighted average, $\Delta G^{\text{assoc}}_{\text{pub, RNA/DNA}}$, between the `rna06` and `dna04.2` association penalties is used for mixed RNA/DNA complexes, centered at $\Delta G^{\text{assoc}}_{\text{pub, hybrid}}$. Let $p_{\text{RNA}}$ and $p_{\text{DNA}}$ be the proportions of RNA and DNA nucleotides in the complex, respectively. Then

$$\Delta G^{\text{assoc}}_{\text{pub,RNA/DNA}} \equiv \Delta G^{\text{assoc}}_{\text{pub,hybrid(RNA:DNA)}} + (2p_\sigma - 1)(\Delta G^{\text{assoc}}_{\text{pub},\sigma} - \Delta G^{\text{assoc}}_{\text{pub,hybrid(RNA:DNA)}}) \tag{S94}$$

$$\sigma = \begin{cases} \text{RNA}, & p_{\text{RNA}} \geq 0.5 \\ \text{DNA}, & \text{otherwise} \end{cases} \tag{S95}$$

which reduces to $\Delta G^{\text{assoc}}_{\text{pub, hybrid(RNA:DNA)}}$ when $p_{\text{RNA}} = p_{\text{DNA}} = 0.5$.

#### S4.5 Free energy model for mixed-material hairpin loops: RNA/DNA

A hairpin loop is defined for a subsequence $\phi_{[i:j]}$ by the single base pair $i \cdot j$ such that there are no nicks or additional base pairs in the range $[i : j]$. Let $n \equiv j - i - 1$ denote the number of unpaired nucleotides in the hairpin loop. Steric effects are assumed to prevent hairpin loops with $n < 3$ as for RNA and DNA. The functional form of the hairpin free energy is as follows:

$$\Delta G^{\text{hairpin}}(\phi_{[i:j]}) = \Delta G^{\text{hairpin}}_{\text{size}}(n) + \Delta G^{\text{hairpin}}_{\text{seq}}(\phi_{[i:j]}). \tag{S96}$$

For the size-dependent term:

$$\Delta G^{\text{hairpin}}_{\text{size}}(n) = \begin{cases} \infty, & n < 3 \\ \Delta G^{\text{hairpinsize}}_{n}, & 3 \leq n \leq 30 \\ \Delta G^{\text{hairpinsize}}_{30} + \log\left(\frac{n}{30}\right) \Delta G^{\text{polymer}}_{\text{entropy}}, & n > 30 \end{cases} \tag{S97}$$

- $\Delta G^{\text{hairpinsize}}_{n}$: a lookup table with empirical and log-extrapolated values for $3 \leq n \leq 30$. The mixed-material model uses a simple average between the `rna06` and `dna04.2` parameters.
- $\Delta G^{\text{polymer}}_{\text{entropy}}$: a logarithmic extrapolation parameter based on Jacobson-Stockmayer polymer theory for $n > 30$. The mixed-material model uses a simple average between the `rna06` and `dna04.2` parameters.

For the sequence-dependent term:

$$\Delta G^{\text{hairpin}}_{\text{seq}}(\phi_{[i:j]}) = \begin{cases} \Delta G^{\text{triloop}}_{\phi_{[i:j]}} + \Delta G^{\text{terminalbp}}_{\phi_j,\phi_i} & n = 3 \\ \Delta G^{\text{tetraloop}}_{\phi_{[i:j]}} & n = 4 \\ \Delta G^{\text{hairpinmm}}_{\phi_{j-1},\phi_j,\phi_i,\phi_{i+1}} & n \geq 5 \end{cases} \tag{S98}$$

- $\Delta G^{\text{triloop}}_{\phi_{[i:j]}}$: an empirical sequence-dependent penalty for a hairpin loop of length $n = 3$. `rna06` and `dna04.2` contain non-overlapping sets of 2 and 16 triloop parameters (each for a different sequence), respectively. The mixed-material model uses the `rna06` or `dna04.2` parameter that matches the sequences of bases of the query, regardless of material. Parameters are stored as a set of 576 mixed-material triloop parameters (18 sequences with each of 5 nucleotides [3 in the loop and 2 in the closing base pair] being either RNA or DNA).
- $\Delta G^{\text{terminalbp}}_{\phi_j,\phi_i}$: an empirical penalty for non-GC terminal base pairs at the end of a duplex. In the mixed-material model, all-RNA terminal base pairs use the `rna06` value and all-DNA terminal base pairs use the `dna04.2` value. If the terminal base pair contains both RNA and DNA, the free energy and enthalpy are set to 0 because the hybrid and chimeric stack values were derived without a terminal base pair penalty.[32,35]
- $\Delta G^{\text{tetraloop}}_{\phi_{[i:j]}}$: a sequence-dependent empirical penalty for hairpins of length $n = 4$. The mixed-material model uses a weighted average if the sequence of bases is in both `rna06` and `dna04.2`:

$$\left(\frac{m_{\text{RNA}}}{6}\right) \Delta G^{\text{tetraloop,RNA}}_{\phi_{[i:j]}} + \left(\frac{m_{\text{DNA}}}{6}\right) \Delta G^{\text{tetraloop,DNA}}_{\phi_{[i:j]}}, \tag{S99}$$

  where $m_{\text{RNA}}$ and $m_{\text{DNA}}$ are the number of RNA and DNA bases in the 6-base input sequence (4 bases in the loop and 2 in the closing base pair). If the input sequence is only in `rna06` or `dna04.2`, then the available value is used regardless of material.
- $\Delta G^{\text{hairpinmm}}_{\phi_{j-1},\phi_j,\phi_i,\phi_{i+1}}$: a sequence-dependent empirical value for mismatched bases adjacent to base pair $i \cdot j$ in a hairpin loop. The mixed-material model uses a weighted average between the `rna06` and `dna04.2` hairpin mismatch with the same sequence:

$$\left(\frac{m_{\text{RNA}}}{4}\right) \Delta G^{\text{hairpinmm,RNA}}_{\phi_{j-1},\phi_j,\phi_i,\phi_{i+1}} + \left(\frac{m_{\text{DNA}}}{4}\right) \Delta G^{\text{hairpinmm,DNA}}_{\phi_{j-1},\phi_j,\phi_i,\phi_{i+1}}, \tag{S100}$$

  where $m_{\text{RNA}}$ and $m_{\text{DNA}}$ are the number of RNA and DNA bases in the 4-base input sequence (2 bases in the mismatch and 2 in the closing base pair).

#### S4.6 Free energy model for mixed-material interior loops: RNA/DNA

An interior loop may be defined via a pair of subsequences $\phi_{[i:d]}$ and $\phi_{[e:j]}$ such that $i < d < e < j$ with base pairs $i \cdot j$ and $d \cdot e$, with no additional paired bases or nicks within the two subsequences.

**Stack loops.** Stack loops are the special case where $d = i + 1$ and $j = e + 1$. The stack loop energy model is as follows:

$$\Delta G^{\text{stackedpair}}(\phi_{[i:i+1]}, \phi_{[j-1:j]}) = \Delta G^{\text{stack}}_{\phi_j,\phi_i,\phi_{i+1},\phi_{j-1}} \tag{S101}$$

- $\Delta G^{\text{stack}}_{\phi_j,\phi_i,\phi_{i+1},\phi_{j-1}}$: empirical values for stack free energies.

- For the mixed-material RNA/DNA model, stack energies are available for 16 hybrid stacks (rr/dd) not including wobble pairs.[32]
- Stack energies are also available for 31 chimeric stacks (rd/rr, rd/dd, dr/rr, dr/rd, dr/dr, and rd/rd).[35] These were rederived here since the derivation in the original paper used RNA and DNA stack energies and strand association penalties from Freier et al.[36] and Sugimoto et al.,[37] respectively, and did not use terminal base-pair penalties (for non-GC terminal base pairs at the end of a duplex); by contrast, `rna06` and `dna04.2` use stack energies, strand association penalties, and terminal base-pair penalties from Xia et al.[20] and SantaLucia Jr.,[11] respectively. The chimeric stack energies were rederived as:

$$\Delta G^{\text{stack,chimeric}}_{\phi_j,\phi_i,\phi_{i+1},\phi_{j-1}} = \Delta G^{\text{duplex}}_{\phi} - \sum_{d,e\neq i,j} \Delta G^{\text{stack,nonchimeric}}_{\phi_e,\phi_d,\phi_{d+1},\phi_{e-1}} - \Delta G^{\text{assoc}}_{\text{pub, RNA/DNA}} - \sum \Delta G^{\text{terminalbp}} - \Delta G^{\text{sym}}(\phi, s) \tag{S102}$$

  where $d$ and $e$ index all stack loops in the duplex other than the chimeric stack, $\Delta G^{\text{duplex}}_{\phi}$ is the observed free energy of the duplex melting, $\Delta G^{\text{terminalbp}}$ is the penalty for any non-GC pair at either end of the duplex, $\Delta G^{\text{assoc}}_{\text{pub, RNA/DNA}}$ is the weighted strand association penalty of equation (S94), and $\Delta G^{\text{sym}}(\phi, s)$ is the symmetry correction of equation (3). For non-chimeric stacks indexed by $d$ and $e$, hybrid stack parameters were taken from Sugimoto et al.[32] and single-material stack parameters were taken from Xia et al.[20] for RNA and SantaLucia Jr.[11] for DNA. Enthalpy parameters were derived in the same way.
- It is noted in the literature that within a stack loop, each chimeric strand in a stack behaves as if the strand is composed solely of the material of the 5′-most base[35] (i.e., an rd/dr stack exhibits similar energetics to an rr/dd stack). An exception to this rule is dr/rr stacks, which are closest to rr/rr stacks rather than dd/rr stacks. This rule is used to fill in the rest of the chimeric stack parameters in the mixed-material model, with the exception of dr/rr stacks which are equated to rr/rr stacks.
- As noted in Section S4.2, rG is considered to form a wobble pair with dT. Melts with duplexes containing internal rG·dT wobble pairs have been reported in literature but parameters were not provided.[34] Here, we derive rG·dT stack parameters from the literature melts using linear regression in the same way interior mismatch parameters were derived for $1\times1$ interior loops in the literature.[33] For each duplex with an internal rG·dT wobble pair, a core duplex without the internal rG·dT pair was melted. Using the nearest-neighbor model, the free energy of the three-nucleotide sequence containing the rG·dT pair and its two flanking pairs ($\Delta G^{\text{trimer}}_{\text{rG}\cdot\text{dT}}$) is derived as follows:

$$\Delta G^{\text{trimer}}_{\text{rG}\cdot\text{dT}} = \Delta G^{\text{rG}\cdot\text{dT duplex}}_{\phi} - \Delta G^{\text{core duplex}} + \Delta G^{\text{stack}}_{\text{closingbp}} \tag{S103}$$

  where $\Delta G^{\text{rG}\cdot\text{dT duplex}}_{\phi}$ is the observed melt free energy of the full duplex with an internal rG·dT wobble pair, $\Delta G^{\text{core duplex}}$ is the observed melt free energy of the core duplex without the rG·dT pair, and $\Delta G^{\text{stack}}_{\text{closingbp}}$ is the free energy of the stack loop composed of the base pairs flanking the rG·dT pair that is broken in the rG·dT duplex (e.g., for the rG·dT trimer rCGC/dGTG, $\Delta G^{\text{stack}}_{\text{closingbp}}$ is $\Delta G^{\text{stack}}_{\text{rCC/dGG}}$). $\Delta G^{\text{rG}\cdot\text{dT}}_{\text{trimer}}$ is further broken down into two stack energies each containing an rG·dT wobble pair. For example:

$$\Delta G^{\text{trimer}}_{\text{rCGC/dGTG}} = \Delta G^{\text{stack}}_{\text{rCG/dTG}} + \Delta G^{\text{stack}}_{\text{rGC/dGT}} \tag{S104}$$

  After $\Delta G^{\text{trimer}}_{\text{rG}\cdot\text{dT}}$ is calculated for all of the melts, the rG·dT stack energies can be solved for by linear regression using singular value decomposition (SVD). Because these melts only included internal rG·dT wobble pairs and not terminal rG·dT wobble pairs, linear regression is done on a rank-deficient matrix and the SVD returns the non-unique solution with the minimum 2-norm.[33] In turn, results from using these stack parameters on sequences with terminal rG·dT wobble pairs are more susceptible to inaccuracy due to non-uniqueness. Enthalpies were derived in the same manner.
- The hybrid stack rGG/dTT comprising two stacked wobble pairs was not measured in the above rG·dT melts. This value was estimated by approximating the decrement in free energy in going from a Watson–Crick hybrid base pair to a rG·dT pair. Specifically, for each of the eight measured $\Delta G^{\text{trimer}}_{\text{rG}\cdot\text{dT}}$ values, the difference in free energy was calculated from $\Delta G^{\text{trimer}}_{\text{rA}\cdot\text{dT}}$, $\Delta G^{\text{trimer}}_{\text{rU}\cdot\text{dA}}$, $\Delta G^{\text{trimer}}_{\text{rC}\cdot\text{dG}}$, and $\Delta G^{\text{trimer}}_{\text{rG}\cdot\text{dC}}$. For example, for $\Delta G^{\text{trimer}}_{\text{rCGC/dGTG}}$ the following were calculated: $\Delta G^{\text{trimer}}_{\text{rCGC/dGTG}} - \Delta G^{\text{trimer}}_{\text{rCAC/dGTG}}$, $\Delta G^{\text{trimer}}_{\text{rCGC/dGTG}} - \Delta G^{\text{trimer}}_{\text{rCUC/dGAG}}$, $\Delta G^{\text{trimer}}_{\text{rCGC/dGTG}} - \Delta G^{\text{trimer}}_{\text{rCCC/dGGG}}$, and $\Delta G^{\text{trimer}}_{\text{rCGC/dGTG}} - \Delta G^{\text{trimer}}_{\text{rCGC/dGCG}}$. $\Delta G^{\text{trimer}}_{\text{rA}\cdot\text{dT}}$, $\Delta G^{\text{trimer}}_{\text{rU}\cdot\text{dA}}$, $\Delta G^{\text{trimer}}_{\text{rC}\cdot\text{dG}}$, and $\Delta G^{\text{trimer}}_{\text{rG}\cdot\text{dC}}$ were calculated by summing published hybrid stack energies from Sugimoto et al.[32] The differences represent the decrement in free energy from switching from two Watson–Crick hybrid stacks to two stacks involving wobble pair rG·dT. For example, trimer free energies can be broken down into two stack energies as follows:

$$\Delta G^{\text{trimer}}_{\text{rCGC/dGCG}} = \Delta G^{\text{stack}}_{\text{rCG/dCG}} + \Delta G^{\text{stack}}_{\text{rGC/dGC}} \tag{S105}$$

$$\Delta G^{\text{stack}}_{\text{rCG/dCG}} + \Delta G^{\text{stack}}_{\text{rGC/dGC}} + (\Delta G^{\text{trimer}}_{\text{rCGC/dGTG}} - \Delta G^{\text{trimer}}_{\text{rCGC/dGCG}}) = \Delta G^{\text{trimer}}_{\text{rCGC/dGTG}} = \Delta G^{\text{stack}}_{\text{rCG/dTG}} + \Delta G^{\text{stack}}_{\text{rGC/dGT}} \tag{S106}$$

If we assume an equal free energy decrement to each Watson–Crick stack, then

$$(\Delta G^{\text{trimer}}_{\text{rCGC/dGTG}} - \Delta G^{\text{trimer}}_{\text{rCGC/dGCG}})/2 \tag{S107}$$

approximates the decrement to one Watson–Crick stack value when switching a rG·dC base pair to rG·dT. $\Delta G^{\text{trimer}}_{\text{rG·dT}} - \Delta G^{\text{trimer}}_{\text{rA·dT}}$, $\Delta G^{\text{trimer}}_{\text{rG·dT}} - \Delta G^{\text{trimer}}_{\text{rU·dA}}$, $\Delta G^{\text{trimer}}_{\text{rG·dT}} - \Delta G^{\text{trimer}}_{\text{rC·dG}}$, and $\Delta G^{\text{trimer}}_{\text{rG·dT}} - \Delta G^{\text{trimer}}_{\text{rG·dC}}$ were calculated for each of the eight measured rG·dT trimers, and averaged across all eight to get $\Delta G^{\text{dec}}_{\text{rA·dT}}$, $\Delta G^{\text{dec}}_{\text{rU·dA}}$, $\Delta G^{\text{dec}}_{\text{rC·dG}}$, and $\Delta G^{\text{dec}}_{\text{rG·dC}}$. These averaged decrements were then applied to each of the eight experimentally derived rG·dT stacks to get eight rGG/dTT estimates. For example, with the stack rCG/TG:

$$\Delta G^{\text{stack}}_{\text{rGG/dTT}} \approx \Delta G^{\text{stack}}_{\text{rCG/dTG}} + \Delta G^{\text{dec}}_{\text{rC·dG}} \tag{S108}$$

These 8 rGG/dTT estimates were averaged to get the final rGG/dTT estimate. The same was done for enthalpy.

- Estimating any missing chimeric stacks comprising a rG·dT and/or rG·rU wobble pair was done using the chimeric stack rule as discussed above, including the exception that dr/rr stacks were equated to the rr/rr value. If the chimeric strand rule resulted in a dG×dT or dG×rU mismatch, a positive constant free energy was used for the stack free energy and enthalpy to discourage base pairing, since dG is not considered to form a wobble pair with dT or rU.

**Bulge loops.** A bulge loop is the special case with either $d = i + 1$ or $j = e + 1$ but not both. Here, we will outline the functional form when $d = i + 1$. Let $n \equiv j - e - 1$ denote the number of unpaired nucleotides in the bulge loop. The bulge loop free energy is modeled as follows:

$$\Delta G^{\text{bulge}}(\phi_{[i:i+1]}, \phi_{[e:j]}) = \Delta G^{\text{bulge}}_{\text{size}}(n) + \Delta G^{\text{bulge}}_{\text{seq}}(\phi_{[i:i+1]}, \phi_{[e:j]}). \tag{S109}$$

The size-dependent term is:

$$\Delta G^{\text{bulge}}_{\text{size}}(n) = \begin{cases} \Delta G^{\text{bulgesize}}_{n}, & n \leq 30 \\ \Delta G^{\text{bulgesize}}_{30} + \log\left(\frac{n}{30}\right) \Delta G^{\text{polymer}}_{\text{entropy}}, & n > 30 \end{cases} \tag{S110}$$

- $\Delta G^{\text{bulge size}}_{n}$: empirical values with logarithmic extrapolation to cover the range $1 \leq n \leq 30$. The mixed-material model uses a simple average of the `rna06` and `dna04.2` values.

The sequence-dependent term:

$$\Delta G^{\text{bulge}}_{\text{seq}}(\phi_{[i:i+1]}, \phi_{[e:j]}) = \begin{cases} \Delta G^{\text{stack}}_{\phi_j,\phi_i,\phi_{i+1},\phi_e}, & e + 2 = j \\ \Delta G^{\text{terminalbp}}_{\phi_j,\phi_i} + \Delta G^{\text{terminalbp}}_{\phi_{i+1},\phi_e}, & \text{otherwise} \end{cases} \tag{S111}$$

remains a look-up table in the mixed-material model, with sequence indicating both material and base (e.g., $\phi_i$ = dT or $\phi_i$ = rA).

**Other small interior loops.**

- **1 × 1 interior loop.** Corresponds to $d - i = 2$ and $j - e = 2$. The free energies for 1×1 interior loops (also referred to as internal mismatches) are kept in a lookup table. Hybrid 1×1 interior loops were measured in Sugimoto et al.[34] (rU×dG, rG×dG, rA×dG, rG×dA, rA×dC, rA×dA, rU×dT, rU×dC, rC×dA, and rC×dT mismatches) and Watkins et al.[33] (rA×dA, rC×dC, rG×dG, and rU×dT mismatches). To ensure compatibility with the hybrid stack parameters of Sugimoto et al.,[32] 1×1 interior loop energies ($\Delta G^{1\times1}$) were rederived in the same way that $\Delta G^{\text{trimer}}_{\text{rG·dT}}$ was derived in equation (S103) as outlined in Watkins et al.[33] For each duplex with a 1×1 interior loop, a core duplex without the 1×1 interior loop was melted. Using the nearest-neighbor model, the free energy of the 1×1 interior loop can be derived as follows:

$$\Delta G^{1\times1} = \Delta G^{1\times1\text{ duplex}}_{\phi} - \Delta G^{\text{core duplex}} + \Delta G^{\text{stack}}_{\text{closingbp}} \tag{S112}$$

where $\Delta G^{1\times1}$ is the observed melt free energy of the full duplex with a 1×1 loop, $\Delta G^{\text{core duplex}}$ is the observed melt free energy of the core duplex without the 1×1 loop, and $\Delta G^{\text{stack}}_{\text{closingbp}}$ is the free energy of the stack loop composed of the base pairs closing the 1×1 loop that is broken by the 1×1 loop (e.g., for the 1×1 loop rCGC/dGGG, $\Delta G^{\text{stack}}_{\text{closingbp}}$ is $\Delta G^{\text{stack}}_{\text{rCC/dGG}}$). $\Delta G^{1\times1}$ is further broken down into two interior mismatch energies, similar to Eq. S104. For example:

$$\Delta G^{1\times1}_{\text{rCGC/dGGG}} = \Delta G^{\text{interior}}_{\text{mm, rCG/dGG}} + \Delta G^{\text{interior}}_{\text{mm, rGC/dGG}} \tag{S113}$$

After $\Delta G^{1\times1}$ is calculated for all of the melts for a particular mismatch (e.g. all rG×dG 1×1 loops), the $\Delta G_{\text{mm}}^{\text{interior}}$ energies for this mismatch can be solved for by linear regression using SVD. Because these melts only include internal and not terminal mismatches, linear regression is done on a rank-deficient matrix and the SVD returns the non-unique solution for $\Delta G_{\text{mm}}^{\text{interior}}$ with the minimum 2-norm.[33] The derivation of parameters for some mismatch types (rA×dA, rG×dG, and rU×dT) used sequences from both Watkins et al.[33] and Sugimoto et al.[32] Each mismatch type had sixteen possible 1×1 loops closed by Watson–Crick base pairs, although no mismatch type had all sixteen measured. For the missing hybrid 1×1 loops closed by Watson–Crick base pairs, the following strategies were used:

- For mismatch types (rA×dA, rC×dC, rG×dG, rU×dT, and rU×dG) that had enough duplexes to derive all eight possible $\Delta G_{\text{mm}}^{\text{interior}}$ energies, the missing $\Delta G^{1\times1}$ values for 1×1 loops that were not profiled experimentally were derived by summing the appropriate $\Delta G_{\text{mm}}^{\text{interior}}$ energies. The enthalpies were estimated in the same way. Below, we refer to this set of five mismatches as the "five-set".
- For mismatch types (rA×dC, rA×dG, rG×dA, and rU×dC) that featured all four 1×1 loops closed by rC·dG or rG·dC at each end, the missing $\Delta G^{1\times1}$ values for 1×1 loops containing at least one rA·dT or rU·dA closing base pair were estimated in the following way. The five-set mismatches were used to estimate free energy decrements for changing a loop closed solely by rC·dG or rG·dC base pairs to one with 1 or 2 rA·dT or rU·dA closing base pairs. For example, the loop closed by 5′-rA·dT-3′ and 3′-rC·dG-5′ can be derived by either changing the 5′-rC·dG-3′ base pair in the loop closed by 5′-rC·dG-3′ and 3′-rC·dG-5′ to 5′-rA·dT-3′, or changing the 5′-rG·dC-3′ base pair in the loop closed by 5′-rG·dC-3′ and 3′-rC·dG-5′ to 5′-rA·dT-3′. Therefore, the decrement for a 1×1 loop closed by 5′-rA·dT-3′ and 3′-rC·dG-5′ is estimated for a particular mismatch type by averaging the free energies of the loop closed by 5′-rC·dG-3′ and 3′-rC·dG-5′ and the loop closed by 5′-rG·dC-3′ and 3′-rC·dG-5′, and subtracting the average from the free energy of the loop closed by 5′-rA·dT-3′ and 3′-rC·dG-5′. This decrement was calculated for every mismatch type in the five-set (rA×dA, rC×dC, rG×dG, rU×dT, and rU×dG) and averaged to yield $\Delta G^{\text{dec}}(5'\text{-rA·dT-}3', 3'\text{-rC·dG-}5')$. These averaged decrements were then used to estimate the missing 1×1 loops with at least one rA·dT or rU·dA closing base pair. For example:

  $$\Delta G^{1\times1}_{\text{rAAC/dGCT}} = (\Delta G^{1\times1}_{\text{rCAC/dGCG}} + \Delta G^{1\times1}_{\text{rGAC/dGCC}})/2 + \Delta G^{\text{dec}}(5'\text{-rA·dT-}3', 3'\text{-rC·dG-}5') \quad \text{(S114)}$$

  For decrements with 2 rA·dT or rU·dA closing base pairs, all four 1×1 loops closed by rC·dG or rG·dC base pairs were used in the average. For example, in calculating the decrement for the loop closed by 5′-rA·dT-3′ and 3′-rA·dT-5′, the loops closed by 5′-rC·dG-3′ and 3′-rC·dG-5′, 5′-rG·dC-3′ and 3′-rC·dG-5′, 5′-rC·dG-3′ and 3′-rG·dC-5′, and 5′-rG·dC-3′ and 3′-rG·dC-5′ were averaged and subtracted from the loop closed by 5′-rA·dT-3′ and 3′-rA·dT-5′. The enthalpies were estimated in the same way.
- The rC×dA mismatch type only has two 1×1 loops that were profiled: rCCG/dCAG and rGCG/dCAC. All mismatch types which had melt data on all four 1×1 loops closed by rC·dG or rG·dC were used to determine the average difference in stability between the loops closed by 5′-rC·dG-3′ and 5′-rG·dC-3′ and the two missing loops closed by rC·dG or rG·dC for rC·dA. This average difference in stability was applied to the rCCG/dCAG free energy to get one estimate for the missing loops closed by rC·dG or rG·dC. The same procedure was done using the loops closed by 5′-rG·dC-3′ and 5′-rG·dC-3′ and the empirical rGCG/dCAC free energy value to get a second estimate. These estimates were averaged to get the final estimate. For the missing 1×1 loops with at least one rA·dT or rU·dA closing base pair, the same procedure was applied, except that only the five-set was used to generate the average differences in stability. Overall this amounts to the same calculation done in the section above for mismatch types that featured all four 1×1 loops, except that only the loops closed by 5′-rC·dG-3′ and 5′-rG·dC-3′, as well as by 5′-rG·dC-3′ and 5′-rG·dC-3′, were used in the averages used to calculate decrements for missing loops since only these two loops were available for rC·dA. The enthalpies were estimated in the same way.
- The rC×dT mismatch type only has one 1×1 loop that was profiled: rCCG/dCTG. The five-set mismatches, as well as any mismatch type that had melt data for both the loop closed by 5′-rC·dG-3′ and 5′-rG·dC-3′ and the loop to be estimated, were used to calculate the average difference in stability between the loop closed by 5′-rC·dG-3′ and 5′-rG·dC-3′ and the loop to be estimated. This average difference in stability was applied to the rCCG/dCTG free energy value to generate an estimate. The enthalpies were estimated in the same way.

For the missing hybrid 1×1 loops closed by at least one rG·dT wobble pair, the following strategy was used:

- The decrements derived above to approximate the rGG/dTT stack ($\Delta G^{\text{dec}}_{\text{rA·dT}}$, $\Delta G^{\text{dec}}_{\text{rU·dA}}$, $\Delta G^{\text{dec}}_{\text{rC·dG}}$, and $\Delta G^{\text{dec}}_{\text{rG·dC}}$) were used to estimate 1×1 loops closed by rG·dT wobble pairs. 1×1 loops with one rG·dT wobble pair and one Watson–Crick pair were estimated by applying the appropriate decrement to a 1×1 loop of the same mismatch type with the rG·dT wobble pair replaced by a Watson–Crick pair. For example:

  $$\Delta G^{1\times1}_{\text{rGAG/dCAT}} \approx \Delta G^{1\times1}_{\text{GAG/dCAC}} + \Delta G^{\text{dec}}_{\text{rG·dC}}/2 \quad \text{(S115)}$$

This was done for the 1×1 loop with rG·dT replaced with rG·dC, rC·dG, rA·dT, and rU·dA, and the results from each were averaged to get the final estimate. For 1×1 loops closed by two rG·dT pairs, all sixteen 1×1 loops of the same mismatch type closed by two Watson–Crick base pairs were used to derive an estimate by applying two decrements to each, and the results were averaged to get the final estimate. The enthalpies were estimated in the same way.

For the missing chimeric 1×1 loops, the following strategies were used:

- The chimeric 1×1 mismatch dT×dG was profiled in Sugimoto et al[34] for the loops closed by 5′-rG·dC-3′ and 3′-rG·dC-5′, 5′-rG·dC-3′ and 3′-rC·dG-5′, 5′-rC·dG-3′ and 3′-rG·dC-5′, and 5′-rC·dG-3′ and 3′-rC·dG-5′ (the same closing base pairs profiled for rA×dC, rA×dG, rG×dA, and rU×dC hybrid 1×1 loops). The differences in free energy between the four measured 1×1 loops for dT×dG closely resembled the differences in free energy between rU×dG loops with the same closing base pairs. Therefore, the 12 rU×dG hybrid loops with closing base pairs that were not profiled for dT×dG were used to estimate those missing dT×dG loops. This includes a combination of experimentally profiled rU×dG hybrid loops as well as loops estimated with the above methods. Specifically, for any of the 12 rU×dG loops that were between the stabilities of two rU×dG loops closed by rC·dG or rG·dC that were also profiled for dT×dG, the dT×dG loop with the same closing base pairs was estimated by taking the average between the analogous dT×dG loops closed by rC·dG or rG·dC. The least stable loop closed by rC·dG or rG·dC that was experimentally profiled for dT×dG and rU×dG was the loop closed by 5′-rC·dG-3′ and 3′-rG·dC-5′. The gaps in free energy between the rU×dG loop closed by 5′-rC·dG-3′ and 3′-rG·dC-5′ and any of the hybrid rU×dG loops that were lower in stability were calculated. These gaps in free energy were then applied to the dT×dG loop closed by 5′-rC·dG-3′ and 3′-rG·dC-5′ to estimate the missing dT×dG loops closed by the same base pairs, so that the differences in free energy for the dT×dG loops matched the hybrid rU×dG loops. The enthalpies were estimated in the same way.
- Any other missing chimeric 1×1 loop was estimated by taking a weighted average between the `rna06` and `dna04.2` free energy values. The same approach was used for enthalpies.

- **1 × 2 interior loop and 2 × 2 interior loops.** For single-material models, these loops empirical parameters are provided in a lookup table. Due to the large storage that would be required for a mixed-material lookup table , the mixed-material model instead uses the same equation used for larger interior loops (see below). For example, 2 × 2 interior loops would require a lookup table with 1,638,400 parameters for the mixed-material case (20 possible identities for each of 2 closing base pairs, 8 possible identities for each of 4 unpaired nucleotides).

**Other large interior loops.** Let $n_1 \equiv d - i - 1$ and $n_2 \equiv j - e - 1$ denote the number of unpaired nucleotides for the two sides of the interior loop. The following formula is used for all other mixed-material interior loops that are not stack loops, bulge loops, or 1×1 interior loops:

$$\Delta G^{\text{interior}}(\phi_{[i:d]}, \phi_{[e:j]}) = \Delta G^{\text{interior}}_{\text{size}}(n_1 + n_2) + \Delta G^{\text{interior}}_{\text{asymm}}(n_1, n_2) + \Delta G^{\text{interior}}_{\text{mm}}(\phi_{[i:d]}, \phi_{[e:j]}) \tag{S116}$$

For the size-dependent term:

$$\Delta G^{\text{interior}}_{\text{size}}(n) = \begin{cases} \Delta G^{\text{interiorsize}}_{n}, & n \leq 30 \\ \Delta G^{\text{interiorsize}}_{30} + \log\left(\frac{n}{30}\right) \Delta G^{\text{polymer}}_{\text{entropy}}, & n > 30 \end{cases} \tag{S117}$$

- $\Delta G^{\text{interiorsize}}_{n}$: empirical values with logarithmic extrapolation to cover the range $1 \leq n \leq 30$. The mixed-material model uses a simple average of the `rna06` and `dna04.2` parameters.[1]

For the asymmetry-based term:

$$\Delta G^{\text{interior}}_{\text{asymm}}(n_1, n_2) = \min(\Delta G^{\text{interiorasymm}}_{5}, |n_1 - n_2| \Delta G^{\text{interiorasymm}}_{\min(4, n_2, n_1)}) \tag{S118}$$

- $\Delta G^{\text{interiorasymm}}_{n}$: The mixed-material model uses a simple average of the empirically regressed values in `rna06` and `dna04.2`.[1]

For the mismatch-based term:

$$\Delta G^{\text{interior}}_{\text{mm}}(\phi_{[i:d]}, \phi_{[e:j]}) = \begin{cases} \Delta G^{\text{interiormm}'}_{\phi_{j-1},\phi_j,\phi_i,\phi_{i+1}} + \Delta G^{\text{interiormm}'}_{\phi_{d-1},\phi_d,\phi_e,\phi_{e+1}}, & i + 2 = d \text{ or } e + 2 = j \\ \Delta G^{\text{interiormm}}_{\phi_{j-1},\phi_j,\phi_i,\phi_{i+1}} + \Delta G^{\text{interiormm}}_{\phi_{d-1},\phi_d,\phi_e,\phi_{e+1}}, & \text{otherwise} \end{cases} \tag{S119}$$

- $\Delta G^{\text{interiormm}}_{\phi_{j-1},\phi_j,\phi_i,\phi_{i+1}}$: An empirical parameter for mismatched bases next to paired bases in an interior loop.
  - Hybrid interior mismatch values were calculated using equation (S113) with SVD linear regression as described earlier. Any interior mismatch represented in hybrid 1×1 loops that were experimentally profiled were regressed first. Then, any remaining Watson–Crick-closed hybrid interior mismatch values were regressed using estimated Watson–Crick-closed hybrid 1×1 loop values while holding the interior mismatch values regressed from experimentally measured 1×1 loops fixed. Finally, all rG·dT-closed hybrid interior mismatch values were regressed using estimated rG·dT-closed hybrid 1×1 loop values while holding all the previously regressed Watson–Crick-closed hybrid interior mismatch values fixed. The missing rG·dT-closed hybrid interior mismatch values were regressed after the missing Watson–Crick-closed hybrid interior mismatch values to reflect increased uncertainty in the estimated rG·dT-closed hybrid 1×1 loop values, given that no rG·dT-closed hybrid 1×1 loops have been experimentally measured.
  - The rest of the mixed-material interior mismatches are a weighted average between the `rna06` and `dna04.2` interior mismatch with the same sequence:

$$\left(\frac{m_{\text{RNA}}}{4}\right)\Delta G^{\text{interiormm,RNA}}_{\phi_{j-1},\phi_j,\phi_i,\phi_{i+1}} + \left(\frac{m_{\text{DNA}}}{4}\right)\Delta G^{\text{interiormm,DNA}}_{\phi_{j-1},\phi_j,\phi_i,\phi_{i+1}}, \tag{S120}$$

    where $m_{\text{RNA}}$ and $m_{\text{DNA}}$ are the number of RNA and DNA bases in the 4-base input sequence (2 bases in the mismatch and 2 in the closing base pair).

- $\Delta G^{\text{interiormm}'}_{\phi_{j-1},\phi_j,\phi_i,\phi_{i+1}}$: a different parameters is used when one side of the interior loop has only one unpaired nucleotide. For single-material interior loops, this parameter was set to $\Delta G^{\text{interiormm}}_{A,\phi_j,\phi_i,A}$ in `rna06` and `dna04.2` corresponding to setting both mismatched bases to A. The mixed-material model uses the same approach, replacing both mismatched bases with A of the corresponding material and evaluating $\Delta G^{\text{interiormm}}_{A,\phi_j,\phi_i,A}$. For example, a rArG/rGdT mismatch is replaced with the rArA/dAdT interior mismatch.

#### S4.7 Free energy model for mixed-material multiloops: RNA/DNA

A multiloop contains 3 or more terminal base pairs and no nicks. It may be defined as a series of bounding subsequences $[\phi]$. The free energy for a multiloop in a specified stacking state, $\omega$, is modeled as follows:

$$\begin{aligned}\Delta G^{\text{multi}}([\phi],\omega) = {} & \Delta G^{\text{multi}}_{\text{init}} + n_{\text{bp}}\Delta G^{\text{multi}}_{\text{bp}} + n_{\text{nt}}\Delta G^{\text{multi}}_{\text{nt}} \\ & + \Delta G^{\text{allterminalbp}}([\phi]) + \Delta G^{\text{allcoax}}([\phi],\omega) + \Delta G^{\text{alldangle}}([\phi],\omega)\end{aligned} \tag{S121}$$

where $n_{\text{bp}}$ is the number of terminal base pairs and $n_{\text{nt}}$ is the number of unpaired nucleotides.

- $\Delta G^{\text{multi}}_{\text{init}}$: penalty for formation of a multiloop. The mixed-material model uses a simple average of the `rna06` and `dna04.2` parameters.
- $\Delta G^{\text{multi}}_{\text{bp}}$: sequence-independent penalty for a terminal base pair in a multiloop. The mixed-material model uses a simple average of the `rna06` and `dna04.2` parameters.
- $\Delta G^{\text{multi}}_{\text{nt}}$: sequence-independent penalty per unpaired nucleotide in a multiloop. The mixed-material model uses the `rna06` parameter for RNA nucleotides and the `dna04.2` parameter for DNA nucleotides.
- $\Delta G^{\text{allterminalbp}}([\phi])$ is the sum of the sequence-dependent free energy $\Delta G^{\text{terminalbp}}_{\phi_i,\phi_j}$ for each terminal base pair $i \cdot j$ in the multiloop, with sequence indicating both material and base (e.g., $\phi_i$ = dT or $\phi_i$ = rA).
- $\Delta G^{\text{allcoax}}([\phi],\omega)$ is the sum of coaxial stacking energies $\Delta G^{\text{coax}}_{\phi_i,\phi_d,\phi_{d+1},\phi_j}$ in the stacking state $\omega$, considered only for adjacent base pairs $i \cdot d$ and $d+1 \cdot j$.
  - $\Delta G^{\text{coax}}_{\phi_i,\phi_d,\phi_{d+1},\phi_j}$: derived for the mixed-material model using the same approach as for stack loops (see Section S4.6), except applied to single-material $\Delta G^{\text{coax}}_{\phi_i,\phi_d,\phi_{d+1},\phi_j}$ values rather than $\Delta G^{\text{stack}}_{\phi_i,\phi_d,\phi_{d+1},\phi_j}$ values.
- $\Delta G^{\text{alldangle}}([\phi],\omega)$ is a sum of the sequence-dependent free energy, $\Delta G^{\text{dangle}}_{i,j}(\phi)$, for each terminal base pair $i \cdot j$ not involved in a coaxial stack in the given stacking state $\omega$, with sequence indicating both material and base (e.g., $\phi_i$ = dT

or $\phi_i$ = rA). For a given terminal base pair $i \cdot j$, $\Delta G^{\text{dangle}}_{i,j}(\phi)$ takes one of four values to match the dangle stacking state for a given $\omega$:

$$\Delta G^{\text{dangle}}_{i,j}(\phi) = \begin{cases} 0 & \text{no dangles} \\ \Delta G^{5'\text{dangle}}_{\phi_i,\phi_{i+1},\phi_j} & 5'\text{ dangle} \\ \Delta G^{3'\text{dangle}}_{\phi_i,\phi_{j-1},\phi_j} & 3'\text{ dangle} \\ \Delta G^{\text{terminalmm}}_{\phi_i,\phi_{i+1},\phi_{j-1},\phi_j} & \text{terminal mismatch} \end{cases} \tag{S122}$$

Note that the state where both $5'$ and $3'$ dangles stack on terminal base pair $i \cdot j$ is classified as a terminal mismatch. For the recursions without dangle stacking (Section S6.3), the term $\Delta G^{\text{alldangle}}([\phi], \omega)$ is neglected.

- $\Delta G^{5'\text{dangle}}_{\phi_i,\phi_j,\phi_k}$: The mixed model uses the weighted average between the $5'$ dangle free energy parametrized for `rna06` and `dna04.2`,[1]

$$\left(\frac{m_{\text{RNA}}}{3}\right) \Delta G^{5'\text{dangle,RNA}}_{\phi_i,\phi_j,\phi_k} + \left(\frac{m_{\text{DNA}}}{3}\right) \Delta G^{5'\text{dangle,DNA}}_{\phi_i,\phi_j,\phi_k}, \tag{S123}$$

  where $m_{\text{RNA}}$ and $m_{\text{DNA}}$ are the number of RNA and DNA bases in the 3-base input sequence (1 dangling base and 2 in the terminal base pair).

- $\Delta G^{3'\text{dangle}}_{\phi_i,\phi_j,\phi_k}$: The mixed model uses the weighted average between the $3'$ dangle free energy parametrized for `rna06` and `dna04.2`,[1]

$$\left(\frac{m_{\text{RNA}}}{3}\right) \Delta G^{3'\text{dangle,RNA}}_{\phi_i,\phi_j,\phi_k} + \left(\frac{m_{\text{DNA}}}{3}\right) \Delta G^{3'\text{dangle,DNA}}_{\phi_i,\phi_j,\phi_k}, \tag{S124}$$

  where $m_{\text{RNA}}$ and $m_{\text{DNA}}$ are the number of RNA and DNA bases in the 3-base input sequence (1 dangling base and 2 in the terminal base pair).

- $\Delta G^{\text{terminalmm}}_{\phi_i,\phi_{i+1},\phi_{j-1},\phi_j}$: An empirical parameter for mismatched bases next to paired bases not in an interior or hairpin context. Terminal mismatches are a weighted average between the `rna06` and `dna04.2` terminal mismatch with the same sequence:

$$\left(\frac{m_{\text{RNA}}}{4}\right) \Delta G^{\text{terminalmm,RNA}}_{\phi_i,\phi_{i+1},\phi_{j-1},\phi_j} + \left(\frac{m_{\text{DNA}}}{4}\right) \Delta G^{\text{terminalmm,DNA}}_{\phi_i,\phi_{i+1},\phi_{j-1},\phi_j}, \tag{S125}$$

  where $m_{\text{RNA}}$ and $m_{\text{DNA}}$ are the number of RNA and DNA bases in the 4-base input sequence (2 in the terminal mismatch and 2 in the terminal base pair).

##### S4.8 Free energy model for mixed-material exterior loops: RNA/DNA

An exterior loop is a loop containing one nick and zero or more terminal base pairs. An exterior loop may be defined as a series of bounding subsequences $[\phi]$ with a given nick location. An unpaired strand is an exterior loop with a free energy of zero, corresponding to the reference state.[2] The free energy of an exterior loop in a specified stacking state $\omega$ is modeled as follows:

$$\Delta G^{\text{exterior}}([\phi], \omega) = 0 + \Delta G^{\text{allterminalbp}}([\phi]) + \Delta G^{\text{allcoax}}([\phi], \omega) + \Delta G^{\text{alldangle}}([\phi], \omega). \tag{S126}$$

For mixed-material exterior loops, the functions $\Delta G^{\text{allterminalbp}}([\phi])$, $\Delta G^{\text{allcoax}}([\phi], \omega)$, and $\Delta G^{\text{alldangle}}([\phi], \omega)$ are defined as above for multiloops. For the recursions without coaxial and dangle stacking (Section S6.3), the terms $\Delta G^{\text{allcoax}}([\phi], \omega)$ and $\Delta G^{\text{alldangle}}([\phi], \omega)$ are neglected.

#### S5 Mixed-material free energy model details: RNA/2′OMe-RNA

The functional form of the free energy model for mixed-material systems is the same as for single-material systems (see Section S3). Here, we describe mixed-material RNA/2′OMe-RNAparameters for each term in the model, taking advantage of empirically derived parameters where available and use single-material RNA parameters where they are not.

##### S5.1 Mixed-material parameter set: RNA/2′OMe-RNA

A mixed-material parameter set contains parameters for mixed-material loops and also specifies the names of two single-material parameter sets with which it is used in conjunction. The following temperature-dependent parameter set is supported:

- `rna_merna06`: RNA/2′OMe-RNA parameters based on hybrid stack loops[30] with additional parameters for mixed-material loops as defined below; for use in conjunction with single-material parameter sets `rna06` and `merna06`.

#### S5.2 Definition of mixed-material Watson–Crick and wobble pairs: RNA/2′OMe-RNA

Mixed-material base pairs are either Watson–Crick pairs:

- RNA/2′OMe-RNA: rG·mC, mG·rC, rA·mU, and mA·rU.

or wobble pairs:

- RNA/2′OMe-RNA: rG·mU and mG·rU.

#### S5.3 Mixed-material salt conditions: RNA/2′OMe-RNA

The default salt condition for our RNA/2′OMe-RNA[30] parameter set is [$Na^+$] = 0.12 M. Due to the unavailability of empirical melt data for RNA/2′OMe-RNA in other salt conditions, no salt correction is supported to enable RNA/2′OMe-RNA calculations at other $Na^+$ concentrations.

#### S5.4 Mixed-material strand association penalty: RNA/2′OMe-RNA

A $\Delta G^{\text{assoc}}_{\text{pub, hybrid(RNA:2}'\text{OMe-RNA)}}$ association penalty derived from RNA/2′OMe-RNA hybrid duplexes is available in the literature.[30] Because `rna06` and `merna06` parameters were derived with different association penalties ($\Delta G^{\text{assoc}}_{\text{pub, RNA}}$ and $\Delta G^{\text{assoc}}_{\text{pub, 2}'\text{OMe-RNA}}$), a weighted average, $\Delta G^{\text{assoc}}_{\text{pub, RNA/2}'\text{OMe-RNA}}$, between the `rna06` and `merna06` association penalties is used for mixed RNA/2′OMe-RNA complexes, centered at $\Delta G^{\text{assoc}}_{\text{pub, hybrid(RNA:2}'\text{OMe-RNA)}}$. Let $p_{\text{RNA}}$ and $p_{2'\text{OMe-RNA}}$ be the proportions of RNA and 2′OMe-RNA nucleotides in the complex, respectively. Then

$$\Delta G^{\text{assoc}}_{\text{pub,RNA/2}'\text{OMe-RNA}} \equiv \Delta G^{\text{assoc}}_{\text{pub,hybrid(RNA:2}'\text{OMe-RNA)}} + (2p_\sigma - 1)(\Delta G^{\text{assoc}}_{\text{pub},\sigma} - \Delta G^{\text{assoc}}_{\text{pub,hybrid(RNA:2}'\text{OMe-RNA)}}) \tag{S127}$$

$$\sigma = \begin{cases} \text{RNA}, & p_{\text{RNA}} \geq 0.5 \\ 2'\text{OMe-RNA}, & \text{otherwise} \end{cases} \tag{S128}$$

which reduces to $\Delta G^{\text{assoc}}_{\text{pub, hybrid(RNA:2}'\text{OMe-RNA)}}$ when $p_{\text{RNA}} = p_{2'\text{OMe-RNA}} = 0.5$.

#### S5.5 Free energy model for mixed-material hairpin loops: RNA/2′OMe-RNA

The terminal base-pair penalty, $\Delta G^{\text{terminalbp}}_{\phi_j,\phi_i}$, is an empirical penalty for non-GC terminal base pairs at the end of a duplex. In the mixed-material model, rA·rU terminal base pairs use the `rna06` value and mA·mU terminal base pairs use the `merna06` value. A single empirically regressed value is used for both rA·mU and mA·rU.[30] Terminal wobble pairs rG·mU and mG·rU use the `rna06` value. Due to the unavailability of additional empirical data, for mixed-material RNA/2′OMe-RNA hairpin loops, with the exception of $\Delta G^{\text{terminalbp}}_{\phi_i,\phi_j}$, all other terms (see Section S3.6) use the `rna06` values.

#### S5.6 Free energy model for mixed-material interior loops: RNA/2′OMe-RNA

An interior loop may be defined via a pair of subsequences $\phi_{[i:d]}$ and $\phi_{[e:j]}$ such that $i < d < e < j$ with base pairs $i \cdot j$ and $d \cdot e$, with no additional paired bases or nicks within the two subsequences.

**Stack loops.** Stack loops are the special case where $d = i + 1$ and $j = e + 1$. The stack loop energy model is as follows:

$$\Delta G^{\text{stackedpair}}(\phi_{[i:i+1]}, \phi_{[j-1:j]}) = \Delta G^{\text{stack}}_{\phi_j,\phi_i,\phi_{i+1},\phi_{j-1}} \tag{S129}$$

- $\Delta G^{\text{stack}}_{\phi_j,\phi_i,\phi_{i+1},\phi_{j-1}}$: empirical values for stack free energies.
  - For the mixed-material RNA/2′OMe-RNA model, empirical stack energies are available for 16 hybrid stacks (rr/mm) not including wobble pairs.[30]
  - For stack loops with a single 2′OMe-RNA nucleotide and three RNA nucleotides, the RNA nucleotide on the same strand as the 2′OMe-RNA nucleotide is substituted with a 2′OMe-RNA nucleotide and the respective hybrid stack value is used.
  - For chimeric stack loops with two 2′OMe-RNA nucleotides and two RNA nucleotides, two hybrid analogs of the stack are made by converting one strand to RNA and the other to 2′OMe-RNA (or vice versa); the average value of the two hybrid stacks is used. If the chimeric stack is symmetric (e.g. mArU/mArU), then there is only one hybrid stack analog.

- ◦ For stack loops with three 2′OMe-RNA nucleotides and one RNA nucleotide, the 2′OMe-RNA nucleotide on the same strand as the RNA nucleotide is substituted with a RNA nucleotide and the respective hybrid stack value is used.
- ◦ Note that for the single-material parameter set `merna06`, for stack loops with four 2′OMe-RNA nucleotides and zero RNA nucleotides, the average value of the two hybrid stacks (rr/mm and mm/rr) containing those nucleotides is used. If the stack is symmetric (e.g., mAmU/mAmU), then there is only one such hybrid stack containing those nucleotides.

**Bulge loops.** A bulge loop is the special case with either $d = i + 1$ or $j = e + 1$ but not both. Here, we will outline the functional form when $d = i + 1$. Let $n \equiv j - e - 1$ denote the number of unpaired nucleotides in the bulge loop. Due to the unavailability of additional empirical data, for mixed-material RNA/2′OMe-RNA bulge loops, with the exception of the terminal base pair penalty, $\Delta G^{\text{terminalbp}}_{\phi_j,\phi_i}$ (defined in Section S5.5), and the stack loop term, $\Delta G^{\text{stack}}_{\phi_j,\phi_i,\phi_{i+1},\phi_{j-1}}$ (defined above in this section), all other terms (see Section S3.7) use the `rna06` values.

**Other small interior loops: 1×1 interior loops, 1×2 interior loops, and 2×2 interior loops.** Due to the unavailability of additional empirical data, for mixed-material RNA/2′OMe-RNA small interior loops, the `rna06` lookup tables are used (see Section S3.7).

**Other large interior loops.** Due to the unavailability of additional empirical data, for mixed-material RNA/2′OMe-RNA large interior loops, the `rna06` values are used for all terms (see Section S3.7).

#### S5.7 Free energy model for mixed-material multiloops: RNA/2′OMe-RNA

A multiloop contains 3 or more terminal base pairs and no nicks. It may be defined as a series of bounding subsequences $[\phi]$. For mixed-material RNA/2′OMe-RNA multiloops, the terminal base-pair penalty, $\Delta G^{\text{terminalbp}}_{\phi_j,\phi_i}$ (defined in Section S5.5) is used and the coaxial stack energy, $\Delta G^{\text{coax}}_{\phi_i,\phi_d,\phi_{d+1},\phi_j}$, is set equal to the analogous stack loop energy $\Delta G^{\text{stack}}_{\phi_i,\phi_d,\phi_{d+1},\phi_j}$ (defined in Section S5.6). Due to the unavailability of additional empirical data, for mixed-material RNA/2′OMe-RNA multiloops, `rna06` values are used for all other terms (see Section S3.8).

#### S5.8 Free energy model for mixed-material exterior loops: RNA/2′OMe-RNA

An exterior loop is a loop containing one nick and zero or more terminal base pairs. An exterior loop may be defined as a series of bounding subsequences $[\phi]$ with a given nick location. An unpaired strand is an exterior loop with a free energy of zero, corresponding to the reference state.[2] For mixed-material RNA/2′OMe-RNA exterior loops, the terminal base-pair penalty, $\Delta G^{\text{terminalbp}}_{\phi_j,\phi_i}$ (defined in Section S5.5), and the coaxial stack energy, $\Delta G^{\text{coax}}_{\phi_i,\phi_d,\phi_{d+1},\phi_j}$ (defined in Section S5.7) are used. Due to the unavailability of additional empirical data, for mixed-material RNA/2′OMe-RNA exterior loops, `rna06` values are used for all other terms (see Section S3.9).

### S6 Mixed-Material Recursions

Recursions specify the dependencies between subproblems and incorporate the details of the complex structural ensemble and the free energy model. The recursions described here can be combined with a quantity-specific evaluation algebra (see Section S3 of Reference 1) and a quantity-specific operation order (see Section S4 of Reference 1) to calculate diverse physical quantities.[1] Each recursion corresponds to an efficient iteration through a *conditional ensemble* of substructures within a given subsequence that are compatible with a specified set of constraints. For a given recursion, a conditional ensemble might include an explicit structural element, which can be considered the base case of the recursion, or a reference to the result of another recursion.

#### S6.1 Separate recursions for intrastrand and interstrand blocks

Reference 2 described dynamic programming recursions for the complex ensemble that checked for a nick next to each nucleotide. This approach enabled treatment of complexes containing an arbitrary number of strands, but caused unnecessary complications in the program flow and eliminated any possibility of vectorization due to the conditional checks within each "for" loop. By contrast, here we follow Reference 1 and employ separate sets of recursions for triangular intrastrand blocks and rectangular interstrand blocks (Figure S4). As a result, each intrastrand and interstrand recursion is kept as simple as possible and both types of recursions can be efficiently vectorized.

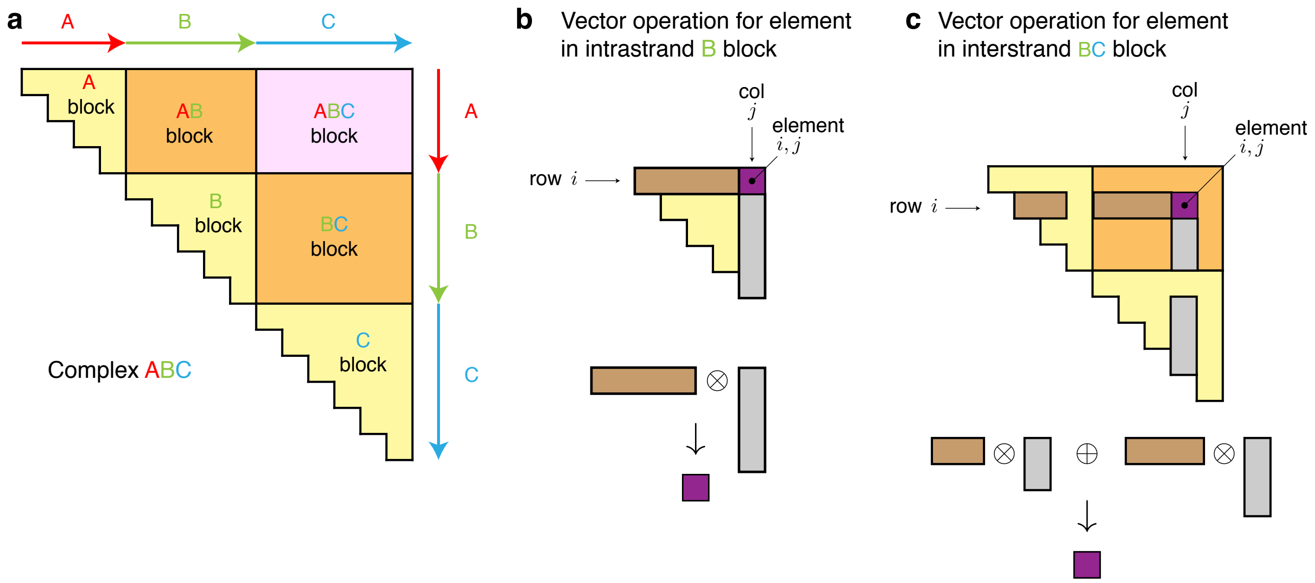

Figure S4: Separate recursions for intrastrand and interstrand blocks. (a) Triangular intrastrand blocks (A, B, C) and rectangular interstrand blocks (AB, BC, ABC) for complex ABC. Element $i, j$ corresponds to a conditional ensemble for subsequence $[i, j]$ which contains no nicks if $i, j$ is in an intrastrand block and one or more nicks if $i, j$ is in an interstrand block. (b) Each recursion operation for calculation of element $i, j$ in an intrastrand block (e.g., $Q_{i,j} \leftarrow \sum_{i\le d<j} Q_{i,d}Q_{d+1,j}$) can be implemented as a vectorized dot product between a subvector of row $i$ (brown) and a subvector of column $j$ (gray) to obtain element $i, j$ (purple). Note that calculation of an element $i, j$ in an intrastrand block uses elements in the same intrastrand block (calculated using intrastrand recursions). (c) Each recursion operation for calculation of element $i, j$ in an interstrand block (e.g., $Q_{i,j} \leftarrow \sum_{i\le d<j,\ \mathrm{strand}(d)=\mathrm{strand}(d+1)} Q_{i,d}Q_{d+1,j}$) can be implemented as multiple vectorized dot products between valid subvectors of row $i$ (brown) and valid subvectors of column $j$ (gray) to obtain element $i, j$ (purple), where valid positions are those that avoid introducing disconnected structures into the complex ensemble (see Algorithm S2). Note that calculation of element $i, j$ in an interstrand block uses elements in one or more interstrand blocks (calculated with interstrand recursions) and two intrastrand blocks (calculated with intrastrand recursions). Used with permission from Fornace et al., *ACS Synth Biol*, 9, 2665-2678, 2020. Copyright 2020 American Chemical Society.

#### S6.2 Conventions for recursion diagrams and equations

In the following sections we will describe recursions corresponding to the complex ensemble without stacking terms (Section S6.3) and with coaxial and dangle stacking subensembles (Section S6.5). Each recursion iterates over all conditional ensembles compatible with the constraints defined for a given recursion type. For a complex of $N$ nucleotides, each full set of recursions is $O(N^3)$ in time and $O(N^2)$ in space. For interior loop recursions, we start by defining an $O(N^4)$ recursion and then describe an exact reduction to $O(N^3)$ time complexity (Section S6.4).

Each recursion is represented in two ways: graphically, as a set of recursion diagrams, and algebraically, as an equation defining the recursion as a specific combination of contributions. The recursion diagrams employ the following conventions:

- Solid circular arcs depict the nucleic acid backbone. An arrowhead denotes the $3'$ end of a strand.
- Dots indicate particular nucleotide positions that define the bounds of recursive contributions. If a dot is labeled with a nucleotide index, the same index is used in the corresponding recursion. If a dot is adjacent to a dot labeled $i$, the implied index of the unlabeled dot is either $i-1$ or $i+1$ (indices increase from $5'$ to $3'$).
- A straight line delimits the boundary for a given contribution. A solid straight line indicates that the connected nucleotides are base-paired. A dashed straight line indicate that the connected nucleotides may or may not be base-paired. A half-solid/half-dashed straight line indicates that the nucleotide connected on the solid side is base-paired to a nucleotide within the demarcated region. A straight line that is solid at both ends and dashed in the middle indicates that the nucleotides at either end are both base-paired but not to each other. A dotted straight line indicates that the connected nucleotides are involved in a stacking state (either a coaxial stacking state or a dangle stacking state). A dotted straight line with hollow dots indicates that there is no base pairing within the demarcated region.
- If a recursive contribution corresponds to a single- or mixed-material contribution, then the nucleic acid backbone within the demarcated region is colored to denote single- or mixed-material. If a terminal base pair in a recursion diagram corresponds to a single- or mixed-material, then the terminal base pair is colored to denote single- or mixed-material.
- Shading indicates that the shaded region in a recursion explicitly incorporates a *recursion energy*, $\Delta G$, representing all or part of a loop free energy (e.g., multiloop recursion energies representing different terms in the multiloop model are incorporated in multiple places in multiple recursions in order to treat the full multiloop model). The color of the shading corresponds to the loop type (and the stacking type when applicable).

A recursion equation provides a mathematical description of the conditional ensemble depicted graphically in a recursion diagram. Recursion equations employ the following conventions:

- For each physical quantity, an appropriate evaluation algebra (see Section S3 of Reference 1) is used to define the generic operators that appear in the recursion equations: $\mathbb{0}$, $\mathbb{1}$, $\oplus$, $\otimes$, $W$, and $Q$. For example, to calculate the partition function, we have:

$$\mathbb{0} \to 0, \quad \mathbb{1} \to 1, \quad \oplus \to +, \quad \otimes \to \times, \quad W(g) \to \exp(-g/kT), \quad Q^a_{i,j} \to Q^a_{i,j}. \tag{S130}$$

  where the last right-hand side indicates that $Q^a_{i,j}$ is a lookup of the relevant stored matrix element.
- A recursion equation for subsequence $[i:j]$ corresponding to element $i,j$ in a triangular intrastrand block is denoted $R^a_{\text{INTRA}}(i,j,\phi)$ for a recursion of type $a$ (e.g., $a \in \{\varnothing, b, m, \dots\}$). A recursion equation for subsequence $[i:j]$ corresponding to element $i,j$ in a rectangular interstrand block is denoted $R^a_{\text{INTER}}(i,j,\phi)$ for a recursion of type $a$. Here, $\phi$ is the sequence of the complex and $i$ and $j$ are nucleotide indices. Note that in the Supporting Information we use $Q^{\varnothing}_{i,j}$ to denote $Q_{i,j}$ so that each recursion has an explicit recursion type $a$. In the mixed-material setting, multiloop and interior loop recursions also have a material type "single" or "mix" (e.g., $R^{a,\text{single}}_{\text{INTRA}}(i,j,\phi)$ and $R^{a,\text{mix}}_{\text{INTRA}}(i,j,\phi)$ for intrastrand blocks and $R^{a,\text{single}}_{\text{INTER}}(i,j,\phi)$ and $R^{a,\text{mix}}_{\text{INTER}}(i,j,\phi)$ for interstrand blocks).
- If a recursion diagram contains a shaded region denoting a recursion energy, $\Delta G$, the corresponding recursion equation will incorporate the recursion energy via the term $W(\Delta G)$.
- In the mixed-material setting, the material of nucleotide $i$ (e.g., RNA or DNA) is denoted $\mu_i$. $\textsc{IsSingle}(i,j,\mu_k)$ checks if subsequence $[i,j]$ is composed entirely of material $\mu_k$. Because there are multiple parameter sets in the mixed-material setting (e.g., RNA, DNA, and RNA/DNA), a parameter can have an additional material subscript to specify which parameter set it belongs to. For example, the multiloop initiation parameter $\Delta G^{\text{multi}}_{\text{init}}$ can be specified as $\Delta G^{\text{multi}}_{\text{init},\mu_i}$ to indicate that it is from the single-material parameter set for material $\mu_i$ and as $\Delta G^{\text{multi}}_{\text{init,mix}}$ to indicate that it is from the mixed-material parameter set.
- After it is evaluated for the first time, $R^a(i,j,\phi)$ is used to yield $Q^a_{i,j}$ in subsequent recursions. In the evaluation algebras that generate scalars (SumProduct, MinSum, Count), the output of $R^a(i,j,\phi)$ is synonymous with the value $Q^a_{i,j}$ that is stored in the recursion matrices. However, other evaluation algebras involve different treatment of the output of $R^a(i,j,\phi)$. For instance, a recursion in the SplitExp evaluation algebra (see Section S3.1.4 of Reference 1) yields a function that must be supplied with a reference exponent $\gamma$ to calculate the mantissa and exponent values that are stored. The ways in which recursion outputs are utilized for each physical quantity are described in Section S4 of Reference 1.

```
DOT(vectors)
n ← LENGTH(vectors_1)
x ← 𝟘
for i ∈ [1 : n]
    t ← 𝟙
    for a ∈ vectors
        t ← t ⊗ a_i
    x ← x ⊕ t
return x
```

Algorithm S1: Generalized dot product over multiple vectors of equal length.

In our pseudocode, we make clear which operations are vectorized using SIMD operations on contiguous arrays via the function DOT (Algorithm S1), which represents a dot product generalized to any number of arguments, each of which is a vector of the same length $n$. The **vectors** argument to this subroutine is composed of row or column subvectors (each a vector of contiguous elements) of the recursion matrices storing the result of previous recursion evaluations (e.g., $Q^{\varnothing}, Q^{b}, Q^{m}, \dots$).

To denote a vector extracted from a matrix block, we replace a scalar index (e.g., $d$) with a vector index (e.g., $\overline{d}$) representing a range of either row or column indices. For example:

$$\overline{d} \equiv [i : j-5] \equiv i, i+1, \dots, j-6, j-5. \tag{S131}$$

represents an ascending range of indices. Any scalar increment is applied to each entry in the range:

$$\overline{d} + 1 \equiv [i+1 : j-4]. \tag{S132}$$

$Q^{\varnothing}_{i,\overline{d}}$ then denotes a subvector of row $i$ from matrix $Q^{\varnothing}$, $Q^{s}_{\overline{d}+1,j}$ denotes a subvector of column $j$ from matrix $Q^{s}$, and

$$\text{DOT}\left(Q^{\varnothing}_{i,\overline{d}}, Q^{s}_{\overline{d}+1,j}\right) \tag{S133}$$

denotes a dot product between these two vectors.

For calculation of matrix elements in interstrand blocks (which by definition involve 2 or more strands), $\eta$ is an array of indices of the nicks between strands within the interstrand block being considered; by convention, each nick is denoted in $\eta$ by the index of the nucleotide following the nick.[†] For example, consider a complex ABC with strands A, B, and C containing 4, 5, and 6 nucleotides respectively. For the AB block, $\eta = [5]$. For the BC block, $\eta = [10]$. For the ABC block, $\eta = [5, 10]$.

To calculate matrix entry $i, j$ for an interstrand block with nicks $\eta$, the function VALID$(i, j, \eta)$ (Algorithm S2) returns the set of valid ranges $\{\overline{d}_1, \overline{d}_2, \dots\}$ for vectorization so as to ensure that all secondary structures are connected and exterior loops appear only when they are explicitly considered by a recursion. There is at most one valid vectorization range per strand, and there may be none for a strand that is too short or for the first or last strand if $i$ or $j$, respectively, is too close to a nick. The ability to identify valid vectorization ranges for calculating each matrix element is a key innovation enabled by using dedicated recursions for intrastrand and interstrand blocks, eliminating the use of "if" statements to identify nick locations (cf. Reference 2), and thus enabling vectorization to achieve dramatic speedups.[1]

Steric requirements require that there be at least three intervening bases between two base-paired nucleotides on the same strand, placing a lower bound on the length of subsequence $[i, j]$ for different recursion types (e.g., a minimum subsequence length to contain a hairpin loop, an interior loop, a multiloop, a terminal base pair, a stacking state, a coaxial stacking state, or a dangle stacking state). Recursions below the minimum subsequence length for a given recursion type return $\mathbb{0}$. For efficiency reasons, we often explicitly specify lower bounds on subsequence length to avoid performing calculations for elements that will evaluate to $\mathbb{0}$.

For exterior loop and multiloop recursions without coaxial and dangle stacking, the elementary recursion entity is a *terminal base pair* (a base pair that terminates a duplex to form a part of the exterior loop or multiloop). For exterior and multiloop recursions with coaxial and dangle stacking, the elementary recursion entity is the *stacking state*, representing either a *coaxial stacking state* (two adjacent terminal base pairs that are coaxially stacked) or a *dangle stacking state* (zero, one, or two unpaired nucleotides dangle stacking on an adjacent terminal base pair).

[†]Note that this definition is unrelated to the use of $\eta$ in Reference 2 to denote the number of nicks in a given subsequence.

```
VALID(i, j, η)
D ← {}
m ← FIRST(η)
n ← LAST(η)
if i + 1 < m and j ≥ n
    d̄ = [i : m − 2]
    D ← D ∪ d̄
if i < m and j − 1 ≥ n
    d̄ = [n : j − 1]
    D ← D ∪ d̄
if i < m and j ≥ n
    for b ∈ [1 : LENGTH(η) − 1]
        if η_{b+1} − η_b > 1
            d̄ = [η_b : η_{b+1} − 2]
            D ← D ∪ d̄
return D
```

Algorithm S2: Enumerate valid positions for vectorization in an interstrand block. $\eta$ is an array of indices of the nicks between strands within the interstrand block being considered; by convention, each nick is denoted in $\eta$ by the index of the nucleotide following the nick. The algorithm identifies at most one valid range $\overline{d}$ for each strand in the block, corresponding to the values of the index $d$ such that $d$ and $d+1$ are on the same strand. This requirement ensures that all secondary structures are connected and that exterior loops only appear when they are being explicitly considered by a recursion. The algorithm returns **D**, the set of valid ranges for the block. For a given $i$ and $j$, each valid range leads to a dot product between range $\overline{d}$ of row $i$ and range $\overline{d}+1$ of column $j$ (e.g., the recursion of Figure S12 contains one dot product for each valid range $\overline{d}$).

#### S6.3 Recursions without coaxial and dangle stacking subensembles

Here, we describe $R^a_{\text{INTRA}}(i,j,\phi)$ recursions for calculating the elements of intrastrand blocks and $R^a_{\text{INTER}}(i,j,\phi)$ recursions for calculating the elements of interstrand blocks for the complex ensemble, $\overline{\Gamma}$, without coaxial and dangle stacking subensembles, in a mixed-material setting. To assist with examining these recursions, the intuition behind the name chosen for each recursion, the nature of the ensemble treated by each recursion, and the dependencies between the different recursions is summarized in Figure S5. For convenience, it may be helpful to consider the recursions from the perspective of partition function calculations since there is a natural correspondence between the generic evaluation algebra nomenclature and the specific operators needed for partition function calculations (see equation (S130)), but the recursions are generic and can be combined with a quantity-specific evaluation algebra and a quantity-specific operation order to calculate diverse physical quantities.[1] The present recursions treat the same structural ensemble $\overline{\Gamma}$ and free energy model $\overline{\Delta G}(\phi, s)$ as our previous implementation (NUPACK 3.2 with dangles option "none").[2]

A recursion $R^a(i,j,\phi)$ operates on subsequence $[i\,:\,j]$ to calculate element $i,j$ for either the unconstrained ensemble $a = \varnothing$ or for one of several constrained ensembles $a \in \{s, b, x, xa, ma, ms, m\}$. Multiloop and interior loop ensembles $a \in \{x, xa, ma, ms, m\}$ are calculated for both single and mixed-material scenarios $b \in \{\text{single}, \text{mix}\}$. This allows the correct free energy parameters to be applied for each loop depending on the material composition of the loop. Briefly, $R^{\varnothing}(i,j,\phi)$ treats the unconstrained ensemble in an exterior loop context where $i$ and $j$ may or may not be paired. $R^s(i,j,\phi)$ serves as an efficiency wrapper over the $3'$-most terminal base pair in an exterior loop context to reduce the time complexity from $O(N^4)$ to $O(N^3)$. $R^b(i,j,\phi)$ treats the constrained ensemble where $i$ and $j$ form base pair $i \cdot j$ in the context of any loop type. $R^{x,\text{single}}(i,j,\phi)$ and $R^{x,\text{mix}}(i,j,\phi)$ treat extensible interior loops (single- or mixed-material, respectively) to reduce the time complexity from $O(N^4)$ to $O(N^3)$. $R^{x,\text{futuremix}}(i,j,\phi)$ treats single-material extensible interior loops with the mixed parameter set to cover the scenario where a single-material extensible interior loop becomes a mixed-material extensible interior loop due to future extension of the loop. $R^{xa,\text{single}}(i,j,\phi)$ and $R^{xa,\text{mix}}(i,j,\phi)$ enable efficient determination of material composition in an interior loop context. $R^{ma,\text{single}}(i,j,\phi)$ and $R^{ma,\text{mix}}(i,j,\phi)$ treat unpaired regions in a multiloop context (for single- and mixed-material unpaired regions, respectively). $R^{ms,\text{single}}(i,j,\phi)$ and $R^{ms,\text{mix}}(i,j,\phi)$ serve as efficiency wrappers over the $3'$-most terminal base pair in a multiloop context (analogous to $R^s$ in an exterior loop context) to reduce the time complexity from $O(N^4)$ to $O(N^3)$ (where the $3'$-most terminal base pair and any unpaired nucleotides are either a single material or mixed-material, respectively). $R^{m,\text{single}}(i,j,\phi)$ and $R^{m,\text{mix}}(i,j,\phi)$ treat the remaining terminal base pairs in a multiloop context (where all remaining terminal base pairs and any unpaired nucleotides are either a single material or mixed-material, respectively).

| Recursion | Naming intuition | Constraint | Context |
|---|---|---|---|
| $\varnothing$ | unconstrained | none | exterior loop |
| $s$ | **s**ummation | efficiency wrapper of $3'$-most terminal base pair | exterior loop |
| $b$ | **b**ase-paired | base pair between $5'$-most and $3'$-most bases of subsequence | any loop |
| $x$, single | e**x**tensible, **single** material | extensible interior loop, single material | interior loop |
| $x$, mix | e**x**tensible, **mix**ed material | extensible interior loop, mixed material | interior loop |
| $x$, futuremix | e**x**tensible, **future** mixed material | extensible interior loop, currently single material but potentially future mixed-material as loop is extended | interior loop |
| $xa$, single | e**x**tensible **a**ccessory, **single** material | extensible interior loop, single-material | interior loop |
| $xa$, mix | e**x**tensible **a**ccessory, **mix**ed material | extensible interior loop, mixed-material | interior loop |
| $ma$, single | **m**ultiloop **a**ccessory, **single** material | unpaired nucleotides, single-material | multiloop |
| $ma$, mix | **m**ultiloop **a**ccessory, **mix**ed material | unpaired nucleotides, mixed-material | multiloop |
| $ms$, single | **m**ultiloop **s**ummation, **single** material | efficiency wrapper of $3'$-most terminal base pair, multiloop is single-material | multiloop |
| $ms$, mix | **m**ultiloop **s**ummation, **mix**ed material | efficiency wrapper of $3'$-most terminal base pair, multiloop is mixed-material | multiloop |
| $m$, single | **m**ultiloop, **single** material | one or more remaining terminal base pairs, multiloop is single-material | multiloop |
| $m$, mix | **m**ultiloop, **mix**ed material | one or more remaining terminal base pairs, multiloop is mixed-material | multiloop |

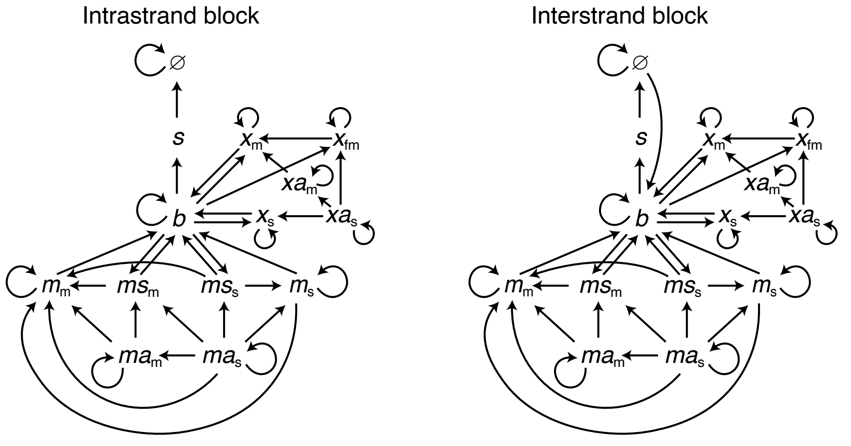

Figure S5: Nomenclature and connectivity for recursions without coaxial and dangle stacking. Top: Nomenclature. Bottom: Dependencies between different recursion types for elements within an intrastrand block (left) or an interstrand block (right). The "single", "mix", and "futuremix" material constraints are abbreviated as the subscripts "s", "m", and "fm", respectively.

##### S6.3.1 Intrastrand dynamic programming recursions without coaxial and dangle stacking

Here, we consider recursions for calculating the entries in a triangular intrastrand block without coaxial and dangle stacking. By definition, there are no nicks between strands in intrastrand recursions since intrastrand blocks involve base-pairing within a single strand.

**$R^{\varnothing}_{\text{INTRA}}$ recursion without coaxial and dangle stacking.** We begin with the recursion $R^{\varnothing}_{\text{INTRA}}(i,j,\phi)$ with the diagram and equation shown in Figure S6. The recursion $R^{\varnothing}_{\text{INTRA}}(i,j,\phi)$ does not change in the mixed-material setting, and is as previously described.[1] $R^{\varnothing}_{\text{INTRA}}(i,j,\phi)$ operates on the unconstrained ensemble for subsequence $[i,j]$ in an exterior loop context where $i$ and $j$ may or may not be paired (depicted with a dashed line between $i$ and $j$ in the recursion diagram). This recursion distinguishes two cases that are combined using $\oplus$ in the recursion equation:

- *No terminal base pairs:* the empty case in an exterior loop context where there are no terminal base pairs in subsequence $[i,j]$ (depicted by the absence of a straight solid line in the recursion diagram). The shading in the recursion diagram represents the recursion energy $\Delta G^{\text{exterior}}_{i,j}(\phi) = 0$ corresponding to the zero reference state for an exterior loop with no base pairs and no coaxial or dangle stacking. The corresponding contribution to the recursion equation is $W(0) = \mathbb{1}$.

- *At least one terminal base pair:* the non-empty case in an exterior loop context where there is at least one terminal base pair (i.e., a base pair terminating a duplex) in subsequence $[i,j]$. The $3'$-most terminal base pair begins at $d+1$ and ends in the interval $[d+2,j]$ (depicted using a half-solid/half-dashed line in the recursion diagram). The contributions for subsequence $[d+1,j]$ are incorporated using a $Q^s_{d+1,j}$ element. Contributions for the remaining subsequence $[i,d]$ are incorporated by a $Q^{\varnothing}_{i,d}$ element. The shading denotes the recursion energy 0 corresponding to the zero reference state in an exterior loop context. Note that the recursion energy $\Delta G^{\text{terminalbp}}(\phi)$ representing one component of the $\Delta G^{\text{exterior}}_{i,j}(\phi)$ free energy is not incorporated here because the full identity of the terminal base pair (i.e., a base pair terminating a duplex) beginning at $d+1$ is not known within the $R^{\varnothing}_{\text{INTRA}}(i,j,\phi)$ recursion (only within the $R^s_{\text{INTRA}}(i,j,\phi)$ recursion). The edge case where the index $d+1=i$ is displayed explicitly to indicate that no $Q^{\varnothing}$ element is accessed in this case. The index limits in the recursion equation reflect the fact that steric effects prevent a hairpin loop with fewer than 3 unpaired nucleotides (hence, $i \cdot j$ cannot form if $j-i<4$).

Note that using the DOT notation (Algorithm S1) and index range notation (Eq S131) to denote vector operations, we have the equivalence:

$$\text{DOT}\left(Q^{\varnothing}_{i,\overline{d}}, Q^s_{\overline{d}+1,j}\right) \equiv \sum_{d=i}^{j-5} Q^{\varnothing}_{i,d} \otimes Q^s_{d+1,j}, \quad j-i>4,$$

$$\text{where } \overline{d} \equiv [i:j-5].$$

We can also recognize that in terms of matrix elements, the dot product

$$\text{DOT}\left(Q^{\varnothing}_{i,\overline{d}}, Q^s_{\overline{d}+1,j}\right) \tag{S134}$$

is between the element range $\overline{d}$ of row $i$ (depicted as brown elements in Figure S4b) and the element range $\overline{d}$+1 of column $j$ (gray elements), yielding element $i,j$ (purple element).

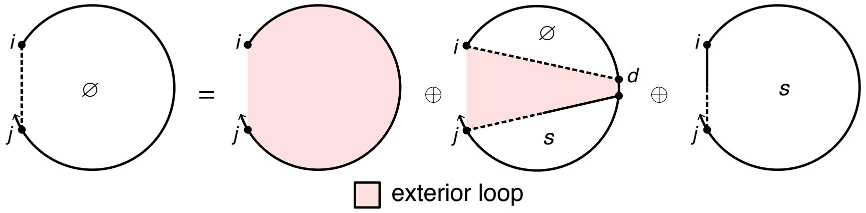

$$R^{\varnothing}_{\text{INTRA}}(i,j,\phi) \equiv \mathbb{1} \oplus \begin{cases} Q^s_{i,j} \oplus \text{DOT}\left(Q^{\varnothing}_{i,\overline{d}}, Q^s_{\overline{d}+1,j}\right), & j-i>4 \\ Q^s_{i,j}, & j-i=4 \\ \mathbb{0}, & \text{otherwise} \end{cases}$$

$$\text{where } \overline{d} \equiv [i:j-5].$$

Figure S6: $R^{\varnothing}_{\text{INTRA}}$ recursion without coaxial and dangle stacking. Top: recursion diagram. Bottom: recursion equation.

**$R^s_{\text{INTRA}}$ recursion without coaxial and dangle stacking.** The $R^{\varnothing}_{\text{INTRA}}(i,j,\phi)$ recursion references $Q^s$ elements that are computed using the $R^s_{\text{INTRA}}$ recursion displayed in Figure S7. The $R^s_{\text{INTRA}}(i,j,\phi)$ recursion does not change in the mixed-material setting and is as previously described.[1] $R^s_{\text{INTRA}}(i,j,\phi)$ operates on a conditional ensemble for subsequence $[i,j]$ in an exterior loop context containing one terminal base pair starting at $i$ and ending in the interval $[i+1,j]$ (depicted as a half-solid/half-dashed line between $i$ and $j$). The contribution for the subsequence $[i,d]$ enclosed by base pair $i \cdot d$ is incorporated using a $Q^b_{i,d}$ element. Shading corresponds to the recursion energy, $\Delta G^{\text{terminalbp}}_{i,d}(\phi)$, representing the sequence-dependent penalty for a terminal base pair in an exterior loop context (dependent on the sequence of base pair $i \cdot d$). The index limits in the recursion equation reflect the fact that steric effects prevent a hairpin loop with fewer than 3 unpaired nucleotides (hence, $i \cdot j$ cannot form if $j-i<4$). Note that the $R^s$ recursion serves as an efficiency wrapper of the $R^b$ recursion (here, representing the 3′-most terminal base pair in an exterior loop context) to reduce the time complexity of the $R^{\varnothing}$ recursion from $O(N^4)$ to $O(N^3)$. This time complexity reduction is achieved by defining the 3′-most base pair using $R^b$ within the $R^s$ efficiency wrapper rather than directly using the $R^b$ recursion within the $R^{\varnothing}$ recursion, so as to avoid introducing a fourth independent index into the $R^{\varnothing}$ recursion.

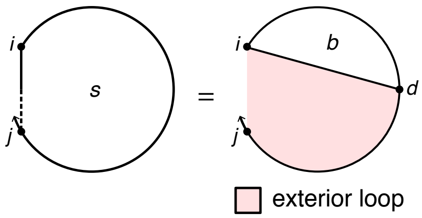

$$R^s_{\text{INTRA}}(i,j,\phi) \equiv \begin{cases} \text{DOT}\left(Q^b_{i,\overline{d}}, W(\Delta G^{\text{terminalbp}}_{i,\overline{d}}(\phi))\right), & j-i \geq 4 \\ \mathbb{0}, & \text{otherwise} \end{cases}$$

where $\overline{d} \equiv [i+4:j]$.

Figure S7: $R^s_{\text{INTRA}}$ recursion without coaxial and dangle stacking. Top: recursion diagram. Bottom: recursion equation.

***$R^b_{\text{INTRA}}$* recursion without coaxial and dangle stacking.** The $R^s_{\text{INTRA}}(i,j,\phi)$ recursion references $Q^b$ elements that are computed using the $R^b_{\text{INTRA}}$ recursion displayed in Figure S8. $R^b_{\text{INTRA}}(i,j,\phi)$ operates on a conditional ensemble for subsequence $[i,j]$ with $i$ and $j$ base paired to each other (depicted with a solid line between $i$ and $j$). The function COMPLEMENTARY$(\phi_i,\phi_j)$ checks if bases $\phi_i$ and $\phi_j$ are complementary (Watson–Crick or wobble pair) without regard to whether $i$ and $j$ are sufficiently separated along the strand to be able to pair sterically. In the mixed-material setting, the appropriate single- or mixed-material sub-recursions and parameters must be applied based on the material composition of the loops. The recursion distinguishes three cases that are combined using $\oplus$ in the recursion equation:

- *Hairpin loop:* the hairpin loop closed by the single base pair $i \cdot j$ (depicted by a straight solid line). The recursion incorporates the recursion energy $\Delta G^{\text{hairpin}}_{i,j}(\phi)$, which takes into account the material of each nucleotide in the subsequence $[i,j]$ (Section S4.5). The index limits in the recursion equation reflect the fact that steric constraints prevent a hairpin loop with fewer than 3 unpaired nucleotides (hence, $i \cdot j$ cannot form if $j-i<4$).
- *Interior loop:* the interior loop closed by the two terminal base pairs $i \cdot j$ and $d \cdot e$ (depicted by straight solid lines). We defer discussion of the calculation of the interior loop contributions using the subroutine INTERIORINTRA until Section S6.4, where we describe both $O(N^4)$ and $O(N^3)$ recursions. The index limits in the recursion equation reflect the fact that steric effects prevent an interior loop with $j-i<6$ due to the steric requirement that there be at least 3 intervening bases between $d$ and $e$.
- *Multiloop:* the multiloop closed by three or more terminal base pairs: 1) the terminal base pair $i \cdot j$ depicted by a straight solid line, 2) a $3'$-most terminal base pair starting at $d$ and ending in the interval $[d+1, j-1]$ (depicted by a straight half-solid/half dashed line between $d$ and $j-1$); the contribution of subsequence $[d, j-1]$ is incorporated by element $Q^{ms}_{d,j-1}$, 3) one or more additional terminal base pairs in the interval $[i+1, d-1]$ (the straight dashed line denotes that $i+1$ and $d-1$ may or may not be paired); the contribution of subsequence $[i+1, d-1]$ is incorporated by element $Q^m_{i+1,d-1}$.
  - *Single- and mixed-material recursions.* Multiloop contributions must be kept separate by material in $R^b_{\text{INTRA}}(i,j,\phi)$ so that the appropriate $\Delta G^{\text{multi}}_{\text{init}}$, $\Delta G^{\text{multi}}_{\text{bp}}$, and $\Delta G^{\text{terminalbp}}_{j,i}(\phi)$ parameters are applied depending on the material of the multiloop. Contributions for single-material and mixed-material multiloops are indicated in the recursion equation as $M_{\text{single}}$ and $M_{\text{mix}}$, respectively. To use the correct physical model for each multiloop as recursion elements are combined across multiple recursion levels, lower level recursions must be calculated separately for the "single" and "mixed" scenarios before they are combined at the $R^b_{\text{INTRA}}(i,j,\phi)$ level. $Q^{ms}_{d+1,j-1}$ is split into $Q^{ms,\text{single}}_{d+1,j-1}$ and $Q^{ms,\text{mix}}_{d+1,j-1}$, while $Q^m_{i+1,d}$ is split into $Q^{m,\text{single}}_{i+1,d}$ and $Q^{m,\text{mix}}_{i+1,d}$. "Single" elements (depicted with a blue backbone in recursion diagrams) only track energy contributions from single-material loop regions, while "mix" elements (depicted with a red background in recursion diagrams) only track contributions from mixed-material loop regions. Because $Q^m_{i+1,d}$ and $Q^{ms}_{d+1,j-1}$ are each split into single-material and mixed-material recursions, there are four possible combinations when multiplying them. Any combination with a mixed element produces an overall mixed multiloop ($Q^{m,\text{mix}}_{i+1,d} \otimes Q^{ms,\text{mix}}_{d+1,j-1}$, $Q^{m,\text{mix}}_{i+1,d} \otimes Q^{ms,\text{single}}_{d+1,j-1}$, $Q^{m,\text{single}}_{i+1,d} \otimes Q^{ms,\text{mix}}_{d+1,j-1}$), which is added to $M_{\text{mix}}$ as the term $C_{1m}$. The remaining combination, $Q^{m,\text{single}}_{i+1,d} \otimes Q^{ms,\text{single}}_{d+1,j-1}$, in term $C_{1s}$ produces a single-material multiloop only if the material of both elements match each other and the material of the closing base pair $i \cdot j$, which is checked using the conditional, $\mu_j = \mu_{j-1} = \mu_{i+1} = \mu_i$. If the conditional is satisfied, the contribution is added

to $M_{\text{single}}$ using term $C_{0s}$. Otherwise, if the conditional fails, the contribution is added to $M_{\text{mix}}$ using term $C_{0m}$. In the conditional, note that the material of any multiloop "single" element, such as $Q^{m,\text{single}}_{i+1,d}$ or $Q^{ms,\text{single}}_{d+1,j-1}$, can be determined by checking the material of the first or last nucleotide of the subsequence spanned by that element; the first and last nucleotides are each guaranteed to appear in the multiloop as either an unpaired nucleotide or as part of a terminal base pair defining the multiloop. In this instance, the material of $Q^{m,\text{single}}_{i+1,\overline{d}}$ is checked at $i+1$, while the material of $Q^{ms,\text{single}}_{\overline{d}+1,j-1}$ is checked at $j-1$, since these indices are constant over vector $\overline{d}$ and apply to all $d$ and $d+1$ in $\overline{d}$. $M_{\text{single}}$ and $M_{\text{mix}}$ are both added to $R^b_{\text{INTRA}}(i,j,\phi)$ to incorporate all single-material and mixed-material multiloops that include terminal base pair $i \cdot j$. The index limits in the recursion equation reflect the fact that steric effects prevent a multiloop with $j-i<11$ due to the steric requirement that there be at least 3 intervening bases between $i+1$ and $d$ and at least 3 intervening bases between $d+1$ and $j-1$.

- *Single- and mixed-material recursion energies.* In recursion diagrams, shading corresponds to three recursion energies: 1) the penalty for formation of a multiloop $\Delta G^{\text{multi}}_{\text{init}}$, 2) the sequence-independent penalty for a terminal base pair in a multiloop $\Delta G^{\text{multi}}_{\text{bp}}$ (corresponding to the sole base pair $i \cdot j$ that is fully defined in this recursion), 3) the sequence-dependent penalty for a terminal base pair in a multiloop context, $\Delta G^{\text{terminalbp}}_{j,i}(\phi)$ (note that the indices are ordered $j$ then $i$ to reflect $5'$ to $3'$ from the perspective of the multiloop). For $M_{\text{single}}$ (single-material multiloops checked using the conditional $\mu_j = \mu_{j-1} = \mu_{i+1} = \mu_i$), the single-material multiloop initiation parameter, $\Delta G^{\text{multi}}_{\text{init},\mu_i}$, is used and the single-material terminal base pair parameter, $\Delta G^{\text{multi}}_{\text{bp},\mu_i}$, is used for terminal base pair $i \cdot j$. For $M_{\text{mix}}$, the mixed-material multiloop initiation parameter, $\Delta G^{\text{multi}}_{\text{init,mix}}$, is used and the terminal base pair parameter, $\Delta G^{\text{multi}}_{\text{bp},\sigma}$, depends on the material of closing base pair $i \cdot j$ and is checked with the $\sigma$ conditional ($\Delta G^{\text{multi}}_{\text{bp},\mu_i}$ is used if $\mu_j = \mu_i$, and $\Delta G^{\text{multi}}_{\text{bp,mix}}$ is used otherwise). The sequence-dependent terminal base pair parameter, $\Delta G^{\text{terminalbp}}_{j,i}(\phi)$, is determined by lookup based on the sequence and material of nucleotides $j$ and $i$ (note that the indices are ordered $j$ then $i$ to reflect $5'$ to $3'$ from the perspective of the multiloop).

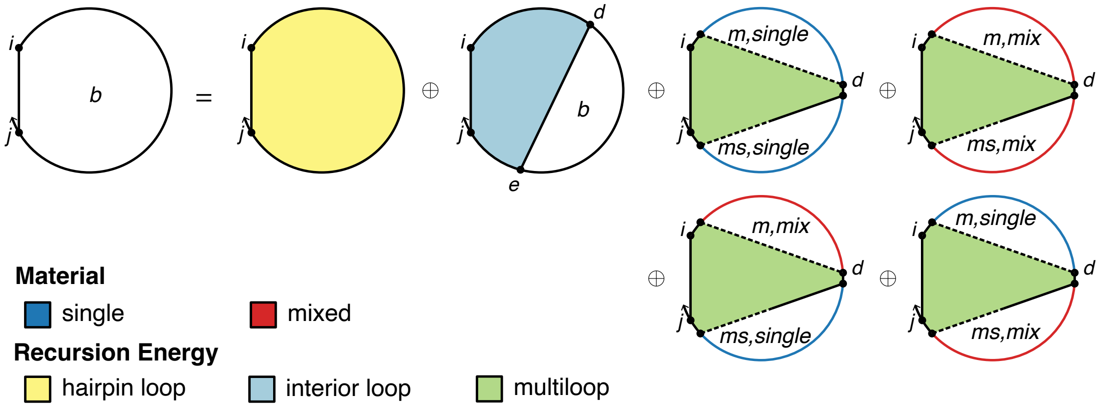

$$R^{b}_{\textsc{Intra}}(i,j,\phi) \equiv \begin{cases} C_{\text{all}}, & \textsc{Complementary}(\phi_i,\phi_j) \\ \mathbb{0}, & \text{otherwise} \end{cases}$$

$$\text{where } C_{\text{all}} \equiv \begin{cases} W(\Delta G^{\text{hairpin}}_{i,j}(\phi)), & j-i \geq 4 \\ \mathbb{0}, & \text{otherwise} \end{cases}$$

$$\oplus \begin{cases} \textsc{InteriorIntra}(i,j,\phi), & j-1 \geq 6 \\ \mathbb{0}, & \text{otherwise} \end{cases}$$

$$\oplus \begin{cases} M_{\text{single}} \oplus M_{\text{mix}}, & j-i \geq 11 \\ \mathbb{0}, & \text{otherwise} \end{cases}$$

$$M_{\text{single}} \equiv C_{0s} \otimes W(\Delta G^{\text{multi}}_{\text{init},\mu_i} + \Delta G^{\text{multi}}_{\text{bp},\mu_i} + \Delta G^{\text{terminalbp}}_{j,i}(\phi))$$

$$C_{0s} \equiv \begin{cases} C_{1s}, & \mu_j = \mu_{j-1} = \mu_{i+1} = \mu_i \\ \mathbb{0}, & \text{otherwise} \end{cases}$$

$$C_{1s} \equiv \text{DOT}\left(Q^{m,\text{single}}_{i+1,\bar{d}}, Q^{ms,\text{single}}_{\bar{d}+1,j-1}\right)$$

$$M_{\text{mix}} \equiv (C_{0m} \oplus C_{1m}) \otimes W(\Delta G^{\text{multi}}_{\text{init,mix}} + \Delta G^{\text{multi}}_{\text{bp},\sigma} + \Delta G^{\text{terminalbp}}_{j,i}(\phi))$$

$$C_{0m} \equiv \begin{cases} \mathbb{0}, & \mu_j = \mu_{j-1} = \mu_{i+1} = \mu_i \\ C_{1s}, & \text{otherwise} \end{cases}$$

$$C_{1m} \equiv \text{DOT}\left(Q^{m,\text{mix}}_{i+1,\bar{d}}, Q^{ms,\text{mix}}_{\bar{d}+1,j-1}\right) \oplus \text{DOT}\left(Q^{m,\text{mix}}_{i+1,\bar{d}}, Q^{ms,\text{single}}_{\bar{d}+1,j-1}\right) \oplus \text{DOT}\left(Q^{m,\text{single}}_{i+1,\bar{d}}, Q^{ms,\text{mix}}_{\bar{d}+1,j-1}\right)$$

$$\sigma \equiv \begin{cases} \mu_i, & \mu_j = \mu_i \\ \text{mix}, & \text{otherwise} \end{cases}$$

$$\text{with } \bar{d} \equiv [i+5 : j-6]$$

Figure S8: $R^{b}_{\textsc{Intra}}$ recursion without coaxial and dangle stacking. Top: recursion diagram. Bottom: recursion equation.

$\boldsymbol{R^{ms}_{\text{INTRA}}}$ **recursion without coaxial and dangle stacking.** The $R^b_{\text{INTRA}}(i,j,\phi)$ recursion references $Q^{ms}$ elements that are computed using the $R^{ms}_{\text{INTRA}}(i,j,\phi)$ recursion shown in Figure S9. $R^{ms}_{\text{INTRA}}(i,j,\phi)$ operates on a conditional ensemble for subsequence $[i,j]$ in a multiloop context containing one terminal base pair starting at $i$ and ending in the interval $[i+1,j]$ (depicted as a half-solid/half-dashed line between $i$ and $j$). A terminal base pair is either a single material (denoted in the recursion diagram by a blue base pair, if the material of the two nucleotides matches) or mixed material (denoted by a red base pair, if the material of the two nucleotides does not match). Like other multiloop recursions in the mixed algorithm, $R^{ms}_{\text{INTRA}}(i,j,\phi)$ is split into $R^{ms,\text{single}}_{\text{INTRA}}(i,j,\phi)$ and $R^{ms,\text{mix}}_{\text{INTRA}}(i,j,\phi)$ (the single and mixed recursions are indicated in the diagram with blue and red backbones, respectively):

- $R^{ms,\text{single}}_{\text{INTRA}}(i,j,\phi)$: This recursion treats the $R^{ms}_{\text{INTRA}}(i,j,\phi)$ conditional ensemble over $[i,j]$ in a multiloop context where terminal base pair $i \cdot d$ and any unpaired nucleotides in the interval $[d+1,j]$ are of a single material. For the edge case where $d=j$, then the whole region $[i,j]$ enclosed by terminal base pair $i \cdot j$ is incorporated using a $Q^b_{i,j}$ element in term $C_2$. This represents a single-material contribution only if $\mu_j = \mu_i$. If the conditional is satisfied, the contribution is added to $R^{ms,\text{single}}_{\text{INTRA}}(i,j,\phi)$ using term $C_{0s}$. Otherwise, if the conditional fails, the contribution is added to $R^{ms,\text{mix}}_{\text{INTRA}}(i,j,\phi)$ using term $C_{0m}$. For the base case where $d$ is in $[i+4,j-1]$, the contribution of terminal base pair $i \cdot d$ is incorporated using $Q^b_{i,d}$ if $\mu_d = \mu_i$ (indicating a single-material terminal base pair), while the unpaired region $[d+1,j]$ is incorporated using $Q^{ma,\text{single}}_{d+1,j}$. The resulting contribution $Q^b_{i,d} \otimes Q^{ma,\text{single}}_{d+1,j}$ of term $C_{1s}$ produces a single-material multiloop only if the material of the terminal base pair $i \cdot d$ matches that of $Q^{ma,\text{single}}_{d+1,j}$, which is checked with conditional $\mu_j = \mu_i$. If the conditional is satisfied, the contribution is added to $R^{ms,\text{single}}_{\text{INTRA}}(i,j,\phi)$ using term $C_{0s}$. Otherwise, if the conditional fails, the contribution is added to $R^{ms,\text{mix}}_{\text{INTRA}}(i,j,\phi)$ using term $C_{0m}$. In the conditional, indices $i$ and $j$ are used to check the material since they are constant over vector $\overline{d}$ and apply to all $d$ and $d+1$ in $\overline{d}$.
- $R^{ms,\text{mix}}_{\text{INTRA}}(i,j,\phi)$: This recursion treats the $R^{ms}_{\text{INTRA}}(i,j,\phi)$ conditional ensemble over $[i,j]$ in a multiloop context where terminal base pair $i \cdot d$ and any unpaired nucleotides overall contain mixed materials. For the edge case where $d=j$, then the whole region $[i,j]$ enclosed by terminal base pair $i \cdot j$ is incorporated using a $Q^b_{i,j}$ element in term $C_2$. As noted above, this represents a mixed-material contribution only if $\mu_j \neq \mu_i$ and is incorporated in $R^{ms,\text{mix}}_{\text{INTRA}}(i,j,\phi)$ using term $C_{0m}$. For the base case where $d$ is in $[i+4,j-1]$, the contribution for the subsequence $[i,d]$ enclosed by terminal base pair $i \cdot d$ is incorporated using a $Q^b_{i,d}$ element, which represents a single-material contribution if $\mu_d = \mu_i$ and a mixed-material contribution otherwise. The contribution for the unpaired region $[d+1,j]$ is incorporated using either $Q^{ma,\text{single}}_{d+1,j}$ or $Q^{ma,\text{mix}}_{d+1,j}$. There are four possible single/mix combinations for $Q^b_{i,d} \otimes Q^{ma}_{d+1,j}$. The three combinations containing at least one mixed element must represent mixed contributions and are included in $R^{ms,\text{mix}}_{\text{INTRA}}(i,j,\phi)$ using term $C_{1m}$. As noted above, the remaining combination calculated in term $C_{1s}$ represents a mixed contribution if $\mu_j \neq \mu_i$, and is incorporated in term $R^{ms,\text{mix}}_{\text{INTRA}}(i,j,\phi)$ using term $C_{0m}$.

Shading corresponds to two recursion energies: 1) the sequence-independent penalty for a terminal base pair in a multiloop (for base pair $i \cdot d$, $\Delta G^{\text{multi}}_{\text{bp},\mu_i}$ if $\mu_d = \mu_i$, and $\Delta G^{\text{multi}}_{\text{bp,mix}}$ otherwise; for base pair $i \cdot j$, $\Delta G^{\text{multi}}_{\text{bp},\sigma}$, where the parameter depends on the material as determined by the $\sigma$ conditional), 2) the sequence-dependent penalty for a terminal base pair in a multiloop context (for base pair $i \cdot d$, $\Delta G^{\text{terminalbp}}_{i,d}(\phi)$, dependent on the sequence and material of base pair $i \cdot d$; for base pair $i \cdot j$, $\Delta G^{\text{terminalbp}}_{i,j}(\phi)$, dependent on the sequence and material of base pair $i \cdot j$). The index limits in the recursion equation reflect the steric requirement that there be at least 3 intervening bases between $i$ and $d$. Note that $R^{ms}$ serves as an efficiency wrapper for $R^b$ in the multiloop context in a completely analogous manner to $R^s$ serving as an efficiency wrapper for $R^b$ in an exterior loop context, with $R^b$ representing the $3'$-most terminal base pair in either context.

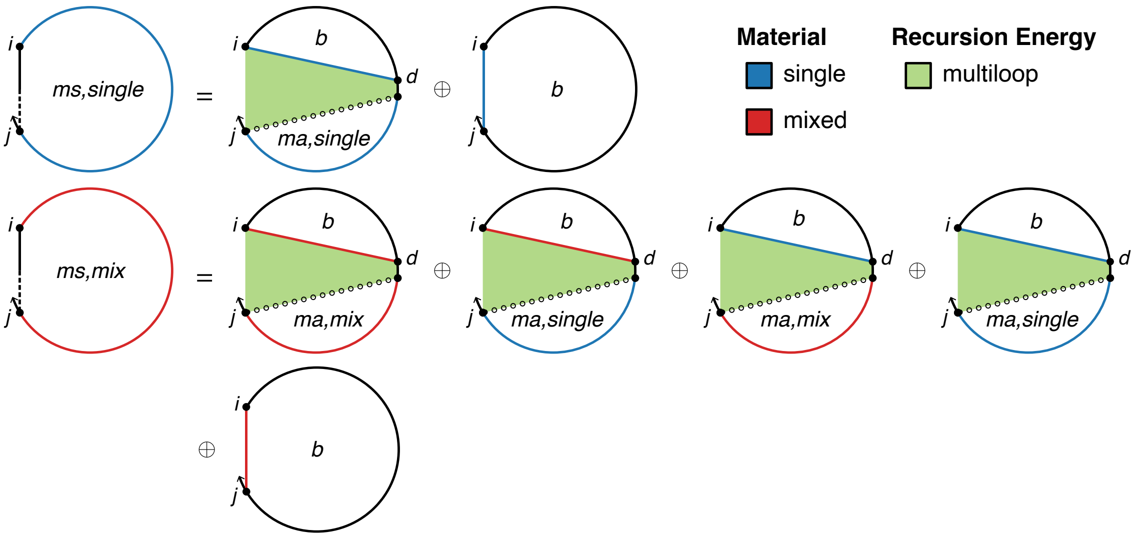

$$R^{ms,\text{single}}_{\text{INTRA}}(i,j,\phi) \equiv C_{0s}$$

$$C_{0s} \equiv \begin{cases} C_{1s} \oplus C_2, & \mu_j = \mu_i \\ \mathbb{0}, & \text{otherwise} \end{cases}$$

$$C_{1s} \equiv \begin{cases} \text{DOT}\left(Q^b_{i,\overline{d}}, Q^{ma,\text{single}}_{\overline{d}+1,j}, W(\Delta G^{\text{multi}}_{\text{bp},\mu_i} + \Delta G^{\text{terminalbp}}_{i,\overline{d}}(\phi))\right), & j-i \geq 5 \text{ and } \mu_d = \mu_i \\ \mathbb{0}, & \text{otherwise} \end{cases}$$

$$C_2 \equiv \begin{cases} Q^b_{i,j} \otimes W(\Delta G^{\text{multi}}_{\text{bp},\sigma} + \Delta G^{\text{terminalbp}}_{i,j}(\phi)), & j-i \geq 4 \\ \mathbb{0}, & \text{otherwise} \end{cases}$$

$$R^{ms,\text{mix}}_{\text{INTRA}}(i,j,\phi) \equiv C_{0m} \oplus C_{1m}$$

$$C_{0m} \equiv \begin{cases} \mathbb{0}, & \mu_j = \mu_i \\ C_{1s} \oplus C_2, & \text{otherwise} \end{cases}$$

$$C_{1m} \equiv \begin{cases} \text{DOT}\left(Q^b_{i,\overline{d}}, (Q^{ma,\text{mix}}_{\overline{d}+1,j} \oplus Q^{ma,\text{single}}_{\overline{d}+1,j}), W(\Delta G^{\text{multi}}_{\text{bp,mix}} + \Delta G^{\text{terminalbp}}_{i,\overline{d}}(\phi))\right), & j-i \geq 5 \text{ and } \mu_d \neq \mu_i \\ \mathbb{0}, & \text{otherwise} \end{cases}$$

$$\oplus \begin{cases} \text{DOT}\left(Q^b_{i,\overline{d}}, Q^{ma,\text{mix}}_{\overline{d}+1,j}, W(\Delta G^{\text{multi}}_{\text{bp},\mu_i} + \Delta G^{\text{terminalbp}}_{i,\overline{d}}(\phi))\right), & j-i \geq 5 \text{ and } \mu_d = \mu_i \\ \mathbb{0}, & \text{otherwise} \end{cases}$$

$$\text{with } \sigma \equiv \begin{cases} \mu_i, & \mu_j = \mu_i \\ \text{mix}, & \text{otherwise} \end{cases}$$

$$\overline{d} \equiv [i+4 : j-1]$$

Figure S9: $R^{ms}_{\text{INTRA}}$ recursion without coaxial and dangle stacking. Top: recursion diagram. Bottom: recursion equation.

$\boldsymbol{R^{ma}_{\text{INTRA}}}$ **recursion without coaxial and dangle stacking.** The $R^{b}_{\text{INTRA}}(i,j,\phi)$ and $R^{ms}_{\text{INTRA}}(i,j,\phi)$ recursions reference $Q^{ma}$ elements that are computed using the $R^{ma}_{\text{INTRA}}$ recursion shown in Figure S10. $R^{ma}_{\text{INTRA}}(i,j,\phi)$ treats the conditional ensemble of subsequence $[i,j]$ in a multiloop context containing no terminal base pairs (the hollow dotted line between $i$ and $j$ indicates that there is no base pairing in $[i,j]$). This recursion is used in the mixed-material algorithm to switch the $\Delta G^{\text{multi}}_{\text{nt}}$ parameter depending on the material of each nucleotide. Note that it is identical to $R^{ma}_{\text{INTRA}}(i,j,\phi)$ in the stacking ensemble (Figure S27). The single-material algorithm does not incorporate this recursion since parameter switching is not necessary. Like other multiloop recursions in the mixed algorithm, $R^{ma}_{\text{INTRA}}(i,j,\phi)$ is split into $R^{ma,\text{single}}_{\text{INTRA}}(i,j,\phi)$ and $R^{ma,\text{mix}}_{\text{INTRA}}(i,j,\phi)$ (the single- and mixed-material recursions are indicated in the recursion diagram by blue and red backbones, respectively):

- $R^{ma,\text{single}}_{\text{INTRA}}(i,j,\phi)$: This recursion calculates contributions from subsequences $[i,j]$ composed of a single material. The shading corresponds to the energy per unpaired nucleotide in a multiloop context for the material of nucleotide $j$, $\Delta G^{\text{multi}}_{\text{nt},\mu_j}$. For the edge case with $i=j$, then the subsequence is a single nucleotide and therefore a single material, yielding a contribution $W(\Delta G^{\text{multi}}_{\text{nt},\mu_i})$ in term $C_{2s}$. For the base case with $i<j$, the $Q^{ma,\text{single}}_{i,j-1}$ element is used to represent the single-material contribution from subsequence $[i,j-1]$ (non-zero only if $[i,j-1]$ is a single material) and the contribution for unpaired nucleotide $j$ is $W(\Delta G^{\text{multi}}_{\text{nt},\mu_j})$. The overall contribution, $Q^{ma,\text{single}}_{i,j-1}\otimes W(\Delta G^{\text{multi}}_{\text{nt},\mu_j})$, represents a single material only if the materials of both terms match, which is checked with the conditional $\mu_j=\mu_{j-1}$. If the conditional is satisfied, the contribution is added to $R^{ma,\text{single}}_{\text{INTRA}}(i,j,\phi)$ using term $C_{0s}$. Otherwise, if the conditional fails, the contribution is added to $R^{ma,\text{mix}}_{\text{INTRA}}(i,j,\phi)$ using term $C_{0m}$.

- $R^{ma,\text{mix}}_{\text{INTRA}}(i,j,\phi)$: This recursion calculates contributions from subsequence $[i,j]$ composed of mixed materials. The shading corresponds to the energy per unpaired nucleotide in a multiloop context for the material of nucleotide $j$, $\Delta G^{\text{multi}}_{\text{nt},\mu_j}$. For the edge case with $i=j$, the contribution is $\mathbb{0}$ since a single nucleotide cannot have more than one material. For the base case with $i<j$, the $Q^{ma,\text{mix}}_{i,j-1}$ element is used to represent the mixed-material contribution of subsequence $[i,j-1]$ (non-zero only if $[i,j-1]$ contains a mixture of materials) and the contribution for unpaired nucleotide $j$ is $W(\Delta G^{\text{multi}}_{\text{nt},\mu_j})$, leading to an overall contribution of $Q^{ma,\text{mix}}_{i,j-1}\otimes W(\Delta G^{\text{multi}}_{\text{nt},\mu_j})$ added to $R^{ma,\text{mix}}_{\text{INTRA}}(i,j,\phi)$ using term $C_{1m}$. As noted above, the contribution, $Q^{ma,\text{single}}_{i,j-1}\otimes W(\Delta G^{\text{multi}}_{\text{nt},\mu_j})$, calculated in term $C_{1s}$ represents a mixed contribution if $\mu_j\neq\mu_{j-1}$, and is incorporated in $R^{ma,\text{mix}}_{\text{INTRA}}(i,j,\phi)$ using term $C_{0m}$.

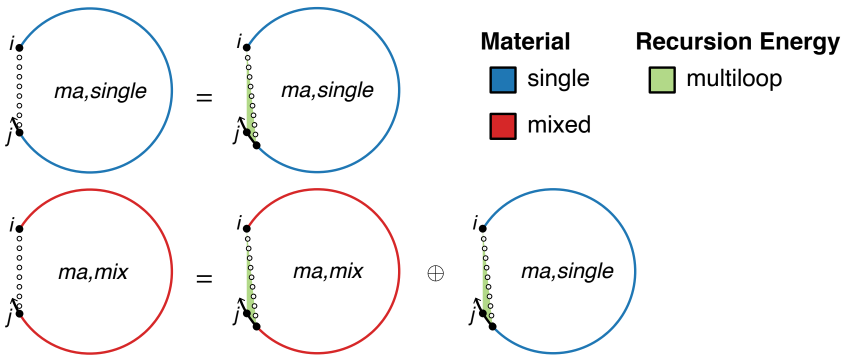

$$R^{ma,\text{single}}_{\text{INTRA}}(i,j,\phi) \equiv C_{0s} \oplus C_{2s}$$

$$C_{0s} \equiv \begin{cases} C_{1s}, & \mu_j = \mu_{j-1} \\ \mathbb{0}, & \text{otherwise} \end{cases}$$

$$C_{1s} \equiv \begin{cases} Q^{ma,\text{single}}_{i,j-1} \otimes W(\Delta G^{\text{multi}}_{\text{nt},\mu_i}), & i < j \\ \mathbb{0}, & \text{otherwise} \end{cases}$$

$$C_{2s} \equiv \begin{cases} W(\Delta G^{\text{multi}}_{\text{nt},\mu_i}), & i = j \\ \mathbb{0}, & \text{otherwise} \end{cases}$$

$$R^{ma,\text{mix}}_{\text{INTRA}}(i,j,\phi) \equiv C_{0m} \oplus C_{1m}$$

$$C_{0m} \equiv \begin{cases} \mathbb{0}, & \mu_j = \mu_{j-1} \\ C_{1s}, & \text{otherwise} \end{cases}$$

$$C_{1m} \equiv \begin{cases} Q^{ma,\text{mix}}_{i,j-1} \otimes W(\Delta G^{\text{multi}}_{\text{nt},\mu_j}), & i < j \\ \mathbb{0}, & \text{otherwise} \end{cases}$$

Figure S10: $R^{ma}_{\text{INTRA}}(i,j,\phi)$ recursion without coaxial and dangle stacking. Top: recursion diagram. Bottom: recursion equation. Note that this recursion is the same regardless of whether coaxial and dangle stacking are included in the structural ensemble.

**$R^m_{\text{INTRA}}$ recursion without coaxial and dangle stacking.** The $R^b_{\text{INTRA}}(i,j,\phi)$ recursion references $Q^m$ elements that are computed using the $R^m_{\text{INTRA}}$ recursion shown in Figure S11. $R^m_{\text{INTRA}}(i,j,\phi)$ operates on a conditional ensemble for subsequence $[i,j]$ in a multiloop context where $i$ and $j$ may or may not be paired (depicted with a dashed line between $i$ and $j$ in the recursion diagram) and where there is at least one terminal base pair. Like other multiloop recursions in the mixed algorithm, $R^m_{\text{INTRA}}(i,j,\phi)$ is split into $R^{m,\text{single}}_{\text{INTRA}}(i,j,\phi)$ and $R^{m,\text{mix}}_{\text{INTRA}}(i,j,\phi)$ (the single and mixed recursions are indicated in the diagram with blue and red backbones, respectively):

- $R^{m,\text{single}}_{\text{INTRA}}(i,j,\phi)$: This recursion operates over the $R^m_{\text{INTRA}}(i,j,\phi)$ conditional ensemble on $[i,j]$ in a multiloop context where any terminal base pair or unpaired nucleotide is of a single material. This recursion distinguishes two cases that are combined using $\oplus$ in the recursion equation:
  - *One terminal base pair*: the case where there is exactly one terminal base pair in subsequence $[i,j]$ in a multiloop context. This terminal base pair starts at $d$ and ends in the interval $[d+1,j]$ (depicted by a straight half-solid/half dashed line between $d$ and $j$). Subsequence $[i,d-1]$ is an unpaired region (depicted by a hollow dotted line between $i$ and $d-1$). For the edge case where $d=i$, then the whole region $[i,j]$ is incorporated using a $Q^{ms,\text{single}}_{i,j}$ element with term $C_{3s}$. No further material checks are needed since the $R^{ms,\text{single}}_{\text{INTRA}}(i,j,\phi)$ recursion only includes single-material contributions. For the base case where $d$ is between $i+1$ and $j-4$, then the unpaired region $[i,d-1]$ is incorporated by a $Q^{ma,\text{single}}_{i,d-1}$ element while the region $[d,j]$ is incorporated by $Q^{ms,\text{single}}_{d,j}$. The resulting contribution $Q^{ma,\text{single}}_{i,d-1} \otimes Q^{ms,\text{single}}_{d,j}$ of term $C_{s1}$ produces a single-material multiloop only if the material matches for both elements, which is checked with the conditional $\mu_j = \mu_i$. If the conditional is satisfied, the contribution is added to $R^{m,\text{single}}_{\text{INTRA}}(i,j,\phi)$ using term $C_{0s}$. Otherwise, if the conditional fails, the contribution is added to $R^{m,\text{mix}}_{\text{INTRA}}(i,j,\phi)$ using term $C_{0m}$. In the conditional, indices $i$ and $j$ are used to check the material since they are constant over vector $\overline{d}$ and apply to all $d-1$ and $d$ in $\overline{d}$.
  - *More than one terminal base pair*: the case where there are two or more terminal base pairs in subsequence $[i,j]$ in a multiloop context. The $3'$-most terminal base pair starts at $e+1$ and ends in the interval $[e+2,j]$ (depicted by a straight half-solid/half dashed line between $e+1$ and $j$); the contribution of subsequence $[e+1,j]$ is incorporated by element $Q^{ms,\text{single}}_{e+1,j}$. There are one or more additional terminal base pairs in the interval $[i,e]$ (the straight dashed line denotes that $i$ and $e$ may or may not be paired); the contribution of subsequence $[i,e]$ is incorporated by element $Q^{m,\text{single}}_{i,e}$. The resulting contribution $Q^{m,\text{single}}_{i,e} \otimes Q^{ms,\text{single}}_{e+1,j}$ of term $C_{2s}$ produces a single-material multiloop only if the material matches for both elements, which is checked with the conditional $\mu_j = \mu_i$. If the conditional is satisfied, the contribution is added to $R^{m,\text{single}}_{\text{INTRA}}(i,j,\phi)$ using term $C_{0s}$. Otherwise, if the conditional fails, the contribution is added to $R^{m,\text{mix}}_{\text{INTRA}}(i,j,\phi)$ using term $C_{0m}$. In the conditional, indices $i$ and $j$ are used to check the material since they are constant over vector $\overline{e}$ and apply to all $e$ and $e+1$ in $\overline{e}$.

- $R^{m,\text{mix}}_{\text{INTRA}}(i,j,\phi)$: This recursion operates over the $R^m_{\text{INTRA}}(i,j,\phi)$ conditional ensemble on $[i,j]$ in a multiloop context where any terminal base pair and unpaired nucleotides overall contain a mixture of materials. This recursion distinguishes two cases that are combined using $\oplus$ in the recursion equation:
  - *One terminal base pair*: the case where there is exactly one terminal base pair in subsequence $[i,j]$ in a multiloop context. This terminal base pair starts at $d$ and ends in the interval $[d+1,j]$ (depicted by a straight half-solid/half dashed line between $d$ and $j$). $[i,d-1]$ is an unpaired region (depicted by a hollow dotted line between $i$ and $d-1$). For the edge case where $d=i$, then whole region $[i,j]$ is incorporated using a $Q^{ms,\text{mix}}_{i,j}$ element with term $C_{3m}$. No further material checks are needed since the $R^{ms,\text{mix}}_{\text{INTRA}}(i,j,\phi)$ recursion only includes mixed-material contributions. For the base case where $d$ is between $i+1$ and $j-4$, then the unpaired region $[i,d-1]$ is incorporated by a $Q^{ma}_{i,d-1}$ element while the region $[d,j]$ is incorporated by a $Q^{ms}_{d,j}$ element. Because $Q^{ma}_{i,d-1}$ and $Q^{ms}_{d,j}$ are each spit into single- and mixed-material recursions, there are 4 possible combinations when multiplying them. The three combinations containing at least one mixed element ($Q^{ma,\text{single}}_{i,d-1} \otimes Q^{ms,\text{mix}}_{d,j}$, $Q^{ma,\text{mix}}_{i,d-1} \otimes Q^{ms,\text{single}}_{d,j}$, $Q^{ma,\text{mix}}_{i,d-1} \otimes Q^{ms,\text{mix}}_{d,j}$) must represent mixed contributions, and are included in $R^{m,\text{mix}}_{\text{INTRA}}(i,j,\phi)$ using term $C_{1m}$. As noted above, the remaining combination, $Q^{ma,\text{single}}_{i,d-1} \otimes Q^{ms,\text{single}}_{d,j}$ calculated in term $C_{1s}$, only represents a mixed contribution if $\mu_j \neq \mu_i$, and is incorporated in $R^{m,\text{mix}}_{\text{INTRA}}(i,j,\phi)$ using term $C_{0m}$.
  - *More than one terminal base pair*: the case where there are two or more terminal base pairs in subsequence $[i,j]$ in a multiloop context. The $3'$-most terminal base pair starts at $e+1$ and ends in the interval $[e+2,j]$ (depicted by a straight half-solid/half dashed line between $e+1$ and $j$); the contribution of subsequence $[e+1,j]$ is incorporated by a $Q^{ms}_{e+1,j}$ element. There are one or more additional terminal base pairs in the interval $[i,e]$ (the straight dashed line denotes that $i$ and $e$ may or may not be paired); the contribution of subsequence $[i,e]$ is incorporated by a $Q^m_{i,e}$ element. Because $Q^m_{i,e}$ and $Q^{ms}_{e+1,j}$ are each spit into single- and mixed-material recursions, there are 4

possible combinations when multiplying them. The three combinations containing at least one mixed element ($Q^{m,\text{single}}_{i,e} \otimes Q^{ms,\text{mix}}_{e+1,j}$, $Q^{m,\text{mix}}_{i,e} \otimes Q^{ms,\text{single}}_{e+1,j}$, $Q^{m,\text{mix}}_{i,e} \otimes Q^{ms,\text{mix}}_{e+1,j}$) must represent mixed contributions, and are included in $R^{m,\text{mix}}_{\text{INTRA}}(i, j, \phi)$ using term $C_{2m}$. As noted above, the remaining combination, $Q^{m,\text{single}}_{i,e} \otimes Q^{ms,\text{single}}_{e+1,j}$ calculated in term $C_{2s}$, only represents a mixed contribution if $\mu_j \neq \mu_i$, and is incorporated in $R^{m,\text{mix}}_{\text{INTRA}}(i, j, \phi)$ using term $C_{0m}$.

Shading in the recursion diagram does not represent any recursion energies because all multiloop contribtributions are handled by other recursions: 1) there are no terminal base pairs in a multiloop context explicitly defined in this case, 2) there are no unpaired bases in a multiloop context explicitly defined in this case. The index limits in the recursion equation reflect the steric requirement that there be at least 3 intervening bases between $d$ and $j$. The index limits in the recursion equation reflect the steric requirement that there be at least 3 intervening bases between $i$ and $e$ and at least 3 intervening bases between $e + 1$ and $j$.

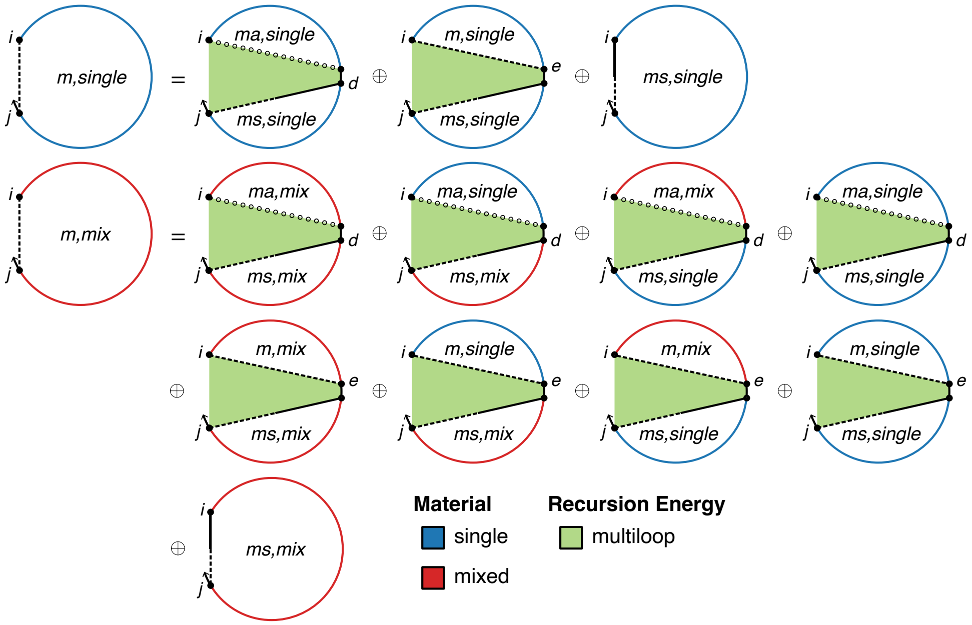

$$R^{m,\text{single}}_{\text{INTRA}}(i,j,\phi) \equiv C_{0s} \oplus C_{3s}$$

$$C_{0s} \equiv \begin{cases} C_{1s} \oplus C_{2s}, & \mu_j = \mu_i \\ \mathbb{0}, & \text{otherwise} \end{cases}$$

$$C_{1s} \equiv \begin{cases} \text{DOT}\left(Q^{ma,\text{single}}_{i,\overline{d}-1}, Q^{ms,\text{single}}_{\overline{d},j}\right), & j-i \geq 5 \\ \mathbb{0}, & \text{otherwise} \end{cases}$$

$$C_{2s} \equiv \begin{cases} \text{DOT}\left(Q^{m,\text{single}}_{i,\overline{e}}, Q^{ms,\text{single}}_{\overline{e}+1,j}\right), & j-i \geq 9 \\ \mathbb{0}, & \text{otherwise} \end{cases}$$

$$C_{3s} \equiv \begin{cases} Q^{ms,\text{single}}_{i,j}, & j-i \geq 4 \\ \mathbb{0}, & \text{otherwise} \end{cases}$$

$$R^{m,\text{mix}}_{\text{INTRA}}(i,j,\phi) \equiv C_{0m} \oplus C_{1m} \oplus C_{2m} \oplus C_{3m}$$

$$C_{0m} \equiv \begin{cases} \mathbb{0}, & \mu_j = \mu_i \\ C_{1s} \oplus C_{2s}, & \text{otherwise} \end{cases}$$

$$C_{1m} \equiv \begin{cases} \text{DOT}\left(Q^{ma,\text{mix}}_{i,\overline{d}-1}, Q^{ms,\text{mix}}_{\overline{d},j}\right) \oplus \text{DOT}\left(Q^{ma,\text{single}}_{i,\overline{d}-1}, Q^{ms,\text{mix}}_{\overline{d},j}\right) \oplus \text{DOT}\left(Q^{ma,\text{mix}}_{i,\overline{d}-1}, Q^{ms,\text{single}}_{\overline{d},j}\right), & j-i \geq 5 \\ \mathbb{0}, & \text{otherwise} \end{cases}$$

$$C_{2m} \equiv \begin{cases} \text{DOT}\left(Q^{m,\text{mix}}_{i,\overline{e}}, Q^{ms,\text{mix}}_{\overline{e}+1,j}\right) \oplus \text{DOT}\left(Q^{m,\text{single}}_{i,\overline{e}}, Q^{ms,\text{mix}}_{\overline{e}+1,j}\right) \oplus \text{DOT}\left(Q^{m,\text{mix}}_{i,\overline{e}}, Q^{ms,\text{single}}_{\overline{e}+1,j}\right), & j-i \geq 9 \\ \mathbb{0}, & \text{otherwise} \end{cases}$$

$$C_{3m} \equiv \begin{cases} Q^{ms,\text{mix}}_{i,j}, & j-i \geq 4 \\ \mathbb{0}, & \text{otherwise} \end{cases}$$

with $\overline{d} \equiv [i+1:j-4], \quad \overline{e} \equiv [i+4:j-5]$

Figure S11: $R^{m}_{\text{INTRA}}$ recursion without coaxial and dangle stacking. Top: recursion diagram. Bottom: recursion equation.

##### S6.3.2 Interstrand dynamic programming recursions without coaxial and dangle stacking

Here, we consider recursions for calculating the entries in a rectangular interstrand block without coaxial and dangle stacking. By definition, interstrand blocks involve 2 or more strands, and hence one or more nicks between strands. For a given interstrand block, $\eta$ stores an array of nick indices between strands within the block, with each nick denoted by the index of the nucleotide following the nick. If $m \equiv \text{FIRST}(\eta)$ and $n \equiv \text{LAST}(\eta)$, then for subsequence $[i, j]$ corresponding to element $i, j$ in the interstrand block, we have by definition $i < m$ (nucleotide $i$ is on the first strand in the block) and $j \geq n$ (nucleotide $j$ is on the last strand in the block).

**$R^{\varnothing}_{\text{INTER}}$ recursion without coaxial and dangle stacking.** The recursion $R^{\varnothing}_{\text{INTER}}(i, j, \phi)$, shown in Figure S12, does not change in the mixed material setting and is as previously described.[1] $R^{\varnothing}_{\text{INTER}}(i, j, \phi)$ operates on the unconstrained ensemble for subsequence $[i, j]$ with $i$ and $j$ on different strands in an exterior loop context where $i$ and $j$ may or may not be paired (depicted with a dashed line between $i$ and $j$ in the recursion diagram). Unlike $R^{\varnothing}_{\text{INTRA}}(i, j, \phi)$, there is no empty case because this would correspond to a disconnected structure (which is not in the multistranded ensemble) due to the presence of one or more nicks between $i$ and $j$. Hence, the only case is *at least one terminal base pair:* the non-empty case in an exterior loop context where there is at least one terminal base pair (i.e., a base pair terminating a duplex) in subsequence $[i, j]$. The $3'$-most terminal base pair begins at $d+1$ and ends in the interval $[d+2, j]$ (depicted using a half-solid/half-dashed line in the recursion diagram). The contributions for subsequence $[d+1, j]$ are incorporated using a $Q^s_{d+1,j}$ element. Contributions for the remaining subsequence $[i, d]$ are incorporated by a $Q^{\varnothing}_{i,d}$ element. The shading denotes the recursion energy 0 corresponding to the zero reference state in an exterior loop context. Note that the recursion energy $\Delta G^{\text{terminalbp}}(\phi)$ representing one component of the $\Delta G^{\text{exterior}}_{i,j}(\phi)$ free energy is not incorporated here because the full identity of the terminal base pair (i.e., a base pair terminating a duplex) beginning at $d+1$ is not known within the $R^{\varnothing}_{\text{INTER}}(i, j, \phi)$ recursion (only within the $R^s(d+1, j, \phi)$ recursion). The edge case where the index $d+1 = i$ is displayed explicitly to indicate that no $Q^{\varnothing}$ element is accessed in this case.

Because there are nicks involved in calculating the elements of interstrand blocks, care must be taken to ensure that no disconnected secondary structures are incorporated in the complex ensemble. For a given interstrand block with nick indices $\eta$, the function VALID returns the set of valid vectorization ranges $\{\overline{d}_1, \overline{d}_2, \ldots\}$, such that for each valid vectorization range, $d$ and $d+1$ are on the same strand (i.e., such that $d$ and $d+1$ do not take on values that would place a nick between them). As is evident from the recursion diagram of Figure S12, if $d$ and $d+1$ were to take on values that placed a nick between them, a disconnected structure would result. There is at most one valid vectorization range per strand, and there may be none for a strand or subsequence that is too short. For each valid vectorization range $\overline{d}$, the resulting dot product

$$\text{DOT}\left(Q^{\varnothing}_{i,\overline{d}}, Q^{s}_{\overline{d}+1,j}\right) \tag{S136}$$

is between the range $\overline{d}$ of row $i$ (depicted as brown elements in Figure S4c) and the range $\overline{d}+1$ of column $j$ (gray elements), yielding element $i, j$ (purple element). Note that Figure S4c depicts two valid vectorization ranges (leading to two dot products that are summed to calculate the purple element); the gap of one element between the two vectorization ranges corresponds to exclusion of the value $d = 3$ which would have placed a nick between nucleotides $d$ and $d+1$ (note that $\eta = 4$ for this interstrand block).

Note that for calculating element $i, j$ in Figure S12, the subsequence submitted to VALID ranges from $i$ to $\max(j-4, n)$,

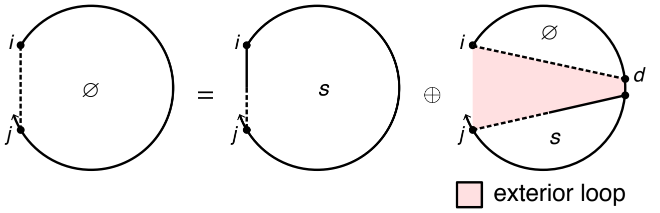

$$R^{\varnothing}_{\text{INTER}}(i, j, \phi) \equiv Q^{s}_{i,j} \oplus \bigoplus_{\overline{d} \in \text{VALID}(i, \max(j-4, n), \eta)} \text{DOT}\left(Q^{\varnothing}_{i,\overline{d}}, Q^{s}_{\overline{d}+1,j}\right) \tag{S135}$$

where $n = \text{LAST}(\eta)$

Figure S12: $R^{\varnothing}_{\text{INTER}}$ recursion without coaxial and dangle stacking. Top: recursion diagram. Bottom: recursion equation.

where $n \equiv \textsc{Last}(\eta)$. This yields two cases:

- If $\max(j-4, n) = j-4$: there is no nick between nucleotide $j-4$ and $j$ (since $n \equiv \textsc{Last}(\eta) < j-4$), so there must be at least 3 intervening bases between $d+1$ and $j$ because steric effects prevent a hairpin loop with fewer than 3 unpaired nucleotides. In this case, each incorporated element $Q^s_{d+1,j}$ results from an $R^s_{\text{INTRA}}(d+1, j, \phi)$ recursion for an intrastrand block.

- If $\max(j-4, n) = n$: there is a nick between nucleotide $j-4$ and $j$ (since $n \geq j-4$), so $d+1$ can be as large as $n-1$ and still pair to any nucleotide in subsequence $[n, j]$. In this case, each incorporated element $Q^s_{d+1,j}$ results from an $R^s_{\text{INTER}}(d+1, j, \phi)$ recursion for an interstrand block.

$\boldsymbol{R^s_{\text{INTER}}}$ **recursion without coaxial and dangle stacking.** The recursion $R^s_{\text{INTER}}(i, j, \phi)$, shown in Figure S13, does not change in the mixed material setting and is as previously described.[1] The $R^{\varnothing}_{\text{INTER}}(i, j, \phi)$ recursion references $Q^s_{d+1,j}$ elements that are computed using either the $R^s_{\text{INTRA}}$ recursion of Figure S7 (if $d{+}1$ and $j$ are on the same strand) or the $R^s_{\text{INTER}}$ recursion of Figure S13 (if $d{+}1$ and $j$ are on different strands). Recursion $R^s_{\text{INTER}}(i, j, \phi)$ operates on a conditional ensemble for subsequence $[i, j]$ with $i$ and $j$ on different strands in an exterior loop context containing one terminal base pair starting at $i$ and ending in the interval $[i+1, j]$ (depicted as a half-solid/half-dashed line between $i$ and $j$). The contribution for the subsequence $[i, d]$ enclosed by base pair $i \cdot d$ is incorporated using a $Q^b_{i,d}$ element. Shading corresponds to the recursion energy, $\Delta G^{\text{terminalbp}}_{i,d}(\phi)$, representing the sequence-dependent penalty for a terminal base pair in an exterior loop context. The index $d$ must always be on the last strand (i.e., $d \geq \textsc{Last}(\eta)$) to ensure there are no strand breaks in the subsequence $[d, j]$, which would correspond to a disconnected structure. Note that the $R^s$ recursion serves as an efficiency wrapper of the $R^b$ recursion (here, representing the 3′-most terminal base pair in an exterior loop context) to reduce the time complexity of the $R^{\varnothing}$ recursion from $O(N^4)$ to $O(N^3)$. This time complexity reduction is achieved by defining the 3′-most terminal base pair using $R^b$ within the $R^s$ efficiency wrapper rather than directly using the $R^b$ recursion within the $R^{\varnothing}$ recursion, so as to avoid introducing a fourth independent index into the $R^{\varnothing}$ recursion.

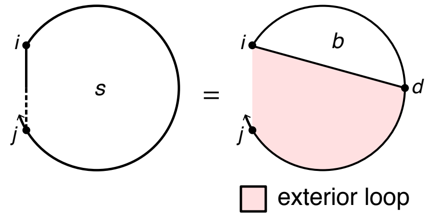

$$R^s_{\text{INTER}}(i, j, \phi) \equiv \text{DOT}\left(Q^b_{i,\overline{d}}, W(\Delta G^{\text{terminalbp}}_{i,\overline{d}}(\phi))\right), \qquad \text{where } \overline{d} \equiv [\textsc{Last}(\eta) : j] \tag{S137}$$

Figure S13: $\boldsymbol{R^s_{\text{INTER}}}$ recursion without coaxial and dangle stacking. Top: recursion diagram. Bottom: recursion equation.

$\boldsymbol{R^b_{\text{INTER}}}$ **recursion without coaxial and dangle stacking.** The $R^s_{\text{INTER}}(i, j, \phi)$ recursion references $Q^b_{i,d}$ elements that are computed using either the $R^b_{\text{INTRA}}$ recursion of Figure S14 (if $i$ and $d$ are on the same strand) or the $R^b_{\text{INTER}}$ recursion of Figure S14 (if $i$ and $d$ are on different strands). $R^b_{\text{INTER}}(i, j, \phi)$ operates on a conditional ensemble for subsequence $[i, j]$ with $i$ and $j$ on different strands and base paired to each other (depicted with a solid line between $i$ and $j$). The function $\textsc{Complementary}(\phi_i, \phi_j)$ checks if bases $\phi_i$ and $\phi_j$ are complementary (Watson–Crick or wobble pair) without regard to whether $i$ and $j$ are sufficiently separated along the strand to be able to pair sterically. In the mixed-material setting, the appropriate single- or mixed-material parameters must be applied based on the material composition of each loop. The recursion distinguishes three cases that are combined using $\oplus$ in the recursion equation:

- *Exterior loop:* The exterior loop is closed by one or more terminal base pairs including terminal base pair $j \cdot i$. The exterior loop treatment does not change in the mixed-material setting and is as previously described.[1]

  - *Base case:* The base case corresponds to the first recursion diagram in the first row of Figure S14 with a nick at $c$. For each nick $c \in \eta$, the contribution of subsequence $[i+1, c-1]$ is incorporated by element $Q^{\varnothing}_{i+1,c-1}$ and the contribution of subsequence $[c, j-1]$ is incorporated by element $Q^{\varnothing}_{c,j-1}$. Shading corresponds to the recursion energy $\Delta G^{\text{terminalbp}}_{j,i}(\phi)$ representing the sequence-dependent penalty for a terminal base pair in an exterior loop context (note that the indices are ordered $j$ then $i$ to reflect $5'$ to $3'$ from the perspective of the exterior loop). $\Delta G^{\text{terminalbp}}_{j,i}(\phi)$ is determined by lookup based on the sequence and material of nucleotides $j$ and $i$.

  ◦ *Edge cases:* In the base case, there is a $Q^{\varnothing}$ element on either side of the nick. In the edge cases treated by the three rightmost diagrams in the first row of Figure S14, one or both of these subsequences is absent because the nick is adjacent to $i$ (diagram 1), adjacent to $j$ (diagram 2), or adjacent to both $i$ and $j$ (diagram 3).

- *Interior loop:* the interior loop closed by the two terminal base pairs $i \cdot j$ and $d \cdot e$ (depicted by straight solid lines). We defer discussion of the calculation of the interior loop contributions using INTERIORINTER until Section S6.4, where we describe both $O(N^4)$ and $O(N^3)$ recursions.

- *Multiloop:* the multiloop closed by three or more terminal base pairs: 1) the terminal base pair $i \cdot j$ depicted by a straight solid line, 2) a $3'$-most terminal base pair starting at $d+1$ and ending in interval $[d+2, j-1]$ (depicted by a straight half-solid/half dashed line between $d+1$ and $j-1$); the contribution of subsequence $[d+1, j-1]$ is incorporated by element $Q^{ms}_{d+1,j-1}$, 3) one or more additional terminal base pairs in the interval $[i+1, d]$ (the straight dashed line denotes that $i+1$ and $d$ may or may not be paired); the contribution of subsequence $[i+1, d]$ is incorporated by element $Q^{m}_{i+1,d}$.

  ◦ *Single- and mixed-material recursions.* Multiloop contributions must be kept separate by material in $R^b_{\text{INTER}}(i,j,\phi)$ so that the appropriate $\Delta G^{\text{multi}}_{\text{init}}$, $\Delta G^{\text{multi}}_{\text{bp}}$, and $\Delta G^{\text{terminalbp}}_{j,i}(\phi)$ parameters are applied depending on the material of the multiloop. Contributions for single- and mixed-material multiloops are indicated in the recursion equation as $M_{\text{single}}$ and $M_{\text{mix}}$, respectively. To use the correct physical model for each multiloop as recursion elements are combined across multiple recursion levels, lower level recursions must be calculated separately for the "single" and "mixed" scenarios before they are combined at the $R^b_{\text{INTER}}(i,j,\phi)$ level. $Q^{ms}_{d+1,j-1}$ is split into $Q^{ms,\text{single}}_{d+1,j-1}$ and $Q^{ms,\text{mix}}_{d+1,j-1}$, while $Q^{m}_{i+1,d}$ is split into $Q^{m,\text{single}}_{i+1,d}$ and $Q^{m,\text{mix}}_{i+1,d}$. "Single" elements (depicted with a blue backbone in recursion diagrams) only track energy contributions from single-material loop regions, while "mix" elements (depicted with a red background in recursion diagrams) only track contributions from mixed-material loop regions. Because $Q^{m}_{i+1,d}$ and $Q^{ms}_{d+1,j-1}$ are each split into single-material and mixed-material recursions, there are four possible combinations when multiplying them. Any combination with a mixed element produces an overall mixed multiloop ($Q^{m,\text{mix}}_{i+1,d} \otimes Q^{ms,\text{mix}}_{d+1,j-1}, Q^{m,\text{mix}}_{i+1,d} \otimes Q^{ms,\text{single}}_{d+1,j-1}, Q^{m,\text{single}}_{i+1,d} \otimes Q^{ms,\text{mix}}_{d+1,j-1}$), which is added to $M_{\text{mix}}$ as the term $C_{1m}$. The remaining combination, $Q^{m,\text{single}}_{i+1,d} \otimes Q^{ms,\text{single}}_{d+1,j-1}$, in term $C_{1s}$ produces a single-material multiloop only if the material of both elements match each other and the material of the closing base pair $i \cdot j$, which is checked using the conditional, $\mu_j = \mu_{j-1} = \mu_{i+1} = \mu_i$. If the conditional is satisfied, the contribution is added to $M_{\text{single}}$ using term $C_{0s}$. Otherwise, if the conditional fails, the contribution is added to $M_{\text{mix}}$ using term $C_{0m}$. In the conditional, note that the material of any multiloop "single" element, such as $Q^{m,\text{single}}_{i+1,d}$ or $Q^{ms,\text{single}}_{d+1,j-1}$, can be determined by checking the material of the first or last nucleotide of the subsequence spanned by that element; the first and last nucleotides are each guaranteed to appear in the multiloop as either an unpaired nucleotide or as part of a terminal base pair defining the multiloop. In this instance, the material of $Q^{m,\text{single}}_{i+1,\overline{d}}$ is checked at $i+1$, while the material of $Q^{ms,\text{single}}_{\overline{d}+1,j-1}$ is checked at $j-1$, since these indices are constant over vector $\overline{d}$ and apply to all $d$ and $d+1$ in $\overline{d}$. To exclude exterior loop states that are not treated by this multiloop recursion, the function VALID returns the set of valid vectorization ranges for which nucleotides $d$ and $d+1$ are on the same strand (i.e., such that $d$ and $d+1$ do not take on values that would place a nick between them).

  ◦ *Single- and mixed-material recursion energies.* In recursion diagrams, shading corresponds to three recursion energies: 1) the penalty for formation of a multiloop $\Delta G^{\text{multi}}_{\text{init}}$, 2) the sequence-independent penalty for a terminal base pair in a multiloop $\Delta G^{\text{multi}}_{\text{bp}}$ (corresponding to the sole base pair $i \cdot j$ that is fully defined in this recursion), 3) the sequence-dependent penalty for a terminal base pair in a multiloop context, $\Delta G^{\text{terminalbp}}_{j,i}(\phi)$ (note that the indices are ordered $j$ then $i$ to reflect $5'$ to $3'$ from the perspective of the multiloop). For $M_{\text{single}}$ (single-material multiloops checked using the conditional $\mu_j = \mu_{j-1} = \mu_{i+1} = \mu_i$), the single-material multiloop initiation parameter, $\Delta G^{\text{multi}}_{\text{init},\mu_i}$, is used and the single-material terminal base pair parameter, $\Delta G^{\text{multi}}_{\text{bp},\mu_i}$, is used for terminal base pair $i \cdot j$. For $M_{\text{mix}}$, the mixed-material multiloop initiation parameter, $\Delta G^{\text{multi}}_{\text{init,mix}}$, is used and the terminal base pair parameter, $\Delta G^{\text{multi}}_{\text{bp},\sigma}$, depends on the material of closing base pair $i \cdot j$ and is checked with the $\sigma$ conditional ($\Delta G^{\text{multi}}_{\text{bp},\mu_i}$ is used if $\mu_j = \mu_i$, and $\Delta G^{\text{multi}}_{\text{bp,mix}}$ is used otherwise). The sequence-dependent terminal base pair parameter, $\Delta G^{\text{terminalbp}}_{j,i}(\phi)$, is determined by lookup based on the sequence and material of nucleotides $j$ and $i$. $M_{\text{single}}$ and $M_{\text{mix}}$ are both added to $R^b_{\text{INTER}}(i,j,\phi)$ to incorporate all single-material and mixed-material multiloops that include terminal base pair $i \cdot j$.

Note that unlike the $R^b_{\text{INTRA}}$ recursion of Figure S8, for $R^b_{\text{INTER}}$ there is no hairpin loop case as $i$ and $j$ are on different strands.

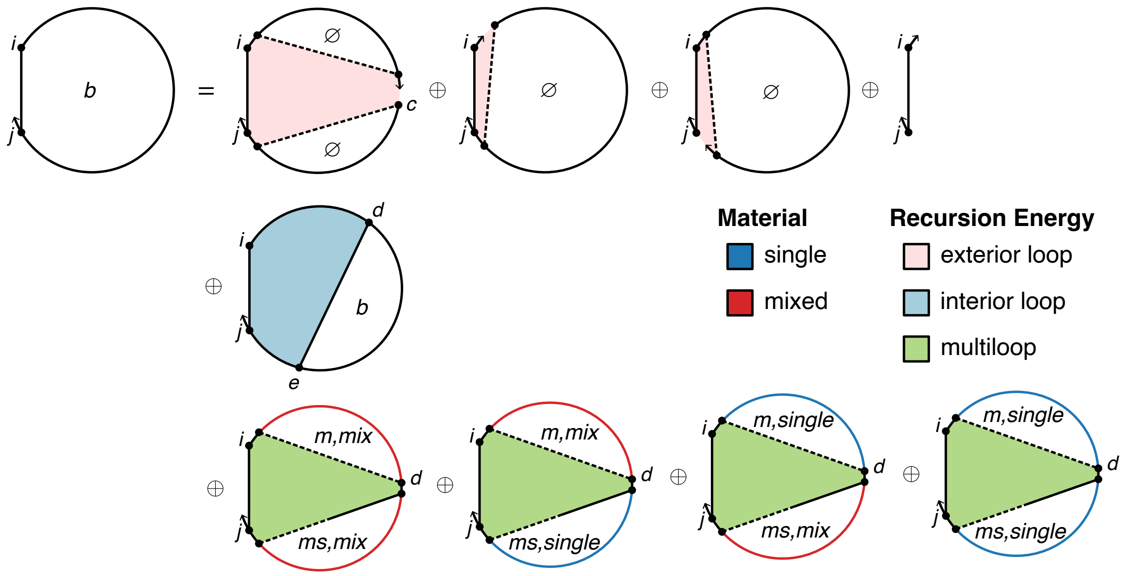

$$R^{b}_{\textsc{Inter}}(i,j,\phi) \equiv \begin{cases} C_{\text{all}}, & \textsc{Complementary}(\phi_i,\phi_j) \\ \mathbb{0}, & \text{otherwise} \end{cases}$$

$$\text{where } C_{\text{all}} \equiv C_{\text{exterior}} \oplus \textsc{InteriorInter}(i,j,\phi) \oplus M_{\text{single}} \oplus M_{\text{mix}}$$

$$C_{\text{exterior}} \equiv \begin{cases} \bigoplus_{c\in\eta} Q^{\varnothing}_{i+1,c-1} \otimes Q^{\varnothing}_{c,j-1} \otimes W(\Delta G^{\text{terminalbp}}_{j,i}(\phi)), & i+1 \neq m \text{ and } j \neq n \\ Q^{\varnothing}_{m,j-1} \otimes W(\Delta G^{\text{terminalbp}}_{j,i}(\phi)), & i+1 = m \text{ and } j \neq n \\ Q^{\varnothing}_{i+1,n-1} \otimes W(\Delta G^{\text{terminalbp}}_{j,i}(\phi)), & i+1 \neq m \text{ and } j = n \\ W(\Delta G^{\text{terminalbp}}_{j,i}(\phi)), & i+1 = j = m = n \end{cases}$$

$$M_{\text{single}} \equiv \bigoplus_{\overline{d}\in\textsc{Valid}(i+1,j-1,\eta)} C_{0s} \otimes W(\Delta G^{\text{multi}}_{\text{init},\mu_i} + \Delta G^{\text{multi}}_{\text{bp},\mu_i} + \Delta G^{\text{terminalbp}}_{j,i}(\phi)),$$

$$C_{0s} \equiv \begin{cases} C_{1s}, & \mu_j = \mu_{j-1} = \mu_{i+1} = \mu_i \\ \mathbb{0}, & \text{otherwise} \end{cases}$$

$$C_{1s} \equiv \textsc{dot}\left(Q^{m,\text{single}}_{i+1,\overline{d}}, Q^{ms,\text{single}}_{\overline{d}+1,j-1}\right)$$

$$M_{\text{mix}} \equiv \bigoplus_{\overline{d}\in\textsc{Valid}(i+1,j-1,\eta)} (C_{0m} \oplus C_{1m}) \otimes W(\Delta G^{\text{multi}}_{\text{init,mix}} + \Delta G^{\text{multi}}_{\text{bp},\sigma} + \Delta G^{\text{terminalbp}}_{j,i}(\phi))$$

$$C_{0m} \equiv \begin{cases} \mathbb{0}, & \mu_j = \mu_{j-1} = \mu_{i+1} = \mu_i \\ C_{1s}, & \text{otherwise} \end{cases}$$

$$C_{1m} \equiv \textsc{dot}\left(Q^{m,\text{mix}}_{i+1,\overline{d}}, Q^{ms,\text{mix}}_{\overline{d}+1,j-1}\right) \oplus \textsc{dot}\left(Q^{m,\text{mix}}_{i+1,\overline{d}}, Q^{ms,\text{single}}_{\overline{d}+1,j-1}\right) \oplus \textsc{dot}\left(Q^{m,\text{single}}_{i+1,\overline{d}}, Q^{ms,\text{mix}}_{\overline{d}+1,j-1}\right)$$

$$\text{with } \sigma \equiv \begin{cases} \mu_i, & \mu_j = \mu_i \\ \text{mix}, & \text{otherwise} \end{cases}$$

$$m = \textsc{First}(\eta)$$

$$n = \textsc{Last}(\eta)$$

Figure S14: $R^{b}_{\textsc{Inter}}$ recursion without coaxial and dangle stacking. Top: recursion diagram. Bottom: recursion equation.

$\boldsymbol{R^{ms}_{\text{INTER}}}$ **recursion without coaxial and dangle stacking.** The $R^b_{\text{INTER}}(i,j,\phi)$ recursion references $Q^{ms}_{d+1,j-1}$ elements that are computed using either the $R^{ms}_{\text{INTRA}}$ recursion shown of Figure S9 (if $d+1$ and $j-1$ are on the same strand) or the $R^{ms}_{\text{INTER}}$ recursion of Figure S15 (if $d+1$ and $j-1$ are on different strands). $R^{ms}_{\text{INTER}}(i,j,\phi)$ operates on a conditional ensemble for subsequence $[i,j]$ in a multiloop context containing one terminal base pair starting at $i$ and ending in the interval $[i+1,j]$ (depicted as a half-solid/half-dashed line between $i$ and $j$). A terminal base pair is either a single material (denoted in the recursion diagram by a blue base pair, if the material of the two nucleotides matches) or mixed material (denoted by a red base pair, if the material of the two nucleotides does not match). Like other multiloop recursions in the mixed algorithm, $R^{ms}_{\text{INTER}}(i,j,\phi)$ is split into $R^{ms,\text{single}}_{\text{INTER}}(i,j,\phi)$ and $R^{ms,\text{mix}}_{\text{INTER}}(i,j,\phi)$ (the single and mixed recursions are indicated in the diagram with blue and red backbones, respectively):

- $R^{ms,\text{single}}_{\text{INTER}}(i,j,\phi)$: This recursion treats the $R^{ms}_{\text{INTER}}(i,j,\phi)$ conditional ensemble over $[i,j]$ where terminal base pair $i \cdot d$ and any unpaired nucleotides in the interval $[d+1,j]$ are of a single material. For the edge case where $d=j$, then the whole region $[i,j]$ enclosed by base pair $i \cdot j$ is incorporated using a $Q^b_{i,j}$ element in term $C_2$. This represents a single-material contribution only if $\mu_j=\mu_i$. If the conditional is satisfied, the contribution is added to $R^{ms,\text{single}}_{\text{INTRA}}(i,j,\phi)$ using term $C_{0s}$. Otherwise, if the conditional fails, the contribution is added to $R^{ms,\text{mix}}_{\text{INTRA}}(i,j,\phi)$ using term $C_{0m}$. For the base case, the contribution of terminal base pair $i \cdot d$ is incorporated using $Q^b_{i,d}$ if $\mu_d=\mu_i$ (indicating a single-material terminal base pair), while the unpaired region $[d+1,j]$ is incorporated using $Q^{ma,\text{single}}_{d+1,j}$. The resulting contribution $Q^b_{i,d} \otimes Q^{ma,\text{single}}_{d+1,j}$ of term $C_{1s}$ produces a single-material multiloop only if the material of the terminal base pair $i \cdot d$ matches that of $Q^{ma,\text{single}}_{d+1,j}$, which is checked with conditional $\mu_j=\mu_i$. If the conditional is satisfied, the contribution is added to $R^{ms,\text{single}}_{\text{INTER}}(i,j,\phi)$ using term $C_{0s}$. Otherwise, if the conditional fails, the contribution is added to $R^{ms,\text{mix}}_{\text{INTER}}(i,j,\phi)$ using term $C_{0m}$.
- $R^{ms,\text{mix}}_{\text{INTER}}(i,j,\phi)$: This recursion treats the $R^{ms}_{\text{INTER}}(i,j,\phi)$ conditional ensemble over $[i,j]$ in a multiloop context where terminal base pair $i \cdot d$ and any unpaired nucleotides overall contain mixed materials. For the edge case where $d=j$ and the whole region $[i,j]$ enclosed by base pair $i \cdot j$ is incorporated using a $Q^b_{i,j}$ element in term $C_2$. As noted above, this represents a mixed-material contribution only if $\mu_j \neq \mu_i$ and is incorporated in $R^{ms,\text{mix}}_{\text{INTER}}(i,j,\phi)$ using term $C_{0m}$. For the base case, the contribution for the subsequence $[i,d]$ enclosed by terminal base pair $i \cdot d$ is incorporated using a $Q^b_{i,d}$ element, which represents a single-material contribution if $\mu_d=\mu_i$ and a mixed-material contribution otherwise. The contribution for the unpaired region $[d+1,j]$ is incorporated using either $Q^{ma,\text{single}}_{d+1,j}$ or $Q^{ma,\text{mix}}_{d+1,j}$. There are four possible single/mix combinations for $Q^b_{i,d} \otimes Q^{ma}_{d+1,j}$. The three combinations that contain at least one mixed element must represent mixed contributions and are included in $R^{ms,\text{mix}}_{\text{INTER}}(i,j,\phi)$ using term $C_{1m}$. As noted above, the remaining combination calculated in term $C_{1s}$ represents a mixed contribution if $\mu_j \neq \mu_i$, and is incorporated in term $R^{ms,\text{mix}}_{\text{INTER}}(i,j,\phi)$ using term $C_{0m}$.

Shading corresponds to two recursion energies: 1) the sequence-independent penalty for a terminal base pair in a multiloop (for base pair $i \cdot d$, $\Delta G^{\text{multi}}_{\text{bp},\mu_i}$ if $\mu_d=\mu_i$, and $\Delta G^{\text{multi}}_{\text{bp,mix}}$ otherwise; for base pair $i \cdot j$, $\Delta G^{\text{multi}}_{\text{bp},\sigma}$, where the parameter depends on the material as determined by the $\sigma$ conditional), 2) the sequence-dependent penalty for a terminal base pair in a multiloop context (for base pair $i \cdot d$, $\Delta G^{\text{terminalbp}}_{i,d}(\phi)$, dependent on the sequence and material of base pair $i \cdot d$; for base pair $i \cdot j$, $\Delta G^{\text{terminalbp}}_{i,j}(\phi)$, dependent on the sequence and material of base pair $i \cdot j$). Nucleotide $d$ must always be on the last strand $(d \geq \textsc{Last}(\eta))$ to ensure that there are no nicks in the subsequence $[d,j]$, which would lead to either a disconnected structure (which is not permitted in the complex ensemble) or an exterior loop state (which is not handled by this multiloop recursion). Also, $d \leq j-1$ in order to provide a minimum of one nucleotide for the $Q^{ma}_{d+1,j}$ recursion, so we must have $j \geq \textsc{Last}(\eta)+1$. Note that $R^{ms}$ serves as an efficiency wrapper for $R^b$ in the multiloop context in a completely analogous manner to $R^s$ serving as an efficiency wrapper for $R^b$ in an exterior loop context, with $R^b$ representing the $3'$-most terminal base pair in either context.

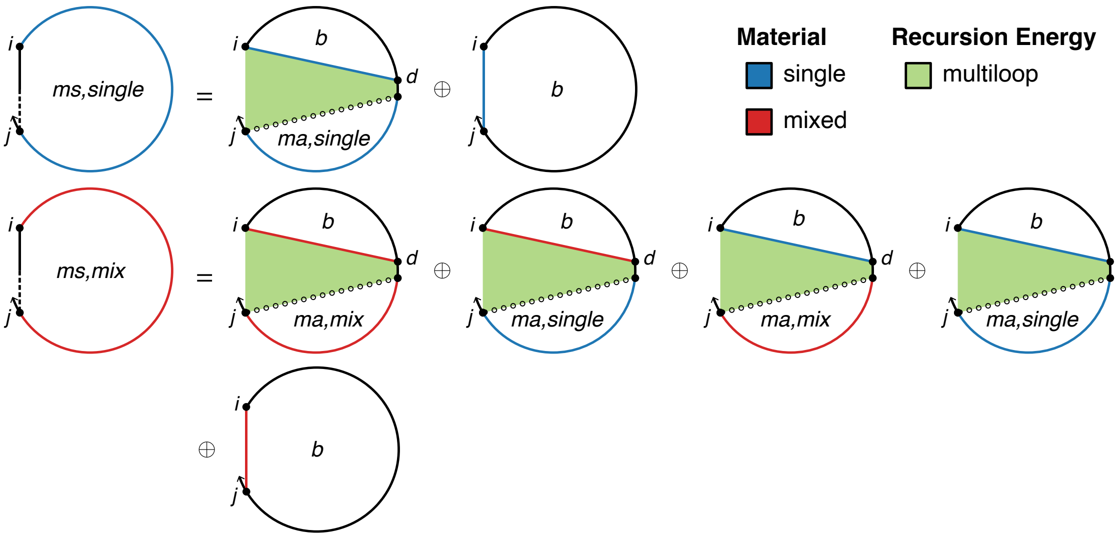

$$R^{ms,\text{single}}_{\text{INTER}}(i,j,\phi) \equiv C_{0s}$$

$$C_{0s} \equiv \begin{cases} C_{1s} \oplus C_2, & \mu_j = \mu_i \\ \mathbb{0}, & \text{otherwise} \end{cases}$$

$$C_{1s} \equiv \begin{cases} \text{DOT}\left(Q^b_{i,\overline{d}}, Q^{ma,\text{single}}_{\overline{d}+1,j}, W(\Delta G^{\text{multi}}_{\text{bp},\mu_i} + \Delta G^{\text{terminalbp}}_{i,\overline{d}}(\phi))\right), & j \geq \textsc{Last}(\eta)+1 \text{ and } \mu_d = \mu_i \\ \mathbb{0}, & \text{otherwise} \end{cases}$$

$$C_2 \equiv Q^b_{i,j} \otimes W(\Delta G^{\text{multi}}_{\text{bp},\sigma} + \Delta G^{\text{terminalbp}}_{i,j}(\phi)),$$

$$R^{ms,\text{mix}}_{\text{INTER}}(i,j,\phi) \equiv C_{0m} \oplus C_{1m}$$

$$C_{0m} \equiv \begin{cases} \mathbb{0}, & \mu_j = \mu_i \\ C_{1s} \oplus C_2, & \text{otherwise} \end{cases}$$

$$\oplus \begin{cases} C_1 + C_2 + C_3, & j \geq \textsc{Last}(\eta)+1 \\ \mathbb{0}, & \text{otherwise} \end{cases}$$

$$\text{where } C_{1m} \equiv \begin{cases} \text{DOT}\left(Q^b_{i,\overline{d}}, (Q^{ma,\text{mix}}_{\overline{d}+1,j} \oplus Q^{ma,\text{single}}_{\overline{d}+1,j}), W(\Delta G^{\text{multi}}_{\text{bp},\text{mix}} + \Delta G^{\text{terminalbp}}_{i,\overline{d}}(\phi))\right), & j \geq \textsc{Last}(\eta)+1 \text{ and } \mu_d \neq \mu_i \\ \mathbb{0}, & \text{otherwise} \end{cases}$$

$$\oplus \begin{cases} \text{DOT}\left(Q^b_{i,\overline{d}}, Q^{ma,\text{mix}}_{\overline{d}+1,j}, W(\Delta G^{\text{multi}}_{\text{bp},\mu_i} + \Delta G^{\text{terminalbp}}_{i,\overline{d}}(\phi))\right), & j \geq \textsc{Last}(\eta)+1 \text{ and } \mu_d = \mu_i \\ \mathbb{0}, & \text{otherwise} \end{cases}$$

$$\text{where } \sigma \equiv \begin{cases} \mu_i, & \mu_j = \mu_i \\ \text{mix}, & \text{otherwise} \end{cases}$$

$$\overline{d} \equiv [\textsc{Last}(\eta) : j-1]$$

Figure S15: $R^{ms}_{\text{INTER}}$ recursion without coaxial and dangle stacking. Top: recursion diagram. Bottom: recursion equation.

$\boldsymbol{R^{ma}_{\text{INTER}}}$ **recursion without coaxial and dangle stacking.** The $R^{ms}_{\text{INTER}}(i,j,\phi)$ recursion references $Q^{ma}_{d+1,j}$ elements that are always computed using the $R^{ma}_{\text{INTRA}}$ recursion of Figure S10 (if $d+1$ and $j$ are on the same strand) and never computed using the $R^{ma}_{\text{INTER}}$ recursion of Figure S16 (if $d+1$ and $j$ are on different strands). $R^{ma}_{\text{INTER}}(i,j,\phi)$ treats the conditional ensemble of subsequence $[i,j]$ in a multiloop context containing no terminal base pairs, where $i$ and $j$ are on different strands. Note that within the $R^{ms}_{\text{INTER}}(i,j,\phi)$ recursion, $d$ must be on the last strand ($d \geq \text{LAST}(\eta)$) to ensure there are no nicks in the subsequence $[d,j]$ (which would otherwise lead to a disconnected structure or an exterior loop state), and $d \leq j-1$ in order to provide a minimum of one nucleotide for the $Q^{ma}_{d+1,j}$ recursion. Hence, $d$ and $j$ are always on the same strand and there are no calls to $R^{ma}_{\text{INTER}}$. Equivalently, the contributions for both single- and mixed-material $R^{ma}_{\text{INTER}}(i,j,\phi)$ recursions are $\mathbb{0}$ (Figure S16).

$$R^{ma,\text{single}}_{\text{INTER}}(i,j,\phi) \equiv \mathbb{0}$$
$$R^{ma,\text{mix}}_{\text{INTER}}(i,j,\phi) \equiv \mathbb{0}$$

Figure S16: $R^{ma}_{\text{INTER}}$ recursion without coaxial and dangle stacking. No recursion diagram is provided because the recursion equations contain no contributions.

$\boldsymbol{R^{m}_{\text{INTER}}}$ **recursion without coaxial and dangle stacking.** The $R^{b}_{\text{INTER}}(i,j,\phi)$ recursion references $Q^{m}_{i+1,d}$ elements that are computed using either the $R^{m}_{\text{INTRA}}$ recursion of Figure S11 (if $i+1$ and $d$ are on the same strand), or the $R^{m}_{\text{INTER}}$ recursion of Figure S17 (if $i+1$ and $d$ are on different strands). $R^{m}_{\text{INTER}}(i,j,\phi)$ operates on a conditional ensemble for subsequence $[i,j]$ in a multiloop context where $i$ and $j$ may or may not be paired (depicted with a dashed line between $i$ and $j$ in the recursion diagram) and where there is at least one terminal base pair. Like other multiloop recursions in the mixed algorithm, $R^{m}_{\text{INTER}}(i,j,\phi)$ is split into $R^{m,\text{single}}_{\text{INTER}}(i,j,\phi)$ and $R^{m,\text{mix}}_{\text{INTER}}(i,j,\phi)$ (the single and mixed recursions are indicated in the diagram with blue and red backbones, respectively):

- $R^{m,\text{single}}_{\text{INTER}}(i,j,\phi)$: This recursion operates over the $R^{m}_{\text{INTER}}(i,j,\phi)$ conditional ensemble on $[i,j]$ in a multiloop context where any terminal base pair or unpaired nucleotide is of a single material. This recursion distinguishes two cases that are combined using $\oplus$ in the recursion equation:
  - *One terminal base pair*: the case where there is exactly one terminal base pair in subsequence $[i,j]$ in a multiloop context. This terminal base pair starts at $d$ and ends in the interval $[d+1,j]$ (depicted by a straight half-solid/half dashed line between $d$ and $j$). Subsequence $[i,d-1]$ is an unpaired region (depicted by a hollow dotted line between $i$ and $d-1$). For the edge case where $d=i$, then all of $[i,j]$ is incorporated by a $Q^{ms,\text{single}}_{i,j}$ element. No further material checks are needed since the $R^{ms,\text{single}}_{\text{INTER}}(i,j,\phi)$ recursion only includes single-material contributions. For the base case where $d$ is between $i+1$ and $\text{FIRST}(\eta)-1$ with $i \leq \text{FIRST}(\eta)-2$, then the unpaired region $[i,d-1]$ is incorporated using a $Q^{ma,\text{single}}_{i,d-1}$ element while the region $[d,j]$ is incorporated using $Q^{ms,\text{single}}_{d,j}$. The resulting contribution $Q^{ma,\text{single}}_{i,d-1} \otimes Q^{ms,\text{single}}_{d,j}$ of term $C_{s1}$ produces a single-material multiloop only if the material matches for both elements, which is checked with conditional $\mu_j = \mu_i$. If the conditional is satisfied, the contribution is added to $R^{m,\text{single}}_{\text{INTER}}(i,j,\phi)$ using term $C_{0s}$. Otherwise, if the conditional fails, the contribution is added to $R^{m,\text{mix}}_{\text{INTER}}(i,j,\phi)$ using term $C_{0m}$. In the conditional, indices $i$ and $j$ are used to check the material since they are constant over vector $\overline{d}$ and apply to all $d-1$ and $d$ in $\overline{d}$.
  - *More than one terminal base pair*: the case where there are two or more terminal base pairs in subsequence $[i,j]$ in a multiloop context. The $3'$-most terminal base pair starts at $e+1$ and ends in the interval $[e+2,j]$ (depicted by a straight half-solid/half dashed line between $e+1$ and $j$); the contribution of subsequence $[e+1,j]$ is incorporated by element $Q^{ms,\text{single}}_{e+1,j}$. There are one or more additional terminal base pairs in the interval $[i,e]$ (the straight dashed line denotes that $i$ and $e$ may or may not be paired); the contribution of subsequence $[i,e]$ is incorporated by element $Q^{m,\text{single}}_{i,e}$. The resulting contribution $Q^{m,\text{single}}_{i,e} \otimes Q^{ms,\text{single}}_{e+1,j}$ of term $C_{2s}$ produces a single-material multiloop only if the material matches for both elements, which is checked with conditional $\mu_j = \mu_i$. If the conditional is satisfied, the contribution is added to $R^{m,\text{single}}_{\text{INTER}}(i,j,\phi)$ using term $C_{0s}$. Otherwise, if the conditional fails, the contribution is added to $R^{m,\text{mix}}_{\text{INTER}}(i,j,\phi)$ using term $C_{0m}$. In the conditional, indices $i$ and $j$ are used to check the material since they are constant over vector $\overline{e}$ and apply to all $e$ and $e+1$ in $\overline{e}$.
- $R^{m,\text{mix}}_{\text{INTER}}(i,j,\phi)$: This recursion operates over the $R^{m}_{\text{INTER}}(i,j,\phi)$ conditional ensemble on $[i,j]$ in a multiloop context where any terminal base pair and unpaired nucleotides overall contain a mixture of materials. This recursion distinguishes two cases that are combined using $\oplus$ in the recursion equation:

- *One terminal base pair*: the case where there is exactly one terminal base pair in subsequence $[i, j]$ in a multiloop context. This terminal base pair starts at $d$ and ends in the interval $[d+1, j]$ (depicted by a straight half-solid/half dashed line between $d$ and $j$). Subsequence $[i, d-1]$ is an unpaired region (depicted by a hollow dotted line between $i$ and $d-1$). For the edge case where $d = i$, then the whole region $[i, j]$ is incorporated by a $Q^{ms,\text{mix}}_{i,j}$ element. No further material checks are needed since the $R^{ms,\text{mix}}_{\text{INTER}}(i, j, \phi)$ recursion only includes mixed material contributions. For the base case where $d$ is between $i+1$ and $\text{FIRST}(\eta)-1$, then the unpaired region $[i, d-1]$ is incorporated using a $Q^{ma}_{i,d-1}$ element while the region $[d, j]$ is incorporated using a $Q^{ms}_{d,j}$ element. Because $Q^{ma}_{i,d-1}$ and $Q^{ms}_{d,j}$ are each spit into single- and mixed-material recursions, there are 4 possible combinations when multiplying them. The three combinations containing at least one mixed element ($Q^{ma,\text{single}}_{i,d-1} \otimes Q^{ms,\text{mix}}_{d,j}$, $Q^{ma,\text{mix}}_{i,d-1} \otimes Q^{ms,\text{single}}_{d,j}$, $Q^{ma,\text{mix}}_{i,d-1} \otimes Q^{ms,\text{mix}}_{d,j}$) must represent overall mixed contributions, and are included in $R^{m,\text{mix}}_{\text{INTER}}(i, j, \phi)$ using term $C_{1m}$. As noted above, the remaining combination, $Q^{ma,\text{single}}_{i,d-1} \otimes Q^{ms,\text{single}}_{d,j}$ calculated in term $C_{1s}$, only represents a mixed contribution if $\mu_j \neq \mu_i$, and is incorporated in $R^{m,\text{mix}}_{\text{INTRA}}(i, j, \phi)$ using term $C_{0m}$.
- *More than one terminal base pair*: the case where there are two or more terminal base pairs in subsequence $[i, j]$ in a multiloop context. The $3'$-most terminal base pair starts at $e+1$ and ends in the interval $[e+2, j]$ (depicted by a straight half-solid/half dashed line between $e+1$ and $j$); the contribution of subsequence $[e+1, j]$ is incorporated by a $Q^{ms}_{e+1,j}$ element. There are one or more additional terminal base pairs in the interval $[i, e]$ (the straight dashed line denotes that $i$ and $e$ may or may not be paired); the contribution of subsequence $[i, e]$ is incorporated by a $Q^{m}_{i,e}$ element. Because $Q^{m}_{i,e}$ and $Q^{ms}_{e+1,j}$ are each spit into single- and mixed-material recursions, there are 4 possible combinations when multiplying them. The three combinations containing at least one mixed element ($Q^{m,\text{single}}_{i,e} \otimes Q^{ms,\text{mix}}_{e+1,j}$, $Q^{m,\text{mix}}_{i,e} \otimes Q^{ms,\text{single}}_{e+1,j}$, $Q^{m,\text{mix}}_{i,e} \otimes Q^{ms,\text{mix}}_{e+1,j}$) must represent mixed contributions, and are included in $R^{m,\text{mix}}_{\text{INTER}}(i, j, \phi)$ using term $C_{2m}$. As noted above, the remaining combination, $Q^{m,\text{single}}_{i,e} \otimes Q^{ms,\text{single}}_{e+1,j}$ calculated in term $C_{2s}$, only represents a mixed contribution if $\mu_j \neq \mu_i$, and is incorporated in $R^{m,\text{mix}}_{\text{INTER}}(i, j, \phi)$ using term $C_{0m}$.

Shading in the recursion diagram does not represent any recursion energies because all multiloop contributions are handled by other recursions: 1) there are no terminal base pairs in a multiloop context explicitly defined in this case, 2) there are no unpaired bases in a multiloop context explicitly defined in this case. Nucleotide $d$ must always be on the first strand ($d \leq \text{FIRST}(\eta) - 1$) to ensure that there are no nicks in the subsequence $[i, d]$, which would lead to either a disconnected structure (which is not permitted in the complex ensemble) or an exterior loop state (which is not handled by this multiloop recursion). Also, $d \geq i+1$ in order to provide a minimum of one nucleotide for the $Q^{ma}_{i,d-1}$ recursion, so we must have $i \leq \text{FIRST}(\eta) - 2$ for terms $C_{1s}$ and $C_{1m}$. To exclude exterior loop states that are not treated by this multiloop recursion, the function VALID returns the set of valid vectorization ranges for which nucleotides $e$ and $e+1$ are on the same strand for terms $C_{2s}$ and $C_{2m}$ (i.e., such that $e$ and $e+1$ do not take on values that would place a nick between them).

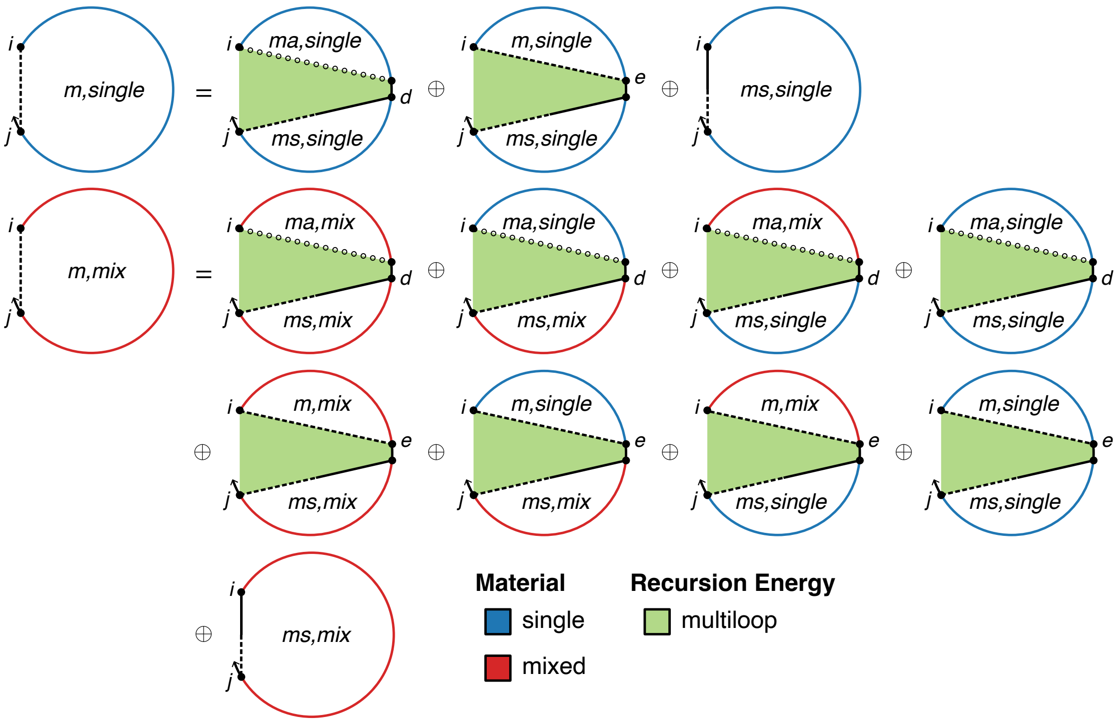

$$R^{m,\text{single}}_{\text{INTER}}(i,j,\phi) \equiv C_{0s} \oplus C_{3s}$$

$$C_{0s} \equiv \begin{cases} C_{1s} \oplus C_{2s}, & \mu_j = \mu_i \\ \mathbb{0}, & \text{otherwise} \end{cases}$$

$$C_{1s} \equiv \begin{cases} \text{DOT}\left(Q^{ma,\text{single}}_{i,\overline{d}-1}, Q^{ms,\text{single}}_{\overline{d},j}\right), & i \leq \text{FIRST}(\eta) - 2 \\ \mathbb{0}, & \text{otherwise} \end{cases}$$

$$C_{2s} \equiv \bigoplus_{\overline{e} \in \text{VALID}(i,j,\eta)} \text{DOT}\left(Q^{m,\text{single}}_{i,\overline{e}}, Q^{ms,\text{single}}_{\overline{e}+1,j}\right)$$

$$C_{3s} \equiv Q^{ms,\text{single}}_{i,j}$$

$$R^{m,\text{mix}}_{\text{INTER}}(i,j,\phi) \equiv C_{0m} \oplus C_{1m} \oplus C_{2m} \oplus C_{3m}$$

$$C_{0m} \equiv \begin{cases} \mathbb{0}, & \mu_j = \mu_i \\ C_{1s} \oplus C_{2s}, & \text{otherwise} \end{cases}$$

$$C_{1m} \equiv \begin{cases} \text{DOT}\left(Q^{ma,\text{mix}}_{i,\overline{d}-1}, Q^{ms,\text{single}}_{\overline{d},j}\right) \oplus \text{DOT}\left(Q^{ma,\text{single}}_{i,\overline{d}-1}, Q^{ms,\text{mix}}_{\overline{d},j}\right) \oplus \text{DOT}\left(Q^{ma,\text{mix}}_{i,\overline{d}-1}, Q^{ms,\text{mix}}_{\overline{d},j}\right), & i \leq \text{FIRST}(\eta) - 2 \\ \mathbb{0}, & \text{otherwise} \end{cases}$$

$$C_{2m} \equiv \bigoplus_{\overline{e} \in \text{VALID}(i,j,\eta)} \text{DOT}\left(Q^{m,\text{mix}}_{i,\overline{e}}, Q^{ms,\text{single}}_{\overline{e}+1,j}\right) \oplus \text{DOT}\left(Q^{m,\text{single}}_{i,\overline{e}}, Q^{ms,\text{mix}}_{\overline{e}+1,j}\right) \oplus \text{DOT}\left(Q^{m,\text{mix}}_{i,\overline{e}}, Q^{ms,\text{mix}}_{\overline{e}+1,j}\right)$$

$$C_{3m} \equiv Q^{ms,\text{mix}}_{i,j}$$

$$\text{with } \overline{d} \equiv [i+1 : \text{FIRST}(\eta) - 1] \tag{S138}$$

Figure S17: $R^m_{\text{INTER}}$ recursion without coaxial and dangle stacking. Top: recursion diagram. Bottom: recursion equation.

#### S6.4 Recursions for interior loop contributions

Interior loop contributions to the recursions $R^b_{\text{INTRA}}(i,j,\phi)$ and $R^b_{\text{INTER}}(i,j,\phi)$ run naively in $O(N^4)$ time, as is evident from the four indices $i,d,e,j$ in the interior loop recursion diagrams in Figures S8 and S14.

##### S6.4.1 Intrastrand interior loop dynamic programming recursions

***O*($N^4$) intrastrand interior loop recursion.** The intrastrand $O(N^4)$ interior loop contribution:

$$O(N^4)\ \textsc{InteriorIntra}(i,j,\phi) \equiv \begin{cases} \bigoplus_{d=i+1}^{j-5} \ \bigoplus_{e=d+4}^{j-1} \{Q^b_{d,e} \otimes W(\Delta G^{\text{interior}}_{i,d,e,j}(\phi,\sigma))\}, & j-i \geq 6 \\ \mathbb{0}, & \text{otherwise} \end{cases}$$
$$\text{where } \sigma \equiv \begin{cases} \mu_i, & \textsc{IsSingle}(i,d,\mu_i) \text{ and } \textsc{IsSingle}(e,j,\mu_i) \\ \text{mix}, & \text{otherwise} \end{cases} \tag{S139}$$

considers interior loops through a nested iteration, first over $d$ in a $5'$ to $3'$ direction and for each $d$ over $e$ in a $5'$ to $3'$ direction. The index limits in the recursion equation reflect the fact that steric effects prevent an interior loop with $j-i<6$ due to the steric requirement that there be at least 3 intervening bases between $d$ and $e$. The region $[d,e]$ enclosed by base pair $d\cdot e$ is represented by a $Q^b_{d,e}$ element and the function $\Delta G^{\text{interior}}_{i,d,e,j}(\phi,\sigma)$ accounts for the free energy of the interior loop with bounding base pairs $i\cdot j$ and $d\cdot e$, substituting in the correct functional form for any of the various interior loop types (stacked pair, bulge, etc; see Section S4.6). $\Delta G^{\text{interior}}_{i,d,e,j}(\phi,\sigma)$ depends on the material of the interior loop as checked with the $\sigma$ conditional ($\Delta G^{\text{interior}}_{i,d,e,j}(\phi,\mu_i)$ is used if all nucleotides are $\mu_i$, and $\Delta G^{\text{interior}}_{i,d,e,j}(\phi,\text{mix})$ is used otherwise).

***O*($N^3$) intrastrand interior loop recursion.** To reduce the complexity of computing interior loop contributions from $O(N^4)$ to $O(N^3)$, we must exploit the functional form of the free energy model for large interior loops (Section S4.6).[38] In References 39 and 40, this optimization was referred to as the "fastiloops" or "fast interior loops" function, and we take a similar approach here following Reference 1. The following optimizations assume the use of a forward operation order (not a backtracking operation order). Interior loops, defined by two bounding base pairs $i\cdot j$ and $d\cdot e$, can be classified by the distances $L_1 = d-i-1$ and $L_2 = j-e-1$; $L_1$ and $L_2$ are the numbers of unpaired nucleotides on each side of the interior loop. In cases where $L_1 < 4$ or $L_2 < 4$, the energy functions generally depend on terms that are nonlinear with respect to $L_1$ and $L_2$. Examples include the special-case energy functions for stack loops and bulge loops, as well as length-dependent asymmetry and size penalties for other interior loops. We term these interior loops *inextensible* because the free energy for a larger loop cannot in general be calculated using the value from a smaller loop. For a given subsequence $[i,j]$, there are only $O(N)$ inextensible interior loops (because of the constant upper bound on $L_1$ or $L_2$) so they do not contribute to the $O(N^4)$ complexity. The remaining interior loops in which $L_1 \geq 4$ and $L_2 \geq 4$ are referred to as *extensible* interior loops because the free energy of a larger loop can be calculated by extending the calculation from a smaller interior loop. For a given subsequence $[i,j]$ there are $O(N^2)$ extensible interior loops, so these are the cases we must deal with efficiently to reduce the time complexity from $O(N^4)$ to $O(N^3)$. For extensible interior loops, (S84) gives:

$$\Delta G^{\text{interior}}_{i,d,e,j}(\phi,\mu) = \Delta G^{\text{interiorsize}}_{L_1+L_2,\mu} + \Delta G^{\text{interiorasymm}}_{|L_1-L_2|,\mu} + \Delta G^{\text{interiormm}}_{j-1,j,i,i+1}(\phi) + \Delta G^{\text{interiormm}}_{d-1,d,e,e+1}(\phi). \tag{S140}$$

where $\mu$ is the material of the loop. Here, the quantity $\Delta G^{\text{interiorsize}}_{L_1+L_2,\mu}$ is a sequence-independent free energy contribution due to the size of the interior loop, $s \equiv L_1 + L_2$ (the sum of the two side lengths), for material $\mu$. The quantity $\Delta G^{\text{interiorasymm}}_{|L_1-L_2|,\mu}$ is a sequence-independent free energy contribution due to the *asymmetry* of the loop, $|L_1 - L_2|$ (the difference of the two side lengths), for material $\mu$. Finally, the two terms $\Delta G^{\text{interiormm}}_{j-1,j,i,i+1}(\phi)$ and $\Delta G^{\text{interiormm}}_{d-1,d,e,e+1}(\phi)$ are sequence-dependent free energy contributions due to mismatch stacking on the base pairs $i\cdot j$ and $d\cdot e$, respectively. Two key insights from (S140) allow us to use this functional form to reduce complexity.[38] First, for every base pair $i\cdot j$, the mismatch term for that base pair is independent of the other quantities and can be factored out. Second, for a given base pair $d\cdot e$, an extensible loop bounded by $i\cdot j$ can be converted to an extensible loop bounded by $i-1\cdot j+1$ by updating $\Delta G^{\text{interiorsize}}_{s,\mu}$ to $\Delta G^{\text{interiorsize}}_{s+2,\mu}$ (if the material $\mu$ stays the same after extension) and replacing $\Delta G^{\text{interiormm}}_{j-1,j,i,i+1}(\phi)$ with $\Delta G^{\text{interiormm}}_{j,j+1,i-1,i}(\phi)$. Thus, we can cache the information specific to the base pair $d\cdot e$ for each given asymmetry the first time it is encountered in an extensible interior loop and then modify only the size information each time it is encountered.

Equation S141 combines the above ideas into a subroutine for computing the interior loop contributions to $R^b_{\text{INTRA}}(i, j, \phi)$.

$$
\begin{aligned}
O(N^3)\ \textsc{InteriorIntra}(i,j,\phi) \equiv & \begin{cases} \bigoplus_{d=i+1}^{\min(i+4,j-5)} \bigoplus_{e=\max(d+4,j-4)}^{j-1} Q^b_{d,e} \otimes W(\Delta G^{\text{interior}}_{i,d,e,j}(\phi,\sigma)), & i+6 \le j \\ \mathbb{0}, & \text{otherwise} \end{cases} \\
\oplus & \begin{cases} \bigoplus_{d=i+1}^{\min(i+4,j-9)} \bigoplus_{e=d+4}^{j-5} Q^b_{d,e} \otimes W(\Delta G^{\text{interior}}_{i,d,e,j}(\phi,\sigma)), & i+10 \le j \\ \mathbb{0}, & \text{otherwise} \end{cases} \\
\oplus & \begin{cases} \bigoplus_{d=i+5}^{j-5} \bigoplus_{e=\max(d+4,j-4)}^{j-1} Q^b_{d,e} \otimes W(\Delta G^{\text{interior}}_{i,d,e,j}(\phi,\sigma)), & i+10 \le j \\ \mathbb{0}, & \text{otherwise} \end{cases} \\
\oplus & \begin{cases} \bigoplus_{s=8}^{j-i-6} (Q^{x,\text{single}}_{i,j,s} \oplus Q^{x,\text{mix}}_{i,j,s}) \otimes W(\Delta G^{\text{interiormm}}_{j-1,j,i,i+1}(\phi)), & i+14 \le j \\ \mathbb{0}, & \text{otherwise} \end{cases} \\
\text{where } \sigma \equiv & \begin{cases} \mu_i, & \textsc{IsSingle}(i,d,\mu_i) \text{ and } \textsc{IsSingle}(e,j,\mu_i) \\ \text{mix}, & \text{otherwise} \end{cases}
\end{aligned}
\tag{S141}
$$

The first three rows handle inextensible interior loops for three cases: 1) $L_1 < 4$ and $L_2 < 4$, 2) $L_1 < 4$ and $L_2 \geq 4$, 3) $L_1 \geq 4$ and $L_2 < 4$. In each case, the contribution of subsequence $[d, e]$ is incorporated using a $Q^b_{d,e}$ element and $\Delta G^{\text{interior}}_{i,d,e,j}(\phi, \sigma)$ is evaluated as for the $O(N^4)$ intrastrand recursion. The fourth row handles extensible interior loops ($L_1 \geq 4$ and $L_2 \geq 4$), by combining a previously computed $Q^x_{i,j,s}$ element for each loop size $s$ with the terminal mismatch free energy, $\Delta G^{\text{interiormm}}_{j-1,j,i,i+1}(\phi)$, corresponding to closing base pair $i \cdot j$. For all four cases, the index limits reflect the steric requirement that there be at least 3 intervening bases between $d$ and $e$. Cached extensible interior loop energies must be kept separate by material so that the appropriate $\Delta G^{\text{interiorsize}}_{s,\mu}$ and $\Delta G^{\text{interiorasymm}}_{|L_1-L_2|,\mu}$ parameters are applied. As a result, $Q^x_{i,j,s}$ is split into $Q^{x,\text{single}}_{i,j,s}$ and $Q^{x,\text{mix}}_{i,j,s}$, which are recombined for each loop size $s$ in Equation S141. There is also a $Q^{x,\text{futuremix}}_{i,j,s}$ element (used to assist in calculating $Q^{x,\text{mix}}_{i,j,s}$) which caches contributions for single-material extensible interior loops using the mixed-material parameter set to cover the scenario where a single-material extensible interior loop becomes a mixed-material extensible interior loop due to a subsequent extension of the loop.

The $R^{x,\text{single}}_{\text{INTRA}}(i, j, s, \phi)$ recursion fills in the three-dimensional tensor $Q^{x,\text{single}}_{i,j,s}$:

$$
\begin{aligned}
R^{x,\text{single}}_{\text{INTRA}}(i,j,s,\phi) \equiv & \begin{cases} C_1 \oplus C_2 \oplus C_3, & j-i > 15 \text{ and } 10 \le s \le j-i-6 \\ C_2 \oplus C_3, & j-i > 14 \text{ and } s = 9 \\ C_2, & j-i > 13 \text{ and } s = 8 \\ \mathbb{0}, & \text{otherwise} \end{cases} \\
\text{where } C_1 \equiv & \begin{cases} Q^{x,\text{single}}_{i+1,j-1,s-2} \otimes W(\Delta G^{\text{interiorsize}}_{s,\mu_i} - \Delta G^{\text{interiorsize}}_{s-2,\mu_i}), & \mu_i = \mu_j = \mu_{i+1} \\ \mathbb{0}, & \text{otherwise} \end{cases} \\
C_2 \equiv & \begin{cases} Q^b_{i+5,j+3-s} \otimes W(\Delta G^{\text{interiorsize}}_{s,\mu_i} + \Delta G^{\text{interiorasymm}}_{s-8,\mu_i} + \Delta G^{\text{interiormm}}_{i+4,i+5,j+3-s,j+4-s}(\phi)), & \textsc{IsSingle}(i,i+5,\mu_i) \text{ and } \\ & \textsc{IsSingle}(j+3-s,j,\mu_i) \\ \mathbb{0}, & \text{otherwise} \end{cases} \\
C_3 \equiv & \begin{cases} Q^b_{s+i-3,j-5} \otimes W(\Delta G^{\text{interiorsize}}_{s,\mu_i} + \Delta G^{\text{interiorasymm}}_{s-8,\mu_i} + \Delta G^{\text{interiormm}}_{s+i-4,s+i-3,j-5,j-4}(\phi)), & \textsc{IsSingle}(i,s+i-3,\mu_i) \text{ and } \\ & \textsc{IsSingle}(j-5,j,\mu_i) \\ \mathbb{0}, & \text{otherwise} \end{cases}
\end{aligned}
\tag{S142}
$$

The $R^{x,\text{futuremix}}_{\text{INTRA}}(i,j,s,\phi)$ recursion fills in the three-dimensional tensor $Q^{x,\text{futuremix}}_{i,j,s}$:

$$R^{x,\text{futuremix}}_{\text{INTRA}}(i,j,s,\phi) \equiv \begin{cases} C_1 \oplus C_2 \oplus C_3, & j-i>15 \text{ and } 10 \le s \le j-i-6 \\ C_2 \oplus C_3, & j-i>14 \text{ and } s=9 \\ C_2, & j-i>13 \text{ and } s=8 \\ \mathbb{0}, & \text{otherwise} \end{cases}$$

$$\text{where } C_1 \equiv \begin{cases} Q^{x,\text{futuremix}}_{i+1,j-1,s-2} \otimes W(\Delta G^{\text{interiorsize}}_{s,\text{mix}} - \Delta G^{\text{interiorsize}}_{s-2,\text{mix}}) & \mu_i = \mu_j = \mu_{i+1} \\ \mathbb{0} & \text{otherwise} \end{cases}$$

$$C_2 \equiv \begin{cases} Q^{b}_{i+5,j+3-s} \otimes W(\Delta G^{\text{interiorsize}}_{s,\text{mix}} + \Delta G^{\text{interiorasymm}}_{s-8,\text{mix}} + \Delta G^{\text{interiormm}}_{i+4,i+5,j+3-s,j+4-s}(\phi)) & \textsc{IsSingle}(i,i+5,\mu_i) \text{ and } \textsc{IsSingle}(j+3-s,j,\mu_i) \\ \mathbb{0} & \text{otherwise} \end{cases}$$

$$C_3 \equiv \begin{cases} Q^{b}_{s+i-3,j-5} \otimes W(\Delta G^{\text{interiorsize}}_{s,\text{mix}} + \Delta G^{\text{interiorasymm}}_{s-8,\text{mix}} + \Delta G^{\text{interiormm}}_{s+i-4,s+i-3,j-5,j-4}(\phi)) & \textsc{IsSingle}(i,s+i-3,\mu_i) \text{ and } \textsc{IsSingle}(j-5,j,\mu_i) \\ \mathbb{0} & \text{otherwise} \end{cases} \tag{S143}$$

The $R^{x,\text{mix}}_{\text{INTRA}}(i,j,s,\phi)$ recursion fills in the three-dimensional tensor $Q^{x,\text{mix}}_{i,j,s}$:

$$R^{x,\text{mix}}_{\text{INTRA}}(i,j,s,\phi) \equiv \begin{cases} C_1 \oplus C_2 \oplus C_3, & j-i>15 \text{ and } 10 \le s \le j-i-6 \\ C_2 \oplus C_3, & j-i>14 \text{ and } s=9 \\ C_2, & j-i>13 \text{ and } s=8 \\ \mathbb{0}, & \text{otherwise} \end{cases}$$

$$\text{where } C_1 \equiv Q^{x,\text{mix}}_{i+1,j-1,s-2} \otimes W(\Delta G^{\text{interiorsize}}_{s,\text{mix}} - \Delta G^{\text{interiorsize}}_{s-2,\text{mix}}) \oplus \begin{cases} \mathbb{0} & \mu_i = \mu_j = \mu_{i+1} \\ Q^{x,\text{futuremix}}_{i+1,j-1,s-2} \otimes W(\Delta G^{\text{interiorsize}}_{s,\text{mix}} - \Delta G^{\text{interiorsize}}_{s-2,\text{mix}}) & \text{otherwise} \end{cases}$$

$$C_2 \equiv \begin{cases} \mathbb{0} & \textsc{IsSingle}(i,i+5,\mu_i) \text{ and } \textsc{IsSingle}(j+3-s,j,\mu_i) \\ Q^{b}_{i+5,j+3-s} \otimes W(\Delta G^{\text{interiorsize}}_{s,\text{mix}} + \Delta G^{\text{interiorasymm}}_{s-8,\text{mix}} + \Delta G^{\text{interiormm}}_{i+4,i+5,j+3-s,j+4-s}(\phi)) & \text{otherwise} \end{cases}$$

$$C_3 \equiv \begin{cases} \mathbb{0} & \textsc{IsSingle}(i,s+i-3,\mu_i) \text{ and } \textsc{IsSingle}(j-5,j,\mu_i) \\ Q^{b}_{s+i-3,j-5} \otimes W(\Delta G^{\text{interiorsize}}_{s,\text{mix}} + \Delta G^{\text{interiorasymm}}_{s-8,\text{mix}} + \Delta G^{\text{interiormm}}_{s+i-4,s+i-3,j-5,j-4}(\phi)) & \text{otherwise} \end{cases} \tag{S144}$$

In $Q^{x,\text{single}}_{i,j,s}$, $Q^{x,\text{futuremix}}_{i,j,s}$, and $Q^{x,\text{mix}}_{i,j,s}$, the indices $i$ and $j$ refer to the closing base pair $i \cdot j$ while the index $s$ refers to the size of the extensible loops (sum of the two side lengths). Contributions can be divided into two classes: previously encountered interior loops (handled in each recursion equation with term $C_1$) and new interior loops (handled in each recursion equation with terms $C_2$ and $C_3$).

In $R^{x,\text{single}}_{\text{INTRA}}(i,j,s,\phi)$, previously encountered single-material loops starting at $i+1$ and $j-1$ that remain single-material loops after extension to start at $i$ and $j$ (checked with the conditional $\mu_i = \mu_j = \mu_{i+1}$) are incorporated by accessing the previously computed element $Q^{x,\text{single}}_{i+1,j-1,s-2}$ and replacing $\Delta G^{\text{interiorsize}}_{s-2,\mu_i}$ with $\Delta G^{\text{interiorsize}}_{s,\mu_i}$ (see term $C_1$ in (S142)). This is the key operation that reduces the complexity of the interior loop recursion to $O(N)$ by capturing all previous loops in $O(1)$. If extending the single-material loop leads to a mixed-material loop (because $i$ and/or $j$ are a different material than the previously cached loops, reflected in the material of $i+1$), the contribution to $R^{x,\text{single}}_{\text{INTRA}}(i,j,s,\phi)$ is $\mathbb{0}$ (see term $C_1$ in (S142)) and the previously cached loops are instead extended as mixed-material loops (see term $C_1$ in (S144)).

This extension of previously single-material loops to become mixed-material loops requires the $R^{x,\text{futuremix}}_{\text{INTRA}}(i,j,s,\phi)$ recursion and $Q^{x,\text{futuremix}}_{i,j,s}$ tensor. $Q^{x,\text{futuremix}}_{i,j,s}$ caches contributions for the same loops as $Q^{x,\text{single}}_{i,j,s}$, but uses mixed-material parameters instead of single-material parameters. The $R^{x,\text{futuremix}}_{\text{INTRA}}(i,j,s,\phi)$ recursion equations (S143) are thus identical to $R^{x,\text{single}}_{\text{INTRA}}(i,j,s,\phi)$ those for the (S142) aside from the free energy parameters. For $R^{x,\text{futuremix}}_{\text{INTRA}}(i,j,s,\phi)$, previously encountered single-material loops starting at $i+1$ and $j-1$ that remain single-material loops after extension to start at $i$ and $j$ (checked with the conditional $\mu_i = \mu_j = \mu_{i+1}$) are incorporated by accessing the previously computed element $Q^{x,\text{futuremix}}_{i+1,j-1,s-2}$ and replacing $\Delta G^{\text{interiorsize}}_{s-2,\text{mix}}$ with $\Delta G^{\text{interiorsize}}_{s,\text{mix}}$ (see term $C_1$ in (S143)). Otherwise, if the conditional does not pass because $i$ and/or $j$ introduce a new material to the loops, previously encountered single-material loops are incorporated in $R^{x,\text{mix}}_{\text{INTRA}}(i,j,s,\phi)$ by accessing the previously computed element $Q^{x,\text{futuremix}}_{i+1,j-1,s-2}$ and replacing $\Delta G^{\text{interiorsize}}_{s-2,\text{mix}}$ with $\Delta G^{\text{interiorsize}}_{s,\text{mix}}$ (see term $C_1$ in (S144)). Caching single-material loop contributions with mixed parameters using $R^{x,\text{futuremix}}_{\text{INTRA}}(i,j,s,\phi)$ is necessary as otherwise there is no convenient way to replace a previously incorporated single-material asymmetry parameter, $\Delta G^{\text{interiorasymm}}_{|L_1-L_2|,\mu_i}$, with the corresponding mixed-material parameter, $\Delta G^{\text{interiorasymm}}_{|L_1-L_2|,\text{mix}}$, when extension leads to a transition from single-material to mixed-material. For $R^{x,\text{mix}}_{\text{INTRA}}(i,j,s,\phi)$, previously encountered mixed-material loops are incorporated by accessing the previously computed element $Q^{x,\text{mix}}_{i+1,j-1,s-2}$ and replacing $\Delta G^{\text{interiorsize}}_{s-2,\text{mix}}$ with $\Delta G^{\text{interiorsize}}_{s,\text{mix}}$ (see term $C_1$ in (S144)).

Note that the terminal mismatch contribution of the closing base pair $i \cdot j$ is not incorporated in the $Q^x$ element, but is combined with $Q^x$ in (S141), so there is never a need to replace one terminal mismatch contribution with another as the loop is extended. New extensible loops that are first encountered for the indices $i, j, s$ (elements that have exactly $L_1 = 4$ or $L_2 = 4$ or both) are handled by the $C_2$ and $C_3$ terms in $R^{x,\text{single}}_{\text{INTRA}}(i,j,s,\phi)$, $R^{x,\text{futuremix}}_{\text{INTRA}}(i,j,s,\phi)$, and $R^{x,\text{mix}}_{\text{INTRA}}(i,j,s,\phi)$. The material of each new loop is checked and the contribution is cached in the appropriate tensor. When $L_1 = 4$ (handled in the $C_2$ terms), the conditional "$\textsc{IsSingle}(i,i+5,\mu_i)$ and $\textsc{IsSingle}(j+3-s,j,\mu_i)$" is checked. When $L_2 = 4$ (handled in the $C_3$ terms), the conditional "$\textsc{IsSingle}(i,s+i-3,\mu_i)$ and $\textsc{IsSingle}(j-5,j,\mu_i)$" is checked. If the conditionals pass, then the new loop contains a single material and the contribution is cached in the $Q^{x,\text{single}}_{i,j,s}$ and $Q^{x,\text{futuremix}}_{i,j,s}$ tensors by the $R^{x,\text{single}}_{\text{INTRA}}(i,j,s,\phi)$ and $R^{x,\text{futuremix}}_{\text{INTRA}}(i,j,s,\phi)$ recursions. If the conditionals fail, then the new loop contains mixed materials and the contribution is cached in the $Q^{x,\text{mix}}_{i,j,s}$ tensor by the $R^{x,\text{mix}}_{\text{INTRA}}(i,j,s,\phi)$ recursion. Note that the subexpressions $C_2$ and $C_3$ are coincident for $L_1 = L_2 = 4$ ($s = 8$).

Crucially, to calculate a new value $Q^x_{i,j,s}$ for a subsequence of length $l = j - i + 1$ for any material, only elements of the form $Q^x_{i+1,j-1,s-2}$ are accessed (for $O(N)$ values of $s$; each value of $l$ corresponds to a diagonal of the intrastrand block). Therefore, we only need to store elements of $Q^x$ for subsequences of length $l, l-1$, and $l-2$ (corresponding to the current diagonal and the two previous diagonals). In other words, only $Q^x$ values corresponding to 3 diagonals need to exist in memory during the forward pass. In moving to the next diagonal $l+1$, we can simply delete all $Q^x_{i,j,s}$ values for diagonal $l-2$ as they will not be accessed again. Hence, only $O(N^2)$ space is necessary to store the needed elements of $Q^x$. Naively storing all of $Q^x_{i,j,s}$ would have needlessly increased the space complexity to $O(N^3)$.

The above recursions enable calculation of interior loop contributions for forward algorithms with $O(N^3)$ time complexity and $O(N^2)$ space complexity. However, this approach is incompatible with backtracking algorithms as the optimization of throwing away $Q^x$ values that are no longer needed during the forward sweep, implies that they are also no longer available for backtracking after the forward sweep is complete. One option is to reconstruct the $Q^x$ values during backtracking, but this incurs $O(N^3)$ time complexity and can lead to loss of precision for large complex ensembles.[40] Following Section S4.5 of Reference 1, another option that we pursue here is to use a different iteration pattern through the $O(N^4)$ interior loop recursions during backtracking. With this option, we exploit the fact that unlike forward algorithms that evaluate recursive elements for all $i$ and $j$ in a forward sweep, backtracking algorithms evaluate only a subset of all possible recursive elements. Hence, the worst-case time complexity can be kept at $O(N^2)$ per structure for our backtracking algorithms.

**$O(N^3)$ intrastrand interior loop recursion with fast conditional checks.** For clarity, the $O(N^3)$ recursions of the previous section include explicit conditionals to check for single- and mixed-material scenarios. However, in practice, this material information can be determined more efficiently by introducing an additional $Q^{xa}$ recursion. This approach to fast conditional checks for interior loops speeds up the overall mixed-material algorithm by approximately a factor of 2 so that it is approximately 2–3.5× slower than the single-material algorithm (rather than 4–7× slower than the single-material algorithm).

With fast conditional checks, instead of checking the material for both sides of every newly encountered loop using $\textsc{IsSingle}$ conditionals as in (S142)–(S144), it suffices to check the smallest possible newly encountered loop for material ($L_1 = L_2 = 4$, checked with conditional "$\textsc{IsSingle}(i,i+5,\mu_i)$ and $\textsc{IsSingle}(j-5,j,\mu_i)$"). All other newly encountered loops must contain this smallest loop. Therefore, if this single-material loop is stored in $R^{x,\text{single}}_{\text{INTRA}}(i,j,s,\phi)$, then only the longer side of newly encountered loops with either $L_1 = 4, L_2 > 4$ or $L_1 > 4, L_2 = 4$ needs to be checked for material (in terms $C_2$ and $C_3$ for $R^{x,\text{single}}_{\text{INTRA}}(i,j,s,\phi)$ and $R^{x,\text{futuremix}}_{\text{INTRA}}(i,j,s,\phi)$) in (S145) and (S146). If the longer side is a single material, then this must be the same material as the shorter side since both sides have already been checked in the smallest possible loop, so the new larger loop is also a single-material loop. Checking one side can be done efficiently without conditionals using a new $Q^{xa}$ recursion that allows for an efficient vectorized implementation. $Q^{xa,\text{single}}_{i,j}$ returns $\mathbb{1}$ if the subsequence $[i,j]$ is composed of a single material, and $\mathbb{0}$ otherwise, so that multiplying by $Q^{xa}$ is equivalent to using a conditional to check whether the subsequence is a single material.

For extending loops, the conditional $\mu_i = \mu_j = \mu_{i+1}$ is already checked by the conditional "$\textsc{IsSingle}(i,i+5,\mu_i)$ and $\textsc{IsSingle}(j-5,j,\mu_i)$". If "$\textsc{IsSingle}(i,i+5,\mu_i)$ and $\textsc{IsSingle}(j-5,j,\mu_i)$" passes, then $\mu_i = \mu_j = \mu_{i+1}$ must pass and single material loops can be extended as single-material (in term $C_1$ for $R^{x,\text{single}}_{\text{INTRA}}(i,j,s,\phi)$ and $R^{x,\text{futuremix}}_{\text{INTRA}}(i,j,s,\phi)$ in (S145) and (S146)). For cases where $\mu_i = \mu_j = \mu_{i+1}$ passes but "$\textsc{IsSingle}(i,i+5,\mu_i)$ and $\textsc{IsSingle}(j-5,j,\mu_i)$" does not pass, the smallest possible loop of the $[i+1, j-1]$ subsequence must contain mixed materials, meaning that $Q^{x,\text{single}}_{i+1,j-1,s-2}$ is $\mathbb{0}$ in any case. Thus, it suffices to check just "$\text{IsSingle}(i, i+5, \mu_i)$ and $\text{IsSingle}(j-5, j, \mu_i)$" in the beginning. If "$\text{IsSingle}(i, i+5, \mu_i)$ and $\text{IsSingle}(j-5, j, \mu_i)$" does not pass, then $R^{x,\text{single}}_{\text{INTRA}}(i, j, s, \phi) = \mathbb{0}$, since all newly encountered loops must be mixed-material, and there are no single-material loops to be extended.

For the $R^{x,\text{mix}}_{\text{INTRA}}(i, j, s, \phi)$ recursion in (S147), if the conditional "$\text{IsSingle}(i, i+5, \mu_i)$ and $\text{IsSingle}(j-5, j, \mu_i)$" passes (handled by term $I_{\text{single}}$), larger loops containing this loop may still be mixed-material. The longer side of newly encountered loops with either $L_1 = 4, L_2 > 4$ or $L_1 > 4, L_2 = 4$ is checked for mixed materials by multiplying by $Q^{xa,\text{mix}}$ (in terms $C_{2s}$ and $C_{3s}$). The recursion $Q^{xa,\text{mix}}_{i,j}$ returns $\mathbb{1}$ if the subsequence $[i, j]$ is mixed-material, and $\mathbb{0}$ otherwise. Mixed-material loops are extended regardless, since once a loop is mixed it is always mixed (in term $C_{1s}$). That is to say, even if "$\text{IsSingle}(i, i+5, \mu_i)$ and $\text{IsSingle}(j-5, j, \mu_i)$" passes, any non-zero contribution from $Q^{x,\text{mix}}_{i+1,j-1,s-2}$ must represent extensible interior loops which contain mixture beyond $[i+1, i+5]$ or $[j-1, j-5]$ which must continue to be extended as mixed when adding nucleotides $i$ and $j$. Adding $i$ and $j$, regardless of their material, would not change the mixed status of these loops. Single-material loops are not extended as mixed in this case, since if "$\text{IsSingle}(i, i+5, \mu_i)$ and $\text{IsSingle}(j-5, j, \mu_i)$" passed then no new materials were introduced by nucleotides $i$ and $j$.

For the $R^{x,\text{mix}}_{\text{INTRA}}(i, j, s, \phi)$ recursion in (S147), if "$\text{IsSingle}(i, i+5, \mu_i)$ and $\text{IsSingle}(j-5, j, \mu_i)$" does not pass (handled by term $I_{\text{mix}}$), then all newly encountered loops are mixed since the smallest loop is mixed. No further conditional checks are needed when computing a newly encountered loop (in terms $C_{2m}$ and $C_{3m}$). Mixed-material loops are extended regardless, since once a loop is mixed it is always mixed (in term $C_{1m}$). Single-material loops are also extended as mixed using $Q^{x,\text{futuremix}}_{i+1,j-1,s-2}$ (in term $C_{1m}$). This is because if "$\text{IsSingle}(i, i+5, \mu_i)$ and $\text{IsSingle}(j-5, j, \mu_i)$" does not pass, then either new materials were introduced by nucleotides $i$ and $j$, requiring single material loops to be extended as mixed, or the smallest possible loop of the $[i+1, j-1]$ subsequence contains mixed materials, meaning that $Q^{x,\text{futuremix}}_{i+1,j-1,s-2}$ is $\mathbb{0}$ in any case.

The $R^{x,\text{single}}_{\text{INTRA}}(i, j, s, \phi)$ recursion fills in the three-dimensional tensor $Q^{x,\text{single}}_{i,j,s}$:

$$R^{x,\text{single}}_{\text{INTRA}}(i,j,s,\phi) \equiv \begin{cases} C_0, & j-i > 13 \\ \mathbb{0}, & \text{otherwise} \end{cases}$$

$$C_0 \equiv \begin{cases} I_{\text{single}}, & \text{IsSingle}(i,i+5,\mu_i) \text{ and } \text{IsSingle}(j-5,j,\mu_i) \\ \mathbb{0}, & \text{otherwise} \end{cases}$$

$$\text{where } I_{\text{single}} \equiv \begin{cases} C_1 \oplus C_2 \oplus C_3, & j-i > 15 \text{ and } 10 \le s \le j-i-6 \\ C_2 \oplus C_3, & j-i > 14 \text{ and } s = 9 \\ C_2, & s = 8 \\ \mathbb{0}, & \text{otherwise} \end{cases} \tag{S145}$$

$$\begin{aligned} \text{with } C_1 &\equiv Q^{x,\text{single}}_{i+1,j-1,s-2} \otimes W(\Delta G^{\text{interiorsize}}_{s,\mu_i} - \Delta G^{\text{interiorsize}}_{s-2,\mu_i}) \\ C_2 &\equiv Q^b_{i+5,j+3-s} \otimes W(\Delta G^{\text{interiorsize}}_{s,\mu_i} + \Delta G^{\text{interiorasymm}}_{s-8,\mu_i} + \Delta G^{\text{interiormm}}_{i+4,i+5,j+3-s,j+4-s}(\phi)) \otimes Q^{xa,\text{single}}_{j+3-s,j} \\ C_3 &\equiv Q^b_{s+i-3,j-5} \otimes W(\Delta G^{\text{interiorsize}}_{s,\mu_i} + \Delta G^{\text{interiorasymm}}_{s-8,\mu_i} + \Delta G^{\text{interiormm}}_{s+i-4,s+i-3,j-5,j-4}(\phi)) \otimes Q^{xa,\text{single}}_{i,s+i-3} \end{aligned}$$

The $R^{x,\text{futuremix}}_{\text{INTRA}}(i,j,s,\phi)$ recursion fills in the three-dimensional tensor $Q^{x,\text{futuremix}}_{i,j,s}$:

$$
\begin{aligned}
R^{x,\text{futuremix}}_{\text{INTRA}}(i,j,s,\phi) &\equiv \begin{cases} C_0, & j-i>13 \\ \mathbb{0}, & \text{otherwise} \end{cases} \\
C_0 &\equiv \begin{cases} I_{\text{futuremix}}, & \textsc{IsSingle}(i,i+5,\mu_i) \text{ and } \textsc{IsSingle}(j-5,j,\mu_i) \\ \mathbb{0}, & \text{otherwise} \end{cases} \\
\text{where } I_{\text{futuremix}} &\equiv \begin{cases} C_1 \oplus C_2 \oplus C_3, & j-i>15 \text{ and } 10 \le s \le j-i-6 \\ C_2 \oplus C_3, & j-i>14 \text{ and } s=9 \\ C_2, & s=8 \\ \mathbb{0}, & \text{otherwise} \end{cases} \\
\text{with } C_1 &\equiv Q^{x,\text{futuremix}}_{i+1,j-1,s-2} \otimes W(\Delta G^{\text{interiorsize}}_{s,\text{mix}} - \Delta G^{\text{interiorsize}}_{s-2,\text{mix}}) \\
C_2 &\equiv Q^{b}_{i+5,j+3-s} \otimes W(\Delta G^{\text{interiorsize}}_{s,\text{mix}} + \Delta G^{\text{interiorasymm}}_{s-8,\text{mix}} + \Delta G^{\text{interiormm}}_{i+4,i+5,j+3-s,j+4-s}(\phi)) \otimes Q^{xa,\text{single}}_{j+3-s,j} \\
C_3 &\equiv Q^{b}_{s+i-3,j-5} \otimes W(\Delta G^{\text{interiorsize}}_{s,\text{mix}} + \Delta G^{\text{interiorasymm}}_{s-8,\text{mix}} + \Delta G^{\text{interiormm}}_{s+i-4,s+i-3,j-5,j-4}(\phi)) \otimes Q^{xa,\text{single}}_{i,s+i-3}
\end{aligned}
\tag{S146}
$$

The $R^{x,\text{mix}}_{\text{INTRA}}(i,j,s,\phi)$ recursion fills in the three-dimensional tensor $Q^{x,\text{mix}}_{i,j,s}$:

$$
\begin{aligned}
R^{x,\text{mix}}_{\text{INTRA}}(i,j,s,\phi) &\equiv \begin{cases} C_0, & j-i>13 \\ \mathbb{0}, & \text{otherwise} \end{cases} \\
C_0 &\equiv \begin{cases} I_{\text{single}}, & \textsc{IsSingle}(i,i+5,\mu_i) \text{ and } \textsc{IsSingle}(j-5,j,\mu_i) \\ I_{\text{mix}}, & \text{otherwise} \end{cases} \\
I_{\text{single}} &\equiv \begin{cases} C_{1s} \oplus C_{2s} \oplus C_{3s}, & j-i>15 \text{ and } 10 \le s \le j-i-6 \\ C_{2s} \oplus C_{3s}, & j-i>14 \text{ and } s=9 \\ C_{2s}, & s=8 \\ \mathbb{0}, & \text{otherwise} \end{cases} \\
\text{where } C_{1s} &\equiv Q^{x,\text{mix}}_{i+1,j-1,s-2} \otimes W(\Delta G^{\text{interiorsize}}_{s,\text{mix}} - \Delta G^{\text{interiorsize}}_{s-2,\text{mix}}) \\
C_{2s} &\equiv Q^{b}_{i+5,j+3-s} \otimes W(\Delta G^{\text{interiorsize}}_{s,\text{mix}} + \Delta G^{\text{interiorasymm}}_{s-8,\text{mix}} + \Delta G^{\text{interiormm}}_{i+4,i+5,j+3-s,j+4-s}(\phi)) \otimes Q^{xa,\text{mix}}_{j+3-s,j} \\
C_{3s} &\equiv Q^{b}_{s+i-3,j-5} \otimes W(\Delta G^{\text{interiorsize}}_{s,\text{mix}} + \Delta G^{\text{interiorasymm}}_{s-8,\text{mix}} + \Delta G^{\text{interiormm}}_{s+i-4,s+i-3,j-5,j-4}(\phi)) \otimes Q^{xa,\text{mix}}_{i,s+i-3} \\
I_{\text{mix}} &\equiv \begin{cases} C_{1m} \oplus C_{2m} \oplus C_{3m}, & j-i>15 \text{ and } 10 \le s \le j-i-6 \\ C_{2m} \oplus C_{3m}, & j-i>14 \text{ and } s=9 \\ C_{2m}, & s=8 \\ \mathbb{0}, & \text{otherwise} \end{cases} \\
\text{where } C_{1m} &\equiv Q^{x,\text{mix}}_{i+1,j-1,s-2} \otimes W(\Delta G^{\text{interiorsize}}_{s,\text{mix}} - \Delta G^{\text{interiorsize}}_{s-2,\text{mix}}) \oplus Q^{x,\text{futuremix}}_{i+1,j-1,s-2} \otimes W(\Delta G^{\text{interiorsize}}_{s,\text{mix}} - \Delta G^{\text{interiorsize}}_{s-2,\text{mix}}) \\
C_{2m} &\equiv Q^{b}_{i+5,j+3-s} \otimes W(\Delta G^{\text{interiorsize}}_{s,\text{mix}} + \Delta G^{\text{interiorasymm}}_{s-8,\text{mix}} + \Delta G^{\text{interiormm}}_{i+4,i+5,j+3-s,j+4-s}(\phi)) \\
C_{3m} &\equiv Q^{b}_{s+i-3,j-5} \otimes W(\Delta G^{\text{interiorsize}}_{s,\text{mix}} + \Delta G^{\text{interiorasymm}}_{s-8,\text{mix}} + \Delta G^{\text{interiormm}}_{s+i-4,s+i-3,j-5,j-4}(\phi))
\end{aligned}
\tag{S147}
$$

***$R^{xa}_{\text{INTRA}}$* recursion.** The $R^{x}_{\text{INTRA}}(i,j,\phi)$ recursion references $Q^{xa}$ elements that are computed using the $R^{xa}_{\text{INTRA}}$ recursion shown in Figure S18. $R^{xa}_{\text{INTRA}}(i,j,\phi)$ treats the conditional ensemble of subsequence $[i,j]$ in interior loop context containing no terminal base pairs (the hollow dotted line between $i$ and $j$ indicates that there is no base pairing in $[i,j]$). This recursion is used in the mixed-material algorithm to efficiently check the material of subsequence $[i,j]$. The recursion equation and diagram are displayed in Figure S18. To allow both single- and mixed-material checks, $R^{xa}_{\text{INTRA}}(i,j,\phi)$ is split into $R^{xa,\text{single}}_{\text{INTRA}}(i,j,\phi)$ and $R^{xa,\text{mix}}_{\text{INTRA}}(i,j,\phi)$ (the single- and mixed-material recursions are indicated in the recursion diagram by blue and red backbones, respectively):

- $R^{xa,\text{single}}_{\text{INTRA}}(i, j, \phi)$: This recursion checks if subsequence $[i, j]$ is composed of a single material, returning $\mathbb{1}$ if true and $\mathbb{0}$ if false. If $i = j$, then the subsequence is a single nucleotide and therefore a single material, yielding $\mathbb{1}$. When $i < j$, the $Q^{xa,\text{single}}_{i,j-1}$ element is used to represent the single-material status of subsequence $[i, j-1]$, which is $\mathbb{1}$ only if $[i, j-1]$ is a single material. $Q^{xa,\text{single}}_{i,j}$ is set equal to $Q^{xa,\text{single}}_{i,j-1}$ if the material of $j$ matches subsequence $[i, j-1]$ (checked with conditional $\mu_j = \mu_{j-1}$). Otherwise, $R^{xa,\text{single}}_{\text{INTRA}}(i, j, \phi)$ is $\mathbb{0}$, since the subsequence is not of a single material; thus future encounters with longer subsequences starting from $i$ will also have a value of $\mathbb{0}$ since they will also be mixed.
- $R^{xa,\text{mix}}_{\text{INTRA}}(i, j, \phi)$: This recursion checks if subsequence $[i, j]$ is composed of mixed materials, returning $\mathbb{1}$ if true and $\mathbb{0}$ if false. If $i = j$, the recursion returns $\mathbb{0}$ since a single nucleotide cannot have more than one material. If $i < j$, the $Q^{xa,\text{mix}}_{i,j-1}$ element is used to represent the mixed-material status of subsequence $[i, j-1]$, and is always included as a contribution to $Q^{xa,\text{mix}}_{i,j}$ since $[i, j]$ must be mixed if $[i, j-1]$ is mixed. If the material of $j$ does not match subsequence $[i, j-1]$, then any single-material contribution from $[i, j-1]$ is now included in $R^{xa,\text{mix}}_{\text{INTRA}}(i, j, \phi)$ (since $[i, j]$ is no longer single-material) using $Q^{xa,\text{single}}_{i,j-1}$ (indicated by term $C_1$).

Note that the $Q^{xa}$ recursion is simply the $Q^{ma}$ recursion with $W(\Delta G^{\text{multi}}_{\text{nt},\mu_i})$ replaced by $\mathbb{1}$ (Figure S18), and applied in an interior loop context instead of a multiloop context.

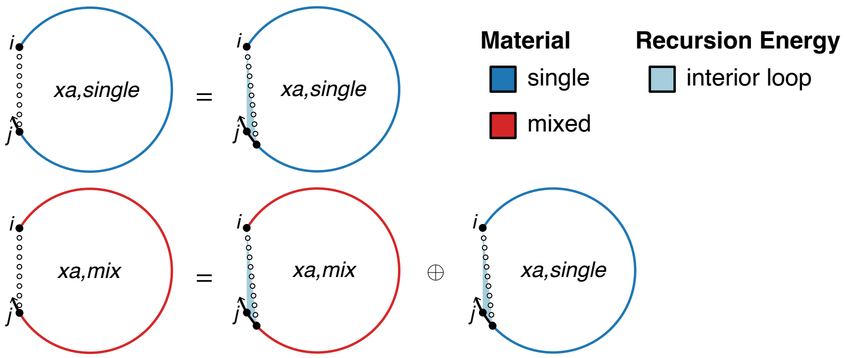

$$R^{xa,\text{single}}_{\text{INTRA}}(i, j, \phi) \equiv \begin{cases} \mathbb{1}, & i = j \\ Q^{xa,\text{single}}_{i,j-1} \otimes \mathbb{1}, & i < j \text{ and } \mu_j = \mu_{j-1} \\ \mathbb{0}, & \text{otherwise} \end{cases}$$

$$R^{xa,\text{mix}}_{\text{INTRA}}(i, j, \phi) \equiv \begin{cases} Q^{xa,\text{mix}}_{i,j-1} \otimes \mathbb{1} \oplus C_1, & i < j \\ \mathbb{0}, & \text{otherwise} \end{cases}$$

$$C_1 \equiv \begin{cases} Q^{xa,\text{single}}_{i,j-1} \otimes \mathbb{1}, & \mu_j \neq \mu_{j-1} \\ \mathbb{0}, & \text{otherwise} \end{cases}$$

Figure S18: $R^{xa}_{\text{INTRA}}$ recursion. Top: recursion diagram. Bottom: recursion equation. This recursion is used for interior loops and does not change depending on whether coaxial and dangle stacking are included for multiloops and exterior loops.

##### S6.4.2 Interstrand interior loop dynamic programming recursions

**$O(N^4)$ interstrand interior loop recursion.** The interstrand $O(N^4)$ interior loop contribution:

$$O(N^4)\ \text{INTERIORINTER}(i, j, \phi) \equiv \begin{cases} \bigoplus_{d=i+1}^{m-1} \bigoplus_{e=n}^{j-1} \{Q^b_{d,e} \otimes W(\Delta G^{\text{interior}}_{i,d,e,j}(\phi, \sigma))\}, & i < m-1 \text{ and } n < j \\ \mathbb{0}, & \text{otherwise} \end{cases}$$
$$\text{where } \sigma \equiv \begin{cases} \mu_i, & \text{ISSINGLE}(i, d, \mu_i) \text{ and ISSINGLE}(e, j, \mu_i) \\ \text{mix}, & \text{otherwise} \end{cases} \tag{S148}$$
$$\text{where } m = \text{FIRST}(\eta)$$
$$n = \text{LAST}(\eta)$$

proceeds in the same general manner as the intrastrand case (S139), considering interior loops in order of ascending $d$ then $e$ indices. However, $d$ is restricted to be on the first strand ($d < m$) and $e$ is restricted to be on the last strand ($e \geq n$), as reflected in the upper summation limit for $d$ and the lower summation limit for $e$. These two requirements ensure that there are no nicks between $i$ and $d$ and between $e$ and $j$, preventing exterior loop states (that are not treated in this interior loop recursion) and disconnected states (that are not part of the complex ensemble).

***O*(*N*³) interstrand interior loop recursion.** Interior loop contributions for elements in interstrand blocks are computed with $O(N^3)$ time complexity using the subroutine:

$$
\begin{aligned}
O(N^3)\ \textsc{InteriorInter}(i,j,\phi) \equiv & \begin{cases} \bigoplus_{d=i+1}^{\min(i+4,m-1)} \bigoplus_{e=\max(n,j-4)}^{j-1} Q^b_{d,e} \otimes W(\Delta G^{\text{interior}}_{i,d,e,j}(\phi,\sigma)), & i+1 < m \text{ and } n < j \\ \mathbb{0}, & \text{otherwise} \end{cases} \\
\oplus & \begin{cases} \bigoplus_{d=i+1}^{\min(i+4,m-1)} \bigoplus_{e=n}^{j-5} Q^b_{d,e} \otimes W(\Delta G^{\text{interior}}_{i,d,e,j}(\phi,\sigma)), & i+1 < m \text{ and } n+4 < j \\ \mathbb{0}, & \text{otherwise} \end{cases} \\
\oplus & \begin{cases} \bigoplus_{d=i+5}^{m-1} \bigoplus_{e=\max(n,j-4)}^{j-1} Q^b_{d,e} \otimes W(\Delta G^{\text{interior}}_{i,d,e,j}(\phi,\sigma)), & i+5 < m \text{ and } n < j \\ \mathbb{0}, & \text{otherwise} \end{cases} \\
\oplus & \begin{cases} \bigoplus_{s=8}^{j-n+m-i-3} (Q^{x,\text{single}}_{i,j,s} \oplus Q^{x,\text{mix}}_{i,j,s}) \otimes W(\Delta G^{\text{interiormm}}_{j-1,j,i,i+1}(\phi)), & i+5 < m \text{ and } n+4 < j \\ \mathbb{0}, & \text{otherwise} \end{cases} \\
\text{where } \sigma \equiv & \begin{cases} \mu_i & \textsc{IsSingle}(i,d,\mu_i) \text{ and } \textsc{IsSingle}(e,j,\mu_i) \\ \text{mix} & \text{otherwise} \end{cases} \\
\text{where } m &= \textsc{First}(\eta) \\
n &= \textsc{Last}(\eta)
\end{aligned}
\tag{S149}
$$

The approach is analogous to that of (S141). Index limits are modified to ensure that $d$ is on the same strand as $i$ and $e$ is on the same strand as $j$, preventing exterior loop states (that are not treated in this interior loop recursion) and disconnected states (that are not part of the complex ensemble).

The recursion $R^{x,\text{single}}_{\textsc{Inter}}(i,j,s,\phi)$ is closely related to (S142):

$$
R^{x,\text{single}}_{\textsc{Inter}}(i,j,s,\phi) \equiv \begin{cases} C_1 \oplus C_2 \oplus C_3, & i+6 < m \text{ and } n+5 < j \text{ and } 10 \leq s \leq j-i+m-n-3 \\ C_2 \oplus C_3, & i+6 < m \text{ and } n+5 < j \text{ and } s = 9 \\ C_3, & i+6 < m \text{ and } n+5 = j \text{ and } s = 9 \\ C_2, & i+6 = m \text{ and } n+5 \leq j \text{ and } s = 9 \\ C_2, & i+6 \leq m \text{ and } n+5 \leq j \text{ and } s = 8 \\ \mathbb{0}, & \text{otherwise} \end{cases}
$$

$$
\begin{aligned}
\text{where } m &= \textsc{First}(\eta) \\
n &= \textsc{Last}(\eta) \\
C_1 &\equiv \begin{cases} Q^{x,\text{single}}_{i+1,j-1,s-2} \otimes W(\Delta G^{\text{interiorsize}}_{s,\mu_i} - \Delta G^{\text{interiorsize}}_{s-2,\mu_i}) & \mu_i = \mu_j = \mu_{i+1} \\ \mathbb{0} & \text{otherwise} \end{cases} \\
C_2 &\equiv \begin{cases} Q^b_{i+5,j+3-s} \otimes W(\Delta G^{\text{interiorsize}}_{s,\mu_i} + \Delta G^{\text{interiorasymm}}_{s-8,\mu_i} + \Delta G^{\text{interiormm}}_{i+4,i+5,j+3-s,j+4-s}(\phi)) & \textsc{IsSingle}(i,i+5,\mu_i) \text{ and } \\ & \textsc{IsSingle}(j+3-s,j,\mu_i) \\ \mathbb{0} & \text{otherwise} \end{cases} \\
C_3 &\equiv \begin{cases} Q^b_{s+i-3,j-5} \otimes W(\Delta G^{\text{interiorsize}}_{s,\mu_i} + \Delta G^{\text{interiorasymm}}_{s-8,\mu_i} + \Delta G^{\text{interiormm}}_{s+i-4,s+i-3,j-5,j-4}(\phi)) & \textsc{IsSingle}(i,s+i-3,\mu_i) \text{ and } \\ & \textsc{IsSingle}(j-5,j,\mu_i) \\ \mathbb{0} & \text{otherwise} \end{cases}
\end{aligned}
\tag{S150}
$$

The recursion $R^{x,\text{futuremix}}_{\text{INTER}}(i,j,s,\phi)$ is closely related to (S143):

$$R^{x,\text{futuremix}}_{\text{INTER}}(i,j,s,\phi) \equiv \begin{cases} C_1 \oplus C_2 \oplus C_3, & i+6<m \text{ and } n+5<j \text{ and } 10 \leq s \leq j-i+m-n-3 \\ C_2 \oplus C_3, & i+6<m \text{ and } n+5<j \text{ and } s=9 \\ C_3, & i+6<m \text{ and } n+5=j \text{ and } s=9 \\ C_2, & i+6=m \text{ and } n+5\leq j \text{ and } s=9 \\ C_2, & i+6\leq m \text{ and } n+5\leq j \text{ and } s=8 \\ \mathbb{0}, & \text{otherwise} \end{cases}$$

$$\begin{aligned} \text{where } m &= \textsc{First}(\eta) \\ n &= \textsc{Last}(\eta) \\ C_1 &\equiv \begin{cases} Q^{x,\text{futuremix}}_{i+1,j-1,s-2} \otimes W(\Delta G^{\text{interiorsize}}_{s,\text{mix}} - \Delta G^{\text{interiorsize}}_{s-2,\text{mix}}) & \mu_i=\mu_j=\mu_{i+1} \\ \mathbb{0} & \text{otherwise} \end{cases} \\ C_2 &\equiv \begin{cases} Q^b_{i+5,j+3-s} \otimes W(\Delta G^{\text{interiorsize}}_{s,\text{mix}} + \Delta G^{\text{interiorasymm}}_{s-8,\text{mix}} + \Delta G^{\text{interiormm}}_{i+4,i+5,j+3-s,j+4-s}(\phi)) & \textsc{IsSingle}(i,i+5,\mu_i) \text{ and } \\ & \textsc{IsSingle}(j+3-s,j,\mu_i) \\ \mathbb{0} & \text{otherwise} \end{cases} \\ C_3 &\equiv \begin{cases} Q^b_{s+i-3,j-5} \otimes W(\Delta G^{\text{interiorsize}}_{s,\text{mix}} + \Delta G^{\text{interiorasymm}}_{s-8,\text{mix}} + \Delta G^{\text{interiormm}}_{s+i-4,s+i-3,j-5,j-4}(\phi)) & \textsc{IsSingle}(i,s+i-3,\mu_i) \text{ and } \\ & \textsc{IsSingle}(j-5,j,\mu_i) \\ \mathbb{0} & \text{otherwise} \end{cases} \end{aligned} \tag{S151}$$

The recursion $R^{x,\text{mix}}_{\text{INTER}}(i,j,s,\phi)$ is closely related to (S144):

$$R^{x,\text{mix}}_{\text{INTER}}(i,j,s,\phi) \equiv \begin{cases} C_1 \oplus C_2 \oplus C_3, & i+6<m \text{ and } n+5<j \text{ and } 10 \leq s \leq j-i+m-n-3 \\ C_2 \oplus C_3, & i+6<m \text{ and } n+5<j \text{ and } s=9 \\ C_3, & i+6<m \text{ and } n+5=j \text{ and } s=9 \\ C_2, & i+6=m \text{ and } n+5\leq j \text{ and } s=9 \\ C_2, & i+6\leq m \text{ and } n+5\leq j \text{ and } s=8 \\ \mathbb{0}, & \text{otherwise} \end{cases}$$

$$\begin{aligned} \text{where } m &= \textsc{First}(\eta) \\ n &= \textsc{Last}(\eta) \\ C_1 &\equiv Q^{x,\text{mix}}_{i+1,j-1,s-2} \otimes W(\Delta G^{\text{interiorsize}}_{s,\text{mix}} - \Delta G^{\text{interiorsize}}_{s-2,\text{mix}}) \\ &\oplus \begin{cases} \mathbb{0} & \mu_i=\mu_j=\mu_{i+1} \\ Q^{x,\text{futuremix}}_{i+1,j-1,s-2} \otimes W(\Delta G^{\text{interiorsize}}_{s,\text{mix}} - \Delta G^{\text{interiorsize}}_{s-2,\text{mix}}) & \text{otherwise} \end{cases} \\ C_2 &\equiv \begin{cases} \mathbb{0} & \textsc{IsSingle}(i,i+5,\mu_i) \text{ and } \\ & \textsc{IsSingle}(j+3-s,j,\mu_i) \\ Q^b_{i+5,j+3-s} \otimes W(\Delta G^{\text{interiorsize}}_{s,\text{mix}} + \Delta G^{\text{interiorasymm}}_{s-8,\text{mix}} + \Delta G^{\text{interiormm}}_{i+4,i+5,j+3-s,j+4-s}(\phi)) & \text{otherwise} \end{cases} \\ C_3 &\equiv \begin{cases} \mathbb{0} & \textsc{IsSingle}(i,s+i-3,\mu_i) \text{ and } \\ & \textsc{IsSingle}(j-5,j,\mu_i) \\ Q^b_{s+i-3,j-5} \otimes W(\Delta G^{\text{interiorsize}}_{s,\text{mix}} + \Delta G^{\text{interiorasymm}}_{s-8,\text{mix}} + \Delta G^{\text{interiormm}}_{s+i-4,s+i-3,j-5,j-4}(\phi)) & \text{otherwise} \end{cases} \end{aligned} \tag{S152}$$

For $R^{x,\text{single}}_{\text{INTER}}(i,j,s,\phi)$, $R^{x,\text{futuremix}}_{\text{INTER}}(i,j,s,\phi)$, and $R^{x,\text{mix}}_{\text{INTER}}(i,j,s,\phi)$ in (S150)–(S152), the recursive component that extends previously encountered extensible loops is shown in $C_1$. Newly encountered extensible loops (elements that have exactly $L_1 = 4$ or $L_2 = 4$ or both) are handled by $C_2$ and $C_3$. Note that $C_2$ and $C_3$ are coincident for $L_1 = L_2 = 4$ ($s = 8$). The conditional checks using $m$ and $n$ prevent exterior loop states (that are not treated in this interior loop recursion) and disconnected states (that are not part of the complex ensemble).

***$O(N^3)$* interstrand interior loop recursion with fast conditional checks.** The recursion $R^{x,\text{single}}_{\text{INTER}}(i,j,s,\phi)$ is closely related to equation S145:

$$R^{x,\text{single}}_{\text{INTER}}(i,j,s,\phi) \equiv \begin{cases} C_0, & i+6 \le m \text{ and } n+5 \le j \\ \mathbb{0}, & \text{otherwise} \end{cases}$$

$$C_0 \equiv \begin{cases} I_{\text{single}}, & \text{IsSingle}(i,i+5,\mu_i) \text{ and IsSingle}(j-5,j,\mu_i) \\ \mathbb{0}, & \text{otherwise} \end{cases}$$

$$\text{where } I_{\text{single}} \equiv \begin{cases} C_1+C_2+C_3, & i+6 < m \text{ and } n+5 < j \text{ and } 10 \le s \le j-i+m-n-3 \\ C_2+C_3, & i+6 < m \text{ and } n+5 < j \text{ and } s=9 \\ C_3, & i+6 < m \text{ and } n+5 = j \text{ and } s=9 \\ C_2, & i+6 = m \text{ and } n+5 \le j \text{ and } s=9 \\ C_2, & s=8 \\ \mathbb{0}, & \text{otherwise} \end{cases} \quad \text{(S153)}$$

$$\begin{aligned} \text{with } m &= \text{First}(\eta) \\ n &= \text{Last}(\eta) \\ C_1 &\equiv Q^{x,\text{single}}_{i+1,j-1,s-2} \otimes W(\Delta G^{\text{interiorsize}}_{s,\mu_i} - \Delta G^{\text{interiorsize}}_{s-2,\mu_i}) \\ C_2 &\equiv Q^{b}_{i+5,j+3-s} \otimes W(\Delta G^{\text{interiorsize}}_{s,\mu_i} + \Delta G^{\text{interiorasymm}}_{s-8,\mu_i} + \Delta G^{\text{interiormm}}_{i+4,i+5,j+3-s,j+4-s}(\phi)) \otimes Q^{xa,\text{single}}_{j+3-s,j} \\ C_3 &\equiv Q^{b}_{s+i-3,j-5} \otimes W(\Delta G^{\text{interiorsize}}_{s,\mu_i} + \Delta G^{\text{interiorasymm}}_{s-8,\mu_i} + \Delta G^{\text{interiormm}}_{s+i-4,s+i-3,j-5,j-4}(\phi)) \otimes Q^{xa,\text{single}}_{i,s+i-3} \end{aligned}$$

The recursion $R^{x,\text{futuremix}}_{\text{INTER}}(i,j,s,\phi)$ is closely related to equation S146:

$$R^{x,\text{futuremix}}_{\text{INTER}}(i,j,s,\phi) \equiv \begin{cases} C_0, & i+6 \le m \text{ and } n+5 \le j \\ \mathbb{0}, & \text{otherwise} \end{cases}$$

$$C_0 \equiv \begin{cases} I_{\text{futuremix}}, & \text{IsSingle}(i,i+5,\mu_i) \text{ and IsSingle}(j-5,j,\mu_i) \\ \mathbb{0}, & \text{otherwise} \end{cases}$$

$$\text{where } I_{\text{futuremix}} \equiv \begin{cases} C_1+C_2+C_3, & i+6 < m \text{ and } n+5 < j \text{ and } 10 \le s \le j-i+m-n-3 \\ C_2+C_3, & i+6 < m \text{ and } n+5 < j \text{ and } s=9 \\ C_3, & i+6 < m \text{ and } n+5 = j \text{ and } s=9 \\ C_2, & i+6 = m \text{ and } n+5 \le j \text{ and } s=9 \\ C_2, & s=8 \\ \mathbb{0}, & \text{otherwise} \end{cases} \quad \text{(S154)}$$

$$\begin{aligned} \text{with } m &= \text{First}(\eta) \\ n &= \text{Last}(\eta) \\ C_1 &\equiv Q^{x,\text{single}}_{i+1,j-1,s-2} \otimes W(\Delta G^{\text{interiorsize}}_{s,\text{mix}} - \Delta G^{\text{interiorsize}}_{s-2,\text{mix}}) \\ C_2 &\equiv Q^{b}_{i+5,j+3-s} \otimes W(\Delta G^{\text{interiorsize}}_{s,\text{mix}} + \Delta G^{\text{interiorasymm}}_{s-8,\text{mix}} + \Delta G^{\text{interiormm}}_{i+4,i+5,j+3-s,j+4-s}(\phi)) \otimes Q^{xa,\text{single}}_{j+3-s,j} \\ C_3 &\equiv Q^{b}_{s+i-3,j-5} \otimes W(\Delta G^{\text{interiorsize}}_{s,\text{mix}} + \Delta G^{\text{interiorasymm}}_{s-8,\text{mix}} + \Delta G^{\text{interiormm}}_{s+i-4,s+i-3,j-5,j-4}(\phi)) \otimes Q^{xa,\text{single}}_{i,s+i-3} \end{aligned}$$

The recursion $R^{x,\text{mix}}_{\text{INTER}}(i,j,s,\phi)$ is closely related to equation S147:

$$R^{x,\text{mix}}_{\text{INTER}}(i,j,s,\phi) \equiv \begin{cases} C_0, & i+6 \le m \text{ and } n+5 \le j \\ \mathbb{0}, & \text{otherwise} \end{cases}$$

$$C_0 \equiv \begin{cases} I_{\text{single}}, & \text{IsSingle}(i,i+5,\mu_i) \text{ and IsSingle}(j-5,j,\mu_i) \\ I_{\text{mix}}, & \text{otherwise} \end{cases}$$

$$\text{where } I_{\text{single}} \equiv \begin{cases} C_{1s}+C_{2s}+C_{3s}, & i+6<m \text{ and } n+5<j \text{ and } 10 \le s \le j-i+m-n-3 \\ C_{2s}+C_{3s}, & i+6<m \text{ and } n+5<j \text{ and } s=9 \\ C_{3s}, & i+6<m \text{ and } n+5=j \text{ and } s=9 \\ C_{2s}, & i+6=m \text{ and } n+5\le j \text{ and } s=9 \\ C_{2s}, & s=8 \\ \mathbb{0}, & \text{otherwise} \end{cases}$$

$$\begin{aligned} \text{with } m &= \text{First}(\eta) \\ n &= \text{Last}(\eta) \\ C_{1s} &\equiv Q^{x,\text{mix}}_{i+1,j-1,s-2} \otimes W(\Delta G^{\text{interiorsize}}_{s,\text{mix}} - \Delta G^{\text{interiorsize}}_{s-2,\text{mix}}) \\ C_{2s} &\equiv Q^{b}_{i+5,j+3-s} \otimes W(\Delta G^{\text{interiorsize}}_{s,\text{mix}} + \Delta G^{\text{interiorasymm}}_{s-8,\text{mix}} + \Delta G^{\text{interiormm}}_{i+4,i+5,j+3-s,j+4-s}(\phi)) \otimes Q^{xa,\text{mix}}_{j+3-s,j} \\ C_{3s} &\equiv Q^{b}_{s+i-3,j-5} \otimes W(\Delta G^{\text{interiorsize}}_{s,\text{mix}} + \Delta G^{\text{interiorasymm}}_{s-8,\text{mix}} + \Delta G^{\text{interiormm}}_{s+i-4,s+i-3,j-5,j-4}(\phi)) \otimes Q^{xa,\text{mix}}_{i,s+i-3} \end{aligned}$$

$$\text{where } I_{\text{mix}} \equiv \begin{cases} C_{1m}+C_{2m}+C_{3m}, & i+6<m \text{ and } n+5<j \text{ and } 10 \le s \le j-i+m-n-3 \\ C_{2m}+C_{3m}, & i+6<m \text{ and } n+5<j \text{ and } s=9 \\ C_{3m}, & i+6<m \text{ and } n+5=j \text{ and } s=9 \\ C_{2m}, & i+6=m \text{ and } n+5\le j \text{ and } s=9 \\ C_{2m}, & s=8 \\ \mathbb{0}, & \text{otherwise} \end{cases}$$

$$\begin{aligned} \text{with } m &= \text{First}(\eta) \\ n &= \text{Last}(\eta) \\ C_{1m} &\equiv Q^{x,\text{mix}}_{i+1,j-1,s-2} \otimes W(\Delta G^{\text{interiorsize}}_{s,\text{mix}} - \Delta G^{\text{interiorsize}}_{s-2,\text{mix}}) \oplus Q^{x,\text{futuremix}}_{i+1,j-1,s-2} \otimes W(\Delta G^{\text{interiorsize}}_{s,\text{mix}} - \Delta G^{\text{interiorsize}}_{s-2,\text{mix}}) \\ C_{2m} &\equiv Q^{b}_{i+5,j+3-s} \otimes W(\Delta G^{\text{interiorsize}}_{s,\text{mix}} + \Delta G^{\text{interiorasymm}}_{s-8,\text{mix}} + \Delta G^{\text{interiormm}}_{i+4,i+5,j+3-s,j+4-s}(\phi)) \\ C_{3m} &\equiv Q^{b}_{s+i-3,j-5} \otimes W(\Delta G^{\text{interiorsize}}_{s,\text{mix}} + \Delta G^{\text{interiorasymm}}_{s-8,\text{mix}} + \Delta G^{\text{interiormm}}_{s+i-4,s+i-3,j-5,j-4}(\phi)) \end{aligned} \tag{S155}$$

***$R^{xa}_{\text{INTER}}$* recursion.** The $R^{x}_{\text{INTER}}(i,j,\phi)$ recursion references $Q^{xa}_{i,s+i-3}$ and $Q^{xa}_{j+3-s,j}$ elements that are always computed using the $R^{xa}_{\text{INTRA}}$ recursion of Figure S18 (if $i$ and $s+i-3$ or $j+3-s$ and $j$ are on the same strand) and never computed using the $R^{xa}_{\text{INTER}}$ recursion of Figure S19 (if $i$ and $s+i-3$ or $j+3-s$ and $j$ are on different strands). $R^{xa}_{\text{INTER}}(i,j,\phi)$ treats the conditional ensemble of subsequence $[i,j]$ in an interior loop context containing no terminal base pairs, where $i$ and $j$ are on different strands. Note that within the $R^{x}_{\text{INTER}}(i,j,\phi)$ recursion, $s+i-3$ must be on the first strand or $j+3-s$ must be on the last strand to ensure there are no nicks in the subsequence $[i,s+i-3]$ or $[j+3-s,j]$ (which would otherwise lead to a disconnected structure or an exterior loop state). Hence, $i$ and $s+i-3$ or $j+3-s$ and $j$ are always on the same strand and there are no calls to $R^{xa}_{\text{INTER}}$. Equivalently, the contributions for both single- and mixed-material $R^{xa}_{\text{INTER}}(i,j,\phi)$ recursions are $\mathbb{0}$ (Figure S19).

$$\begin{aligned} R^{xa,\text{single}}_{\text{INTER}}(i,j,\phi) &\equiv \mathbb{0} \\ R^{xa,\text{mix}}_{\text{INTER}}(i,j,\phi) &\equiv \mathbb{0} \end{aligned}$$

Figure S19: $R^{xa}_{\text{INTER}}$ recursion. No recursion diagram is provided because the recursion equations contain no contributions. This recursion is used for interior loops and does not change depending on whether coaxial and dangle stacking are included for multiloops and exterior loops.

#### S6.5 Recursions with coaxial and dangle stacking subensembles

Here, we describe $R^a_{\text{INTRA}}(i,j,\phi)$ recursions for calculating the elements of intrastrand blocks and $R^a_{\text{INTER}}(i,j,\phi)$ recursions for calculating the elements of interstrand blocks for the complex ensemble, $\overline{\Gamma}^{\shortparallel}$, including coaxial and dangle stacking subensembles in a mixed-materials setting. For the previously defined exterior loop and multiloop recursions without coaxial and dangle stacking (see Section S6.3), the elementary recursion entity was a *terminal base pair* (a base pair that terminates a duplex to form a part of the exterior loop or multiloop). For example, a recursion might contain exactly one terminal base pair, a $3'$-most terminal base pair, or one or more terminal base pairs. Here, for exterior loop and multiloop recursions with coaxial and dangle stacking, we make use of three new elementary recursion entities:

- *Coaxial stacking state:* two adjacent terminal base pairs that are coaxially stacked. Hence, a coaxial stacking state involves exactly two terminal base pairs.
- *Dangle stacking state:* zero, one, or two unpaired nucleotides dangle stacking on an adjacent terminal base pair. Hence, a dangle stacking state involves exactly one terminal base pair.
- *Stacking state:* a coaxial stacking state or a dangle stacking state (two adjacent terminal base pairs that are coaxially stacked or zero, one, or two unpaired nucleotides dangle stacking on an adjacent terminal base pair). Hence, a stacking state involves either two or one terminal base pairs.

For example, a recursion might contain exactly one stacking state, a $3'$-most stacking state, or one or more stacking states. Note that a terminal base pair without coaxial and dangle stacking corresponds to the subset of a dangle stacking state where there are zero nucleotides dangle stacking, so the complex ensemble without coaxial and dangle stacking is a subset of the complex ensemble with coaxial and dangle stacking.

To assist with examining the recursions with coaxial and dangle stacking, the intuition behind the name chosen for each recursion, the nature of the ensemble treated by each recursion, and the dependencies between the different recursions is summarized in Figure S20. To limit proliferation of new names and facilitate comparison to the non-stacking recursions of Section S6.3 (that treat complex ensemble $\overline{\Gamma}$ without coaxial and dangle stacking), we re-use the names of the non-stacking recursions but with updated recursion diagrams and recursion equations. Additionally, we introduce new recursions as needed to treat the coaxial and dangle stacking states in ensemble $\overline{\Gamma}^{\shortparallel}$.

A recursion $R^a(i,j,\phi)$ operates on subsequence $[i:j]$ to calculate element $i,j$ for either the unconstrained ensemble $a=\varnothing$ or for one of several constrained ensembles $a \in \{s, cd, b, n, x, xa, ma, ms, mcs, mc, md, m\}$. Multiloop and interior loop ensembles ($\{x, xa, ma, ms, mcs, mc, md, m\}$) are each divided into subensembles constrained by material $b \in \{single, \text{mix}\}$ so that loop contributions can be kept separate by material. This allows the correct free energy parameters to be applied for each material. Briefly, $R^{\varnothing}(i,j,\phi)$ treats the unconstrained ensemble in an exterior loop context where $i$ and $j$ may or may not be paired. $R^s(i,j,\phi)$ serves as an efficiency wrapper over the $3'$-most stacking state in an exterior loop context to reduce the time complexity from $O(N^4)$ to $O(N^3)$. $R^{cd}(i,j,\phi)$ treats a single stacking state (a coaxial stacking state or a dangle stacking state) in an exterior loop context. $R^b(i,j,\phi)$ treats the constrained ensemble where $i$ and $j$ form base pair $i\cdot j$ in the context of any loop type.

$R^{x,\text{single}}(i,j,\phi)$ and $R^{x,\text{mix}}(i,j,\phi)$ treat extensible interior loops (single- or mixed-material, respectively) to reduce the time complexity from $O(N^4)$ to $O(N^3)$. $R^{x,\text{futuremix}}(i,j,\phi)$ treats single-material extensible interior loops with the mixed parameter set to cover the scenario where a single-material extensible interior loop becomes a mixed-material extensible interior loop due to future extension of the loop. $R^{xa,\text{single}}(i,j,\phi)$ and $R^{xa,\text{mix}}(i,j,\phi)$ enable efficient determination of material composition in an interior loop context.

$R^{ma,\text{single}}(i,j,\phi)$ and $R^{ma,\text{mix}}(i,j,\phi)$ treat unpaired regions in a multiloop context (for single- and mixed-material unpaired regions, respectively). $R^{ms,\text{single}}(i,j,\phi)$ and $R^{ms,\text{mix}}(i,j,\phi)$ serves as efficiency wrappers over the $3'$-most stacking state in a multiloop context (analogous to $R^s$ in an exterior loop context) to reduce the time complexity from $O(N^4)$ to $O(N^3)$ (where the $3'$-most terminal base pair and any unpaired nucleotides are either a single material or mixed-material, respectively). $R^{mcs}(i,j,\phi)$ serves as an efficiency wrapper over the $3'$-most coaxial stacking state in a multiloop context to reduce the time complexity from $O(N^4)$ to $O(N^3)$ (where the $3'$-most terminal base pair and any unpaired nucleotides are either a single material or mixed-material, respectively). $R^{mc,\text{single}}(i,j,\phi)$ and $R^{mc,\text{mix}}(i,j,\phi)$ treat a single coaxial stacking state in a multiloop context (where both terminal base pairs are either a single material or mixed-material, respectively). $R^{md,\text{single}}(i,j,\phi)$ and $R^{md,\text{mix}}(i,j,\phi)$ treat a single dangle stacking state in a multiloop context (where the terminal base pair and any dangling nucleotides are a single material or mixed-material, respectively). $R^{m,\text{single}}(i,j,\phi)$ and $R^{m,\text{mix}}(i,j,\phi)$ treat one or more remaining stacking states in a multiloop context (where all remaining terminal base pairs and any unpaired nucleotides are a single material or mixed-material, respectively). When combined, $R^{mc}$ and $R^{md}$ constitute the multiloop equivalent to $R^{cd}$ in an exterior loop context; they are kept separate to allow proper treatment of a multiloop edge case. In the exterior loop context, the efficiency wrapper $R^s$ wraps $R^{cd}$ to treat coaxial and dangle stacking simultaneously. In the multiloop context, because of the edge case, the efficiency wrapper $R^{mcs}$ wraps $R^{mc}$ (to treat coaxial stacking alone) and then the efficiency wrapper $R^{ms}$

incorporates $R^{mcs}$ in addition to wrapping $R^{md}$ (to treat dangle stacking alone). Hence, the efficiency wrapper $R^{ms}$ (treating both coaxial and dangle stacking) is the multiloop equivalent to $R^{s}$ in an exterior loop context.

| Recursion | Naming intuition | Constraint | Context |
|---|---|---|---|
| $\varnothing$ | unconstrained | none | exterior loop |
| $s$ | **s**um | efficiency wrapper for $3'$-most stacking state | exterior loop |
| $cd$ | **c**oaxial and **d**angle | one stacking state (coaxial or dangle stacking state) | exterior loop |
| $b$ | **b**ase-paired | base pair between $5'$-most and $3'$-most bases of subsequence | any loop |
| $n$ | **n**ick | nick between strands | exterior loop |
| $x$, single | e**x**tensible, **single** material | extensible interior loop, single-material | interior loop |
| $x$, mix | e**x**tensible, **mix**ed material | extensible interior loop, mixed-material | interior loop |
| $x$, futuremix | e**x**tensible, **single** material | extensible interior loop, currently single material but potentially future mixed-material as loop is extended | interior loop |
| $xa$, single | e**x**tensible **a**ccessory, **single** material | extensible interior loop, single-material | interior loop |
| $xa$, mix | e**x**tensible **a**ccessory, **mix**ed material | extensible interior loop, mixed-material | interior loop |
| $ma$, single | **m**ultiloop **a**ccessory, **single** material | unpaired nucleotides, single-material | multiloop |
| $ma$, mix | **m**ultiloop **a**ccessory, **mix**ed material | unpaired nucleotides, mixed-material | multiloop |
| $ms$, single | **m**ultiloop **s**ummation, **single** material | efficiency wrapper for $3'$-most stacking state, multiloop is single-material | multiloop |
| $ms$, mix | **m**ultiloop **s**ummation, **mix**ed material | efficiency wrapper for $3'$-most stacking state multiloop is mixed-material | multiloop |
| $mcs$, single | **m**ultiloop **c**oaxial **s**um, **single** material | efficiency wrapper for $3'$-most coaxial stacking state, multiloop is single-material | multiloop |
| $mcs$, mix | **m**ultiloop **c**oaxial **s**um, **mix**ed material | efficiency wrapper for $3'$-most coaxial stacking state, multiloop is mixed-material | multiloop |
| $mc$, single | **m**ultiloop **c**oaxial, **single** material | one coaxial stacking state, multiloop is single-material | multiloop |
| $mc$, mix | **m**ultiloop **c**oaxial, **mix**ed material | one coaxial stacking state, multiloop is mixed-material | multiloop |
| $md$, single | **m**ultiloop **d**angle, **single** material | one dangle stacking state, multiloop is single-material | multiloop |
| $md$, mix | **m**ultiloop **d**angle, **mix**ed material | one dangle stacking state, multiloop is mixed-material | multiloop |
| $m$, single | **m**ultiloop, **single** material | one or more remaining stacking states, multiloop is single-material | multiloop |
| $m$, mix | **m**ultiloop, **mix**ed material | one or more remaining stacking states, multiloop is mixed-material | multiloop |

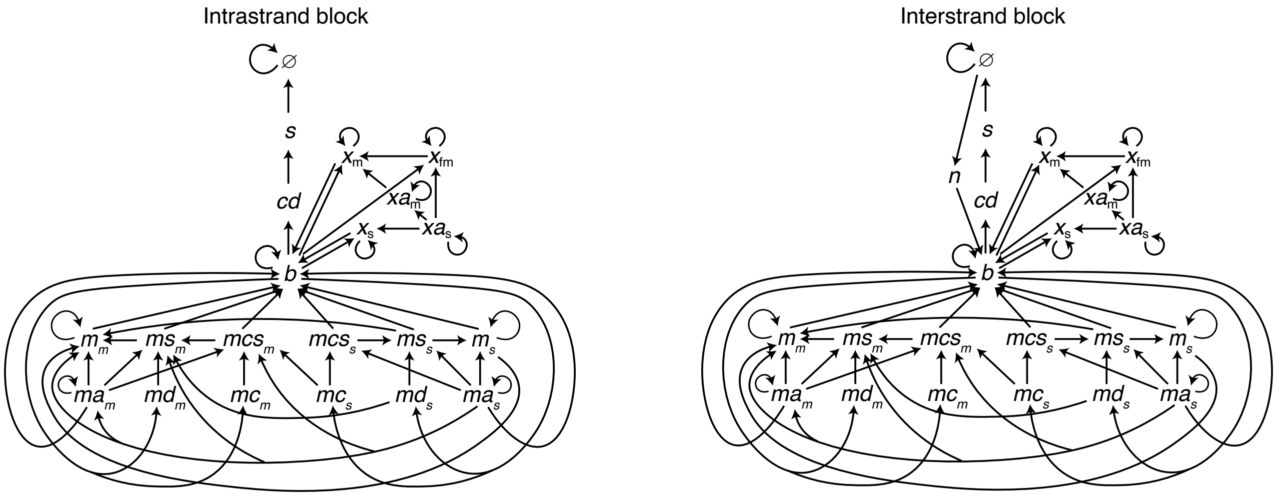

Figure S20: Nomenclature and connectivity for recursions with coaxial and dangle stacking. Top: Nomenclature. Bottom: Dependencies between different recursion types for elements within an intrastrand block (left) or an interstrand block (right). The "single", "mix", and "futuremix" material constraints are abbreviated as the subscripts $s$, $m$, and $fm$, respectively.

##### S6.5.1 Summation over dangle stacking states

Recursions that incorporate dangle stacking use a standardized approach to sum (using the $\oplus$ operator) over the subensemble of dangle stacking states on an adjacent terminal base pair. An example sum is depicted in the recursion diagram of Figure S21a by the dotted line between unpaired bases adjacent to a solid line denoting paired bases. The shading indicates explicit incorporation of a dangle free energy $\Delta G^{\text{dangle}}_{i,i+k,j-l,j}(\phi)$ (different for each dangle stacking state in the subensemble) and a terminal base pair free energy $\Delta G^{\text{terminalbp}}_{i+k,j-l}(\phi)$ (dependent on the sequence of base pair $i+k \cdot j-l$). The corresponding recursion equation of Figure S21b uses the indices $k \in \{0,1\}$ and $l \in \{0,1\}$ to sum over the four dangle stacking states, which are illustrated in Figure S21c. For $k = l = 0$, there are no unpaired bases dangle stacking on terminal base pair $i \cdot j$. For $k = 1,\ l = 0$, there is a $5'$ dangle stack on terminal base pair $i \cdot j$. For $k = 0,\ l = 1$, there is a $3'$ dangle stack on terminal base pair $i \cdot j$. For $k = l = 1$, there are both $5'$ and $3'$ dangle stacks on terminal base pair $i \cdot j$; this stacking state is referred to as a *terminal mismatch*. For clarity, recursion equations incorporate the generic dangle free energy function $\Delta G^{\text{dangle}}_{i,i+k,j-l,j}(\phi)$ which returns the appropriate free energy for each of the four stacking states:

$$\Delta G^{\text{dangle}}_{i,i+k,j-l,j}(\phi) = \begin{cases} 0 & k=0,\ l=0, \quad \text{no dangles} \\ \Delta G^{5'\text{dangle}}_{i,i+1,j}(\phi) & k=1,\ l=0, \quad 5' \text{ dangle} \\ \Delta G^{3'\text{dangle}}_{i\,j-1,j}(\phi) & k=0,\ l=1, \quad 3' \text{ dangle} \\ \Delta G^{\text{terminalmm}}_{i,i+1,j-1,j}(\phi) & k=1,\ l=1, \quad \text{terminal mismatch} \end{cases} \tag{S156}$$

Terminal mismatch free energies $\Delta G^{\text{terminalmm}}_{i,i+1,j-1,j}(\phi)$ have been published for RNA[4,7] and are included in the `rna95` and `rna06` parameter sets. However, terminal mismatch parameters for DNA are not public.[12] As a result, the `dna04` parameter set assigns the terminal mismatch free energy to be the sum of the published $5'$ and $3'$ dangle free energies:[11,12]

$$\Delta G^{\text{terminalmm}}_{i,i+1,j-1,j}(\phi) \equiv \Delta G^{5'\text{ dangle}}_{i,i+1,j}(\phi) + \Delta G^{3'\text{ dangle}}_{i\,j-1,j}(\phi). \tag{S157}$$

**a** Recursion diagram

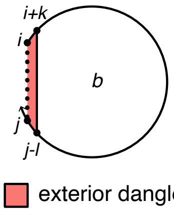

**b** Recursion equation

$$\bigoplus_{\substack{k\in\{0,1\}\\ l\in\{0,1\}}} \begin{cases} Q^b_{i+k,j-l} \otimes W(\Delta G^{\text{dangle}}_{i,i+k,j-l,j}(\phi) + \Delta G^{\text{terminalbp}}_{i+k,j-l}(\phi)), & i+k < j-l \\ 0, & \text{otherwise} \end{cases}$$

**c** Four possible dangle stacking states

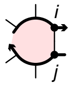

$k=0,\ l=0$ no dangles $\qquad Q^b_{i,j} \otimes W(\Delta G^{\text{terminalbp}}_{i,j}(\phi)), \qquad i<j$

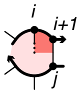

$k=1,\ l=0$ $5'$ dangle $\qquad Q^b_{i+1,j} \otimes W(\Delta G^{5'\text{dangle}}_{i,i+1,j}(\phi) + \Delta G^{\text{terminalbp}}_{i+1,j}(\phi)), \qquad i+1<j$

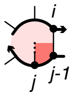

$k=0,\ l=1$ $3'$ dangle $\qquad Q^b_{i,j-1} \otimes W(\Delta G^{3'\text{dangle}}_{i,j-1,j}(\phi) + \Delta G^{\text{terminalbp}}_{i,j-1}(\phi)), \qquad i<j-1$

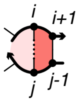

$k=1,\ l=1$ terminal mismatch $\qquad Q^b_{i+1,j-1} \otimes W(\Delta G^{\text{terminalmm}}_{i,i+1,j-1,j}(\phi) + \Delta G^{\text{terminalbp}}_{i+1,j-1}(\phi))\ , \qquad i+1<j-1$

Figure S21: (a) Example recursion diagram taken from the definition of $R^{cd}_{\text{INTRA}}(i,j,\phi)$. Note that the considered base pair is always between bases $i+k$ and $j-l$. (b) Equivalent recursion expression which specifies the specific free energy parameter contributions. (c) Decomposition of the sum in (b) into terms from each of 4 specific dangle stacking states.

##### S6.5.2 Intrastrand dynamic programming recursions with coaxial and dangle stacking

Here, we consider recursions for calculating the entries in a triangular intrastrand block with coaxial and dangle stacking. By definition, there are no nicks between strands in intrastrand recursions since intrastrand blocks involve base-pairing within a single strand.

**$R^{\varnothing}_{\text{INTRA}}$ recursion with coaxial and dangle stacking.** We begin with the recursion $R^{\varnothing}_{\text{INTRA}}(i,j,\phi)$ with the diagram and equation shown in Figure S22. The recursion $R^{\varnothing}_{\text{INTRA}}(i,j,\phi)$ does not change in the mixed-material setting, and is as previously described.[1] $R^{\varnothing}_{\text{INTRA}}(i,j,\phi)$ operates on the unconstrained ensemble for subsequence $[i,j]$ in an exterior loop context where $i$ and $j$ may or may not be paired (depicted with a dashed line between $i$ and $j$ in the recursion diagram). This recursion distinguishes two cases that are combined using $\oplus$ in the recursion equation:

- *No stacking states:* the empty case in an exterior loop context where there are no stacking states in subsequence $[i,j]$ (depicted by the absence of a straight solid line in the recursion diagram). The exterior loop shading in the recursion diagram represents the recursion energy $\Delta G^{\text{exterior}}_{i,j}(\phi) = 0$ corresponding to the zero reference state for an exterior loop with no coaxial stacking states or dangle stacking states (and hence, no terminal base pairs). The corresponding contribution to the recursion equation is $W(0) = \mathbb{1}$.
- *At least one stacking state:* the non-empty case in an exterior loop context where there is at least one stacking state (i.e., two adjacent terminal base pairs coaxially stacking, or zero, one, or two unpaired nucleotides dangle stacking on an adjacent terminal base pair) in subsequence $[i,j]$. The $3'$-most stacking state begins at $d+1$ and ends in the interval $[d+2,j]$ (depicted using a dashed line in the recursion diagram). The contributions for subsequence $[d+1,j]$ are incorporated using a $Q^s_{d+1,j}$ element. Contributions for the remaining subsequence $[i,d]$ are incorporated by a $Q^{\varnothing}_{i,d}$ element. The shading denotes the recursion energy 0 corresponding to the zero reference state in an exterior loop context. The edge case where the index $d+1=i$ is displayed explicitly to indicate that no $Q^{\varnothing}$ element is accessed in this case. The index limits in the recursion equation reflect the fact that steric effects prevent a hairpin loop with fewer than 3 unpaired nucleotides (hence, $i \cdot j$ cannot form if $j-i<4$).

Note that using the DOT notation (Algorithm S1) and index range notation (S131) to denote vector operations, we have the equivalence:

$$\text{DOT}\left(Q^{\varnothing}_{i,\overline{d}}, Q^{s}_{\overline{d}+1,j}\right) \equiv \sum_{d=i}^{j-5} Q^{\varnothing}_{i,d} \otimes Q^{s}_{d+1,j}, \quad j-i>4.$$

$$\text{where } \overline{d} \equiv [i : j-5].$$

We can also recognize that in terms of matrix elements, the dot product

$$\text{DOT}\left(Q^{\varnothing}_{i,\overline{d}}, Q^{s}_{\overline{d}+1,j}\right) \tag{S158}$$

is between the element range $\overline{d}$ of row $i$ (depicted as brown elements in Figure S4b) and the element range $\overline{d}$+1 of column $j$ (gray elements), yielding element $i,j$ (purple element).

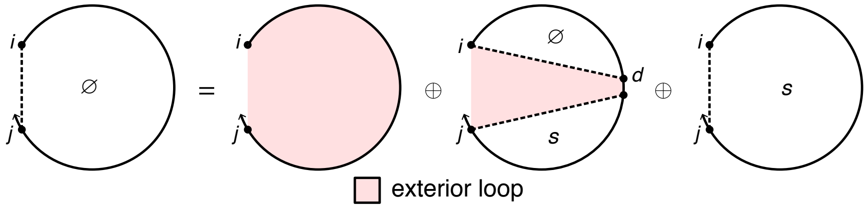

$$R^{\varnothing}_{\text{INTRA}}(i,j,\phi) \equiv \mathbb{1} \oplus \begin{cases} Q^{s}_{i,j} \oplus \text{DOT}\left(Q^{\varnothing}_{i,\overline{d}}, Q^{s}_{\overline{d}+1,j}\right), & j-i>4 \\ Q^{s}_{i,j}, & j-i=4 \\ \mathbb{0}, & \text{otherwise} \end{cases} \tag{S159}$$

$$\text{where } \overline{d} \equiv [i : j-5].$$

Figure S22: $R^{\varnothing}_{\text{INTRA}}$ recursion with coaxial and dangle stacking. Top: recursion diagram. Bottom: recursion equation.

***$R^s_{\text{INTRA}}$* recursion with coaxial and dangle stacking.** The $R^{\varnothing}_{\text{INTRA}}(i,j,\phi)$ recursion references $Q^s$ elements that are computed using the $R^s_{\text{INTRA}}$ recursion displayed in Figure S23. The $R^s_{\text{INTRA}}(i,j,\phi)$ recursion does not change in the mixed-material setting and is as previously described.[1] $R^s_{\text{INTRA}}(i,j,\phi)$ operates on a conditional ensemble for subsequence $[i,j]$ in an exterior loop context containing one stacking state starting at $i$ and ending in the interval $[i+1,j]$ (depicted as a dashed line between $i$ and $j$). The contribution for the stacking state in subsequence $[i,d]$ is incorporated using a $Q^{cd}_{i,d}$ element. Shading denotes no recursion energy as stacking energies and terminal base pair penalties are handled in $R^{cd}$. The index limits in the recursion equation reflect the steric requirement that there be at least 3 intervening bases between $i$ and $d$ (because the $R^{cd}$ recursion incorporates a minimum of one terminal base pair in subsequence $[i:d]$). Note that the $R^s$ recursion serves as an efficiency wrapper of the $R^{cd}$ recursion (here, representing the 3′-most stacking state in an exterior loop context) to reduce the time complexity of the $R^{\varnothing}$ recursion from $O(N^4)$ to $O(N^3)$. This time complexity reduction is achieved by defining the 3′-most stacking state using $R^{cd}$ within the $R^s$ efficiency wrapper rather than directly using the $R^{cd}$ recursion within the $R^{\varnothing}$ recursion, so as to avoid introducing a fourth independent index into the $R^{\varnothing}$ recursion.

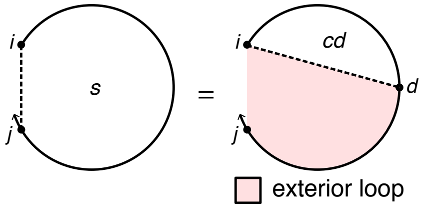

$$R^s_{\text{INTRA}}(i,j,\phi) \equiv \begin{cases} \text{DOT}\left(Q^{cd}_{i,\overline{d}}\right), & j-i \geq 4 \\ \mathbb{0}, & \text{otherwise} \end{cases}$$

where $\overline{d} \equiv [i+4:j]$.

Figure S23: $R^s_{\text{INTRA}}$ recursion with coaxial and dangle stacking. Top: recursion diagram. Bottom: recursion equation.

***$R^{cd}_{\text{INTRA}}$* recursion with coaxial and dangle stacking.** The $R^s_{\text{INTRA}}(i,j,\phi)$ recursion references $Q^{cd}$ elements that are computed using the $R^{cd}_{\text{INTRA}}$ recursion displayed in Figure S24. The $R^{cd}_{\text{INTRA}}(i,j,\phi)$ recursion does not change in the mixed-material setting and is as previously described.[1] $R^{cd}_{\text{INTRA}}(i,j,\phi)$ treats a single stacking state in an exterior loop context, corresponding to either of two cases that are combined using $\oplus$ in the recursion equation:

- *Coaxial stacking state:* two adjacent terminal base pairs ($i \cdot d$ and $d+1 \cdot j$) coaxially stack on each other. The contributions of subsequences $[i,d]$ and $[d+1,j]$ are incorporated using $Q^b_{i,d}$ and $Q^b_{d+1,j}$ elements. Shading corresponds to two kinds of recursion energy: 1) the sequence-dependent penalties for two terminal base pairs in an exterior loop context, $\Delta G^{\text{terminalbp}}_{i,d}(\phi)$ and $\Delta G^{\text{terminalbp}}_{d+1,j}(\phi)$ (dependent on the sequence of base pairs $i \cdot d$ and $d+1 \cdot j$), 2) the sequence-dependent coaxial stacking free energy $\Delta G^{\text{coax}}_{i,d,j}(\phi)$ (dependent on the sequences of base pairs $i \cdot d$ and $d+1 \cdot j$). Note that $\Delta G^{\text{coax}}_{i,d,j}(\phi)$ requires only 3 indices because $d+1$ is implied by $d$. The index limits in the recursion equation reflect the steric requirement that there be at least 3 intervening bases between $i$ and $d$ and at least 3 intervening bases between $d+1$ and $j$.

- *Dangle stacking state:* zero, one, or two unpaired nucleotides dangle stack on an adjacent terminal base pair ($i+k \cdot j-l$). The recursion diagram summarizes four dangle stacking states (depicted as a dotted line between $i$ and $j$) corresponding to no dangles, 5′ dangle, 3′ dangle, or terminal mismatch (see Figure S21 for details). The contribution of subsequence $[i+k, j-l]$ is incorporated using $Q^b_{i+k,j-l}$ element. Shading corresponds to two recursion energies: 1) the sequence-dependent penalty for a terminal base pair in an exterior loop context, $\Delta G^{\text{terminalbp}}_{i+k,j-l}(\phi)$ (dependent on the sequence of base pair $i+k \cdot j-l$), 2) the sequence-dependent dangle stacking free energy $\Delta G^{\text{dangle}}_{i,i+k,j-l,j}(\phi)$ which takes on one of four values corresponding to the four dangle stacking states (see Figure S21). The index limits in the recursion equation reflect the steric requirement that there be at least 3 intervening bases between $i+k$ and $j-l$.

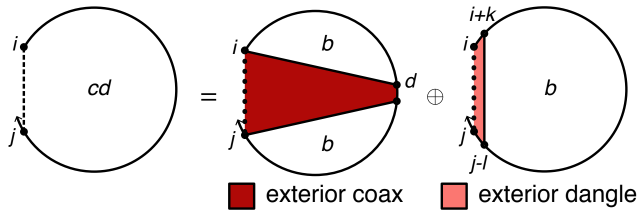

$$R^{cd}_{\text{INTRA}}(i,j,\phi) \equiv \begin{cases} \text{DOT}\left(Q^b_{i,\overline{d}}, Q^b_{\overline{d}+1,j}, W(\Delta G^{\text{coax}}_{i,\overline{d},j}(\phi) + \Delta G^{\text{terminalbp}}_{i,\overline{d}}(\phi) + \Delta G^{\text{terminalbp}}_{\overline{d}+1,j}(\phi))\right), & j-i \geq 9 \\ \mathbb{0}, & \text{otherwise} \end{cases}$$

$$\oplus \bigoplus_{\substack{k\in\{0,1\}\\ l\in\{0,1\}}} \begin{cases} Q^b_{i+k,j-l} \otimes W(\Delta G^{\text{dangle}}_{i,i+k,j-l,j}(\phi) + \Delta G^{\text{terminalbp}}_{i+k,j-l}(\phi)), & (j-l)-(i+k) \geq 4 \\ \mathbb{0}, & \text{otherwise} \end{cases}$$

where $\overline{d} \equiv [i+4 : j-5]$

Figure S24: $R^{cd}_{\text{INTRA}}$ recursion with coaxial and dangle stacking. Top: recursion diagram. Bottom: recursion equation.

**$R^b_{\text{INTRA}}$ recursion with coaxial and dangle stacking.** The $R^{cd}_{\text{INTRA}}(i,j,\phi)$ recursion references $Q^b$ elements that are computed using the $R^b_{\text{INTRA}}$ recursion displayed in Figure S25. $R^b_{\text{INTRA}}(i,j,\phi)$ operates on a conditional ensemble for subsequence $[i,j]$ with $i$ and $j$ base paired to each other (depicted with a solid line between $i$ and $j$). In the mixed-material setting, the appropriate single- or mixed-material sub-recursions and parameters must be applied based on the material composition of the loops. The recursion distinguishes four cases that are combined using $\oplus$ in the recursion equation:

- *Hairpin loop:* the hairpin loop closed by the single base pair $i \cdot j$ (depicted by a straight solid line). The recursion incorporates the recursion energy $\Delta G^{\text{hairpin}}_{i,j}(\phi)$, which takes into account the material of each nucleotide in the subsequence $[i,j]$ (Section S4.5). This treatment of hairpin loops is the same as for the non-stacking recursions. The index limits in the recursion equation reflect the fact that steric constraints prevent a hairpin loop with fewer than 3 unpaired nucleotides (hence, $i \cdot j$ cannot form if $j-i<4$).

- *Interior loop:* the interior loop closed by the two terminal base pairs $i \cdot j$ and $d \cdot e$ (depicted by straight solid lines). Calculation of the interior loop contributions using an $O(N^4)$ or $O(N^3)$ version of the InteriorIntra recursion is described in Section S6.4. This treatment of interior loops is the same as for the non-stacking recursions. The index limits in the recursion equation reflect the fact that steric effects prevent an interior loop with $j-i<6$ due to the steric requirement that there be at least 3 intervening bases between $d$ and $e$.

- *Multiloop with coaxial stacking on terminal base pair $j \cdot i$:* the multiloop closed by three or more terminal base pairs with coaxial stacking on base pair $j \cdot i$. This case corresponds to the eight recursion diagrams on the second and third rows of Figure S25 and is treated by the subroutine MultiCoaxIntra (recursion equation S160). The recursions in the second row treat the case where the terminal base pair $j \cdot i$ forms a coaxial stack with adjacent terminal base pair $d+1 \cdot j-1$, depicted as a dotted straight line between $i$ and $d+1$. The contribution of subsequence $[i+1,d]$ is incorporated by element $Q^b_{i+1,d}$. The contributions of one or more remaining stacking states in subsequence $[i+1,d]$ are incorporated by element $Q^m_{i+1,d}$. The recursions in the third row treat the analogous case where the terminal base pair $j \cdot i$ forms a coaxial stack with adjacent terminal base pair $i+1 \cdot d$; for simplicity, here we focus on describing the recursions in the second row:
    - *Single- and mixed-material recursions.* Multiloop contributions must be kept separate by material in $R^b_{\text{INTRA}}(i,j,\phi)$, so that the appropriate $\Delta G^{\text{multi}}_{\text{init}}$, $\Delta G^{\text{multi}}_{\text{bp}}$, and $\Delta G^{\text{terminalbp}}_{j,i}(\phi)$ parameters are applied depending on the material of the multiloop. Contributions for single-material and mixed-material multiloops are indicated in the recursion equation as $M_{\text{single}}$ and $M_{\text{mix}}$, respectively. To use the correct physical model for each multiloop as recursion elements are combined across multiple recursion levels, lower level recursions must be calculated separately for the "single" and "mixed" scenarios before they are combined at the $R^b_{\text{INTRA}}(i,j,\phi)$ level. $Q^m_{i+1,d}$ is split into $Q^{m,\text{single}}_{i+1,d}$ and $Q^{m,\text{mix}}_{i+1,d}$. "Single" elements (depicted with a blue backbone in recursion diagrams) only track energy contributions from single-material loop regions, while "mix" elements (depicted with a red backbone in recursion diagrams) only track

contributions from mixed-material loop regions. The contribution for the subsequence $[d+1, j-1]$ enclosed by base pair $d+1 \cdot j-1$ is incorporated using a $Q^b_{d+1,j-1}$ element, representing either a single-material or mixed-material terminal base pair depending on whether $\mu_{j-1} = \mu_{d+1}$ (single-material; denoted by a blue base pair in the recursion diagram) or $\mu_{j-1} \neq \mu_{d+1}$ (mixed-material; denoted by a red base pair in the recursion diagram). Because $Q^m_{i+1,d}$ is split into single-material and mixed-material recursions and $Q^b_{d+1,j-1}$ also has both single- and mixed-material cases, there are four possible single/mix combinations when multiplying them. The three combinations that contain at least one mixed element must represent mixed contributions and are included in $M_{\text{mix}}$ with the term $C_{1m}$. The remaining combination in term $C_{1s}$, produces a single-material multiloop only if the material of both elements match each other and the material of the closing base pair $j \cdot i$, which is checked using the conditional, $\mu_j = \mu_{j-1} = \mu_{i+1} = \mu_i$. If the conditional is satisfied, the contribution is added to $M_{\text{single}}$ using term $C_{0s}$. Otherwise, if the conditional fails, the contribution is added to $M_{\text{mix}}$ using term $C_{0m}$. In the conditional, note that the material of the multiloop "single" element, $Q^{m,\text{single}}_{i+1,d}$, can be determined by checking the material of the first or last nucleotide of the subsequence spanned by that element; the first and last nucleotides are each guaranteed to appear in the multiloop as either an unpaired nucleotide or as part of a terminal base pair defining the multiloop. In this instance, the material of $Q^{m,\text{single}}_{i+1,\overline{d}}$ is checked at $i+1$ since this index is constant over vector $\overline{d}$ and apply to all $d$ in $\overline{d}$. $M_{\text{single}}$ and $M_{\text{mix}}$ are combined to incorporate all single- and mixed-material multiloops that include coaxial stacking on terminal base pair $j \cdot i$. The index limits in the recursion equation reflect the fact that steric effects prevent a multiloop with $j - i < 11$ due to the steric requirement that there be at least 3 intervening bases between $i+1$ and $d$ (which must contain one or more stacking states and hence one or more terminal base pairs) and at least 3 intervening bases between $d+1$ and $j-1$.

- *Single- and mixed-material recursion energies.* In the recursion diagrams, pale green shading corresponds to three recursion energies: 1) the penalty for formation of a multiloop $\Delta G^{\text{multi}}_{\text{init}}$, 2) the sequence-independent penalties for two terminal base pairs in a multiloop, $\Delta G^{\text{multi}}_{\text{bp}}$ (corresponding to base pairs $d+1 \cdot j-1$ and $j \cdot i$), 3) the sequence-dependent penalties for two terminal base pairs in a multiloop context, $\Delta G^{\text{terminalbp}}_{d+1,j-1}(\phi)$ and $\Delta G^{\text{terminalbp}}_{j,i}(\phi)$ (note that the indices are ordered $j$ then $i$ to reflect $5'$ to $3'$ from the perspective of the multiloop). For $M_{\text{single}}$ (single-material multiloops checked using the conditional $\mu_j = \mu_{j-1} = \mu_{i+1} = \mu_i$), the single-material multiloop initiation parameter, $\Delta G^{\text{multi}}_{\text{init},\mu_i}$, is used and the single-material terminal base pair parameter, $\Delta G^{\text{multi}}_{\text{bp},\mu_i}$, is used for each of terminal base pairs $i \cdot j$ and $d+1 \cdot j-1$. For $M_{\text{mix}}$, the mixed-material multiloop initiation parameter $\Delta G^{\text{multi}}_{\text{init,mix}}$ is used; for terminal base pair $j \cdot i$, the terminal base pair parameter, $\Delta G^{\text{multi}}_{\text{bp},\sigma}$, depends on the material of $j$ and $i$ and is checked with the $\sigma$ conditional ($\Delta G^{\text{multi}}_{\text{bp},\mu_i}$ is used if $\mu_j = \mu_i$, and $\Delta G^{\text{multi}}_{\text{bp,mix}}$ is used otherwise); for terminal base pair $d+1 \cdot j-1$, the terminal base pair parameter depends on the material of $d+1$ and $j-1$ ($\Delta G^{\text{multi}}_{\text{bp},\mu_{j-1}}$ is used if $\mu_{d+1} = \mu_{j-1}$, and $\Delta G^{\text{multi}}_{\text{bp,mix}}$ is used otherwise). For $M_{\text{single}}$ and $M_{\text{mix}}$, the sequence-dependent terminal base pair parameter, $\Delta G^{\text{terminalbp}}_{j,i}(\phi)$, is determined by lookup based on the sequence and material of nucleotides $j$ and $i$ (note that the indices are ordered $j$ then $i$ to reflect $5'$ to $3'$ from the perspective of the multiloop); the sequence-dependent terminal base pair parameter, $\Delta G^{\text{terminalbp}}_{d+1,j-1}(\phi)$, is determined by lookup based on the sequence and material of nucleotides $d+1$ and $j-1$. In the recursion diagrams, dark green shading corresponds to the sequence-dependent coaxial stacking recursion energy $\Delta G^{\text{coax}}_{d+1,j-1,i}(\phi)$ (dependent on the sequences of terminal base pairs $d+1 \cdot j-1$ and $j \cdot i$). Note that $\Delta G^{\text{coax}}_{d+1,j-1,i}(\phi)$ requires only 3 indices because $j$ is implied by $j-1$. $\Delta G^{\text{coax}}_{\overline{d}+1,j-1,i}(\phi)$ is determined by lookup based on the sequence and material of nucleotides $d+1$, $j-1$, $j$ and $i$.

- *Multiloop with dangle stacking on terminal base pair $j \cdot i$:* the multiloop closed by three or more terminal base pairs with dangle stacking on terminal base pair $j \cdot i$. This case corresponds to the ten recursion diagrams on the fourth, fifth, and sixth rows of Figure S25 and is treated by the subroutine MULTIDANGLEINTRA (recursion equation S161). Multiloop contributions must be kept separate by material in $R^b_{\text{INTRA}}(i, j, \phi)$, so that the appropriate $\Delta G^{\text{multi}}_{\text{init}}$, $\Delta G^{\text{multi}}_{\text{bp}}$, and $\Delta G^{\text{terminalbp}}_{j,i}(\phi)$ parameters are applied depending on the material of the multiloop. Contributions for single-material and mixed-material multiloops are indicated in the recursion equation as $M_{\text{single}}$ and $M_{\text{mix}}$, respectively. To use the correct physical model for each multiloop as recursion elements are combined across multiple recursion levels, lower level recursions ($Q^m$, $Q^{ms}$, $Q^{ma}$, $Q^{mcs}$) must be calculated separately for the "single" and "mixed" scenarios before they are combined at the $R^b_{\text{INTRA}}(i, j, \phi)$ level. "Single" elements (depicted with a blue backbone in recursion diagrams) only track energy contributions from single-material loop regions, while "mix" elements (depicted with a red backbone in recursion diagrams) only track contributions from mixed-material loop regions.

  - *Base case with two or more additional stacking states.* The recursions on the fourth row treat the case where there is a dangle stacking state involving the terminal base pair $j \cdot i$ (depicted as a dotted straight line between $i+k$ and $j-l$), a $3'$-most stacking state in subsequence $[d+1, j-l-1]$ (depicted as a dashed line between $d+1$ and $j-l-1$), and one or more stacking states in subsequence $[i+k+1, d]$ (depicted as a dashed line between $i+k+1$

and $d$). The contributions for subsequence $[d+1, j-l-1]$ are incorporated using a $Q^{ms}_{d+1,j-l-1}$ element, while the contributions for $[i+k+1, d]$ are incorporated using a $Q^{m}_{i+k+1,d}$ element. Because $Q^{m}_{i+k+1,d}$ and $Q^{ms}_{d+1,j-l-1}$ are each split into single- and mixed-material recursions, there are 4 possible combinations when multiplying them. Any combination with a mixed element produces an overall mixed multiloop ($Q^{m,\text{mix}}_{i+k+1,d} \otimes Q^{ms,\text{mix}}_{d+1,j-l-1}, Q^{m,\text{mix}}_{i+k+1,d} \otimes Q^{ms,\text{single}}_{d+1,j-l-1}, Q^{m,\text{single}}_{i+k+1,d} \otimes Q^{ms,\text{mix}}_{d+1,j-l-1}$), which is added to $M_{\text{mix}}$ as term $C_{1m}$ in (S161). The remaining combination, $Q^{m,\text{single}}_{i+k+1,d} \otimes Q^{ms,\text{single}}_{d+1,j-l-1}$, in term $C_{1s}$ produces a single-material multiloop only if the material matches for both elements, the closing base pair $j \cdot i$, and any unpaired dangling nucleotide $i+k$ and $j-l$, which is checked using the conditional $\text{IsSingle}(i, i+k+1, \mu_i)$ and $\text{IsSingle}(j-l-1, j, \mu_i)$. Recall that the function $\text{IsSingle}(x, y, \mu_i)$ checks if subsequence $[x, y]$ is composed entirely of material $\mu_i$. If the conditional is satisfied, the contribution is added to $M_{\text{single}}$ using term $C_{0s}$. Otherwise, if the conditional fails, the contribution is added to $M_{\text{mix}}$ using term $C_{0m}$. In the conditional, note that the material of any multiloop single element, like $Q^{m,\text{single}}_{i+k+1,d}$ or $Q^{ms,\text{single}}_{d+1,j-l-1}$, can be known by checking the material of the first or last nucleotide of the subsequence spanned by that element; the first and last nucleotides are each guaranteed to appear in the multiloop as either an unpaired nucleotide or as part of a terminal base pair defining the multiloop. In this instance, the material of $Q^{m,\text{single}}_{i+k+1,\overline{d}}$ is checked at $i+k+1$, while the material of $Q^{ms,\text{single}}_{\overline{d}+1,j-l-1}$ is checked at $j-l-1$, since these indices are constant over vector $\overline{d}$ and apply to all $d$ and $d+1$ in $\overline{d}$. Note that $k, l \in \{0, 1\}$ determine whether unpaired nucleotides dangle stack on the adjacent terminal base pair $j \cdot i$ in a multiloop context. The situation is analogous to that in an exterior loop context with $R^{cd}_{\text{INTRA}}(i, j, \phi)$ (as detailed in Figure S21) with the only difference being that in the exterior loop context, $i+k$ and $j-l$ index the paired bases and in the multiloop context $i+k$ and $j-l$ index the unpaired bases. The index limits in the recursion equation reflect the fact that steric effects prevent a multiloop with $(j-l)-(i+k) < 11$ due to the steric requirement that there be at least 3 intervening bases between $i+k+1$ and $d$ (which must contain one or more stacking states and hence one or more terminal base pairs) and at least 3 intervening bases between $d+1$ and $j-l-1$ (which must contain a 3′-most stacking state and hence one or two terminal base pairs). *Single- and mixed-material recursion energies.* In the recursion diagrams, the pale green shading corresponds to four multiloop recursion energies: 1) the penalty for formation of a multiloop $\Delta G^{\text{multi}}_{\text{init}}$, 2) the sequence-independent penalty for one terminal base pair in a multiloop, $\Delta G^{\text{multi}}_{\text{bp}}$ (corresponding to base pair $j \cdot i$), 3) the penalty per unpaired nucleotide in a multiloop $\Delta G^{\text{multi}}_{\text{nt}}$ (a total of $k+l$ dangling nucleotides; as a result this term is zeroed out when $k = l = 0$), 4) the sequence-dependent penalty for a terminal base pair in a multiloop context, $\Delta G^{\text{terminalbp}}_{j,i}(\phi)$ (note that the indices are ordered $j$ then $i$ to reflect 5′ to 3′ from the perspective of the multiloop). For $M_{\text{single}}$, the single-material multiloop initiation parameter $\Delta G^{\text{multi}}_{\text{init},\mu_i}$ is used; the single-material terminal base pair parameter $\Delta G^{\text{multi}}_{\text{bp},\mu_i}$ is used for terminal base pair $j \cdot i$; the unpaired nucleotide parameter $\Delta G^{\text{multi}}_{\text{nt},\mu_i}$ is used for any unpaired nucleotide $i+k$ or $j-l$. For $M_{\text{mix}}$, the mixed-material multiloop initiation parameter $\Delta G^{\text{multi}}_{\text{init,mix}}$ is used; for terminal base pair $j \cdot i$, the terminal base pair parameter, $\Delta G^{\text{multi}}_{\text{bp},\sigma}$, depends on the material of $j$ and $i$ and is checked with the $\sigma$ conditional ($\Delta G^{\text{multi}}_{\text{bp},\mu_i}$ is used if $\mu_j = \mu_i$, and $\Delta G^{\text{multi}}_{\text{bp,mix}}$ is used otherwise); the unpaired nucleotide parameter $\Delta G^{\text{multi}}_{\text{nt}}$ depends on the material of each dangling nucleotide ($\Delta G^{\text{multi}}_{\text{nt},\mu_{i+k}}$ for $i+k$ and $\Delta G^{\text{multi}}_{\text{nt},\mu_{j-l}}$ for $j-l$). For $M_{\text{single}}$ and $M_{\text{mix}}$, the sequence-dependent terminal base pair parameter, $\Delta G^{\text{terminalbp}}_{j,i}(\phi)$, is determined by lookup based on the sequence and material of nucleotides $j$ and $i$ (note that the indices are ordered $j$ then $i$ to reflect 5′ to 3′ from the perspective of the multiloop). In the recursion diagrams, the medium green shading corresponds to the sequence-dependent dangle stacking recursion energy $\Delta G^{\text{dangle}}_{j-l,j,i,i+k}(\phi)$ (dependent on the sequence and material of the participating nucleotides).

- *Edge case with one additional coaxial stacking state.*The recursions on the fifth row treat the case where there is a dangle stacking state involving the terminal base pair $j \cdot i$ (depicted as a dotted straight line between $i+k$ and $j-l$), a single coaxial stacking state in subsequence $[e, j-l-1]$ (depicted as a half-solid/half-dashed line between $e$ and $j-l-1$), and an unpaired region in subsequence $[i+k+1, e-1]$ (depicted as a hollow dotted line between $i+k+1$ and $e-1$). The contributions for subsequence $[e, j-l-1]$ are incorporated using a $Q^{mcs}_{e,j-l-1}$ element, while the contributions for unpaired region $[i+k+1, e-1]$ are incorporated using a $Q^{ma}_{i+k+1,e-1}$ element. Because $Q^{ma}_{i+k+1,e-1}$ and $Q^{mcs}_{e,j-l-1}$ are each split into single- and mixed-material recursions, there are 4 possible combinations when multiplying them. Any combination with a mixed element produces an overall mixed multiloop ($Q^{ma,\text{mix}}_{i+k+1,e-1} \otimes Q^{mcs,\text{mix}}_{e,j-l-1}, Q^{ma,\text{mix}}_{i+k+1,e-1} \otimes Q^{mcs,\text{single}}_{e,j-l-1}, Q^{ma,\text{single}}_{i+k+1,e-1} \otimes Q^{mcs,\text{mix}}_{e,j-l-1}$), which is added to $M_{\text{mix}}$ as term $C_{2m}$ in (S161). The remaining combination, $Q^{ma,\text{single}}_{i+k+1,e-1} \otimes Q^{mcs,\text{single}}_{e,j-l-1}$, in term $C_{2s}$ produces a single-material multiloop only if the material matches for both elements, the closing base pair $j \cdot i$, and any unpaired dangling nucleotide $i+k$ and $j-l$, which is checked with the conditional $\text{IsSingle}(i, i+k+1, \mu_i)$ and $\text{IsSingle}(j-l-1, j, \mu_i)$. Recall that the function $\text{IsSingle}(x, y, \mu_i)$ checks if subsequence $[x, y]$ is composed entirely of material $\mu_i$. If the conditional is satisfied, the contribution is added to $M_{\text{single}}$ using term $C_{02}$. Otherwise, if the conditional fails, the contribution is added to $M_{\text{mix}}$ using term $C_{0m}$. In the conditional, note that the material of $Q^{ma,\text{single}}_{i+k+1,\overline{e}-1}$ is checked at $i+k+1$, while the material of $Q^{mcs,\text{single}}_{\overline{e},j-l-1}$ is checked at $j-l-1$, since these indices are constant over vector $\overline{e}$ and apply to all $e-1$ and $e$ in $\overline{e}$. The recursions on the sixth row treat the special case where $e=i+k+1$, in which the contributions for subsequence $[i+k+1, j-l-1]$ are incorporated using a $Q^{mcs}_{i+k+1,j-l-1}$ element, for which there are both single- and mixed-material recursions. The mixed-material contribution $Q^{mcs,\text{mix}}_{i+k+1,j-l-1}$ is added to $M_{\text{mix}}$ as term $C_{3m}$. The contribution of $Q^{mcs,\text{single}}_{i+k+1,j-l-1}$ in term $C_{3s}$ produces a single-material multiloop only if the material matches for the $Q^{mcs,\text{single}}_{i+k+1,j-l-1}$ element, the closing base pair $j \cdot i$, and any unpaired dangling nucleotide $i+k$ and $j-l$, which is checked with the conditional $\textsc{IsSingle}(i, i+k+1, \mu_i)$ and $\textsc{IsSingle}(j-l-1, j, \mu_i)$. If the conditional is satisfied, the contribution is added to $M_{\text{single}}$ using term $C_{0s}$. Otherwise, if the conditional fails, the contribution is added to $M_{\text{mix}}$ using term $C_{0m}$. The index limits in the recursion equation reflect the fact that steric effects prevent a multiloop with $(j-l)-(i+k) < 11$ due to the steric requirement that there be at least 8 intervening bases between $e$ and $j-l-1$ (which must contain a coaxial stacking state and hence two adjacent terminal base pairs). Note that this edge case covers the scenario where there are exactly three terminal base pairs and the two terminal base pairs that are not $j \cdot i$ are coaxially stacked. That situation is not covered by the base case because for that recursion, a multiloop with 3 terminal base pairs would have one terminal base pair in the $Q^{ms}$ element, and one terminal base pair in the $Q^{m}$ element (hence, those two terminal base pairs cannot coaxially stack since they are in different recursions). *Single- and mixed-material recursion energies.* In the recursion diagrams, the pale green shading corresponds to four multiloop recursion energies: 1) the penalty for formation of a multiloop $\Delta G^{\text{multi}}_{\text{init}}$, 2) the sequence-independent penalty for one terminal base pair in a multiloop, $\Delta G^{\text{multi}}_{\text{bp}}$ (corresponding to base pair $j \cdot i$), 3) the penalty per unpaired nucleotide in a multiloop $\Delta G^{\text{multi}}_{\text{nt}}$ ($k+l$ dangling nucleotides plus the unpaired nucleotides $k+l+1, \ldots, e-1$; as a result this term is zeroed out when $k=l=0$ and $e=i+1$), 4) the sequence-dependent penalty for a terminal base pair in a multiloop context, $\Delta G^{\text{terminalbp}}_{j,i}(\phi)$ (note that the indices are ordered $j$ then $i$ to reflect $5'$ to $3'$ from the perspective of the multiloop). For $M_{\text{single}}$, the single-material multiloop initiation parameter $\Delta G^{\text{multi}}_{\text{init},\mu_i}$ is used; the single-material terminal base pair parameter $\Delta G^{\text{multi}}_{\text{bp},\mu_i}$ is used for terminal base pair $j \cdot i$; the unpaired nucleotide parameter $\Delta G^{\text{multi}}_{\text{nt},\mu_i}$ is used for any unpaired nucleotide $i+k$ or $j-l$. For $M_{\text{mix}}$, the mixed-material multiloop initiation parameter $\Delta G^{\text{multi}}_{\text{init,mix}}$ is used; for terminal base pair $j \cdot i$, the terminal base pair parameter, $\Delta G^{\text{multi}}_{\text{bp},\sigma}$, depends on the material of $j$ and $i$ and is checked with the $\sigma$ conditional ($\Delta G^{\text{multi}}_{\text{bp},\mu_i}$ is used if $\mu_j = \mu_i$, and $\Delta G^{\text{multi}}_{\text{bp,mix}}$ is used otherwise); the unpaired nucleotide parameter $\Delta G^{\text{multi}}_{\text{nt}}$ depends on the material of each dangling nucleotide ($\Delta G^{\text{multi}}_{\text{nt},\mu_{i+k}}$ for $i+k$ and $\Delta G^{\text{multi}}_{\text{nt},\mu_{j-l}}$ for $j-l$). For $M_{\text{single}}$ and $M_{\text{mix}}$, the sequence-dependent terminal base pair parameter, $\Delta G^{\text{terminalbp}}_{j,i}(\phi)$, is determined by lookup based on the sequence and material of nucleotides $j$ and $i$ (note that the indices are ordered $j$ then $i$ to reflect $5'$ to $3'$ from the perspective of the multiloop). In the recursion diagrams, the medium green shading corresponds to the sequence-dependent dangle stacking recursion energy $\Delta G^{\text{dangle}}_{j-l,j,i,i+k}(\phi)$ (dependent on the sequence and material of the participating nucleotides).

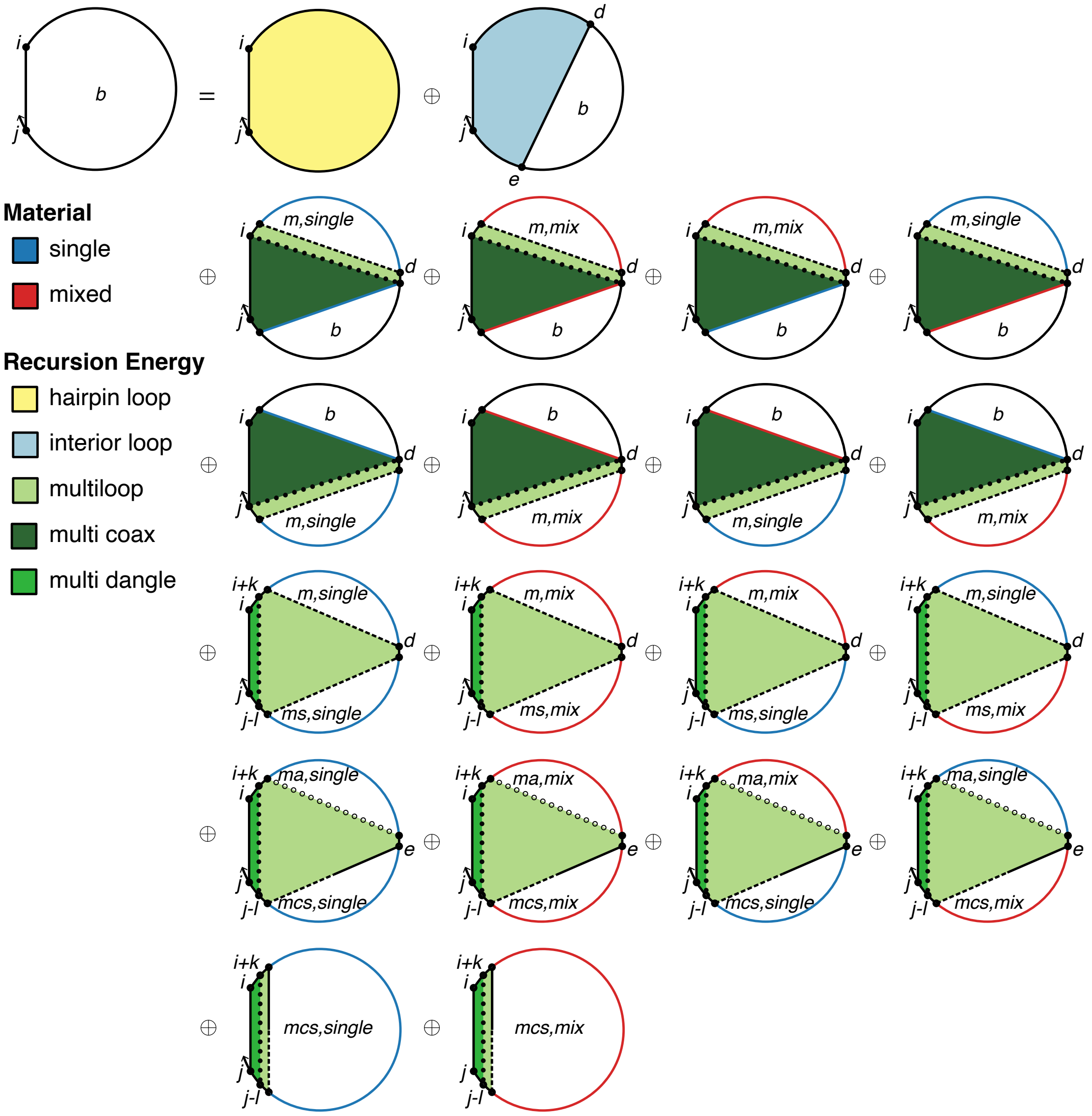

$$R^b_{\text{INTRA}}(i,j,\phi) \equiv \begin{cases} C_{\text{all}}, & \textsc{Complementary}(\phi_i,\phi_j) \\ \mathbb{0}, & \text{otherwise} \end{cases}$$

$$\text{where } C_{\text{all}} \equiv \begin{cases} W(\Delta G^{\text{hairpin}}_{i,j}(\phi)), & j-i \geq 4 \\ \mathbb{0}, & \text{otherwise} \end{cases}$$

$$\oplus \begin{cases} \textsc{InteriorIntra}(i,j,\phi), & j-1 \geq 6 \\ \mathbb{0}, & \text{otherwise} \end{cases}$$

$$\oplus \begin{cases} \textsc{MultiCoaxIntra}(i,j,\phi) \oplus \textsc{MultiDangleIntra}(i,j,\phi), & j-i \geq 11 \\ \mathbb{0}, & \text{otherwise} \end{cases}$$

Figure S25: $R^b_{\text{INTRA}}$ recursion with coaxial and dangle stacking. Top: recursion diagram. Bottom: recursion equation.

$$\textsc{MultiCoaxIntra}(i,j,\phi) \equiv M_{\text{single}} \oplus M_{\text{mix}}$$

$$
\begin{aligned}
\text{where } M_{\text{single}} &\equiv C_{0s} \otimes W(\Delta G^{\text{multi}}_{\text{init},\mu_i} + 2\Delta G^{\text{multi}}_{\text{bp},\mu_i} + \Delta G^{\text{terminalbp}}_{j,i}(\phi)) \\
C_{0s} &\equiv \begin{cases} C_{1s} \oplus C_{2s}, & \mu_j = \mu_{j-1} = \mu_{i+1} = \mu_i \\ \mathbb{0}, & \text{otherwise} \end{cases} \\
C_{1s} &\equiv \begin{cases} \text{DOT}\left(Q^{m,\text{single}}_{i+1,\overline{d}}, Q^{b}_{\overline{d}+1,j-1}, W(\Delta G^{\text{coax}}_{\overline{d}+1,j-1,i}(\phi) + \Delta G^{\text{terminalbp}}_{\overline{d}+1,j-1}(\phi))\right), & \mu_{d+1} = \mu_{j-1} \\ \mathbb{0}, & \text{otherwise} \end{cases} \\
C_{2s} &\equiv \begin{cases} \text{DOT}\left(Q^{b}_{i+1,\overline{d}}, Q^{m,\text{single}}_{\overline{d}+1,j-1}, W(\Delta G^{\text{coax}}_{j,i,\overline{d}}(\phi) + \Delta G^{\text{terminalbp}}_{i+1,\overline{d}}(\phi))\right), & \mu_{d} = \mu_{i+1} \\ \mathbb{0}, & \text{otherwise} \end{cases} \\
\text{and } M_{\text{mix}} &\equiv (C_{0m} \oplus C_{1m} \oplus C_{2m}) \otimes W(\Delta G^{\text{multi}}_{\text{init,mix}} + \Delta G^{\text{multi}}_{\text{bp},\sigma} + \Delta G^{\text{terminalbp}}_{j,i}(\phi)) \\
C_{0m} &\equiv \begin{cases} \mathbb{0}, & \mu_j = \mu_{j-1} = \mu_{i+1} = \mu_i \\ C_{1s} \otimes W(\Delta G^{\text{multi}}_{\text{bp},\mu_{j-1}}) \oplus C_{2s} \otimes W(\Delta G^{\text{multi}}_{\text{bp},\mu_{i+1}}), & \text{otherwise} \end{cases} \\
C_{1m} &\equiv \begin{cases} \text{DOT}\left(Q^{m,\text{mix}}_{i+1,\overline{d}}, Q^{b}_{\overline{d}+1,j-1}, W(\Delta G^{\text{coax}}_{\overline{d}+1,j-1,i}(\phi) + \Delta G^{\text{terminalbp}}_{\overline{d}+1,j-1}(\phi) + \Delta G^{\text{multi}}_{\text{bp},\mu_{j-1}})\right), & \mu_{d+1} = \mu_{j-1} \\ \mathbb{0}, & \text{otherwise} \end{cases} \\
&\oplus \begin{cases} \text{DOT}\left(Q^{m,\text{mix}}_{i+1,\overline{d}}, Q^{b}_{\overline{d}+1,j-1}, W(\Delta G^{\text{coax}}_{\overline{d}+1,j-1,i}(\phi) + \Delta G^{\text{terminalbp}}_{\overline{d}+1,j-1}(\phi) + \Delta G^{\text{multi}}_{\text{bp,mix}})\right), & \mu_{d+1} \neq \mu_{j-1} \\ \mathbb{0}, & \text{otherwise} \end{cases} \\
&\oplus \begin{cases} \text{DOT}\left(Q^{m,\text{single}}_{i+1,\overline{d}}, Q^{b}_{\overline{d}+1,j-1}, W(\Delta G^{\text{coax}}_{\overline{d}+1,j-1,i}(\phi) + \Delta G^{\text{terminalbp}}_{\overline{d}+1,j-1}(\phi) + \Delta G^{\text{multi}}_{\text{bp,mix}})\right), & \mu_{d+1} \neq \mu_{j-1} \\ \mathbb{0}, & \text{otherwise} \end{cases} \\
C_{2m} &\equiv \begin{cases} \text{DOT}\left(Q^{b}_{i+1,\overline{d}}, Q^{m,\text{mix}}_{\overline{d}+1,j-1}, W(\Delta G^{\text{coax}}_{j,i,\overline{d}}(\phi) + \Delta G^{\text{terminalbp}}_{i+1,\overline{d}}(\phi) + \Delta G^{\text{multi}}_{\text{bp},\mu_{i+1}})\right), & \mu_{d} = \mu_{i+1} \\ \mathbb{0}, & \text{otherwise} \end{cases} \\
&\oplus \begin{cases} \text{DOT}\left(Q^{b}_{i+1,\overline{d}}, Q^{m,\text{mix}}_{\overline{d}+1,j-1}, W(\Delta G^{\text{coax}}_{j,i,\overline{d}}(\phi) + \Delta G^{\text{terminalbp}}_{i+1,\overline{d}}(\phi) + \Delta G^{\text{multi}}_{\text{bp,mix}})\right), & \mu_{d} \neq \mu_{i+1} \\ \mathbb{0}, & \text{otherwise} \end{cases} \\
&\oplus \begin{cases} \text{DOT}\left(Q^{b}_{i+1,\overline{d}}, Q^{m,\text{single}}_{\overline{d}+1,j-1}, W(\Delta G^{\text{coax}}_{j,i,\overline{d}}(\phi) + \Delta G^{\text{terminalbp}}_{i+1,\overline{d}}(\phi) + \Delta G^{\text{multi}}_{\text{bp,mix}})\right), & \mu_{d} \neq \mu_{i+1} \\ \mathbb{0}, & \text{otherwise} \end{cases} \\
\text{with } \sigma &\equiv \begin{cases} \mu_i, & \mu_j = \mu_i \\ \text{mix}, & \text{otherwise} \end{cases} \\
\overline{d} &\equiv [i+5 : j-6]
\end{aligned}
\tag{S160}
$$

$$\textsc{MultiDangleIntra}(i,j,\phi) \equiv \bigoplus_{\substack{k\in\{0,1\}\\ l\in\{0,1\}}} \begin{cases} M_{\text{single}} \oplus M_{\text{mix}}, & (j-l)-(i+k) \geq 11 \\ \mathbb{0}, & \text{otherwise} \end{cases}$$

$$\begin{aligned}
\text{where } M_{\text{single}} &\equiv C_{0s} \otimes W(\Delta G^{\text{dangle}}_{j-l,j,i,i+k}(\phi) + \Delta G^{\text{multi}}_{\text{init},\mu_i} + \Delta G^{\text{multi}}_{\text{bp},\mu_i} + (k+l)\Delta G^{\text{multi}}_{\text{nt},\mu_i} + \Delta G^{\text{terminalbp}}_{j,i}(\phi)) \\
C_{0s} &\equiv \begin{cases} C_{1s} \oplus C_{2s} \oplus C_{3s}, & \textsc{IsSingle}(i, i+k+1, \mu_i) \text{ and } \textsc{IsSingle}(j-l-1, j, \mu_i) \\ \mathbb{0}, & \text{otherwise} \end{cases} \\
C_{1s} &\equiv \text{DOT}\left(Q^{m,\text{single}}_{i+k+1,\overline{d}}, Q^{ms,\text{single}}_{\overline{d}+1,j-l-1}\right) \\
C_{2s} &\equiv \begin{cases} \text{DOT}\left(Q^{ma,\text{single}}_{i+k+1,\overline{e}-1}, Q^{mcs,\text{single}}_{\overline{e},j-l-1}\right), & (j-l)-(i+k) \geq 12 \\ \mathbb{0}, & \text{otherwise} \end{cases} \\
C_{3s} &\equiv Q^{mcs,\text{single}}_{i+k+1,j-l-1} \\
\text{and } M_{\text{mix}} &\equiv (C_{0m} \oplus C_{1m} \oplus C_{2m} \oplus C_{3m}) \\
&\quad \otimes W(\Delta G^{\text{dangle}}_{j-l,j,i,i+k}(\phi) + \Delta G^{\text{multi}}_{\text{init},\text{mix}} + \Delta G^{\text{multi}}_{\text{bp},\sigma} + k\Delta G^{\text{multi}}_{\text{nt},\mu_{i+k}} + l\Delta G^{\text{multi}}_{\text{nt},\mu_{j-l}} + \Delta G^{\text{terminalbp}}_{j,i}(\phi)) \\
C_{0m} &\equiv \begin{cases} \mathbb{0}, & \textsc{IsSingle}(i, i+k+1, \mu_i) \text{ and } \textsc{IsSingle}(j-l-1, j, \mu_i) \\ C_{1s} \oplus C_{2s} \oplus C_{3s}, & \text{otherwise} \end{cases} \\
C_{1m} &\equiv \text{DOT}\left(Q^{m,\text{mix}}_{i+k+1,\overline{d}}, Q^{ms,\text{single}}_{\overline{d}+1,j-l-1}\right) \oplus \text{DOT}\left(Q^{m,\text{single}}_{i+k+1,\overline{d}}, Q^{ms,\text{mix}}_{\overline{d}+1,j-l-1}\right) \oplus \text{DOT}\left(Q^{m,\text{mix}}_{i+k+1,\overline{d}}, Q^{ms,\text{mix}}_{\overline{d}+1,j-l-1}\right) \\
C_{2m} &\equiv \begin{cases} \text{DOT}\left(Q^{ma,\text{single}}_{i+k+1,\overline{e}-1}, Q^{mcs,\text{mix}}_{\overline{e},j-l-1}\right) \oplus \text{DOT}\left(Q^{ma,\text{mix}}_{i+k+1,\overline{e}-1}, Q^{mcs,\text{single}}_{\overline{e},j-l-1}\right) & \\ \oplus\, \text{DOT}\left(Q^{ma,\text{mix}}_{i+k+1,\overline{e}-1}, Q^{mcs,\text{mix}}_{\overline{e},j-l-1}\right), & (j-l)-(i+k) \geq 12 \\ \mathbb{0}, & \text{otherwise} \end{cases} \\
C_{3m} &\equiv Q^{mcs,\text{mix}}_{i+k+1,j-l-1} \\
\text{with } \sigma &\equiv \begin{cases} \mu_i, & \mu_j = \mu_i \\ \text{mix}, & \text{otherwise} \end{cases} \\
\overline{d} &\equiv [i+k+5 : j-l-6], \quad \overline{e} \equiv [i+k+2 : j-l-10]
\end{aligned} \tag{S161}$$

$\boldsymbol{R^{ms}_{\textbf{INTRA}}}$ **recursion with coaxial and dangle stacking.** The $R^{b}_{\text{INTRA}}(i,j,\phi)$ recursion references $Q^{ms}$ elements that are computed using the $R^{ms}_{\text{INTRA}}$ recursion shown in Figure S26. $R^{ms}_{\text{INTRA}}(i,j,\phi)$ operates on a conditional ensemble for subsequence $[i,j]$ in a multiloop context containing one stacking state starting at $i$ and ending in the interval $[i+1,j]$ (depicted as a dashed line between $i$ and $j$). Like other multiloop recursions in the mixed algorithm, $R^{ms}_{\text{INTRA}}(i,j,\phi)$ is split into $R^{ms,\text{single}}_{\text{INTRA}}(i,j,\phi)$ and $R^{ms,\text{mix}}_{\text{INTRA}}(i,j,\phi)$ (the single and mixed recursions are indicated in the diagram with blue and red backbones, respectively):

- $R^{ms,\text{single}}_{\text{INTRA}}(i,j,\phi)$: This recursion treats the $R^{ms}_{\text{INTRA}}(i,j,\phi)$ conditional ensemble over $[i,j]$ in a multiloop context where all terminal base pairs and unpaired nucleotides are of a single material. There are two cases that are combined using $\oplus$ in the recursion equation:
  - *Coaxial stacking state:* The contribution for the coaxial stacking state in subsequence $[i,j]$ is calculated using a $Q^{mcs,\text{single}}_{i,j}$ element using term $C_{3s}$. Because $Q^{mcs,\text{single}}_{i,j}$ only includes single-material contributions, no further material checks are needed.
  - *Dangle stacking state:* The contribution for the dangle stacking state in subsequence $[i,d]$ is incorporated using a $Q^{md}_{i,d}$ element. For the edge case where $d=j$, the whole region $[i,j]$ is incorporated using a $Q^{md,\text{single}}_{i,j}$ element using term $C_{2s}$. Because $Q^{md,\text{single}}_{i,j}$ only includes single-material contributions, no further material checks are needed. For the base case where $d$ is in $[i+4,j-1]$, then $Q^{md,\text{single}}_{i,d}$ is used to represent the dangle stacking state in $[i,d]$ while the unpaired region $[d+1,j]$ is incorporated using $Q^{ma,\text{single}}_{d+1,j}$. The resulting contribution $Q^{md,\text{single}}_{i,d} \otimes Q^{ma,\text{single}}_{d+1,j}$ of term $C_{1s}$ produces a single-material multiloop only if the material matches for both elements, which is checked with the conitional $\mu_j = \mu_i$. If the conditional is satisfied, the contribution is added to $R^{ms,\text{single}}_{\text{INTRA}}(i,j,\phi)$ using term $C_{0s}$. Otherwise, if the conditional fails, the contribution is added to $R^{ms,\text{mix}}_{\text{INTRA}}(i,j,\phi)$ using term $C_{0m}$. In the conditional, indices $i$ and $j$ are used to check the material since they are constant over vector $\overline{d}$ and apply to all $d$ and $d+1$ in $\overline{d}$.
- $R^{ms,\text{mix}}_{\text{INTRA}}(i,j,\phi)$: This recursion treats the $R^{ms}_{\text{INTRA}}(i,j,\phi)$ conditional ensemble over $[i,j]$ in a multiloop context where the terminal base pairs and unpaired nucleotides overall contain a mixture of materials. There are two cases that are combined using $\oplus$ in the recursion equation:
  - *Coaxial stacking state:* The contribution for the coaxial stacking state in subsequence $[i,j]$ is calculated using a $Q^{mcs,\text{mix}}_{i,j}$ element using term $C_{3m}$. Because $Q^{mcs,\text{mix}}_{i,j}$ only includes mixed contributions, no further material checks are needed.
  - *Dangle stacking state:* The contribution for the dangle stacking state in subsequence $[i,d]$ is incorporated using a $Q^{md}_{i,d}$ element. For the edge case where $d=j$, the whole region $[i,j]$ is incorporated using a $Q^{md,\text{mix}}_{i,j}$ element using term $C_{2m}$. Because $Q^{md,\text{mix}}_{i,j}$ only includes mixed contributions, no further material checks are needed. For the base case where $d$ is in $[i+4,j-1]$, the contributions for the dangle stacking state in subsequence $[i,d]$ are incorporated with a $Q^{md}_{i,d}$ element while the contributions for the unpaired region $[d+1,j]$ are incorporated with a $Q^{ma}_{d+1,j}$ element. Because $Q^{md}_{i,d}$ and $Q^{ma}_{d+1,j}$ are each split into single- and mixed-material recursions, there are 4 possible combinations when multiplying them. The three combinations containing at least one mixed element ($Q^{md,\text{single}}_{i,d} \otimes Q^{ma,\text{mix}}_{d+1,j}$, $Q^{md,\text{mix}}_{i,d} \otimes Q^{ma,\text{single}}_{d+1,j}$, $Q^{md,\text{mix}}_{i,d} \otimes Q^{ma,\text{mix}}_{d+1,j}$) must represent mixed contributions, and are included in $R^{ms,\text{mix}}_{\text{INTRA}}(i,j,\phi)$ using term $C_{1m}$. As noted above, the remaining combination, $Q^{md,\text{single}}_{i,d} \otimes Q^{ma,\text{single}}_{d+1,j}$, calculated in term $C_{1s}$ represents a mixed contribution if $\mu_j \neq \mu_i$, and is incorporated in $R^{ms,\text{mix}}_{\text{INTRA}}(i,j,\phi)$ using term $C_{0m}$.

The shading does not represent any recursion energies as all multiloop contributions are handled by other recursions: 1) there are no terminal base pairs in a multiloop context explicitly defined in this case, 2) there are no unpaired bases in a multiloop context explicitly defined in this case. Note that $R^{ms}$ directly incorporates the $R^{mcs}$ recursion which serves as an efficiency wrapper of the $R^{mc}$ recursion, and hence, $R^{ms}$ is an efficiency wrapper of $R^{mc}$ (the $3'$-most coaxial stacking state in a multiloop context). Note also that $R^{ms}$ is an efficiency wrapper of the $R^{md}$ recursion (the $3'$-most dangle stacking state in a multiloop context). Taken together $R^{mc}$ and $R^{md}$ represent the $3'$-most stacking state in a multiloop context, analogous to $R^{cd}$ representing the $3'$-most stacking state (coaxial or dangle) in an exterior loop context. The reason that $R^{mc}$ (coaxial stacking states) and $R^{md}$ (dangle stacking states) are calculated and stored separately in a multiloop context is that coaxial-only information (stored in element $Q^{mcs}$) is needed for the previously described multiloop edge case (recursion diagrams in the fifth and sixth rows of Figure S25). As a result, coaxial-only information is calculated using the efficiency wrapper $R^{mcs}$ for use in that edge case, and then coaxial-only and dangle-only information are combined by the $R^{ms}$ efficiency wrapper (which is fully analogous to the $R^{s}$ efficiency wrapper in the exterior loop context). With this approach, the operations spent calculating coaxial stacking information for $Q^{mcs}$ elements are not repeated when calculating both coaxial and dangle stacking for $Q^{ms}$ elements.

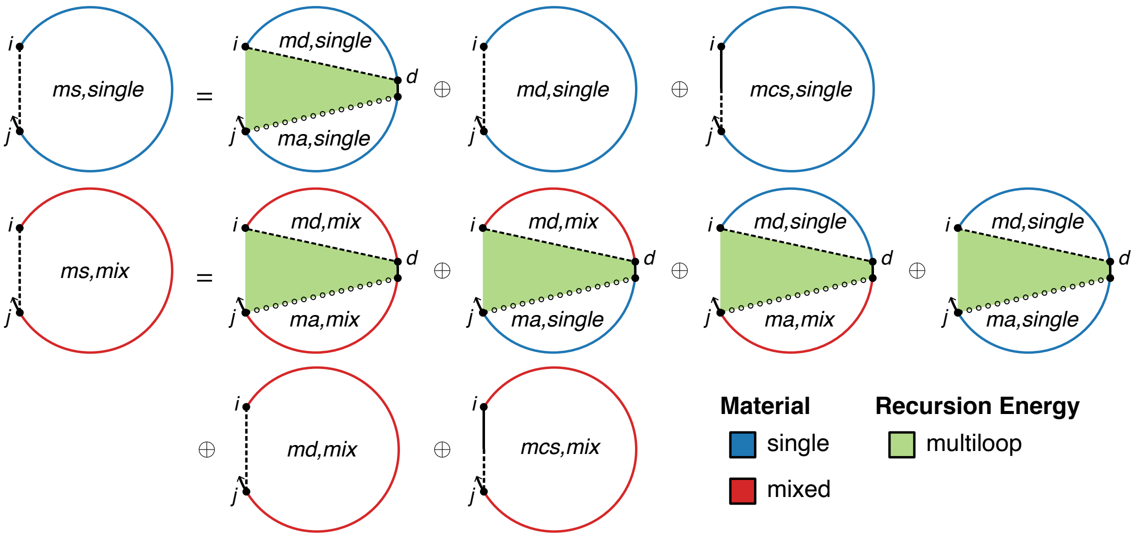

$$R^{ms,\text{single}}_{\text{INTRA}}(i,j,\phi) \equiv C_{0s} \oplus C_{2s} \oplus C_{3s}$$

$$C_{0s} \equiv \begin{cases} C_{1s}, & \mu_j = \mu_i \\ \mathbb{0}, & \text{otherwise} \end{cases}$$

$$C_{1s} \equiv \begin{cases} \text{DOT}\left(Q^{md,\text{single}}_{i,\overline{d}}, Q^{ma,\text{single}}_{\overline{d}+1,j}\right), & j-i \geq 5 \\ \mathbb{0}, & \text{otherwise} \end{cases}$$

$$C_{2s} \equiv \begin{cases} Q^{md,\text{single}}_{i,j}, & j-i \geq 4 \\ \mathbb{0}, & \text{otherwise} \end{cases}$$

$$C_{3s} \equiv Q^{mcs,\text{single}}_{i,j}$$

$$R^{ms,\text{mix}}_{\text{INTRA}}(i,j,\phi) \equiv C_{0m} \oplus C_{1m} \oplus C_{2m} \oplus C_{3m}$$

$$C_{0m} \equiv \begin{cases} \mathbb{0}, & \mu_j = \mu_i \\ C_{1s}, & \text{otherwise} \end{cases}$$

$$C_{1m} \equiv \begin{cases} \text{DOT}\left(Q^{md,\text{mix}}_{i,\overline{d}}, Q^{ma,\text{single}}_{\overline{d}+1,j}\right) \oplus \text{DOT}\left(Q^{md,\text{single}}_{i,\overline{d}}, Q^{ma,\text{mix}}_{\overline{d}+1,j}\right) \oplus \text{DOT}\left(Q^{md,\text{single}}_{i,\overline{d}}, Q^{ma,\text{mix}}_{\overline{d}+1,j}\right), & j-i \geq 5 \\ \mathbb{0}, & \text{otherwise} \end{cases}$$

$$C_{2m} \equiv \begin{cases} Q^{md,\text{mix}}_{i,j}, & j-i \geq 4 \\ \mathbb{0}, & \text{otherwise} \end{cases}$$

$$C_{3m} \equiv Q^{mcs,\text{mix}}_{i,j}$$

where $\overline{d} \equiv [i+4 : j-1]$

Figure S26: $R^{ms}_{\text{INTRA}}$ recursion with coaxial and dangle stacking. Top: recursion diagram. Bottom: recursion equation.

$\boldsymbol{R^{ma}_{\text{INTRA}}}$ **recursion with coaxial and dangle stacking.** The $R^{b}_{\text{INTRA}}(i,j,\phi)$ and $R^{ms}_{\text{INTRA}}(i,j,\phi)$ recursions reference $Q^{ma}$ elements that are computed using the $R^{ma}_{\text{INTRA}}$ recursion shown in Figure S27. $R^{ma}_{\text{INTRA}}(i,j,\phi)$ treats the conditional ensemble of subsequence $[i,j]$ in a multiloop context containing no terminal base pairs (the hollow dotted line between $i$ and $j$ indicates that there is no base pairing in $[i,j]$). This recursion is used in the mixed-material algorithm to switch the $\Delta G^{\text{multi}}_{\text{nt}}$ parameter depending on the material of each nucleotide. Note that it is identical to $R^{ma}_{\text{INTRA}}(i,j,\phi)$ in the non-stacking ensemble (Figure S10). The single-material algorithm does not incorporate this recursion since parameter switching is not necessary. Like other multiloop recursions in the mixed algorithm, $R^{ma}_{\text{INTRA}}(i,j,\phi)$ is split into $R^{ma,\text{single}}_{\text{INTRA}}(i,j,\phi)$ and $R^{ma,\text{mix}}_{\text{INTRA}}(i,j,\phi)$ (the single- and mixed-material recursions are indicated in the recursion diagram by blue and red backbones, respectively):

- $R^{ma,\text{single}}_{\text{INTRA}}(i,j,\phi)$: This recursion calculates contributions from subsequences $[i,j]$ composed of a single material. The shading corresponds to the energy per unpaired nucleotide in a multiloop context for the material of nucleotide $j$, $\Delta G^{\text{multi}}_{\text{nt},\mu_j}$. For the edge case with $i=j$, then the subsequence is a single nucleotide and therefore a single material, yielding a contribution $W(\Delta G^{\text{multi}}_{\text{nt},\mu_i})$ in term $C_{2s}$. For the base case with $i<j$, the $Q^{ma,\text{single}}_{i,j-1}$ element is used to represent the single-material contribution from subsequence $[i,j-1]$ (non-zero only if $[i,j-1]$ is a single material) and the contribution for unpaired nucleotide $j$ is $W(\Delta G^{\text{multi}}_{\text{nt},\mu_j})$. The overall contribution, $Q^{ma,\text{single}}_{i,j-1} \otimes W(\Delta G^{\text{multi}}_{\text{nt},\mu_j})$, represents a single material only if the materials of both terms match, which is checked with the conditional $\mu_j = \mu_{j-1}$. If the conditional is satisfied, the contribution is added to $R^{ma,\text{single}}_{\text{INTRA}}(i,j,\phi)$ using term $C_{0s}$. Otherwise, if the conditional fails, the contribution is added to $R^{ma,\text{mix}}_{\text{INTRA}}(i,j,\phi)$ using term $C_{0m}$.
- $R^{ma,\text{mix}}_{\text{INTRA}}(i,j,\phi)$: This recursion calculates contributions from subsequence $[i,j]$ composed of mixed materials. The shading corresponds to the energy per unpaired nucleotide in a multiloop context for the material of nucleotide $j$, $\Delta G^{\text{multi}}_{\text{nt},\mu_j}$. For the edge case with $i=j$, the contribution is $\mathbb{0}$ since a single nucleotide cannot have more than one material. For the base case with $i<j$, the $Q^{ma,\text{mix}}_{i,j-1}$ element is used to represent the mixed-material contribution of subsequence $[i,j-1]$ (non-zero only if $[i,j-1]$ contains a mixture of materials) and the contribution for unpaired nucleotide $j$ is $W(\Delta G^{\text{multi}}_{\text{nt},\mu_j})$, leading to an overall contribution of $Q^{ma,\text{mix}}_{i,j-1} \otimes W(\Delta G^{\text{multi}}_{\text{nt},\mu_j})$ added to $R^{ma,\text{mix}}_{\text{INTRA}}(i,j,\phi)$ using term $C_{1m}$. As noted above, the contribution, $Q^{ma,\text{single}}_{i,j-1} \otimes W(\Delta G^{\text{multi}}_{\text{nt},\mu_j})$, calculated in term $C_{1s}$ represents a mixed contribution if $\mu_j \neq \mu_{j-1}$, and is incorporated in $R^{ma,\text{mix}}_{\text{INTRA}}(i,j,\phi)$ using term $C_{0m}$.

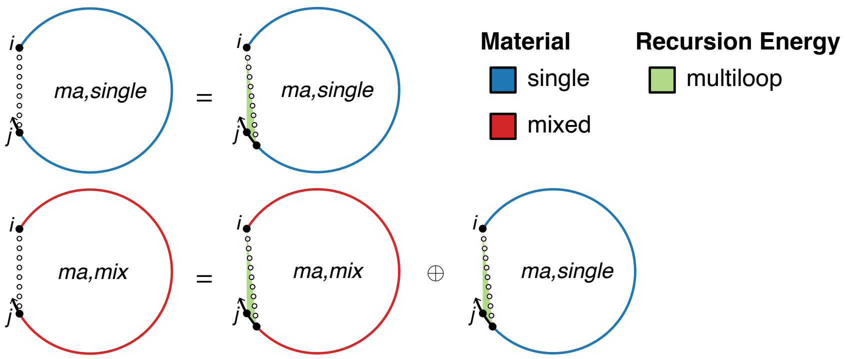

$$R^{ma,\text{single}}_{\text{INTRA}}(i,j,\phi) \equiv C_{0s} \oplus C_{2s}$$

$$C_{0s} \equiv \begin{cases} C_{1s}, & \mu_j = \mu_{j-1} \\ \mathbb{0}, & \text{otherwise} \end{cases}$$

$$C_{1s} \equiv \begin{cases} Q^{ma,\text{single}}_{i,j-1} \otimes W(\Delta G^{\text{multi}}_{\text{nt},\mu_i}), & i < j \\ \mathbb{0}, & \text{otherwise} \end{cases}$$

$$C_{2s} \equiv \begin{cases} W(\Delta G^{\text{multi}}_{\text{nt},\mu_i}), & i = j \\ \mathbb{0}, & \text{otherwise} \end{cases}$$

$$R^{ma,\text{mix}}_{\text{INTRA}}(i,j,\phi) \equiv C_{0m} \oplus C_{1m}$$

$$C_{0m} \equiv \begin{cases} \mathbb{0}, & \mu_j = \mu_{j-1} \\ C_{1s}, & \text{otherwise} \end{cases}$$

$$C_{1m} \equiv \begin{cases} Q^{ma,\text{mix}}_{i,j-1} \otimes W(\Delta G^{\text{multi}}_{\text{nt},\mu_j}), & i < j \\ \mathbb{0}, & \text{otherwise} \end{cases}$$

Figure S27: $R^{ma}_{\text{INTRA}}(i,j,\phi)$ recursion with coaxial and dangle stacking. Top: recursion diagram. Bottom: recursion equation. Note that this recursion is the same regardless of whether coaxial and dangle stacking are included in the structural ensemble.

$\boldsymbol{R^{mcs}_{\text{INTRA}}}$ **recursion with coaxial and dangle stacking.** The $R^{ms}_{\text{INTRA}}(i,j,\phi)$ recursion references $Q^{mcs}$ elements that are computed using the $R^{mcs}_{\text{INTRA}}$ recursion displayed in Figure S28. $R^{mcs}_{\text{INTRA}}(i,j,\phi)$ operates on a conditional ensemble for subsequence $[i,j]$ in a multiloop context containing one coaxial stacking state starting at $i$ and ending in the interval $[i+1,j]$ (depicted as a half-solid/half-dashed line between $i$ and $j$). Like other multiloop recursions in the mixed algorithm, $R^{mcs}_{\text{INTRA}}(i,j,\phi)$ is split into $R^{mcs,\text{single}}_{\text{INTRA}}(i,j,\phi)$ and $R^{mcs,\text{mix}}_{\text{INTRA}}(i,j,\phi)$ (the single and mixed recursions are indicated in the diagram with blue and red backbones, respectively):

- $R^{mcs,\text{single}}_{\text{INTRA}}(i,j,\phi)$: This recursion treats the $R^{mcs}_{\text{INTRA}}(i,j,\phi)$ conditional ensemble over $[i,j]$ in a multiloop context where all terminal base pairs and unpaired nucleotides are of a single material. For the edge case where $d=j$, then whole region $[i,j]$ is incorporated using a $Q^{mc,\text{single}}_{i,j}$ element using term $C_{2s}$. No further material checks are needed since the $R^{mc,\text{single}}_{\text{INTRA}}(i,j,\phi)$ recursion only includes single-material contributions. For the base case where $d$ is in $[i+9,j-1]$, then $Q^{mc,\text{single}}_{i,d}$ is used to represent the coaxial stacking state in $[i,d]$ while the unpaired region $[d+1,j]$ is incorporated using $Q^{ma,\text{single}}_{d+1,j}$. The resulting contribution $Q^{mc,\text{single}}_{i,d}\otimes Q^{ma,\text{single}}_{d+1,j}$ of term $C_{1s}$ produces a single-material multiloop only if the material matches for both elements, which is checked with the conitional $\mu_j=\mu_i$. If the conditional is satisfied, the contribution is added to $R^{mcs,\text{single}}_{\text{INTRA}}(i,j,\phi)$ using term $C_{0s}$. Otherwise, if the conditional fails, the contribution is added to $R^{mcs,\text{mix}}_{\text{INTRA}}(i,j,\phi)$ using term $C_{0m}$. In the conditional, indices $i$ and $j$ are used to check the material since they are constant over vector $\bar{d}$ and apply to all $d$ and $d+1$ in $\bar{d}$.

- $R^{mcs,\text{mix}}_{\text{INTRA}}(i,j,\phi)$: This recursion treats the $R^{mcs}_{\text{INTRA}}(i,j,\phi)$ conditional ensemble over $[i,j]$ in a multiloop context where the terminal base pairs and unpaired nucleotides overall contain a mixture of materials. For the edge case where $d=j$, then whole region $[i,j]$ is incorporated using a $Q^{mc,\text{mix}}_{i,j}$ element using term $C_{2m}$. No further material checks are needed since the $R^{mc,\text{mix}}_{\text{INTRA}}(i,j,\phi)$ recursion only includes mixed material contributions. For the base case where $d$ is in $[i+9,j-1]$, then $Q^{mc}_{i,d}$ is used to represent the coaxial stacking state in $[i,d]$ while the unpaired region $[d+1,j]$ is incorporated using a $Q^{ma}_{d+1,j}$ element. Because $Q^{mc}_{i,d}$ and $Q^{ma}_{d+1,j}$ are each split into single- and mixed-material recursions, there are 4 possible combinations when multiplying them. The three combinations containing at least one mixed element ($Q^{mc,\text{single}}_{i,d}\otimes Q^{ma,\text{mix}}_{d+1,j}$, $Q^{mc,\text{mix}}_{i,d}\otimes Q^{ma,\text{single}}_{d+1,j}$, $Q^{mc,\text{mix}}_{i,d}\otimes Q^{ma,\text{mix}}_{d+1,j}$) must represent mixed contributions, and are included in $R^{mcs,\text{mix}}_{\text{INTRA}}(i,j,\phi)$ using term $C_{1m}$. As noted above, the remaining combination, $Q^{mc,\text{single}}_{i,d}\otimes Q^{ma,\text{single}}_{d+1,j}$, calculated in term $C_{1s}$ represents a mixed contribution only if $\mu_j\neq\mu_i$, and is incorporated in $R^{mcs,\text{mix}}_{\text{INTRA}}(i,j,\phi)$ using term $C_{0m}$.

The shading does not represent any recursion energies as all multiloop contributions are handled by other recursions: 1) there are no terminal base pairs in a multiloop context explicitly defined in this case, 2) there are no unpaired bases in a multiloop context explicitly defined in this case. Note that the $R^{mcs}$ recursion serves as an efficiency wrapper of the $R^{mc}$ recursion (here, representing the $3'$-most coaxial stacking state in a multiloop context). The index limits in the recursion equation reflect the steric requirement that there be at least 8 intervening bases between $i$ and $d$ (because the $R^{mc}$ recursion incorporates a coaxial stack involving two adjacent terminal base pairs such that $i$ and $d$ are paired to intervening adjacent bases).

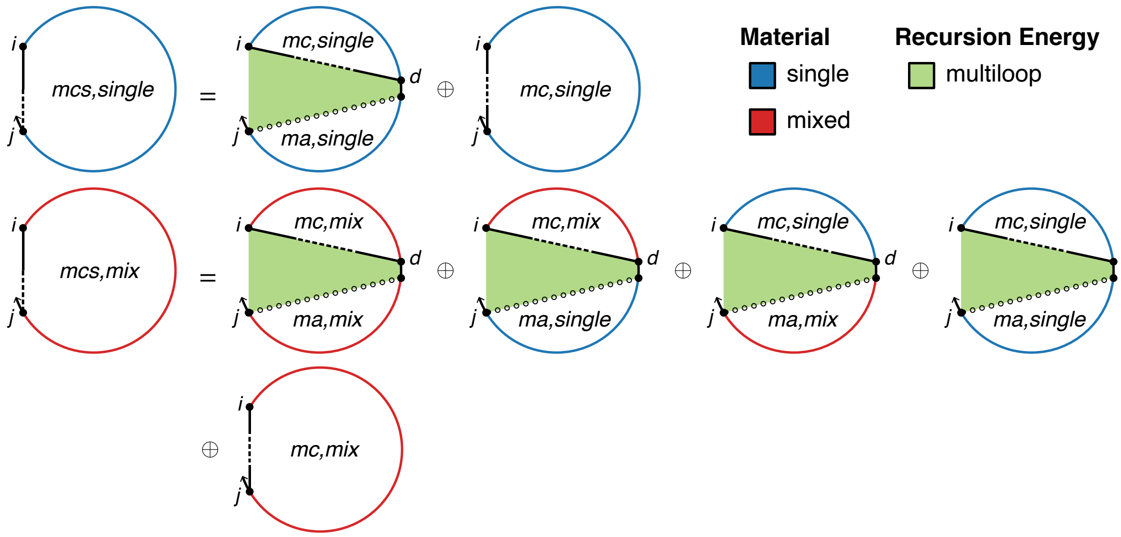

$$R^{mcs,\text{single}}_{\text{INTRA}}(i,j,\phi) \equiv C_{0s} \oplus C_{2s}$$

$$C_{0s} \equiv \begin{cases} C_{1s}, & \mu_j = \mu_i \\ \mathbb{0}, & \text{otherwise} \end{cases}$$

$$C_{1s} \equiv \begin{cases} \text{DOT}\left(Q^{mc,\text{single}}_{i,\overline{d}}, Q^{ma,\text{single}}_{\overline{d}+1,j}\right), & j-i \geq 10 \\ \mathbb{0}, & \text{otherwise} \end{cases}$$

$$C_{2s} \equiv \begin{cases} Q^{mc,\text{single}}_{i,j}, & j-i \geq 9 \\ \mathbb{0}, & \text{otherwise} \end{cases}$$

$$R^{mcs,\text{mix}}_{\text{INTRA}}(i,j,\phi) \equiv C_{0m} \oplus C_{1m} \oplus C_{2m}$$

$$C_{0m} \equiv \begin{cases} \mathbb{0}, & \mu_j = \mu_i \\ C_{1s}, & \text{otherwise} \end{cases}$$

$$C_{1m} \equiv \begin{cases} \text{DOT}\left(Q^{mc,\text{mix}}_{i,\overline{d}}, Q^{ma,\text{single}}_{\overline{d}+1,j}\right) \oplus \text{DOT}\left(Q^{mc,\text{single}}_{i,\overline{d}}, Q^{ma,\text{mix}}_{\overline{d}+1,j}\right) \oplus \text{DOT}\left(Q^{mc,\text{mix}}_{i,\overline{d}}, Q^{ma,\text{mix}}_{\overline{d}+1,j}\right), & j-i \geq 10 \\ \mathbb{0}, & \text{otherwise} \end{cases}$$

$$C_{2m} \equiv \begin{cases} Q^{mc,\text{mix}}_{i,j}, & j-i \geq 9 \\ \mathbb{0}, & \text{otherwise} \end{cases}$$

where $\overline{d} \equiv [i+9 : j-1]$

Figure S28: $R^{mcs}_{\text{INTRA}}$ recursion with coaxial and dangle stacking. Top: recursion diagram. Bottom: recursion equation.

$\boldsymbol{R^{mc}_{\text{INTRA}}}$ **recursion with coaxial and dangle stacking.** The $R^{mcs}_{\text{INTRA}}(i, j, \phi)$ recursion references $Q^{mc}$ elements that are computed using the $R^{mc}_{\text{INTRA}}$ recursion displayed in Figure S29. This recursion treats a single coaxial stacking state in a multiloop context (depicted as a straight line between $i$ and $j$ that is solid at both ends and dashed in the middle to indicate that $i$ and $j$ are both base-paired but not to each other). Two adjacent terminal base pairs ($i \cdot d$ and $d+1 \cdot j$) coaxially stack on each other. Each terminal base pair is either a single material (denoted in the recursion diagram by a blue base pair, if the material of the two nucleotides matches) or mixed material (denoted by a red base pair, if the material of the two nucleotides does not match). The contributions of subsequences $[i, d]$ and $[d+1, j]$ are incorporated using $Q^b_{i,d}$ and $Q^b_{d+1,j}$ elements in term $C_{\text{stack}}$. The index limits in the recursion equation reflect the steric requirement that there be at least 3 intervening bases between $i$ and $d$ and at least 3 intervening bases between $d+1$ and $j$. Like other multiloop recursions in the mixed algorithm, $R^{mc}_{\text{INTRA}}(i, j, \phi)$ is split into $R^{mc,\text{single}}_{\text{INTRA}}(i, j, \phi)$ and $R^{mc,\text{mix}}_{\text{INTRA}}(i, j, \phi)$ (the single and mixed recursions are indicated in the diagram with blue and red backbones, respectively):

- $R^{mc,\text{single}}_{\text{INTRA}}(i, j, \phi)$: This recursion treats the $R^{mc}_{\text{INTRA}}(i, j, \phi)$ conditional ensemble over $[i, j]$ where terminal base pairs $i \cdot d$ and $d+1 \cdot j$ are of a single material. The case where both terminal base pairs are single-material (checked with conditional "$\mu_d = \mu_i$ and $\mu_{d+1} = \mu_j$" in term $C_{1s}$) leads to a single-material multiloop only if the material matches for both terminal base pairs, which is checked with the conditional $\mu_j = \mu_i$. If the conditional is satisfied, the contribution is added to $R^{mc,\text{single}}_{\text{INTRA}}(i, j, \phi)$ using term $C_{0s}$. Otherwise, if the conditional fails, the contribution is added to $R^{mc,\text{mix}}_{\text{INTRA}}(i, j, \phi)$ using term $C_{0m}$. In the conditional, indices $i$ and $j$ are used to check the material since they are constant over vector $\overline{d}$ and apply to all $d$ and $d+1$ in $\overline{d}$.

- $R^{mc,\text{mix}}_{\text{INTRA}}(i, j, \phi)$: This recursion treats the $R^{mc}_{\text{INTRA}}(i, j, \phi)$ conditional ensemble over $[i, j]$ where the terminal base pairs $i \cdot d$ and $d+1 \cdot j$ overall contain a mixture of materials. Because each of the terminal base pairs can be single- or mixed-material, there are four possible single/mix combinations for $Q^b_{i,d} \otimes Q^b_{d+1,j}$. The three combinations containing at least one mixed base pair are included in $R^{mc,\text{mix}}_{\text{INTRA}}(i, j, \phi)$ using term $C_{1m}$. As noted above, the remaining combination of term $C_{1s}$ where each terminal base pair is single-material represents a mixed contribution only if $\mu_j \neq \mu_i$ and is incorporated in $R^{mc,\text{mix}}_{\text{INTRA}}(i, j, \phi)$ using term $C_{0m}$.

Shading corresponds to three kinds of recursion energy: 1) the sequence-independent penalties for two terminal base pairs in a multiloop context (for base pair $i \cdot d$, $\Delta G^{\text{multi}}_{\text{bp},\mu_i}$ if $\mu_d = \mu_i$, and $\Delta G^{\text{multi}}_{\text{bp,mix}}$ otherwise; for base pair $d+1 \cdot j$, $\Delta G^{\text{multi}}_{\text{bp},\mu_j}$ if $\mu_{d+1} = \mu_j$, and $\Delta G^{\text{multi}}_{\text{bp,mix}}$ otherwise), 2) the sequence-dependent penalties for two terminal base pairs in a multiloop context ($\Delta G^{\text{terminalbp}}_{i,d}(\phi)$ and $\Delta G^{\text{terminalbp}}_{d+1,j}(\phi)$, dependent on the sequence and material of base pairs $i \cdot d$ and $d+1 \cdot j$), 3) the sequence-dependent coaxial stacking free energy $\Delta G^{\text{coax}}_{i,d,j}(\phi)$ (dependent on the sequence and material of base pairs $i \cdot d$ and $d+1 \cdot j$); note that $\Delta G^{\text{coax}}_{i,d,j}(\phi)$ requires only 3 indices because $d+1$ is implied by $d$.

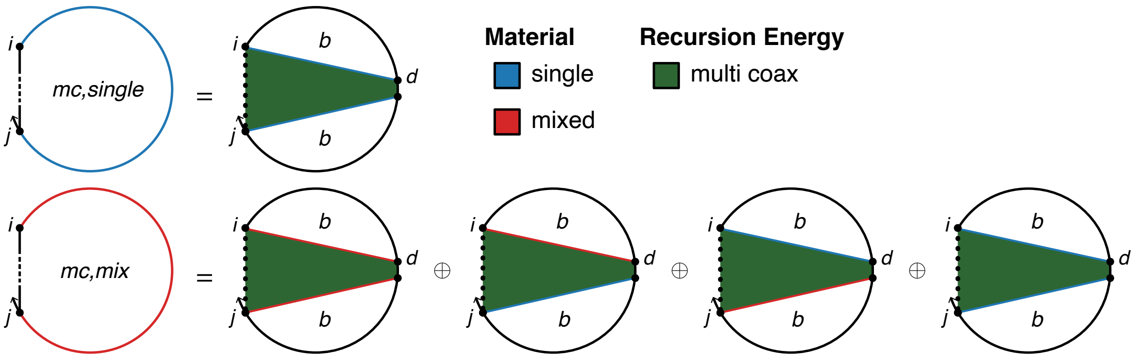

$$R^{mc,\text{single}}_{\text{INTRA}}(i,j,\phi) \equiv C_{0s}$$

$$\text{where } C_{0s} \equiv \begin{cases} C_{1s}, & \mu_j = \mu_i \\ \mathbb{0}, & \text{otherwise} \end{cases}$$

$$C_{1s} \equiv \begin{cases} C_{\text{stack}} \otimes W(\Delta G^{\text{multi}}_{\text{bp},\mu_i} + \Delta G^{\text{multi}}_{\text{bp},\mu_j}), & \mu_d = \mu_i \text{ and } \mu_{d+1} = \mu_j \\ \mathbb{0}, & \text{otherwise} \end{cases}$$

$$C_{\text{stack}} \equiv \begin{cases} \text{DOT}\left(Q^b_{i,\overline{d}}, Q^b_{\overline{d}+1,j}, W(\Delta G^{\text{coax}}_{i,\overline{d},j}(\phi) + \Delta G^{\text{terminalbp}}_{i,\overline{d}}(\phi) + \Delta G^{\text{terminalbp}}_{\overline{d}+1,j}(\phi))\right), & j-i \geq 9 \\ \mathbb{0}, & \text{otherwise} \end{cases}$$

$$R^{mc,\text{mix}}_{\text{INTRA}}(i,j,\phi) \equiv C_{0m} \oplus C_{1m}$$

$$\text{where } C_{0m} \equiv \begin{cases} \mathbb{0}, & \mu_j = \mu_i \\ C_{1s}, & \text{otherwise} \end{cases}$$

$$C_{1m} \equiv \begin{cases} C_{\text{stack}} \otimes W(\Delta G^{\text{multi}}_{\text{bp},\mu_i} + \Delta G^{\text{multi}}_{\text{bp,mix}}), & \mu_d = \mu_i \text{ and } \mu_{d+1} \neq \mu_j \\ \mathbb{0}, & \text{otherwise} \end{cases}$$

$$\oplus \begin{cases} C_{\text{stack}} \otimes W(\Delta G^{\text{multi}}_{\text{bp,mix}} + \Delta G^{\text{multi}}_{\text{bp},\mu_j}), & \mu_d \neq \mu_i \text{ and } \mu_{d+1} = \mu_j \\ \mathbb{0}, & \text{otherwise} \end{cases}$$

$$\oplus \begin{cases} C_{\text{stack}} \otimes W(2\Delta G^{\text{multi}}_{\text{bp,mix}}), & \mu_d \neq \mu_i \text{ and } \mu_{d+1} \neq \mu_j \\ \mathbb{0}, & \text{otherwise} \end{cases}$$

$$\text{with } \overline{d} \equiv [i+4 : j-5]$$

Figure S29: $R^{mc}_{\text{INTRA}}$ recursion with coaxial and dangle stacking. Top: recursion diagram. Bottom: recursion equation.

$\boldsymbol{R^{md}_{\text{INTRA}}}$ **recursion with coaxial and dangle stacking.** The $R^{ms}_{\text{INTRA}}(i, j, \phi)$ recursion references $Q^{md}$ elements that are computed using the $R^{md}_{\text{INTRA}}$ recursion displayed in Figure S30. This recursion treats a single dangle stacking state (depicted as a dashed line between $i$ and $j$) in a multiloop context with either zero, one, or two unpaired nucleotides dangle stacking on an adjacent terminal base pair ($i + k \cdot j - l$). The recursion diagram represents these four alternative dangle stacking states corresponding to no dangles, $5'$ dangle, $3'$ dangle, or terminal mismatch (see Figure S21 for details). The terminal base pair is either a single material (denoted in the recursion diagram with a blue base pair, if $\mu_{i+k} = \mu_{j-l}$), or mixed-material (denoted by a red base pair, if $\mu_{i+k} \neq \mu_{j-l}$). The contribution of subsequence $[i + k, j - l]$ is incorporated using a $Q^b_{i+k,j-l}$ element in term $C_{\text{stack}}$. The index limits in the recursion equation reflect the steric requirement that there be at least 3 intervening bases between $i + k$ and $j - l$. Like other multiloop recursions in the mixed algorithm, $R^{md}_{\text{INTRA}}(i, j, \phi)$ is split into $R^{md,\text{single}}_{\text{INTRA}}(i, j, \phi)$ and $R^{md,\text{mix}}_{\text{INTRA}}(i, j, \phi)$ (the single and mixed recursions are indicated in the diagram with blue and red backbones, respectively):

- $R^{md,\text{single}}_{\text{INTRA}}(i, j, \phi)$: This recursion treats the $R^{md}_{\text{INTRA}}(i, j, \phi)$ conditional ensemble over $[i, j]$ where terminal base pair $i + k \cdot j - l$ and any unpaired dangling nucleotides are of a single material.
- $R^{md,\text{mix}}_{\text{INTRA}}(i, j, \phi)$: This recursion treats the $R^{md}_{\text{INTRA}}(i, j, \phi)$ conditional ensemble over $[i, j]$ where terminal base pair $i + k \cdot j - l$ and any unpaired dangling nucleotides overall contain a mixture of materials.

The conditional "$\mu_{i+k} = \mu_{j-l} = \mu_j = \mu_i$" is used to check if the terminal base pair $i + k \cdot j - l$ and any dangling nucleotides are of a single material. If the conditional is satisfied, $C_{\text{stack}}$ is added to $R^{md,\text{single}}_{\text{INTRA}}(i, j, \phi)$. Otherwise, if the conditional is not satisfied, $C_{\text{stack}}$ is added to $R^{md,\text{mix}}_{\text{INTRA}}(i, j, \phi)$. Shading corresponds to four recursion energies: 1) the sequence-independent penalty for one terminal base pair in a multiloop context, $\Delta G^{\text{multi}}_{\text{bp},\sigma}$ (corresponding to base pair $j - l \cdot i + k$), where the parameter depends on the material as determined by the $\sigma$ conditional, 2) the penalty per unpaired nucleotide in a multiloop context $\Delta G^{\text{multi}}_{\text{nt},\mu_i}$ or $\Delta G^{\text{multi}}_{\text{nt},\mu_j}$ (a total of $k + l$ dangling nucleotides; as a result this term is zeroed out when $k = l = 0$), where the parameter depends on the material of each dangling nucleotide ($\mu_i$ or $\mu_j$), 3) the sequence-dependent penalty for a terminal base pair in a multiloop loop context, $\Delta G^{\text{terminalbp}}_{i+k,j-l}(\phi)$ (dependent on the sequence and material of base pair $i + k \cdot j - l$), 4) the sequence-dependent dangle stacking free energy $\Delta G^{\text{dangle}}_{i,i+k,j-l,j}(\phi)$ (dependent on the sequence and material of the terminal base pair and dangle stacking bases) which takes on one of four values corresponding to the four dangle stacking states (see Figure S21).

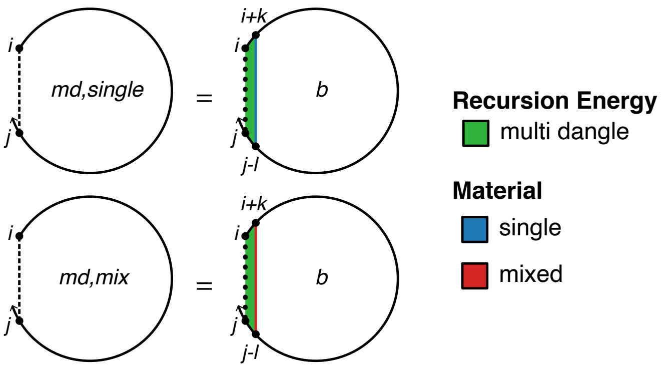

$$R^{md,\text{single}}_{\text{INTRA}}(i,j,\phi) \equiv \bigoplus_{\substack{k\in\{0,1\}\\ l\in\{0,1\}}} \begin{cases} C_{\text{stack}}, & \mu_{i+k}=\mu_{j-l}=\mu_j=\mu_i \\ \mathbb{0}, & \text{otherwise} \end{cases}$$

$$R^{md,\text{mix}}_{\text{INTRA}}(i,j,\phi) \equiv \bigoplus_{\substack{k\in\{0,1\}\\ l\in\{0,1\}}} \begin{cases} \mathbb{0}, & \mu_{i+k}=\mu_{j-l}=\mu_j=\mu_i \\ C_{\text{stack}}, & \text{otherwise} \end{cases}$$

$$\text{where } C_{\text{stack}} \equiv \begin{cases} Q^b_{i+k,j-l} \otimes \\ W(\Delta G^{\text{dangle}}_{i,i+k,j-l,j}(\phi) + \Delta G^{\text{multi}}_{\text{bp},\sigma} + k\Delta G^{\text{multi}}_{\text{nt},\mu_i} + l\Delta G^{\text{multi}}_{\text{nt},\mu_j} + \Delta G^{\text{terminalbp}}_{i+k,j-l}(\phi)), & (j-l)-(i+k) \geq 4 \\ \mathbb{0}, & \text{otherwise} \end{cases}$$

$$\sigma \equiv \begin{cases} \mu_{i+k}, & \mu_{i+k}=\mu_{j-l} \\ \text{mix}, & \text{otherwise} \end{cases}$$

Figure S30: $R^{md}_{\text{INTRA}}$ recursion with coaxial and dangle stacking. Top: recursion diagram. Bottom: recursion equation.

**$R^m_{\text{INTRA}}$ recursion with coaxial and dangle stacking.** The $R^b_{\text{INTRA}}(i,j,\phi)$ recursion references $Q^m$ elements that are computed using the $R^m_{\text{INTRA}}$ recursion shown in Figure S31. $R^m_{\text{INTRA}}(i,j,\phi)$ operates on a conditional ensemble for subsequence $[i,j]$ in a multiloop context where $i$ and $j$ may or may not be paired (depicted with a dashed line between $i$ and $j$ in the recursion diagram) and where there is at least one stacking state. Like other multiloop recursions in the mixed algorithm, $R^m_{\text{INTRA}}(i,j,\phi)$ is split into $R^{m,\text{single}}_{\text{INTRA}}(i,j,\phi)$ and $R^{m,\text{mix}}_{\text{INTRA}}(i,j,\phi)$ (the single and mixed recursions are indicated in the diagram with blue and red backbones, respectively):

- $R^{m,\text{single}}_{\text{INTRA}}(i,j,\phi)$: This recursion operates over the $R^m_{\text{INTRA}}(i,j,\phi)$ conditional ensemble on $[i,j]$ in a multiloop context where any terminal base pair or unpaired nucleotide is of a single material. This recursion distinguishes two cases that are combined using $\oplus$ in the recursion equation:
  - *One stacking state*: the case where there is exactly one stacking state in subsequence $[i,j]$ in a multiloop context. This stacking state starts at $d$ and ends in the interval $[d+1,j]$ (depicted by a straight dashed line between $d$ and $j$); subsequence $[i,d-1]$ does not have any stacking states (depicted by a hollow dotted line between $i$ and $d-1$). For the edge case with $d=i$, the entire region $[i,j]$ is incorporated using a $Q^{ms,\text{single}}_{i,j}$ element with term $C_{3s}$. No further material checks are needed since the $R^{ms,\text{single}}_{\text{INTRA}}(i,j,\phi)$ recursion only includes single-material contributions. For the base case where $d$ is between $i+1$ and $j-4$, then the unpaired region $[i,d-1]$ is incorporated using a $Q^{ma,\text{single}}_{i,d-1}$ element while the region $[d,j]$ is incorporated using $Q^{ms,\text{single}}_{d,j}$. The resulting contribution $Q^{ma,\text{single}}_{i,d-1} \otimes Q^{ms,\text{single}}_{d,j}$ of term $C_{1s}$ produces a single-material multiloop only if the material matches for both elements, which is checked with the conditional $\mu_j = \mu_i$. If the conditional is satisfied, the contribution is added to $R^{m,\text{single}}_{\text{INTRA}}(i,j,\phi)$ using term $C_{0s}$. Otherwise, if the conditional fails, the contribution is added to $R^{m,\text{mix}}_{\text{INTRA}}(i,j,\phi)$ using term $C_{0m}$. In the conditional, indices $i$ and $j$ are used to check the material since they are constant over vector $\overline{d}$ and apply to all $d-1$ and $d$ in $\overline{d}$.

- ◦ *More than one stacking state*: the case where there are two or more stacking states in subsequence $[i, j]$ in a multiloop context. The $3'$-most stacking state starts at $e + 1$ and ends in the interval $[e + 2, j]$ (depicted by a straight dashed line between $e + 1$ and $j$); the contribution of subsequence $[e + 1, j]$ is incorporated by element $Q^{ms,\text{single}}_{e+1,j}$. There are one or more additional stacking states in the interval $[i, e]$ (the straight dashed line denotes that $i$ and $e$ may or may not be paired); the contribution of subsequence $[i, e]$ is incorporated by element $Q^{m,\text{single}}_{i,e}$. The resulting contribution $Q^{ma,\text{single}}_{i,e} \otimes Q^{ms,\text{single}}_{e+1,j}$ of term $C_{2s}$ produces a single-material multiloop only if the material matches for both elements, which is checked with the conditional $\mu_j = \mu_i$. If the conditional is satisfied, the contribution is added to $R^{m,\text{single}}_{\text{INTRA}}(i, j, \phi)$ using term $C_{0s}$. Otherwise, if the conditional fails, the contribution is added to $R^{m,\text{mix}}_{\text{INTRA}}(i, j, \phi)$ using term $C_{0m}$. In the conditional, indices $i$ and $j$ are used to check the material since they are constant over vector $\overline{e}$ and apply to all $e$ and $e + 1$ in $\overline{e}$.

- $R^{m,\text{mix}}_{\text{INTRA}}(i, j, \phi)$: This recursion operates over the $R^{m}_{\text{INTRA}}(i, j, \phi)$ conditional ensemble on $[i, j]$ in a multiloop context where the terminal base pairs and unpaired nucleotides overall contain a mixture of materials. This recursion distinguishes two cases that are combined using $\oplus$ in the recursion equation:
  - ◦ *One stacking state*: the case where there is exactly one stacking state in subsequence $[i, j]$ in a multiloop context. This stacking state starts at $d$ and ends in the interval $[d + 1, j]$ (depicted by a straight dashed line between $d$ and $j$); subsequence $[i, d - 1]$ does not have any stacking states (depicted by a hollow dotted line between $i$ and $d - 1$). For the edge case with $d = i$, the entire region $[i, j]$ is incorporated using a $Q^{ms,\text{mix}}_{i,j}$ element with term $C_{3m}$. No further material checks are needed since the $R^{ms,\text{mix}}_{\text{INTRA}}(i, j, \phi)$ recursion only includes mixed material contributions. For the base case where $d$ is between $i + 1$ and $j - 4$, then the unpaired region $[i, d - 1]$ is incorporated using a $Q^{ma}_{i,d-1}$ element while the region $[d, j]$ is incorporated using a $Q^{ms}_{d,j}$ element. Because $Q^{ma}_{i,d-1}$ and $Q^{ms}_{d,j}$ are each split into single- and mixed-material recursions, there are 4 possible combinations when multiplying them. The three combinations containing at least one mixed element ($Q^{ma,\text{single}}_{i,d-1} \otimes Q^{ms,\text{mix}}_{d,j}$, $Q^{ma,\text{mix}}_{i,d-1} \otimes Q^{ms,\text{single}}_{d,j}$, $Q^{ma,\text{mix}}_{i,d-1} \otimes Q^{ms,\text{mix}}_{d,j}$) must represent mixed contributions, and are included in $R^{m,\text{mix}}_{\text{INTRA}}(i, j, \phi)$ using term $C_{1m}$. As noted above, the remaining combination, $Q^{ma,\text{single}}_{i,d-1} \otimes Q^{ms,\text{single}}_{d,j}$, calculated in term $C_{1s}$ represents a mixed contribution only if $\mu_j \neq \mu_i$, and is incorporated in $R^{m,\text{mix}}_{\text{INTRA}}(i, j, \phi)$ using term $C_{0m}$.
  - ◦ *More than one stacking state*: the case where there are two or more stacking states in subsequence $[i, j]$ in a multiloop context. The $3'$-most stacking state starts at $e+1$ and ends in the interval $[e+2, j]$ (depicted by a straight dashed line between $e + 1$ and $j$); the contribution of subsequence $[e + 1, j]$ is incorporated by element $Q^{ms}_{e+1,j}$. There are one or more additional stacking states in the interval $[i, e]$ (the straight dashed line denotes that $i$ and $e$ may or may not be paired); the contribution of subsequence $[i, e]$ is incorporated by element $Q^{m}_{i,e}$. Because $Q^{m}_{i,e}$ and $Q^{ms}_{e+1,j}$ are each split into single- and mixed-material recursions, there are 4 possible combinations when multiplying them. The three combinations containing at least one mixed element ($Q^{m,\text{single}}_{i,e} \otimes Q^{ms,\text{mix}}_{e+1,j}$, $Q^{m,\text{mix}}_{i,e} \otimes Q^{ms,\text{single}}_{e+1,j}$, $Q^{m,\text{mix}}_{i,e} \otimes Q^{ms,\text{mix}}_{e+1,j}$) must represent overall mixed contributions, and are included in $R^{m,\text{mix}}_{\text{INTRA}}(i, j, \phi)$ using term $C_{2m}$. As noted above, the remaining combination, $Q^{m,\text{single}}_{i,e} \otimes Q^{ms,\text{single}}_{e+1,j}$, calculated in term $C_{2s}$ represents a mixed contribution only if $\mu_j \neq \mu_i$, and is incorporated in $R^{m,\text{mix}}_{\text{INTRA}}(i, j, \phi)$ using term $C_{0m}$.

The shading does not represent any recursion energies as all multiloop contributions are handled by other recursions: 1) there are no terminal base pairs in a multiloop context explicitly defined in this case, 2) there are no unpaired bases in a multiloop context explicitly defined in this case. The index limits in the recursion equation reflect the steric requirement that there be at least 3 intervening bases between $d$ and $j$ (because the $R^{ms}$ incorporates a minimum of one terminal base pair in subsequence $[d : j]$) and the steric requirement that there be at least 3 intervening bases between $i$ and $e$ and at least 3 intervening bases between $e + 1$ and $j$.

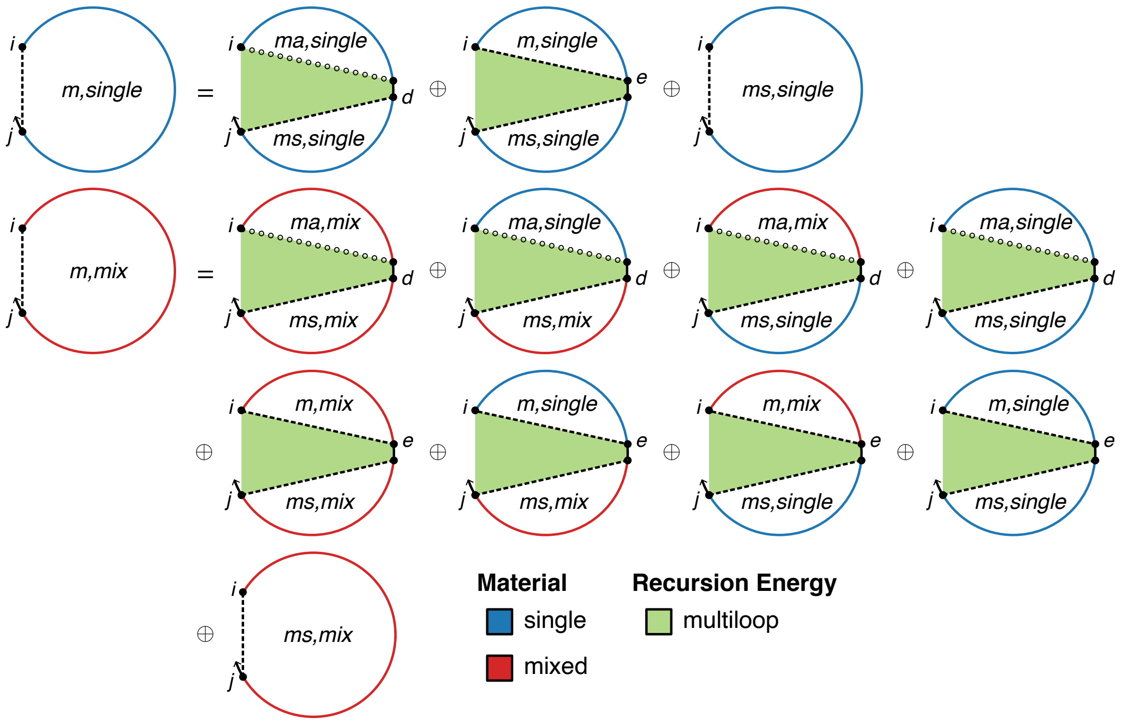

$$R^{m,\text{single}}_{\text{INTRA}}(i,j,\phi) \equiv C_{0s} \oplus C_{3s}$$

$$C_{0s} \equiv \begin{cases} C_{1s} \oplus C_{2s}, & \mu_j = \mu_i \\ \mathbb{0}, & \text{otherwise} \end{cases}$$

$$C_{1s} \equiv \begin{cases} \text{DOT}\left(Q^{ma,\text{single}}_{i,\bar{d}-1}, Q^{ms,\text{single}}_{\bar{d},j}\right), & j-i \geq 5 \\ \mathbb{0}, & \text{otherwise} \end{cases}$$

$$C_{2s} \equiv \begin{cases} \text{DOT}\left(Q^{m,\text{single}}_{i,\bar{e}}, Q^{ms,\text{single}}_{\bar{e}+1,j}\right), & j-i \geq 9 \\ \mathbb{0}, & \text{otherwise} \end{cases}$$

$$C_{3s} \equiv \begin{cases} Q^{ms,\text{single}}_{i,j}, & j-i \geq 4 \\ \mathbb{0}, & \text{otherwise} \end{cases}$$

$$R^{m,\text{mix}}_{\text{INTRA}}(i,j,\phi) \equiv C_{0m} \oplus C_{1m} \oplus C_{2m} \oplus C_{3m}$$

$$C_{0m} \equiv \begin{cases} \mathbb{0}, & \mu_j = \mu_i \\ C_{1s} \oplus C_{2s}, & \text{otherwise} \end{cases}$$

$$C_{1m} \equiv \begin{cases} \text{DOT}\left(Q^{ma,\text{mix}}_{i,\bar{d}-1}, Q^{ms,\text{mix}}_{\bar{d},j}\right) \oplus \text{DOT}\left(Q^{ma,\text{single}}_{i,\bar{d}-1}, Q^{ms,\text{mix}}_{\bar{d},j}\right) \oplus \text{DOT}\left(Q^{ma,\text{mix}}_{i,\bar{d}-1}, Q^{ms,\text{single}}_{\bar{d},j}\right), & j-i \geq 5 \\ \mathbb{0}, & \text{otherwise} \end{cases}$$

$$C_{2m} \equiv \begin{cases} \text{DOT}\left(Q^{m,\text{mix}}_{i,\bar{e}}, Q^{ms,\text{mix}}_{\bar{e}+1,j}\right) \oplus \text{DOT}\left(Q^{m,\text{single}}_{i,\bar{e}}, Q^{ms,\text{mix}}_{\bar{e}+1,j}\right) \oplus \text{DOT}\left(Q^{m,\text{mix}}_{i,\bar{e}}, Q^{ms,\text{single}}_{\bar{e}+1,j}\right), & j-i \geq 9 \\ \mathbb{0}, & \text{otherwise} \end{cases}$$

$$C_{3m} \equiv \begin{cases} Q^{ms,\text{mix}}_{i,j}, & j-i \geq 4 \\ \mathbb{0}, & \text{otherwise} \end{cases}$$

where $\bar{d} \equiv [i+1 : j-4], \quad \bar{e} \equiv [i+4 : j-5]$

Figure S31: $R^{m}_{\text{INTRA}}$ recursion with coaxial and dangle stacking. Top: recursion diagram. Bottom: recursion equation.

##### S6.5.3 Interstrand dynamic programming recursions with coaxial and dangle stacking

Here, we consider recursions for calculating the entries in a rectangular interstrand block with coaxial and dangle stacking. By definition, interstrand blocks involve 2 or more strands, and hence one or more nicks between strands. For a given interstrand block, $\eta$ stores an array of nick indices between strands within the block, with each nick denoted by the index of the nucleotide following the nick. If $m \equiv \textsc{First}(\eta)$ and $n \equiv \textsc{Last}(\eta)$, then for subsequence $[i, j]$ corresponding to element $i, j$ in the interstrand block, we have by definition $i < m$ (nucleotide $i$ is on the first strand in the block) and $j \geq n$ (nucleotide $j$ is on the last strand in the block).

**$R^{\varnothing}_{\text{INTER}}$ recursion with coaxial and dangle stacking.** We begin with the $R^{\varnothing}_{\text{INTER}}(i, j, \phi)$ recursion shown in Figure S32, which does not change in the mixed-material setting and is as previously described.[1] $R^{\varnothing}_{\text{INTER}}(i, j, \phi)$ operates on the unconstrained ensemble for subsequence $[i, j]$ with $i$ and $j$ on different strands in an exterior loop context where $i$ and $j$ may or may not be paired (depicted with a dashed line between $i$ and $j$ in the recursion diagram). Unlike $R^{\varnothing}_{\text{INTRA}}(i, j, \phi)$, there is no empty case because this would correspond to a disconnected structure (which is not in the ensemble) due to the presence of one or more nicks between $i$ and $j$. Hence, the only case is *at least one stacking state:* the non-empty case in an exterior loop context where there is at least one stacking state (i.e., two adjacent terminal base pairs coaxially stacked or zero, one, or two unpaired nucleotides dangle stacking on an adjacent terminal base pair) in subsequence $[i, j]$. The $3'$-most stacking state begins at $d + 1$ and ends in the interval $[d + 2, j]$ (depicted using a dashed line in the recursion diagram). The contributions for subsequence $[d + 1, j]$ are incorporated using a $Q^s_{d+1,j}$ element. Contributions for the remaining subsequence $[i, d]$ are incorporated by a $Q^{\varnothing}_{i,d}$ element. The shading denotes the recursion energy 0 corresponding to the zero reference state in an exterior loop context. The edge case where the index $d + 1 = i$ is displayed explicitly to indicate that no $Q^{\varnothing}$ element is accessed in this case.

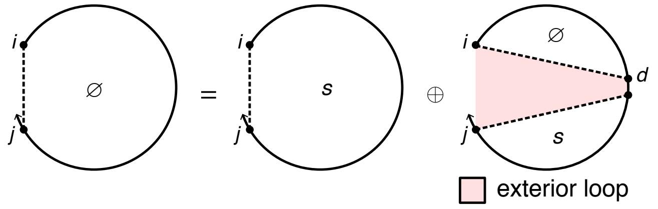

$$R^{\varnothing}_{\text{INTER}}(i, j, \phi) \equiv Q^s_{i,j} \oplus \bigoplus_{\overline{d} \in \textsc{Valid}(i, \max(j-4, n), \eta)} \textsc{dot}\left(Q^{\varnothing}_{i,\overline{d}}, Q^s_{\overline{d}+1,j}\right)$$

where $n = \textsc{Last}(\eta)$

Figure S32: $R^{\varnothing}_{\text{INTER}}$ recursion with coaxial and dangle stacking. Top: recursion diagram. Bottom: recursion equation.

Because there are nicks involved in calculating the elements of interstrand blocks, care must be taken to ensure that no disconnected secondary structures are incorporated in the complex ensemble. For a given interstrand block with nick indices $\eta$, the function $\textsc{Valid}$ returns the set of valid vectorization ranges $\{\overline{d}_1, \overline{d}_2, \dots\}$, such that for each valid vectorization range, $d$ and $d + 1$ are on the same strand (i.e., such that $d$ and $d + 1$ do not take on values that would place a nick between them). As is evident from the recursion diagram of Figure S32, if $d$ and $d + 1$ were to take on values that placed a nick between them, a disconnected structure would result. There is at most one valid vectorization range per strand, and there may be none for a strand or subsequence that is too short. For each valid vectorization range $\overline{d}$, the resulting dot product

$$\textsc{dot}\left(Q^{\varnothing}_{i,\overline{d}}, Q^s_{\overline{d}+1,j}\right) \tag{S162}$$

is between the range $\overline{d}$ of row $i$ (depicted as brown elements in Figure S4c) and the range $\overline{d} + 1$ of column $j$ (gray elements), yielding element $i, j$ (purple element). Note that Figure S4c depicts two valid vectorization ranges (leading to two dot products that are summed to calculate the purple element); the gap of one element between the two vectorization ranges corresponds to exclusion of the value $d = 3$ which would have placed a nick between nucleotides $d$ and $d + 1$ (note that $\eta = 4$ for this interstrand block).

Note that for calculating element $i, j$, the subsequence submitted to $\textsc{Valid}$ ranges from $i$ to $\max(j - 4, n)$, where $n \equiv \textsc{Last}(\eta)$. This yields two cases:

- If $\max(j-4, n) = j-4$: there is no nick between nucleotide $j-4$ and $j$ (since $n \equiv \textsc{Last}(\eta) < j-4$), so there must be at least 3 intervening bases between $d+1$ and $j$ because steric effects prevent a hairpin loop with fewer than 3 unpaired nucleotides. In this case, each incorporated element $Q^s_{d+1,j}$ results from an $R^s_{\text{INTRA}}(d+1, j, \phi)$ recursion for an intrastrand block.

- If $\max(j-4, n) = n$: there is a nick between nucleotide $j-4$ and $j$ (since $n \geq j-4$), so $d+1$ can be as large as $n-1$ and still pair to any nucleotide in subsequence $[n, j]$. In this case, each incorporated element $Q^s_{d+1,j}$ results from an $R^s_{\text{INTER}}(d+1, j, \phi)$ recursion for an interstrand block.

**$R^s_{\text{INTER}}$ recursion with coaxial and dangle stacking.** The recursion $R^s_{\text{INTER}}(i, j, \phi)$, shown in Figure S33, does not change in the mixed-material setting and is as previously described.[1] The $R^{\varnothing}_{\text{INTER}}(i, j, \phi)$ recursion references $Q^s_{d+1,j}$ elements that are computed using either the $R^s_{\text{INTRA}}$ recursion of Figure S23 (if $d+1$ and $j$ are on the same strand) or the $R^s_{\text{INTER}}$ recursion of Figure S33 (if $d+1$ and $j$ are on different strands). Recursion $R^s_{\text{INTER}}(i, j, \phi)$ operates on a conditional ensemble for subsequence $[i, j]$ with $i$ and $j$ on different strands in an exterior loop context containing one stacking state starting at $i$ and ending in the interval $[i+1, j]$ (depicted as a dashed line between $i$ and $j$). The contribution for the stacking state in subsequence $[i, d]$ is incorporated using a $Q^{cd}_{i,d}$ element. Shading denotes no recursion energy as stacking energies and terminal base pair penalties are handled in $R^{cd}$. The index $d$ must always be on the last strand (i.e., $d \geq \textsc{Last}(\eta)$) to ensure there are no strand breaks in the subsequence $[d, j]$, which would correspond to a disconnected structure. Note that the $R^s$ recursion serves as an efficiency wrapper of the $R^{cd}$ recursion (here, representing the 3′-most stacking state in an exterior loop context) to reduce the time complexity of the $R^{\varnothing}$ recursion from $O(N^4)$ to $O(N^3)$. This time complexity reduction is achieved by defining the 3′-most stacking state using $R^{cd}$ within the $R^s$ efficiency wrapper rather than directly using the $R^{cd}$ recursion within the $R^{\varnothing}$ recursion, so as to avoid introducing a fourth independent index into the $R^{\varnothing}$ recursion.

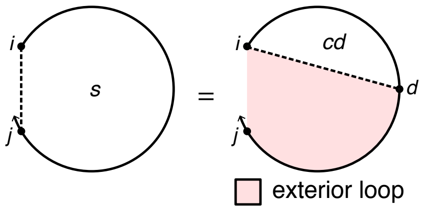

$$R^s_{\text{INTER}}(i, j, \phi) \equiv \text{DOT}\left(Q^{cd}_{i,\overline{d}}\right)$$

$$\text{where } \overline{d} \equiv [\textsc{Last}(\eta) : j]$$

Figure S33: $R^s_{\text{INTER}}$ recursion with coaxial and dangle stacking. Top: recursion diagram. Bottom: recursion equation.

**$R^{cd}_{\text{INTER}}$ recursion with coaxial and dangle stacking.** The recursion $R^{cd}_{\text{INTER}}(i, j, \phi)$, shown in Figure S34, does not change in the mixed-material setting and is as previously described.[1] The $R^s_{\text{INTER}}(i, j, \phi)$ recursion references $Q^{cd}$ elements that are computed using either the $R^{cd}_{\text{INTRA}}$ recursion of Figure S24 (if $i$ and $d$ are on the same strand) or the $R^{cd}_{\text{INTER}}$ recursion of Figure S34 (if $i$ and $d$ are on different strands). Recursion $R^{cd}_{\text{INTER}}(i, j, \phi)$ treats a single stacking state in an exterior loop context, corresponding to either of two cases that are combined using $\oplus$ in the recursion equation:

- *Coaxial stacking state:* two adjacent terminal base pairs ($i \cdot d$ and $d+1 \cdot j$) coaxially stack on each other. The contributions of subsequences $[i, d]$ and $[d+1, j]$ are incorporated using $Q^b_{i,d}$ and $Q^b_{d+1,j}$ elements. Shading corresponds to two kinds of recursion energy: 1) the sequence-dependent penalties for two terminal base pairs in an exterior loop context, $\Delta G^{\text{terminalbp}}_{i,d}(\phi)$ and $\Delta G^{\text{terminalbp}}_{d+1,j}(\phi)$ (dependent on the sequence of base pairs $i \cdot d$ and $d+1 \cdot j$), 2) the sequence-dependent coaxial stacking free energy $\Delta G^{\text{coax}}_{i,d,j}(\phi)$ (dependent on the sequences of base pairs $i \cdot d$ and $d+1 \cdot j$). Note that $\Delta G^{\text{coax}}_{i,d,j}(\phi)$ requires only 3 indices because $d+1$ is implied by $d$. To ensure that disconnected structures are excluded from the ensemble, the function $\textsc{Valid}$ returns the set of valid vectorization ranges for which nucleotides $d$ and $d+1$ are on the same strand (i.e., such that $d$ and $d+1$ do not take on values that would place a nick between them).

- *Dangle stacking state:* zero, one, or two unpaired nucleotides dangle stack on an adjacent terminal base pair ($i+k \cdot j-l$). The recursion diagram summarizes four dangle stacking states (depicted as a dotted line between $i$ and $j$) corresponding to no dangles, 5′ dangle, 3′ dangle, or terminal mismatch (see Figure S21 for details). The contribution of subsequence $[i+k, j-l]$ is incorporated using $Q^b_{i+k,j-l}$ element. Shading corresponds to two recursion energies: 1) the sequence-dependent penalty for a terminal base pair in an exterior loop context, $\Delta G^{\text{terminalbp}}_{i+k,j-l}(\phi)$ (dependent on the sequence of base pair $i+k \cdot j-l$), 2) the sequence-dependent dangle stacking free energy $\Delta G^{\text{dangle}}_{i,i+k,j-l,j}(\phi)$ which takes on one of four values corresponding to the four dangle stacking states (see Figure S21). To ensure that disconnected structures are excluded from the ensemble, the index limits in the recursion equation prevent a nick between a dangling nucleotide and the base pair on which it stacks (i.e., no nick between $i$ and $i+k$, and no nick between $j$ and $j-l$).

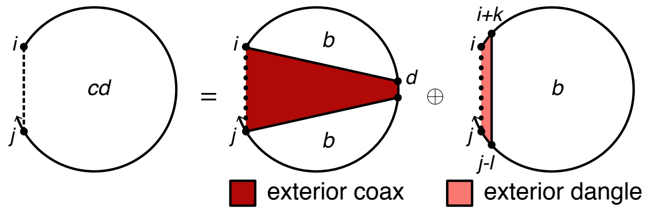

$$R^{cd}_{\text{INTER}}(i,j,\phi) \equiv \bigoplus_{\overline{d}\in\text{VALID}(i,j,\eta)} \text{DOT}\left(Q^b_{i,\overline{d}}, Q^b_{\overline{d}+1,j}, W(\Delta G^{\text{coax}}_{i,\overline{d},j}(\phi) + \Delta G^{\text{terminalbp}}_{i,\overline{d}}(\phi) + \Delta G^{\text{terminalbp}}_{\overline{d}+1,j}(\phi))\right)$$

$$\oplus \bigoplus_{\substack{k\in\{0,1\}\\ l\in\{0,1\}}} \begin{cases} Q^b_{i+k,j-l} \otimes W(\Delta G^{\text{dangle}}_{i,i+k,j-l,j}(\phi) + \Delta G^{\text{terminalbp}}_{i+k,j-l}(\phi)), & i+k<m \text{ and } j-l\geq n \\ \mathbb{0}, & \text{otherwise} \end{cases}$$

$$\text{where } m = \text{FIRST}(\eta)$$
$$n = \text{LAST}(\eta)$$

Figure S34: $R^{cd}_{\text{INTER}}$ recursion with coaxial and dangle stacking. Top: recursion diagram. Bottom: recursion equation.

**$R^b_{\text{INTER}}$ recursion with coaxial and dangle stacking.** The $R^{cd}_{\text{INTER}}(i,j,\phi)$ recursion references $Q^b_{i,d}$ elements that are computed using either the $R^b_{\text{INTRA}}$ recursion of Figure S25 (if $i$ and $d$ are on the same strand) or the $R^s_{\text{INTER}}$ recursion of Figure S35 (if $i$ and $d$ are on different strands). $R^b_{\text{INTER}}(i,j,\phi)$ operates on a conditional ensemble for subsequence $[i,j]$ with $i$ and $j$ on different strands and base paired to each other (depicted with a solid line between $i$ and $j$). The function COMPLEMENTARY$(\phi_i,\phi_j)$ checks if bases $\phi_i$ and $\phi_j$ are complementary (Watson–Crick or wobble pair) without regard to whether $i$ and $j$ are sufficiently separated along the strand to be able to pair sterically. In the mixed-material setting, the appropriate single- or mixed-material parameters must be applied based on the material composition of each loop. The recursion distinguishes five cases that are combined using $\oplus$ in the recursion equation:

- *Exterior loop with coaxial stacking on terminal base pair $j \cdot i$:* This case describes the exterior loop with two or more terminal base pairs and coaxial stacking on terminal base pair $j \cdot i$. The exterior loop treatment does not change in the mixed-material setting and is as previously described:[1]
  - *Base cases.* The base cases correspond to the recursion diagrams on the first row of Figure S35 and are treated using term $C_1$ in the subroutine MULTICOAXINTER (recursion equation S165). The diagram on the left treats the case where terminal base pair $j \cdot i$ forms a coaxial stack with adjacent terminal base pair $d+1 \cdot j-1$, depicted as a dotted straight line between $i$ and $d+1$. The contribution of subsequence $[i+1,d]$ is incorporated by element $Q^n_{i+1,d}$. The pale pink shading corresponds to the sequence-dependent penalties for two terminal base pairs in an exterior loop context, $\Delta G^{\text{terminalbp}}_{d+1,j-1}(\phi)$ and $\Delta G^{\text{terminalbp}}_{j,i}(\phi)$ (note that the indices are ordered $j$ then $i$ to reflect $5'$ to $3'$ from the perspective of the exterior loop). $\Delta G^{\text{terminalbp}}_{d+1,j-1}(\phi)$ and $\Delta G^{\text{terminalbp}}_{j,i}(\phi)$ are determined by lookup based on the sequence and material of the two nucleotides participating in each terminal base pair. The dark pink shading corresponds to the sequence-dependent coaxial stacking recursion energy $\Delta G^{\text{coax}}_{d+1,j-1,i}(\phi)$ (dependent on the sequences of base pairs $d+1 \cdot j-1$ and $j \cdot i$). Note that $\Delta G^{\text{coax}}_{d+1,j-1,i}(\phi)$ requires only 3 indices because $j$ is implied by $j-1$. $\Delta G^{\text{coax}}_{d+1,j-1,i}(\phi)$ is determined by lookup based on the sequence and material of nucleotides $d+1$, $j-1$, $j$ and $i$. The recursion on the right treats the analogous case where terminal base pair $j \cdot i$ forms a coaxial stack with adjacent terminal base pair $i+1 \cdot d$. To ensure that disconnected structures are excluded from the ensemble, the function VALID returns the set of valid vectorization ranges for which nucleotides $d$ and $d+1$ are on the same strand (i.e., such that $d$ and $d+1$ do not take on values that would place a nick between them).

- *Edge cases.* In the base case, the $Q^n$ element incorporates a nick with a neighboring $Q^\varnothing$ element on either side. The edge cases diagrammed in rows 2 to 4 of Figure S35 correspond to states where either: one of the neighboring $Q^\varnothing$ elements is omitted because the nick is adjacent to one of the two coaxially-stacking base pairs (row 2 corresponding to term $C_2$ and row 3 corresponding to terms $C_3$ and $C_4$), or where both of the neighboring $Q^\varnothing$ elements are omitted because the nick is adjacent to both of the coaxially stacking base pairs (row 4 corresponding to terms $C_5$ and $C_6$). To ensure that disconnected structures are excluded from the ensemble for terms $C_3$ and $C_4$, the function VALID returns the set of valid vectorization ranges for which nucleotides $d$ and $d+1$ are on the same strand (i.e., such that $d$ and $d+1$ do not take on values that would place a nick between them).

- *Exterior loop with a dangle stacking state involving terminal base pair $j \cdot i$:* This case describes the exterior loop with one or more terminal base pairs and a dangle stacking state involving terminal base pair $j \cdot i$. The exterior loop treatment does not change in the mixed-material setting and is as previously described:[1]

  - *Base case.* The base case corresponds to the leftmost recursion diagram in row 5 of Figure S35 and is treated by term $C_1$ in the subroutine MULTIDANGLEINTER (recursion equation S160). The contribution of subsequence $[i+k+1, j-l-1]$ is incorporated by element $Q^n_{i+k+1,j-l-1}$. The pale pink shading corresponds to the sequence-dependent penalty for a terminal base pair in an exterior loop context, $\Delta G^{\text{terminalbp}}_{j,i}(\phi)$ (note that the indices are ordered $j$ then $i$ to reflect $5'$ to $3'$ from the perspective of the multiloop). $\Delta G^{\text{terminalbp}}_{j,i}(\phi)$ is determined by lookup based on the sequence and material of nucleotides $j$ and $i$. The medium pink shading corresponds to the sequence-dependent dangle stacking energy $\Delta G^{\text{dangle}}_{j-l,j,i,i+k}(\phi)$. Note that $k, l \in \{0, 1\}$ determine whether unpaired nucleotides dangle stack on the adjacent base pair $j \cdot i$. The situation is analogous to that in $R^{cd}_{\text{INTER}}$ (as detailed in Figure S21) with the only difference being that in $R^{cd}_{\text{INTER}}$, $i+k$ and $j-l$ index the paired bases and here $i+k$ and $j-l$ index the unpaired bases. $\Delta G^{\text{dangle}}_{j-l,j,i,i+k}(\phi)$ is determined by lookup based on the sequence and material of nucleotides $j-l$, $j$, $i$, $i+k$.
  - *Edge cases.* In the base case, the $Q^n$ element incorporates a nick with a neighboring $Q^\varnothing$ element on either side. The edge cases in diagrams 2 to 4 of row 5 of Figure S35 correspond to states where either: one of the neighboring $Q^\varnothing$ elements is omitted because the nick is adjacent to the dangle stacking state on one side (diagrams 2 and 3 corresponding to term $C_2$), or where both of the neighboring $Q^\varnothing$ elements are omitted because the nick is adjacent to the dangle stacking state on both sides (diagram 4 corresponding to term $C_3$).

- *Interior loop:* the interior loop closed by the two terminal base pairs $i \cdot j$ and $d \cdot e$ (depicted by straight solid lines). Calculation of the interior loop contributions using an $O(N^4)$ or $O(N^3)$ version of the INTERIORINTER recursion is described in Section S6.4. This treatment of interior loops is the same as for the non-stacking recursions.

- *Multiloop with coaxial stacking on terminal base pair $j \cdot i$:* the multiloop closed by three or more terminal base pairs with coaxial stacking on base pair $j \cdot i$. This case corresponds to the eight recursion diagrams on the seventh and eighth rows of Figure S35 and is treated by the subroutine MULTICOAXINTER (recursion equation S165). The recursions in the seventh row treat the case where the terminal base pair $j \cdot i$ forms a coaxial stack with adjacent terminal base pair $d+1 \cdot j-1$, depicted as a dotted straight line between $i$ and $d+1$. The contribution of subsequence $[i+1, d]$ is incorporated by element $Q^b_{i+1,d}$. The contributions of one or more remaining stacking states in subsequence $[i+1, d]$ are incorporated by element $Q^m_{i+1,d}$. The recursions in the eighth row treat the analogous case where the terminal base pair $j \cdot i$ forms a coaxial stack with adjacent terminal base pair $i+1 \cdot d$; for simplicity, here we focus on describing the recursions in the seventh row:

  - *Single- and mixed-material recursions.* Multiloop contributions must be kept separate by material in $R^b_{\text{INTRA}}(i, j, \phi)$, so that the appropriate $\Delta G^{\text{multi}}_{\text{init}}$, $\Delta G^{\text{multi}}_{\text{bp}}$, and $\Delta G^{\text{terminalbp}}_{j,i}(\phi)$ parameters are applied depending on the material of the multiloop. Contributions for single-material and mixed-material multiloops are indicated in the recursion equation as $M_{\text{single}}$ and $M_{\text{mix}}$, respectively. To use the correct physical model for each multiloop as recursion elements are combined across multiple recursion levels, lower level recursions must be calculated separately for the "single" and "mixed" scenarios before they are combined at the $R^b_{\text{INTRA}}(i, j, \phi)$ level. $Q^m_{i+1,d}$ is split into $Q^{m,\text{single}}_{i+1,d}$ and $Q^{m,\text{mix}}_{i+1,d}$. "Single" elements (depicted with a blue backbone in recursion diagrams) only track energy contributions from single-material loop regions, while "mix" elements (depicted with a red backbone in recursion diagrams) only track contributions from mixed-material loop regions. The contribution for the subsequence $[d+1, j-1]$ enclosed by base pair $d+1 \cdot j-1$ is incorporated using a $Q^b_{d+1,j-1}$ element, representing either a single-material or mixed-material terminal base pair depending on whether $\mu_{j-1} = \mu_{d+1}$ (single-material; denoted by a blue base pair in the recursion diagram) or $\mu_{j-1} \neq \mu_{d+1}$ (mixed-material; denoted by a red base pair in the recursion diagram). Because

$Q^m_{i+1,d}$ is split into single-material and mixed-material recursions and $Q^b_{d+1,j-1}$ also has both single- and mixed-material cases, there are four possible single/mix combinations when multiplying them. The three combinations that contain at least one mixed element must represent mixed contributions and are included in $M_{\text{mix}}$ with the term $C_{1m}$. The remaining combination in term $C_{1s}$, produces a single-material multiloop only if the material of both elements match each other and the material of the closing base pair $j \cdot i$, which is checked using the conditional, $\mu_j = \mu_{j-1} = \mu_{i+1} = \mu_i$. If the conditional is satisfied, the contribution is added to $M_{\text{single}}$ using term $C_{0s}$. Otherwise, if the conditional fails, the contribution is added to $M_{\text{mix}}$ using term $C_{0m}$. In the conditional, note that the material of the multiloop "single" element, $Q^{m,\text{single}}_{i+1,\overline{d}}$, can be determined by checking the material of the first or last nucleotide of the subsequence spanned by that element; the first and last nucleotides are each guaranteed to appear in the multiloop as either an unpaired nucleotide or as part of a terminal base pair defining the multiloop. In this instance, the material of $Q^{m,\text{single}}_{i+1,\overline{d}}$ is checked at $i+1$ since this index is constant over vector $\overline{d}$ and apply to all $d$ in $\overline{d}$. $M_{\text{single}}$ and $M_{\text{mix}}$ are combined to incorporate all single- and mixed-material multiloops that include coaxial stacking on terminal base pair $j \cdot i$. To exclude exterior loop states that are not treated by this multiloop recursion, the function VALID returns the set of valid vectorization ranges for which nucleotides $d$ and $d+1$ are on the same strand (i.e., such that $d$ and $d+1$ do not take on values that would place a nick between them).

  - *Single- and mixed-material recursion energies.* In the recursion diagrams, pale green shading corresponds to three recursion energies: 1) the penalty for formation of a multiloop $\Delta G^{\text{multi}}_{\text{init}}$, 2) the sequence-independent penalties for two terminal base pairs in a multiloop, $\Delta G^{\text{multi}}_{\text{bp}}$ (corresponding to base pairs $d+1 \cdot j-1$ and $j \cdot i$), 3) the sequence-dependent penalties for two terminal base pairs in a multiloop context, $\Delta G^{\text{terminalbp}}_{d+1,j-1}(\phi)$ and $\Delta G^{\text{terminalbp}}_{j,i}(\phi)$ (note that the indices are ordered $j$ then $i$ to reflect $5'$ to $3'$ from the perspective of the multiloop). For $M_{\text{single}}$ (single-material multiloops checked using the conditional $\mu_j = \mu_{j-1} = \mu_{i+1} = \mu_i$), the single-material multiloop initiation parameter, $\Delta G^{\text{multi}}_{\text{init},\mu_i}$, is used and the single-material terminal base pair parameter, $\Delta G^{\text{multi}}_{\text{bp},\mu_i}$, is used for each of terminal base pairs $i \cdot j$ and $d+1 \cdot j-1$. For $M_{\text{mix}}$, the mixed-material multiloop initiation parameter $\Delta G^{\text{multi}}_{\text{init,mix}}$ is used; for terminal base pair $j \cdot i$, the terminal base pair parameter, $\Delta G^{\text{multi}}_{\text{bp},\sigma}$, depends on the material of $j$ and $i$ and is checked with the $\sigma$ conditional ($\Delta G^{\text{multi}}_{\text{bp},\mu_i}$ is used if $\mu_j = \mu_i$, and $\Delta G^{\text{multi}}_{\text{bp,mix}}$ is used otherwise); for terminal base pair $d+1 \cdot j-1$, the terminal base pair parameter depends on the material of $d+1$ and $j-1$ ($\Delta G^{\text{multi}}_{\text{bp},\mu_{j-1}}$ is used if $\mu_{d+1} = \mu_{j-1}$, and $\Delta G^{\text{multi}}_{\text{bp,mix}}$ is used otherwise). For $M_{\text{single}}$ and $M_{\text{mix}}$, the sequence-dependent terminal base pair parameter, $\Delta G^{\text{terminalbp}}_{j,i}(\phi)$, is determined by lookup based on the sequence and material of nucleotides $j$ and $i$ (note that the indices are ordered $j$ then $i$ to reflect $5'$ to $3'$ from the perspective of the multiloop); the sequence-dependent terminal base pair parameter, $\Delta G^{\text{terminalbp}}_{d+1,j-1}(\phi)$, is determined by lookup based on the sequence and material of nucleotides $d+1$ and $j-1$. In the recursion diagrams, dark green shading corresponds to the sequence-dependent coaxial stacking recursion energy $\Delta G^{\text{coax}}_{d+1,j-1,i}(\phi)$ (dependent on the sequences of terminal base pairs $d+1 \cdot j-1$ and $j \cdot i$). Note that $\Delta G^{\text{coax}}_{d+1,j-1,i}(\phi)$ requires only 3 indices because $j$ is implied by $j-1$. $\Delta G^{\text{coax}}_{\overline{d}+1,j-1,i}(\phi)$ is determined by lookup based on the sequence and material of nucleotides $d+1$, $j-1$, $j$ and $i$.

- *Multiloop with dangle stacking on terminal base pair $j \cdot i$:* the multiloop closed by three or more terminal base pairs with dangle stacking on terminal base pair $j \cdot i$. This case corresponds to the ten recursion diagrams on the ninth, tenth, and eleventh rows of Figure S35 and is treated by the subroutine MULTIDANGLEINTER (recursion equation S166). Multiloop contributions must be kept separate by material in $R^b_{\text{INTRA}}(i,j,\phi)$, so that the appropriate $\Delta G^{\text{multi}}_{\text{init}}$, $\Delta G^{\text{multi}}_{\text{bp}}$, and $\Delta G^{\text{terminalbp}}_{j,i}(\phi)$ parameters are applied depending on the material of the multiloop. Contributions for single-material and mixed-material multiloops are indicated in the recursion equation as $M_{\text{single}}$ and $M_{\text{mix}}$, respectively. To use the correct physical model for each multiloop as recursion elements are combined across multiple recursion levels, lower level recursions ($Q^m$, $Q^{ms}$, $Q^{ma}$, $Q^{mcs}$) must be calculated separately for the "single" and "mixed" scenarios before they are combined at the $R^b_{\text{INTRA}}(i,j,\phi)$ level. "Single" elements (depicted with a blue backbone in recursion diagrams) only track energy contributions from single-material loop regions, while "mix" elements (depicted with a red backbone in recursion diagrams) only track contributions from mixed-material loop regions.

  - *Base case with two or more additional stacking states.* The recursions on the ninth row treat the case where there is a dangle stacking state involving the terminal base pair $j \cdot i$ (depicted as a dotted straight line between $i+k$ and $j-l$), a $3'$-most stacking state in subsequence $[d+1, j-l-1]$ (depicted as a dashed line between $d+1$ and $j-l-1$), and one or more stacking states in subsequence $[i+k+1, d]$ (depicted as a dashed line between $i+k+1$ and $d$). The contributions for subsequence $[d+1, j-l-1]$ are incorporated using a $Q^{ms}_{d+1,j-l-1}$ element, while the contributions for $[i+k+1, d]$ are incorporated using a $Q^m_{i+k+1,d}$ element. Because $Q^m_{i+k+1,d}$ and $Q^{ms}_{d+1,j-l-1}$ are each split into single- and mixed-material recursions, there are 4 possible combinations when multiplying them. Any combination with a mixed element produces an overall mixed multiloop ($Q^{m,\text{mix}}_{i+k+1,d} \otimes Q^{ms,\text{mix}}_{d+1,j-l-1}$, $Q^{m,\text{mix}}_{i+k+1,d} \otimes$

$Q^{ms,\text{single}}_{d+1,j-l-1}$, $Q^{m,\text{single}}_{i+k+1,d} \otimes Q^{ms,\text{mix}}_{d+1,j-l-1}$), which is added to $M_{\text{mix}}$ as term $C_{1m}$ in (S166). The remaining combination, $Q^{m,\text{single}}_{i+k+1,d} \otimes Q^{ms,\text{single}}_{d+1,j-l-1}$, in term $C_{1s}$ produces a single-material multiloop only if the material matches for both elements, the closing base pair $j \cdot i$, and any unpaired dangling nucleotide $i+k$ and $j-l$, which is checked using the conditional $\text{IsSingle}(i, i+k+1, \mu_i)$ and $\text{IsSingle}(j-l-1, j, \mu_i)$. Recall that the function $\text{IsSingle}(x, y, \mu_i)$ checks if subsequence $[x, y]$ is composed entirely of material $\mu_i$. If the conditional is satisfied, the contribution is added to $M_{\text{single}}$ using term $C_{0s}$. Otherwise, if the conditional fails, the contribution is added to $M_{\text{mix}}$ using term $C_{0m}$. In the conditional, note that the material of any multiloop single element, like $Q^{m,\text{single}}_{i+k+1,d}$ or $Q^{ms,\text{single}}_{d+1,j-l-1}$, can be known by checking the material of the first or last nucleotide of the subsequence spanned by that element; the first and last nucleotides are each guaranteed to appear in the multiloop as either an unpaired nucleotide or as part of a terminal base pair defining the multiloop. In this instance, the material of $Q^{m,\text{single}}_{i+k+1,\overline{d}}$ is checked at $i+k+1$, while the material of $Q^{ms,\text{single}}_{\overline{d}+1,j-l-1}$ is checked at $j-l-1$, since these indices are constant over vector $\overline{d}$ and apply to all $d$ and $d+1$ in $\overline{d}$. Note that $k, l \in \{0, 1\}$ determine whether unpaired nucleotides dangle stack on the adjacent terminal base pair $j \cdot i$ in a multiloop context. The situation is analogous to that in an exterior loop context with $R^{cd}_{\text{INTRA}}(i, j, \phi)$ (as detailed in Figure S21) with the only difference being that in the exterior loop context, $i+k$ and $j-l$ index the paired bases and in the multiloop context $i+k$ and $j-l$ index the unpaired bases. To exclude exterior loop states that are not treated by this multiloop recursion, the function $\text{Valid}$ returns the set of valid vectorization ranges for which nucleotides $d$ and $d+1$ are on the same strand (i.e., such that $d$ and $d+1$ do not take on values that would place a nick between them). *Single- and mixed-material recursion energies.* In the recursion diagrams, the pale green shading corresponds to four multiloop recursion energies: 1) the penalty for formation of a multiloop $\Delta G^{\text{multi}}_{\text{init}}$, 2) the sequence-independent penalty for one terminal base pair in a multiloop, $\Delta G^{\text{multi}}_{\text{bp}}$ (corresponding to base pair $j \cdot i$), 3) the penalty per unpaired nucleotide in a multiloop $\Delta G^{\text{multi}}_{\text{nt}}$ (a total of $k+l$ dangling nucleotides; as a result this term is zeroed out when $k = l = 0$), 4) the sequence-dependent penalty for a terminal base pair in a multiloop context, $\Delta G^{\text{terminalbp}}_{j,i}(\phi)$ (note that the indices are ordered $j$ then $i$ to reflect $5'$ to $3'$ from the perspective of the multiloop). For $M_{\text{single}}$, the single-material multiloop initiation parameter $\Delta G^{\text{multi}}_{\text{init},\mu_i}$ is used; the single-material terminal base pair parameter $\Delta G^{\text{multi}}_{\text{bp},\mu_i}$ is used for terminal base pair $j \cdot i$; the unpaired nucleotide parameter $\Delta G^{\text{multi}}_{\text{nt},\mu_i}$ is used for any unpaired nucleotide $i+k$ or $j-l$. For $M_{\text{mix}}$, the mixed-material multiloop initiation parameter $\Delta G^{\text{multi}}_{\text{init,mix}}$ is used; for terminal base pair $j \cdot i$, the terminal base pair parameter, $\Delta G^{\text{multi}}_{\text{bp},\sigma}$, depends on the material of $j$ and $i$ and is checked with the $\sigma$ conditional ($\Delta G^{\text{multi}}_{\text{bp},\mu_i}$ is used if $\mu_j = \mu_i$, and $\Delta G^{\text{multi}}_{\text{bp,mix}}$ is used otherwise); the unpaired nucleotide parameter $\Delta G^{\text{multi}}_{\text{nt}}$ depends on the material of each dangling nucleotide ($\Delta G^{\text{multi}}_{\text{nt},\mu_{i+k}}$ for $i+k$ and $\Delta G^{\text{multi}}_{\text{nt},\mu_{j-l}}$ for $j-l$). For $M_{\text{single}}$ and $M_{\text{mix}}$, the sequence-dependent terminal base pair parameter, $\Delta G^{\text{terminalbp}}_{j,i}(\phi)$, is determined by lookup based on the sequence and material of nucleotides $j$ and $i$ (note that the indices are ordered $j$ then $i$ to reflect $5'$ to $3'$ from the perspective of the multiloop). In the recursion diagrams, the medium green shading corresponds to the sequence-dependent dangle stacking recursion energy $\Delta G^{\text{dangle}}_{j-l,j,i,i+k}(\phi)$ (dependent on the sequence and material of the participating nucleotides).

- *Edge case with one additional coaxial stacking state.*The recursions on the tenth row treat the case where there is a dangle stacking state involving the terminal base pair $j \cdot i$ (depicted as a dotted straight line between $i+k$ and $j-l$), a single coaxial stacking state in subsequence $[e, j-l-1]$ (depicted as a half-solid/half-dashed line between $e$ and $j-l-1$), and an unpaired region in subsequence $[i+k+1, e-1]$ (depicted as a hollow dotted line between $i+k+1$ and $e-1$). The contributions for subsequence $[e, j-l-1]$ are incorporated using a $Q^{mcs}_{e,j-l-1}$ element, while the contributions for unpaired region $[i+k+1, e-1]$ are incorporated using a $Q^{ma}_{i+k+1,e-1}$ element. Because $Q^{ma}_{i+k+1,e-1}$ and $Q^{mcs}_{e,j-l-1}$ are each split into single- and mixed-material recursions, there are 4 possible combinations when multiplying them. Any combination with a mixed element produces an overall mixed multiloop ($Q^{ma,\text{mix}}_{i+k+1,e-1} \otimes Q^{mcs,\text{mix}}_{e,j-l-1}$, $Q^{ma,\text{mix}}_{i+k+1,e-1} \otimes Q^{mcs,\text{single}}_{e,j-l-1}$, $Q^{ma,\text{single}}_{i+k+1,e-1} \otimes Q^{mcs,\text{mix}}_{e,j-l-1}$), which is added to $M_{\text{mix}}$ as term $C_{2m}$ in (S166). The remaining combination, $Q^{ma,\text{single}}_{i+k+1,e-1} \otimes Q^{mcs,\text{single}}_{e,j-l-1}$, in term $C_{2s}$ produces a single-material multiloop only if the material matches for both elements, the closing base pair $j \cdot i$, and any unpaired dangling nucleotide $i+k$ and $j-l$, which is checked with the conditional $\text{IsSingle}(i, i+k+1, \mu_i)$ and $\text{IsSingle}(j-l-1, j, \mu_i)$. Recall that the function $\text{IsSingle}(x, y, \mu_i)$ checks if subsequence $[x, y]$ is composed entirely of material $\mu_i$. If the conditional is satisfied, the contribution is added to $M_{\text{single}}$ using term $C_{0s}$. Otherwise, if the conditional fails, the contribution is added to $M_{\text{mix}}$ using term $C_{0m}$. In the conditional, note that the material of $Q^{ma,\text{single}}_{i+k+1,\overline{e}-1}$ is checked at $i+k+1$, while the material of $Q^{mcs,\text{single}}_{\overline{e},j-l-1}$ is checked at $j-l-1$, since these indices are constant over vector $\overline{e}$ and apply to all $e-1$ and $e$ in $\overline{e}$. The recursions on the eleventh row treat the special case where $e = i+k+1$, in which the contributions for subsequence $[i+k+1, j-l-1]$ are incorporated using a $Q^{mcs}_{i+k+1,j-l-1}$ element, for which there are both single- and mixed-material recursions. The mixed-material contribution $Q^{mcs,\text{mix}}_{i+k+1,j-l-1}$ is added to $M_{\text{mix}}$ as term $C_{3m}$. The contribution of $Q^{mcs,\text{single}}_{i+k+1,j-l-1}$ in term $C_{3s}$ produces a single-material multiloop only if the material matches for the $Q^{mcs,\text{single}}_{i+k+1,j-l-1}$ element, the closing base pair $j \cdot i$, and any unpaired dangling nucleotide $i+k$ and $j-l$, which is checked with the conditional $\textsc{IsSingle}(i, i+k+1, \mu_i)$ and $\textsc{IsSingle}(j-l-1, j, \mu_i)$. If the conditional is satisfied, the contribution is added to $M_{\text{single}}$ using term $C_{0s}$. Otherwise, if the conditional fails, the contribution is added to $M_{\text{mix}}$ using term $C_{0m}$. Note that this edge case covers the scenario where there are exactly three terminal base pairs and the two terminal base pairs that are not $j \cdot i$ are coaxially stacked. That situation is not covered by the base case because for that recursion, a multiloop with 3 terminal base pairs would have one terminal base pair in the $Q^{ms}$ element, and one terminal base pair in the $Q^m$ element (hence, those two terminal base pairs cannot coaxially stack since they are in different recursions). Nucleotide $e$ must always be on the first strand ($e \leq m-1$) to ensure that there are no nicks in the subsequence $[i, e]$, which would lead to either a disconnected structure (which is not permitted in the complex ensemble) or an exterior loop state (which is not handled by this multiloop recursion). Also, $e \geq i+k+2$ in order to provide a minimum of one nucleotide for the $Q^{ma}_{i+k+1,e-1}$ recursion, so we must have $i+k+2 < m$ for terms $C_{2s}$ and $C_{2m}$. *Single- and mixed-material recursion energies.* In the recursion diagrams, the pale green shading corresponds to four multiloop recursion energies: 1) the penalty for formation of a multiloop $\Delta G^{\text{multi}}_{\text{init}}$, 2) the sequence-independent penalty for one terminal base pair in a multiloop, $\Delta G^{\text{multi}}_{\text{bp}}$ (corresponding to base pair $j \cdot i$), 3) the penalty per unpaired nucleotide in a multiloop $\Delta G^{\text{multi}}_{\text{nt}}$ ($k+l$ dangling nucleotides plus the unpaired nucleotides $k+l+1, \ldots, e-1$; as a result this term is zeroed out when $k = l = 0$ and $e = i+1$), 4) the sequence-dependent penalty for a terminal base pair in a multiloop context, $\Delta G^{\text{terminalbp}}_{j,i}(\phi)$ (note that the indices are ordered $j$ then $i$ to reflect $5'$ to $3'$ from the perspective of the multiloop). For $M_{\text{single}}$, the single-material multiloop initiation parameter $\Delta G^{\text{multi}}_{\text{init},\mu_i}$ is used; the single-material terminal base pair parameter $\Delta G^{\text{multi}}_{\text{bp},\mu_i}$ is used for terminal base pair $j \cdot i$; the unpaired nucleotide parameter $\Delta G^{\text{multi}}_{\text{nt},\mu_i}$ is used for any unpaired nucleotide $i+k$ or $j-l$. For $M_{\text{mix}}$, the mixed-material multiloop initiation parameter $\Delta G^{\text{multi}}_{\text{init,mix}}$ is used; for terminal base pair $j \cdot i$, the terminal base pair parameter, $\Delta G^{\text{multi}}_{\text{bp},\sigma}$, depends on the material of $j$ and $i$ and is checked with the $\sigma$ conditional ($\Delta G^{\text{multi}}_{\text{bp},\mu_i}$ is used if $\mu_j = \mu_i$, and $\Delta G^{\text{multi}}_{\text{bp,mix}}$ is used otherwise); the unpaired nucleotide parameter $\Delta G^{\text{multi}}_{\text{nt}}$ depends on the material of each dangling nucleotide ($\Delta G^{\text{multi}}_{\text{nt},\mu_{i+k}}$ for $i+k$ and $\Delta G^{\text{multi}}_{\text{nt},\mu_{j-l}}$ for $j-l$). For $M_{\text{single}}$ and $M_{\text{mix}}$, the sequence-dependent terminal base pair parameter, $\Delta G^{\text{terminalbp}}_{j,i}(\phi)$, is determined by lookup based on the sequence and material of nucleotides $j$ and $i$ (note that the indices are ordered $j$ then $i$ to reflect $5'$ to $3'$ from the perspective of the multiloop). In the recursion diagrams, the medium green shading corresponds to the sequence-dependent dangle stacking recursion energy $\Delta G^{\text{dangle}}_{j-l,j,i,i+k}(\phi)$ (dependent on the sequence and material of the participating nucleotides).

Note that unlike the $R^b_{\text{INTRA}}$ recursion of Figure S25, for $R^b_{\text{INTER}}$ there is no hairpin loop case as $i$ and $j$ are on different strands.

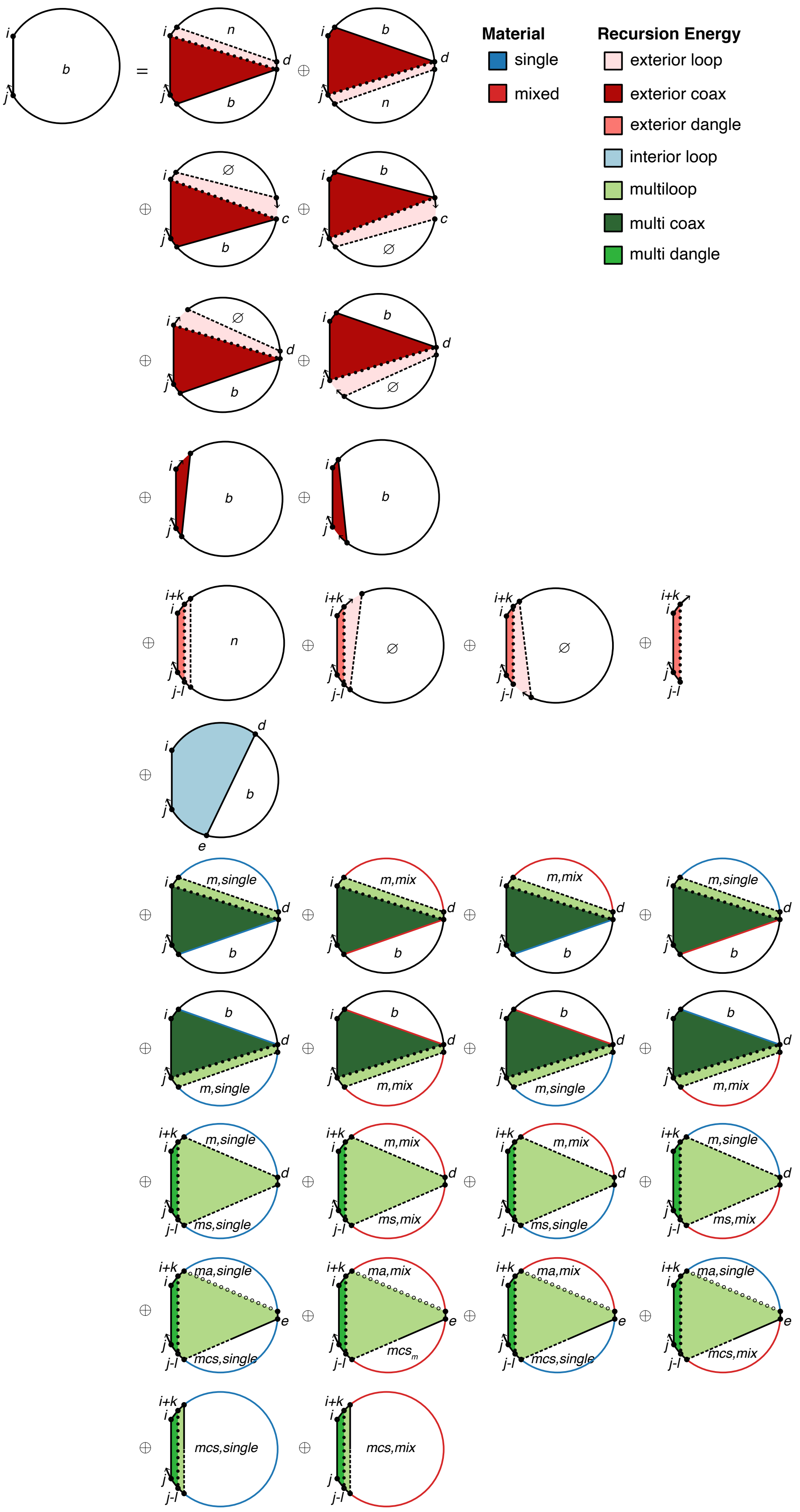
Material
single
mixed
Recursion Energy
exterior loop
exterior coax
exterior dangle
interior loop
multiloop
multi coax
multi dangle

$$R^b_{\text{INTER}}(i,j,\phi) \equiv \begin{cases} C_{\text{all}}, & \textsc{Complementary}(\phi_i,\phi_j) \\ \mathbb{0}, & \text{otherwise} \end{cases}$$

$$\begin{aligned} C_{\text{all}} \equiv\ & \textsc{ExteriorCoaxInter}(i,j,\phi,\eta) \\ & \oplus \textsc{ExteriorDangleInter}(i,j,\phi,\eta) \\ & \oplus \textsc{InteriorInter}(i,j,\phi,\eta) \\ & \oplus \textsc{MultiCoaxInter}(i,j,\phi,\eta) \\ & \oplus \textsc{MultiDangleInter}(i,j,\phi,\eta) \end{aligned}$$

Figure S35: $R^b_{\text{INTER}}$ recursion with coaxial and dangle stacking. Top: recursion diagram. Bottom: recursion equation.

$$
\begin{aligned}
\textsc{ExteriorCoaxInter}(i,j,\phi,\eta) \equiv\ & C_1 \otimes W(\Delta G^{\text{terminalbp}}_{j,i}(\phi)) \\
& \oplus \begin{cases} C_2 \otimes W(\Delta G^{\text{terminalbp}}_{j,i}(\phi)), & i+1<m \text{ and } j-1 \geq n \\ (C_3 \oplus C_5) \otimes W(\Delta G^{\text{terminalbp}}_{j,i}(\phi)), & i+1=m \text{ and } j-1 \geq n \\ (C_4 \oplus C_6) \otimes W(\Delta G^{\text{terminalbp}}_{j,i}(\phi)), & i+1<m \text{ and } j=n \\ \mathbb{0}, & \text{otherwise} \end{cases} \\
\text{where } m &= \textsc{First}(\eta) \\
n &= \textsc{Last}(\eta) \\
C_1 \equiv & \bigoplus_{\overline{d} \in \textsc{Valid}(i+1,j-1,\eta)} \Big[ \text{DOT}\left(Q^n_{i+1,\overline{d}}, Q^b_{\overline{d}+1,j-1}, W(\Delta G^{\text{coax}}_{\overline{d}+1,j-1,i}(\phi) + \Delta G^{\text{terminalbp}}_{\overline{d}+1,j-1}(\phi))\right) \\
& \oplus \text{DOT}\left(Q^b_{i+1,\overline{d}}, Q^n_{\overline{d}+1,j-1}, W(\Delta G^{\text{coax}}_{j,i,\overline{d}}(\phi) + \Delta G^{\text{terminalbp}}_{i+1,\overline{d}}(\phi))\right)\Big] \\
C_2 \equiv & \bigoplus_{c \in \eta} \Big[ Q^{\varnothing}_{i+1,c-1} \otimes Q^b_{c,j-1} \otimes W(\Delta G^{\text{coax}}_{c,j-1,i}(\phi) + \Delta G^{\text{terminalbp}}_{c,j-1}(\phi)) \\
& \oplus Q^b_{i+1,c-1} \otimes Q^{\varnothing}_{c,j-1} \otimes W(\Delta G^{\text{coax}}_{j,i,c-1}(\phi) + \Delta G^{\text{terminalbp}}_{i+1,c-1}(\phi))\Big] \\
C_3 \equiv & \bigoplus_{\overline{d} \in \textsc{Valid}(i,j-1,\eta)} \text{DOT}\left(Q^{\varnothing}_{i+1,\overline{d}}, Q^b_{\overline{d}+1,j-1}, W(\Delta G^{\text{coax}}_{\overline{d}+1,j-1,i}(\phi) + \Delta G^{\text{terminalbp}}_{\overline{d}+1,j-1}(\phi))\right) \\
C_4 \equiv & \bigoplus_{\overline{d} \in \textsc{Valid}(i+1,j,\eta)} \text{DOT}\left(Q^b_{i+1,\overline{d}}, Q^{\varnothing}_{\overline{d}+1,j-1}, W(\Delta G^{\text{coax}}_{j,i,\overline{d}}(\phi) + \Delta G^{\text{terminalbp}}_{i+1,\overline{d}}(\phi))\right) \\
C_5 \equiv\ & Q^b_{i+1,j-1} \otimes W(\Delta G^{\text{coax}}_{i+1,j-1,i}(\phi) + \Delta G^{\text{terminalbp}}_{i+1,j-1}(\phi)) \\
C_6 \equiv\ & Q^b_{i+1,j-1} \otimes W(\Delta G^{\text{coax}}_{j,i,j-1}(\phi) + \Delta G^{\text{terminalbp}}_{i+1,j-1}(\phi))
\end{aligned}
\tag{S163}
$$

$$
\begin{aligned}
\textsc{ExteriorDangleInter}(i,j,\phi,\eta) \equiv & \bigoplus_{\substack{k\in\{0,1\} \\ l\in\{0,1\}}} \begin{cases} C_1, & i+k+1<m \text{ and } j-l-1 \geq n \\ C_2, & i+k+1=m \text{ and } j-l-1 \geq n \\ C_2, & i+k+1<m \text{ and } j-l=n \\ C_3, & i+k+1=m \text{ and } j-l=n \text{ and } m=n \\ \mathbb{0}, & \text{otherwise} \end{cases} \\
\text{where } m &= \textsc{First}(\eta) \\
n &= \textsc{Last}(\eta) \\
C_1 &\equiv Q^n_{i+k+1,j-l-1} \otimes C_3 \\
C_2 &\equiv Q^{\varnothing}_{i+k+1,j-l-1} \otimes C_3 \\
C_3 &\equiv W(\Delta G^{\text{dangle}}_{j-l,j,i,i+k}(\phi) + \Delta G^{\text{terminalbp}}_{j,i}(\phi))
\end{aligned}
\tag{S164}
$$

$$\textsc{MultiCoaxInter}(i,j,\phi,\eta) \equiv \bigoplus_{\overline{d}\in \textsc{Valid}(i+1,j-1,\eta)} M_{\text{single}} \oplus M_{\text{multi}}$$

$$\begin{aligned}
\text{where } M_{\text{single}} &\equiv C_{0s} \otimes W(\Delta G^{\text{multi}}_{\text{init},\mu_i} + 2\Delta G^{\text{multi}}_{\text{bp},\mu_i} + \Delta G^{\text{terminalbp}}_{j,i}(\phi)) \\
C_{0s} &\equiv \begin{cases} C_{1s} \oplus C_{2s}, & \mu_{i+1} = \mu_{j-1} = \mu_j = \mu_i \\ \mathbb{0}, & \text{otherwise} \end{cases} \\
C_{1s} &\equiv \begin{cases} \text{DOT}\left(Q^{m,\text{single}}_{i+1,\overline{d}}, Q^{b}_{\overline{d}+1,j-1}, W(\Delta G^{\text{coax}}_{\overline{d}+1,j-1,i}(\phi) + \Delta G^{\text{terminalbp}}_{\overline{d}+1,j-1}(\phi))\right), & \mu_{d+1} = \mu_{j-1} \\ \mathbb{0}, & \text{otherwise} \end{cases} \\
C_{2s} &\equiv \begin{cases} \text{DOT}\left(Q^{b}_{i+1,\overline{d}}, Q^{m,\text{single}}_{\overline{d}+1,j-1}, W(\Delta G^{\text{coax}}_{j,i,\overline{d}}(\phi) + \Delta G^{\text{terminalbp}}_{i+1,\overline{d}}(\phi))\right), & \mu_{d} = \mu_{i+1} \\ \mathbb{0}, & \text{otherwise} \end{cases} \\
\text{and } M_{\text{mix}} &\equiv (C_{0m} \oplus C_{1m} \oplus C_{2m}) \otimes W(\Delta G^{\text{multi}}_{\text{init},\text{mix}} + \Delta G^{\text{multi}}_{\text{bp},\sigma} + \Delta G^{\text{terminalbp}}_{j,i}(\phi)) \\
C_{0m} &\equiv \begin{cases} \mathbb{0}, & \mu_j = \mu_{j-1} = \mu_{i+1} = \mu_i \\ C_{1s} \otimes W(\Delta G^{\text{multi}}_{\text{bp},\mu_{j-1}}) \oplus C_{2s} \otimes W(\Delta G^{\text{multi}}_{\text{bp},\mu_{i+1}}), & \text{otherwise} \end{cases} \\
C_{1m} &\equiv \begin{cases} \text{DOT}\left(Q^{m,\text{mix}}_{i+1,\overline{d}}, Q^{b}_{\overline{d}+1,j-1}, W(\Delta G^{\text{coax}}_{\overline{d}+1,j-1,i}(\phi) + \Delta G^{\text{terminalbp}}_{\overline{d}+1,j-1}(\phi) + \Delta G^{\text{multi}}_{\text{bp},\mu_{j-1}})\right), & \mu_{d+1} = \mu_{j-1} \\ \mathbb{0}, & \text{otherwise} \end{cases} \\
&\oplus \begin{cases} \text{DOT}\left(Q^{m,\text{single}}_{i+1,\overline{d}}, Q^{b}_{\overline{d}+1,j-1}, W(\Delta G^{\text{coax}}_{\overline{d}+1,j-1,i}(\phi) + \Delta G^{\text{terminalbp}}_{\overline{d}+1,j-1}(\phi) + \Delta G^{\text{multi}}_{\text{bp},\text{mix}})\right), & \mu_{d+1} \neq \mu_{j-1} \\ \mathbb{0}, & \text{otherwise} \end{cases} \\
&\oplus \begin{cases} \text{DOT}\left(Q^{m,\text{mix}}_{i+1,\overline{d}}, Q^{b}_{\overline{d}+1,j-1}, W(\Delta G^{\text{coax}}_{\overline{d}+1,j-1,i}(\phi) + \Delta G^{\text{terminalbp}}_{\overline{d}+1,j-1}(\phi) + \Delta G^{\text{multi}}_{\text{bp},\text{mix}})\right), & \mu_{d+1} \neq \mu_{j-1} \\ \mathbb{0}, & \text{otherwise} \end{cases} \\
C_{2m} &\equiv \begin{cases} \text{DOT}\left(Q^{b}_{i+1,\overline{d}}, Q^{m,\text{mix}}_{\overline{d}+1,j-1}, W(\Delta G^{\text{coax}}_{j,i,\overline{d}}(\phi) + \Delta G^{\text{terminalbp}}_{i+1,\overline{d}}(\phi) + \Delta G^{\text{multi}}_{\text{bp},\mu_{i+1}})\right), & \mu_{d} = \mu_{i+1} \\ \mathbb{0}, & \text{otherwise} \end{cases} \\
&\oplus \begin{cases} \text{DOT}\left(Q^{b}_{i+1,\overline{d}}, Q^{m,\text{single}}_{\overline{d}+1,j-1}, W(\Delta G^{\text{coax}}_{j,i,\overline{d}}(\phi) + \Delta G^{\text{terminalbp}}_{i+1,\overline{d}}(\phi) + \Delta G^{\text{multi}}_{\text{bp},\text{mix}})\right), & \mu_{d} \neq \mu_{i+1} \\ \mathbb{0}, & \text{otherwise} \end{cases} \\
&\oplus \begin{cases} \text{DOT}\left(Q^{b}_{i+1,\overline{d}}, Q^{m,\text{mix}}_{\overline{d}+1,j-1}, W(\Delta G^{\text{coax}}_{j,i,\overline{d}}(\phi) + \Delta G^{\text{terminalbp}}_{i+1,\overline{d}}(\phi) + \Delta G^{\text{multi}}_{\text{bp},\text{mix}})\right), & \mu_{d} \neq \mu_{i+1} \\ \mathbb{0}, & \text{otherwise} \end{cases} \\
\text{with } \sigma &\equiv \begin{cases} \mu_i, & \mu_j = \mu_i \\ \text{mix}, & \text{otherwise} \end{cases}
\end{aligned} \tag{S165}$$

$$\textsc{MultiDangleInter}(i,j,\phi,\eta) \equiv \bigoplus_{\substack{k\in\{0,1\}\\ l\in\{0,1\}}} \begin{cases} M_{\text{single}} \oplus M_{\text{mix}}, & i+k+1 < m \text{ and } j-l-1 \geq n \\ \mathbb{0}, & \text{otherwise} \end{cases}$$

$$\begin{aligned}
\text{where } M_{\text{single}} &\equiv C_{0s} \otimes W(\Delta G^{\text{dangle}}_{j-l,j,i,i+k}(\phi) + \Delta G^{\text{multi}}_{\text{init},\mu_i} + \Delta G^{\text{multi}}_{\text{bp},\mu_i} + (k+l)\Delta G^{\text{multi}}_{\text{nt},\mu_i} + \Delta G^{\text{terminalbp}}_{j,i}(\phi)) \\
C_{0s} &\equiv \begin{cases} C_{1s} \oplus C_{2s} \oplus C_{3s}, & \textsc{IsSingle}(i,i+k+1,\mu_i) \text{ and } \textsc{IsSingle}(j-l-1,j,\mu_i) \\ \mathbb{0}, & \text{otherwise} \end{cases} \\
C_{1s} &\equiv \bigoplus_{\overline{d}\in\textsc{Valid}(i+k+1,j-l-1,\eta)} \textsc{dot}\left(Q^{m,\text{single}}_{i+k+1,\overline{d}}, Q^{ms,\text{single}}_{\overline{d}+1,j-l-1}\right) \\
C_{2s} &\equiv \begin{cases} \textsc{dot}\left(Q^{ma,\text{single}}_{i+k+1,\overline{e}-1}, Q^{mcs,\text{single}}_{\overline{e},j-l-1}\right), & i+k+2 < m \\ \mathbb{0}, & \text{otherwise} \end{cases} \\
C_{3s} &\equiv Q^{mcs,\text{single}}_{i+k+1,j-l-1} \\
\text{and } M_{\text{mix}} &\equiv (C_{0m} \oplus C_{1m} \oplus C_{2m} \oplus C_{3m}) \\
&\quad \otimes W(\Delta G^{\text{dangle}}_{j-l,j,i,i+k}(\phi) + \Delta G^{\text{multi}}_{\text{init,mix}} + \Delta G^{\text{multi}}_{\text{bp},\sigma} + k\Delta G^{\text{multi}}_{\text{nt},\mu_{i+k}} + l\Delta G^{\text{multi}}_{\text{nt},\mu_{j-l}} + \Delta G^{\text{terminalbp}}_{j,i}(\phi)) \\
C_{0m} &\equiv \begin{cases} \mathbb{0}, & \textsc{IsSingle}(i,i+k+1,\mu_i) \text{ and } \textsc{IsSingle}(j-l-1,j,\mu_i) \\ C_{1s} \oplus C_{2s} \oplus C_{3s}, & \text{otherwise} \end{cases} \\
C_{1m} &\equiv \bigoplus_{\overline{d}\in\textsc{Valid}(i+k+1,j-l-1,\eta)} \left[\left(\textsc{dot}\left(Q^{m,\text{mix}}_{i+k+1,\overline{d}}, Q^{ms,\text{single}}_{\overline{d}+1,j-l-1}\right) \oplus \textsc{dot}\left(Q^{m,\text{single}}_{i+k+1,\overline{d}}, Q^{ms,\text{mix}}_{\overline{d}+1,j-l-1}\right) \oplus \textsc{dot}\left(Q^{m,\text{mix}}_{i+k+1,\overline{d}}, Q^{ms,\text{mix}}_{\overline{d}+1,j-l-1}\right)\right]\right. \\
C_{2m} &\equiv \begin{cases} \textsc{dot}\left(Q^{ma,\text{mix}}_{i+k+1,\overline{e}-1}, Q^{mcs,\text{single}}_{\overline{e},j-l-1}\right) \oplus \textsc{dot}\left(Q^{ma,\text{single}}_{i+k+1,\overline{e}-1}, Q^{mcs,\text{mix}}_{\overline{e},j-l-1}\right) & \\ \oplus\textsc{dot}\left(Q^{ma,\text{mix}}_{i+k+1,\overline{e}-1}, Q^{mcs,\text{mix}}_{\overline{e},j-l-1}\right), & i+k+2 < m \\ \mathbb{0}, & \text{otherwise} \end{cases} \\
C_{3m} &\equiv Q^{mcs,\text{mix}}_{i+k+1,j-l-1} \\
\text{with } \sigma &\equiv \begin{cases} \mu_i, & \mu_j = \mu_i \\ \text{mix}, & \text{otherwise} \end{cases} \\
m &= \textsc{First}(\eta) \\
n &= \textsc{Last}(\eta) \\
\overline{e} &\equiv [i+k+2 : m-1]
\end{aligned} \tag{S166}$$

$\boldsymbol{R^{n}_{\text{INTER}}}$ **recursion with coaxial and dangle stacking.** The recursion $R^{n}_{\text{INTER}}(i,j,\phi)$, shown in Figure S36, does not change in the mixed-material setting and is as previously described.[1] The $R^{b}_{\text{INTER}}(i,j,\phi)$ recursion references $Q^{n}$ elements that are computed using the $R^{n}_{\text{INTER}}$ recursion. $R^{n}_{\text{INTER}}(i,j,\phi)$ operates on a conditional ensemble in an exterior loop context with a nick at $c$. For each nick $c \in \eta$, the contribution of subsequence $[i,c-1]$ is incorporated by element $Q^{\varnothing}_{i,c-1}$ and the contribution of subsequence $[c,j]$ is incorporated by element $Q^{\varnothing}_{c,j}$. Shading denotes no recursion energies. Note that while the subsequence $[i,j]$ is disonnected within the $R^{n}_{\text{INTER}}(i,j,\phi)$ recursion due to the nick at $c$, these states are connected when used in the context of the $R^{b}_{\text{INTER}}$ recursion.

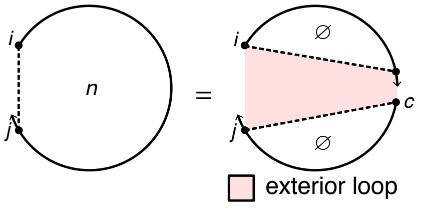

$$R^{n}_{\text{INTER}}(i,j,\phi) \equiv \bigoplus_{c\in\eta} Q^{\varnothing}_{i,c-1} \otimes Q^{\varnothing}_{c,j}$$

Figure S36: $R^{n}_{\text{INTER}}$ recursion with coaxial and dangle stacking. Top: recursion diagram. Bottom: recursion equation.

$\boldsymbol{R^{ms}_{\text{INTER}}}$ **recursion with coaxial and dangle stacking.** The $R^{b}_{\text{INTER}}(i,j,\phi)$ recursion references $Q^{ms}_{d+1,j-l-1}$ elements that are computed using either the $R^{ms}_{\text{INTRA}}$ recursion shown of Figure S26 (if $d+1$ and $j-l-1$ are on the same strand) or the $R^{ms}_{\text{INTER}}$ recursion of Figure S37 (if $d+1$ and $j-l-1$ are on different strands). $R^{ms}_{\text{INTER}}(i,j,\phi)$ operates on a conditional ensemble for subsequence $[i,j]$ in a multiloop context containing one stacking state starting at $i$ and ending in the interval $[i+1,j]$ (depicted as a dashed line between $i$ and $j$). Like other multiloop recursions in the mixed algorithm, $R^{ms}_{\text{INTER}}(i,j,\phi)$ is split into $R^{ms,\text{single}}_{\text{INTER}}(i,j,\phi)$ and $R^{ms,\text{mix}}_{\text{INTER}}(i,j,\phi)$ (the single and mixed recursions are indicated in the diagram with blue and red backbones, respectively):

- $R^{ms,\text{single}}_{\text{INTER}}(i,j,\phi)$: This recursion treats the $R^{ms}_{\text{INTER}}(i,j,\phi)$ conditional ensemble over $[i,j]$ in a multiloop context where all terminal base pairs and unpaired nucleotides are of a single material. There are two cases that are combined using $\oplus$ in the recursion equation:
  - *Coaxial stacking state:* The contribution for the coaxial stacking state in subsequence $[i,j]$ is calculated using a $Q^{mcs,\text{single}}_{i,j}$ element using term $C_{3s}$. Because $Q^{mcs,\text{single}}_{i,j}$ only includes single-material contributions, no further material checks are needed.
  - *Dangle stacking state:* The contribution for the dangle stacking state in subsequence $[i,d]$ is incorporated using a $Q^{md}_{i,d}$ element. For the edge case where $d=j$, the whole region $[i,j]$ is incorporated using a $Q^{md,\text{single}}_{i,j}$ element using term $C_{2s}$. Because $Q^{md,\text{single}}_{i,j}$ only includes single-material contributions, no further material checks are needed. For the base case where $d$ is in $[i+4,j-1]$, then $Q^{md,\text{single}}_{i,d}$ is used to represent the dangle stacking state in $[i,d]$ while the unpaired region $[d+1,j]$ is incorporated using $Q^{ma,\text{single}}_{d+1,j}$. The resulting contribution $Q^{md,\text{single}}_{i,d} \otimes Q^{ma,\text{single}}_{d+1,j}$ of term $C_{1s}$ produces a single-material multiloop only if the material matches for both elements, which is checked with the conitional $\mu_j = \mu_i$. If the conditional is satisfied, the contribution is added to $R^{ms,\text{single}}_{\text{INTER}}(i,j,\phi)$ using term $C_{0s}$. Otherwise, if the conditional fails, the contribution is added to $R^{ms,\text{mix}}_{\text{INTER}}(i,j,\phi)$ using term $C_{0m}$. In the conditional, indices $i$ and $j$ are used to check the material since they are constant over vector $\overline{d}$ and apply to all $d$ and $d+1$ in $\overline{d}$.

- $R^{ms,\text{mix}}_{\text{INTER}}(i,j,\phi)$: This recursion treats the $R^{ms}_{\text{INTER}}(i,j,\phi)$ conditional ensemble over $[i,j]$ in a multiloop context where the terminal base pairs and unpaired nucleotides overall contain a mixture of materials. There are two cases that are combined using $\oplus$ in the recursion equation:
  - *Coaxial stacking state:* The contribution for the coaxial stacking state in subsequence $[i,j]$ is calculated using a $Q^{mcs,\text{mix}}_{i,j}$ element using term $C_{3m}$. Because $Q^{mcs,\text{mix}}_{i,j}$ only includes mixed contributions, no further material checks are needed.

- *Dangle stacking state:* The contribution for the dangle stacking state in subsequence $[i, d]$ is incorporated using a $Q^{md}_{i,d}$ element. For the edge case where $d = j$, the whole region $[i, j]$ is incorporated using a $Q^{md,\text{mix}}_{i,j}$ element using term $C_{2m}$. Because $Q^{md,\text{mix}}_{i,j}$ only includes mixed contributions, no further material checks are needed. For the base case, the contributions for the dangle stacking state in subsequence $[i, d]$ are incorporated with a $Q^{md}_{i,d}$ element while the contributions for the unpaired region $[d + 1, j]$ are incorporated with a $Q^{ma}_{d+1,j}$ element. Because $Q^{md}_{i,d}$ and $Q^{ma}_{d+1,j}$ are each split into single- and mixed-material recursions, there are 4 possible combinations when multiplying them. The three combinations containing at least one mixed element ($Q^{md,\text{single}}_{i,d} \otimes Q^{ma,\text{mix}}_{d+1,j}$, $Q^{md,\text{mix}}_{i,d} \otimes Q^{ma,\text{single}}_{d+1,j}$, $Q^{md,\text{mix}}_{i,d} \otimes Q^{ma,\text{mix}}_{d+1,j}$) must represent mixed contributions, and are included in $R^{ms,\text{mix}}_{\text{INTER}}(i, j, \phi)$ using term $C_{1m}$. As noted above, the remaining combination, $Q^{md,\text{single}}_{i,d} \otimes Q^{ma,\text{single}}_{d+1,j}$, calculated in term $C_{1s}$ represents a mixed contribution if $\mu_j \neq \mu_i$, and is incorporated in $R^{ms,\text{mix}}_{\text{INTER}}(i, j, \phi)$ using term $C_{0m}$.

The shading does not represent any recursion energies as all multiloop contributions are handled by other recursions: 1) there are no terminal base pairs in a multiloop context explicitly defined in this case, 2) there are no unpaired bases in a multiloop context explicitly defined in this case. Nucleotide $d$ must always be on the last strand ($d \geq \text{LAST}(\eta)$) to ensure that there are no nicks in the subsequence $[d, j]$, which would lead to either a disconnected structure (which is not permitted in the complex ensemble) or an exterior loop state (which is not handled by this multiloop recursion). Also, $d \leq j - 1$ in order to provide a minimum of one nucleotide for the $Q^{ma}_{d+1,j}$ recursion, so we must have $j \geq \text{LAST}(\eta) + 1$. Note that $R^{ms}$ directly incorporates the $R^{mcs}$ recursion which serves as an efficiency wrapper of the $R^{mc}$ recursion, and hence, $R^{ms}$ is an efficiency wrapper of $R^{mc}$ (the $3'$-most coaxial stacking state in a multiloop context). Note also that $R^{ms}$ is an efficiency wrapper of the $R^{md}$ recursion (the $3'$-most dangle stacking state in a multiloop context). Taken together $R^{mc}$ and $R^{md}$ represent the $3'$-most stacking state in a multiloop context, analogous to $R^{cd}$ representing the $3'$-most stacking state (coaxial or dangle) in an exterior loop context. The reason that $R^{mc}$ (coaxial stacking states) and $R^{md}$ (dangle stacking states) are calculated and stored separately in a multiloop context is that coaxial-only information (stored in element $Q^{mcs}$) is needed for the previously described multiloop edge case (recursion diagrams in the tenth and eleventh rows of Figure S25). As a result, coaxial-only information is calculated using the efficiency wrapper $R^{mcs}$ for use in that edge case, and then coaxial-only and dangle-only information are combined by the $R^{ms}$ efficiency wrapper (which is fully analogous to the $R^s$ efficiency wrapper in the exterior loop context). With this approach, the operations spent calculating coaxial stacking information for $Q^{mcs}$ elements are not repeated when calculating both coaxial and dangle stacking for $Q^{ms}$ elements.

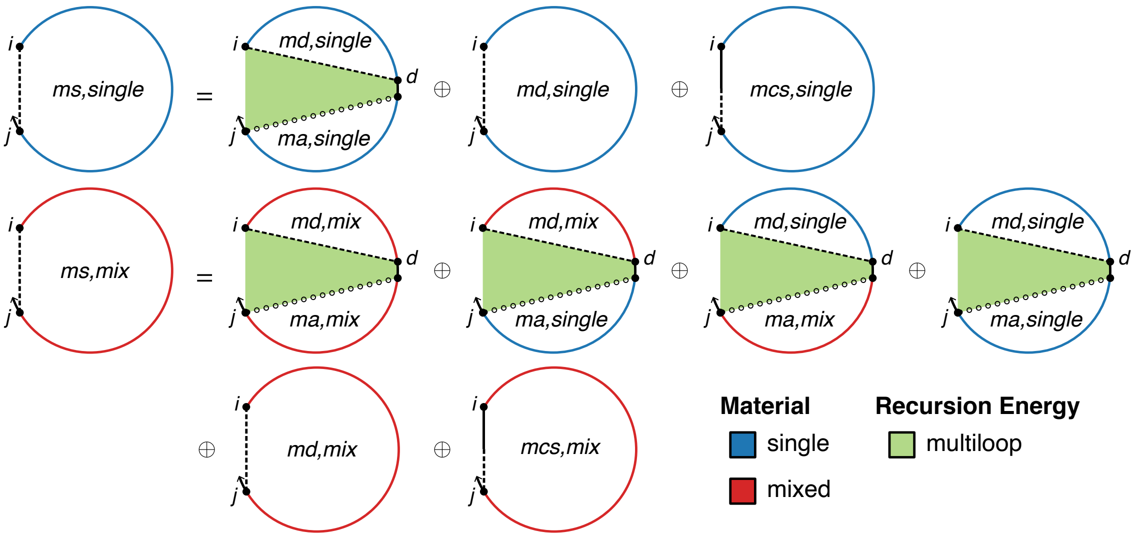

$$R^{ms,\text{single}}_{\text{INTER}}(i,j,\phi) \equiv C_{0s} \oplus C_{2s} \oplus C_{3s}$$

$$C_{0s} \equiv \begin{cases} C_{1s}, & \mu_j = \mu_i \\ \mathbb{0}, & \text{otherwise} \end{cases}$$

$$C_{1s} \equiv \begin{cases} \text{DOT}\left(Q^{md,\text{single}}_{i,\overline{d}}, Q^{ma,\text{single}}_{\overline{d}+1,j}\right), & j > \textsc{Last}(\eta) \\ \mathbb{0}, & \text{otherwise} \end{cases}$$

$$C_{2s} \equiv Q^{md,\text{single}}_{i,j}$$

$$C_{3s} \equiv Q^{mcs,\text{single}}_{i,j}$$

$$R^{ms,\text{mix}}_{\text{INTER}}(i,j,\phi) \equiv C_{0m} \oplus C_{1m} \oplus C_{2m} \oplus C_{3m}$$

$$C_{0m} \equiv \begin{cases} \mathbb{0}, & \mu_j = \mu_i \\ C_{1s}, & \text{otherwise} \end{cases}$$

$$C_{1m} \equiv \begin{cases} \text{DOT}\left(Q^{md,\text{mix}}_{i,\overline{d}}, Q^{ma,\text{single}}_{\overline{d}+1,j}\right) \oplus \text{DOT}\left(Q^{md,\text{single}}_{i,\overline{d}}, Q^{ma,\text{mix}}_{\overline{d}+1,j}\right) \oplus \text{DOT}\left(Q^{md,\text{single}}_{i,\overline{d}}, Q^{ma,\text{mix}}_{\overline{d}+1,j}\right), & j > \textsc{Last}(\eta) \\ \mathbb{0}, & \text{otherwise} \end{cases}$$

$$C_{2m} \equiv Q^{md,\text{mix}}_{i,j}$$

$$C_{3m} \equiv Q^{mcs,\text{mix}}_{i,j}$$

where $\overline{d} \equiv [\textsc{Last}(\eta) : j-1]$

Figure S37: $R^{ms}_{\text{INTER}}$ recursion with coaxial and dangle stacking. Top: recursion diagram. Bottom: recursion equation.

$\boldsymbol{R^{ma}_{\text{INTER}}}$ **recursion with coaxial and dangle stacking.** The $R^{ms}_{\text{INTER}}(i,j,\phi)$ recursion references $Q^{ma}_{d+1,j}$ elements that are always computed using the $R^{ma}_{\text{INTRA}}$ recursion of Figure S27 (if $d+1$ and $j$ are on the same strand) and never computed using the $R^{ma}_{\text{INTER}}$ recursion of Figure S38 (if $d+1$ and $j$ are on different strands). $R^{ma}_{\text{INTER}}(i,j,\phi)$ treats the conditional ensemble of subsequence $[i,j]$ in a multiloop context containing no stacking states, where $i$ and $j$ are on different strands. Note that within the $R^{ms}_{\text{INTER}}(i,j,\phi)$ recursion, $d$ must be on the last strand $(d \geq \textsc{Last}(\eta))$ to ensure there are no nicks in the subsequence $[d,j]$ (which would otherwise lead to a disconnected structure or an exterior loop state), and $d \leq j-1$ in order to provide a minimum of one nucleotide for the $Q^{ma}_{d+1,j}$ recursion. Hence, $d$ and $j$ are always on the same strand and there are no calls to $R^{ma}_{\text{INTER}}$. Equivalently, the contributions for both single- and mixed-material $R^{ma}_{\text{INTER}}(i,j,\phi)$ recursions are $\mathbb{0}$ (Figure S38).

$$R^{ma,\text{single}}_{\text{INTER}}(i,j,\phi) \equiv \mathbb{0}$$
$$R^{ma,\text{mix}}_{\text{INTER}}(i,j,\phi) \equiv \mathbb{0}$$

Figure S38: $R^{ma}_{\text{INTER}}$ recursion with coaxial and dangle stacking. No recursion diagram is provided because the recursion equations contain no contributions.

$\boldsymbol{R^{mcs}_{\text{INTER}}}$ **recursion with coaxial and dangle stacking.** The $R^{ms}_{\text{INTER}}(i,j,\phi)$ recursion references $Q^{mcs}$ elements that are computed using the $R^{mcs}_{\text{INTER}}$ recursion displayed in Figure S39. $R^{mcs}_{\text{INTER}}(i,j,\phi)$ operates on a conditional ensemble for subsequence $[i,j]$ in a multiloop context containing one coaxial stacking state starting at $i$ and ending in the interval $[i+1,j]$ (depicted as a half-solid/half-dashed line between $i$ and $j$). Like other multiloop recursions in the mixed algorithm, $R^{mcs}_{\text{INTER}}(i,j,\phi)$ is split into $R^{mcs,\text{single}}_{\text{INTER}}(i,j,\phi)$ and $R^{mcs,\text{mix}}_{\text{INTER}}(i,j,\phi)$ (the single and mixed recursions are indicated in the diagram with blue and red backbones, respectively):

- $R^{mcs,\text{single}}_{\text{INTER}}(i,j,\phi)$: This recursion treats the $R^{mcs}_{\text{INTER}}(i,j,\phi)$ conditional ensemble over $[i,j]$ in a multiloop context where all terminal base pairs and unpaired nucleotides are of a single material. For the edge case where $d=j$, then whole region $[i,j]$ is incorporated using a $Q^{mc,\text{single}}_{i,j}$ element using term $C_{2s}$. No further material checks are needed since the $R^{mc,\text{single}}_{\text{INTER}}(i,j,\phi)$ recursion only includes single-material contributions. For the base case, then $Q^{mc,\text{single}}_{i,d}$ is used to represent the coaxial stacking state in $[i,d]$ while the unpaired region $[d+1,j]$ is incorporated using $Q^{ma,\text{single}}_{d+1,j}$. The resulting contribution $Q^{mc,\text{single}}_{i,d} \otimes Q^{ma,\text{single}}_{d+1,j}$ of term $C_{1s}$ produces a single-material multiloop only if the material matches for both elements, which is checked with the conitional $\mu_j = \mu_i$. If the conditional is satisfied, the contribution is added to $R^{mcs,\text{single}}_{\text{INTER}}(i,j,\phi)$ using term $C_{0s}$. Otherwise, if the conditional fails, the contribution is added to $R^{mcs,\text{mix}}_{\text{INTER}}(i,j,\phi)$ using term $C_{0m}$. In the conditional, indices $i$ and $j$ are used to check the material since they are constant over vector $\overline{d}$ and apply to all $d$ and $d+1$ in $\overline{d}$.
- $R^{mcs,\text{mix}}_{\text{INTER}}(i,j,\phi)$: This recursion treats the $R^{mcs}_{\text{INTER}}(i,j,\phi)$ conditional ensemble over $[i,j]$ in a multiloop context where the terminal base pairs and unpaired nucleotides overall contain a mixture of materials. For the edge case where $d=j$, then whole region $[i,j]$ is incorporated using a $Q^{mc,\text{mix}}_{i,j}$ element using term $C_{2m}$. No further material checks are needed since the $R^{mc,\text{mix}}_{\text{INTER}}(i,j,\phi)$ recursion only includes mixed material contributions. For the base case, then $Q^{mc}_{i,d}$ is used to represent the coaxial stacking state in $[i,d]$ while the unpaired region $[d+1,j]$ is incorporated using a $Q^{ma}_{d+1,j}$ element. Because $Q^{mc}_{i,d}$ and $Q^{ma}_{d+1,j}$ are each split into single- and mixed-material recursions, there are 4 possible combinations when multiplying them. The three combinations containing at least one mixed element ($Q^{mc,\text{single}}_{i,d} \otimes Q^{ma,\text{mix}}_{d+1,j}$, $Q^{mc,\text{mix}}_{i,d} \otimes Q^{ma,\text{single}}_{d+1,j}$, $Q^{mc,\text{mix}}_{i,d} \otimes Q^{ma,\text{mix}}_{d+1,j}$) must represent mixed contributions, and are included in $R^{mcs,\text{mix}}_{\text{INTER}}(i,j,\phi)$ using term $C_{1m}$. As noted above, the remaining combination, $Q^{mc,\text{single}}_{i,d} \otimes Q^{ma,\text{single}}_{d+1,j}$, calculated in term $C_{1s}$ represents a mixed contribution only if $\mu_j \neq \mu_i$, and is incorporated in $R^{mcs,\text{mix}}_{\text{INTER}}(i,j,\phi)$ using term $C_{0m}$.

The shading does not represent any recursion energies as all multiloop contributions are handled by other recursions: 1) there are no terminal base pairs in a multiloop context explicitly defined in this case, 2) there are no unpaired bases in a multiloop context explicitly defined in this case. Nucleotide $d$ must always be on the last strand $(d \geq \textsc{Last}(\eta))$ to ensure that there are no nicks in the subsequence $[d,j]$, which would lead to either a disconnected structure (which is not permitted in the complex ensemble) or an exterior loop state (which is not handled by this multiloop recursion). Also, $d \leq j-1$ in order to provide a minimum of one nucleotide for the $Q^{ma}_{d+1,j}$ recursion, so we must have $j \geq \textsc{Last}(\eta)+1$. Note that the $R^{mcs}$ recursion serves as an efficiency wrapper of the $R^{mc}$ recursion (here, representing the $3'$-most coaxial stacking state in a multiloop context). The index limits in the recursion equation reflect the steric requirement that there be at least 8 intervening bases between $i$ and $d$ (because the $R^{mc}$ recursion incorporates a coaxial stack involving two adjacent terminal base pairs such that $i$ and $d$ are paired to intervening adjacent bases).

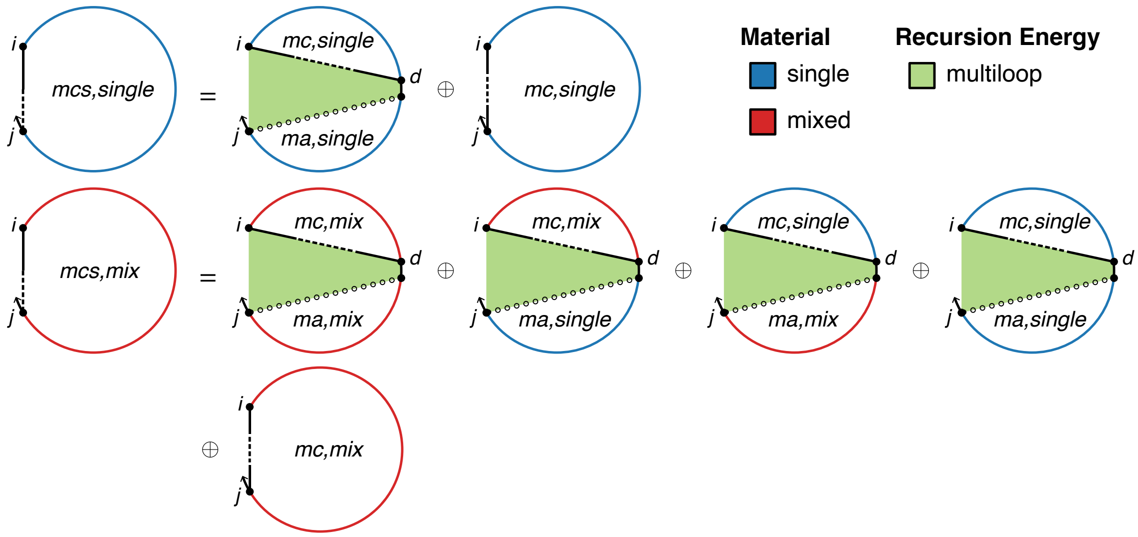

$$R^{mcs,\text{single}}_{\text{INTER}}(i,j,\phi) \equiv C_{0s} \oplus C_{2s}$$

$$C_{0s} \equiv \begin{cases} C_{1s}, & \mu_j = \mu_i \\ \mathbb{0}, & \text{otherwise} \end{cases}$$

$$C_{1s} \equiv \begin{cases} \text{DOT}\left(Q^{mc,\text{single}}_{i,\overline{d}}, Q^{ma,\text{single}}_{\overline{d}+1,j}\right), & j > \textsc{Last}(\eta) \\ \mathbb{0}, & \text{otherwise} \end{cases}$$

$$C_{2s} \equiv Q^{mc,\text{single}}_{i,j}$$

$$R^{mcs,\text{mix}}_{\text{INTER}}(i,j,\phi) \equiv C_{0m} \oplus C_{1m} \oplus C_{2m}$$

$$C_{0m} \equiv \begin{cases} \mathbb{0}, & \mu_j = \mu_i \\ C_{1s}, & \text{otherwise} \end{cases}$$

$$C_{1m} \equiv \begin{cases} \text{DOT}\left(Q^{mc,\text{mix}}_{i,\overline{d}}, Q^{ma,\text{single}}_{\overline{d}+1,j}\right) \oplus \text{DOT}\left(Q^{mc,\text{single}}_{i,\overline{d}}, Q^{ma,\text{mix}}_{\overline{d}+1,j}\right) \oplus \text{DOT}\left(Q^{mc,\text{mix}}_{i,\overline{d}}, Q^{ma,\text{mix}}_{\overline{d}+1,j}\right), & j > \textsc{Last}(\eta) \\ \mathbb{0}, & \text{otherwise} \end{cases}$$

$$C_{2m} \equiv Q^{mc,\text{mix}}_{i,j}$$

where $\overline{d} \equiv [\textsc{Last}(\eta) : j-1]$

Figure S39: $R^{mcs}_{\text{INTER}}$ recursion with coaxial and dangle stacking. Top: recursion diagram. Bottom: recursion equation.

**$R^{mc}_{\text{INTER}}$ recursion with coaxial and dangle stacking.** The $R^{mcs}_{\text{INTER}}(i,j,\phi)$ recursion references $Q^{mc}$ elements that are computed using the $R^{mc}_{\text{INTER}}$ recursion displayed in Figure S40. This recursion treats a single coaxial stacking state in a multiloop context (depicted as a straight line between $i$ and $j$ that is solid at both ends and dashed in the middle to indicate that $i$ and $j$ are both base-paired but not to each other). Two adjacent terminal base pairs ($i \cdot d$ and $d+1 \cdot j$) coaxially stack on each other. Each terminal base pair is either a single material (denoted in the recursion diagram by a blue base pair, if the material of the two nucleotides matches) or mixed material (denoted by a red base pair, if the material of the two nucleotides does not match). The contributions of subsequences $[i,d]$ and $[d+1,j]$ are incorporated using $Q^b_{i,d}$ and $Q^b_{d+1,j}$ elements in term $C_{\text{stack}}$. The index limits in the recursion equation reflect the steric requirement that there be at least 3 intervening bases between $i$ and $d$ and at least 3 intervening bases between $d+1$ and $j$. Like other multiloop recursions in the mixed algorithm, $R^{mc}_{\text{INTER}}(i,j,\phi)$ is split into $R^{mc,\text{single}}_{\text{INTER}}(i,j,\phi)$ and $R^{mc,\text{mix}}_{\text{INTER}}(i,j,\phi)$ (the single and mixed recursions are indicated in the diagram with blue and red backbones, respectively):

- $R^{mc,\text{single}}_{\text{INTER}}(i,j,\phi)$: This recursion treats the $R^{mc}_{\text{INTER}}(i,j,\phi)$ conditional ensemble over $[i,j]$ where terminal base pairs $i \cdot d$ and $d+1 \cdot j$ are of a single material. The case where both terminal base pairs are single-material (checked with conditional "$\mu_d = \mu_i$ and $\mu_{d+1} = \mu_j$" in term $C_{1s}$) leads to a single-material multiloop only if the material matches for both terminal base pairs, which is checked with the conditional $\mu_j = \mu_i$. If the conditional is satisfied, the contribution is added to $R^{mc,\text{single}}_{\text{INTER}}(i,j,\phi)$ using term $C_{0s}$. Otherwise, if the conditional fails, the contribution is added to $R^{mc,\text{mix}}_{\text{INTER}}(i,j,\phi)$ using term $C_{0m}$. In the conditional, indices $i$ and $j$ are used to check the material since they are constant over vector $\overline{d}$ and apply to all $d$ and $d+1$ in $\overline{d}$.

- $R^{mc,\text{mix}}_{\text{INTER}}(i,j,\phi)$: This recursion treats the $R^{mc}_{\text{INTER}}(i,j,\phi)$ conditional ensemble over $[i,j]$ where the terminal base pairs $i \cdot d$ and $d+1 \cdot j$ overall contain a mixture of materials. Because each of the terminal base pairs can be single- or mixed-material, there are four possible single/mix combinations for $Q^b_{i,d} \otimes Q^b_{d+1,j}$. The three combinations containing at least one mixed base pair are included in $R^{mc,\text{mix}}_{\text{INTER}}(i,j,\phi)$ using term $C_{1m}$. As noted above, the remaining combination of term $C_{1s}$ where each terminal base pair is single-material represents a mixed contribution only if $\mu_j \neq \mu_i$ and is incorporated in $R^{mc,\text{mix}}_{\text{INTER}}(i,j,\phi)$ using term $C_{0m}$.

Shading corresponds to three kinds of recursion energy: 1) the sequence-independent penalties for two terminal base pairs in a multiloop context (for base pair $i \cdot d$, $\Delta G^{\text{multi}}_{\text{bp},\mu_i}$ if $\mu_d = \mu_i$, and $\Delta G^{\text{multi}}_{\text{bp,mix}}$ otherwise; for base pair $d+1 \cdot j$, $\Delta G^{\text{multi}}_{\text{bp},\mu_j}$ if $\mu_{d+1} = \mu_j$, and $\Delta G^{\text{multi}}_{\text{bp,mix}}$ otherwise), 2) the sequence-dependent penalties for two terminal base pairs in a multiloop context ($\Delta G^{\text{terminalbp}}_{i,d}(\phi)$ and $\Delta G^{\text{terminalbp}}_{d+1,j}(\phi)$, dependent on the sequence and material of base pairs $i \cdot d$ and $d+1 \cdot j$), 3) the sequence-dependent coaxial stacking free energy $\Delta G^{\text{coax}}_{i,d,j}(\phi)$ (dependent on the sequence and material of base pairs $i \cdot d$ and $d+1 \cdot j$); note that $\Delta G^{\text{coax}}_{i,d,j}(\phi)$ requires only 3 indices because $d+1$ is implied by $d$. To exclude exterior loop states that are not treated by this multiloop recursion, the function VALID returns the set of valid vectorization ranges for which nucleotides $d$ and $d+1$ are on the same strand (i.e., such that $d$ and $d+1$ do not take on values that would place a nick between them).

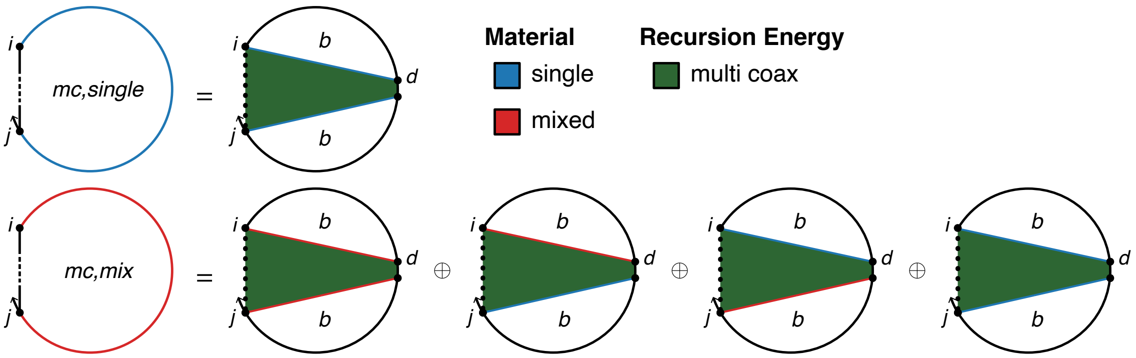

$$R^{mc,\text{single}}_{\text{INTER}}(i,j,\phi) \equiv C_{0s}$$

$$\text{where } C_{0s} \equiv \begin{cases} C_{1s}, & \mu_j = \mu_i \\ \mathbb{0}, & \text{otherwise} \end{cases}$$

$$C_{1s} \equiv \begin{cases} C_{\text{stack}} \otimes W(\Delta G^{\text{multi}}_{\text{bp},\mu_i} + \Delta G^{\text{multi}}_{\text{bp},\mu_j}), & \mu_d = \mu_i \text{ and } \mu_{d+1} = \mu_j \\ \mathbb{0}, & \text{otherwise} \end{cases}$$

$$C_{\text{stack}} \equiv \bigoplus_{\overline{d} \in \textsc{Valid}(i,j,\eta)} \textsc{dot}\left(Q^b_{i,\overline{d}}, Q^b_{\overline{d}+1,j}, W(\Delta G^{\text{coax}}_{i,\overline{d},j}(\phi) + \Delta G^{\text{terminalbp}}_{i,\overline{d}}(\phi) + \Delta G^{\text{terminalbp}}_{\overline{d}+1,j}(\phi))\right)$$

$$R^{mc,\text{mix}}_{\text{INTER}}(i,j,\phi) \equiv C_{0m} \oplus C_{1m}$$

$$\text{where } C_{0m} \equiv \begin{cases} \mathbb{0}, & \mu_j = \mu_i \\ C_{1s}, & \text{otherwise} \end{cases}$$

$$C_{1m} \equiv \begin{cases} C_{\text{stack}} \otimes W(\Delta G^{\text{multi}}_{\text{bp},\mu_i} + \Delta G^{\text{multi}}_{\text{bp,mix}}), & \mu_d = \mu_i \text{ and } \mu_{d+1} \neq \mu_j \\ \mathbb{0}, & \text{otherwise} \end{cases}$$

$$\oplus \begin{cases} C_{\text{stack}} \otimes W(\Delta G^{\text{multi}}_{\text{bp,mix}} + \Delta G^{\text{multi}}_{\text{bp},\mu_j}), & \mu_d \neq \mu_i \text{ and } \mu_{d+1} = \mu_j \\ \mathbb{0}, & \text{otherwise} \end{cases}$$

$$\oplus \begin{cases} C_{\text{stack}} \otimes W(2\Delta G^{\text{multi}}_{\text{bp,mix}}), & \mu_d \neq \mu_i \text{ and } \mu_{d+1} \neq \mu_j \\ \mathbb{0}, & \text{otherwise} \end{cases}$$

Figure S40: $R^{mc}_{\text{INTER}}$ recursion with coaxial and dangle stacking. Top: recursion diagram. Bottom: recursion equation.

$\boldsymbol{R^{md}_{\text{INTER}}}$ **recursion with coaxial and dangle stacking.** The $R^{ms}_{\text{INTER}}(i,j,\phi)$ recursion references $Q^{md}$ elements that are computed using the $R^{md}_{\text{INTER}}$ recursion displayed in Figure S41. This recursion treats a single dangle stacking state (depicted as a dashed line between $i$ and $j$) in a multiloop context with either zero, one, or two unpaired nucleotides dangle stacking on an adjacent terminal base pair ($i+k \cdot j-l$). The recursion diagram represents these four alternative dangle stacking states corresponding to no dangles, $5'$ dangle, $3'$ dangle, or terminal mismatch (see Figure S21 for details). The terminal base pair is either a single material (denoted in the recursion diagram with a blue base pair, if $\mu_{i+k} = \mu_{j-l}$), or mixed-material (denoted by a red base pair, if $\mu_{i+k} \neq \mu_{j-l}$). The contribution of subsequence $[i+k, j-l]$ is incorporated using a $Q^b_{i+k,j-l}$ element in term $C_{\text{stack}}$. The index limits in the recursion equation reflect the steric requirement that there be at least 3 intervening bases between $i+k$ and $j-l$. Like other multiloop recursions in the mixed algorithm, $R^{md}_{\text{INTER}}(i,j,\phi)$ is split into $R^{md,\text{single}}_{\text{INTER}}(i,j,\phi)$ and $R^{md,\text{mix}}_{\text{INTER}}(i,j,\phi)$ (the single and mixed recursions are indicated in the diagram with blue and red backbones, respectively):

- $R^{md,\text{single}}_{\text{INTER}}(i,j,\phi)$: This recursion treats the $R^{md}_{\text{INTER}}(i,j,\phi)$ conditional ensemble over $[i,j]$ where terminal base pair $i+k \cdot j-l$ and any unpaired dangling nucleotides are of a single material.

- $R^{md,\text{mix}}_{\text{INTER}}(i,j,\phi)$: This recursion treats the $R^{md}_{\text{INTER}}(i,j,\phi)$ conditional ensemble over $[i,j]$ where terminal base pair $i+k \cdot j-l$ and any unpaired dangling nucleotides overall contain a mixture of materials.

The conditional "$\mu_{i+k} = \mu_{j-l} = \mu_j = \mu_i$" is used to check if the terminal base pair $i+k \cdot j-l$ and any dangling nucleotides are of a single material. If the conditional is satisfied, $C_{\text{stack}}$ is added to $R^{md,\text{single}}_{\text{INTER}}(i,j,\phi)$. Otherwise, if the conditional is not satisfied, $C_{\text{stack}}$ is added to $R^{md,\text{mix}}_{\text{INTER}}(i,j,\phi)$. Shading corresponds to four recursion energies: 1) the sequence-independent penalty for one terminal base pair in a multiloop context, $\Delta G^{\text{multi}}_{\text{bp},\sigma}$ (corresponding to base pair $j-l \cdot i+k$), where the parameter depends on the material as determined by the $\sigma$ conditional, 2) the penalty per unpaired nucleotide in a multiloop context $\Delta G^{\text{multi}}_{\text{nt},\mu_i}$ or $\Delta G^{\text{multi}}_{\text{nt},\mu_j}$ (a total of $k+l$ dangling nucleotides; as a result this term is zeroed out when $k = l = 0$), where the parameter depends on the material of each dangling nucleotide ($\mu_i$ or $\mu_j$), 3) the sequence-dependent penalty for a terminal base pair in a multiloop loop context, $\Delta G^{\text{terminalbp}}_{i+k,j-l}(\phi)$ (dependent on the sequence and material of base pair $i+k \cdot j-l$), 4) the sequence-dependent dangle stacking free energy $\Delta G^{\text{dangle}}_{i,i+k,j-l,j}(\phi)$ (dependent on the sequence and material of the terminal base pair and dangle stacking bases) which takes on one of four values corresponding to the four dangle stacking states (see Figure S21). To exclude exterior loop states that are not treated by this multiloop recursion, the index limits in the recursion equation prevent a nick between a dangling nucleotide and the base pair on which it stacks (i.e., no nick between $i$ and $i+k$, and no nick between $j$ and $j-l$).

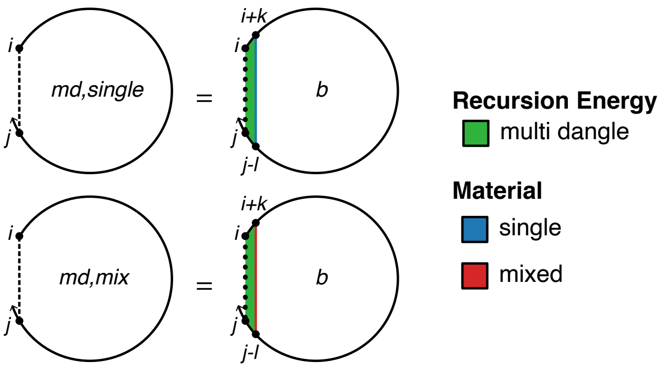

$$R^{md,\text{single}}_{\text{INTER}}(i,j,\phi) \equiv \bigoplus_{\substack{k\in\{0,1\}\\ l\in\{0,1\}}} \begin{cases} C_{\text{stack}}, & \mu_{i+k} = \mu_{j-l} = \mu_j = \mu_i \\ \mathbb{0}, & \text{otherwise} \end{cases}$$

$$R^{md,\text{mix}}_{\text{INTER}}(i,j,\phi) \equiv \bigoplus_{\substack{k\in\{0,1\}\\ l\in\{0,1\}}} \begin{cases} \mathbb{0}, & \mu_{i+k} = \mu_{j-l} = \mu_j = \mu_i \\ C_{\text{stack}}, & \text{otherwise} \end{cases}$$

$$\text{where } C_{\text{stack}} \equiv \begin{cases} Q^b_{i+k,j-l} \otimes \\ W(\Delta G^{\text{dangle}}_{i,i+k,j-l,j}(\phi) + \Delta G^{\text{multi}}_{\text{bp},\sigma} + k\Delta G^{\text{multi}}_{\text{nt},\mu_i} + l\Delta G^{\text{multi}}_{\text{nt},\mu_j} + \Delta G^{\text{terminalbp}}_{i+k,j-l}(\phi)), & i+k < m \text{ and } j-l \geq n \\ \mathbb{0}, & \text{otherwise} \end{cases}$$

$$\text{with } \sigma \equiv \begin{cases} \mu_{i+k}, & \mu_{i+k} = \mu_{j-l} \\ \text{mix}, & \text{otherwise} \end{cases}$$

$$m = \textsc{First}(\eta)$$

$$n = \textsc{Last}(\eta)$$

Figure S41: $R^{md}_{\text{INTER}}$ recursion with coaxial and dangle stacking. Top: recursion diagram. Bottom: recursion equation.

**$R^{m}_{\text{INTER}}$ recursion with coaxial and dangle stacking.** The $R^{b}_{\text{INTER}}(i,j,\phi)$ recursion references $Q^m$ elements that are computed using either the $R^{m}_{\text{INTRA}}$ recursion of Figure S31 (if the end points of the referenced subsequence are on the same strand), or the $R^{m}_{\text{INTER}}$ recursion of Figure S42 (if the end points of the referenced subsequence are on different strands). $R^{m}_{\text{INTER}}(i,j,\phi)$ operates on a conditional ensemble for subsequence $[i,j]$ in a multiloop context where $i$ and $j$ may or may not be paired (depicted with a dashed line between $i$ and $j$ in the recursion diagram) and where there is at least one stacking state in the subsequence. Like other multiloop recursions in the mixed algorithm, $R^{m}_{\text{INTER}}(i,j,\phi)$ is split into $R^{m,\text{single}}_{\text{INTER}}(i,j,\phi)$ and $R^{m,\text{mix}}_{\text{INTER}}(i,j,\phi)$ (the single and mixed recursions are indicated in the diagram with blue and red backbones, respectively):

- $R^{m,\text{single}}_{\text{INTER}}(i,j,\phi)$: This recursion operates over the $R^{m}_{\text{INTER}}(i,j,\phi)$ conditional ensemble on $[i,j]$ in a multiloop context where any terminal base pair or unpaired nucleotide is of a single material. This recursion distinguishes two cases that are combined using $\oplus$ in the recursion equation:
  - *One stacking state*: the case where there is exactly one stacking state in subsequence $[i,j]$ in a multiloop context. This stacking state starts at $d$ and ends in the interval $[d+1,j]$ (depicted by a straight dashed line between $d$ and $j$); subsequence $[i,d-1]$ does not have any stacking states (depicted by a hollow dotted line between $i$ and $d-1$). For the edge case with $d=i$, the entire region $[i,j]$ is incorporated using a $Q^{ms,\text{single}}_{i,j}$ element with term $C_{3s}$. No further material checks are needed since the $R^{ms,\text{single}}_{\text{INTER}}(i,j,\phi)$ recursion only includes single-material contributions. For the base case, then the unpaired region $[i,d-1]$ is incorporated using a $Q^{ma,\text{single}}_{i,d-1}$ element while the region $[d,j]$ is incorporated using $Q^{ms,\text{single}}_{d,j}$. The resulting contribution $Q^{ma,\text{single}}_{i,d-1} \otimes Q^{ms,\text{single}}_{d,j}$ of term $C_{1s}$ produces a single-material multiloop only if the material matches for both elements, which is checked with the conditional $\mu_j = \mu_i$. If the conditional is satisfied, the contribution is added to $R^{m,\text{single}}_{\text{INTER}}(i,j,\phi)$ using term $C_{0s}$. Otherwise, if

the conditional fails, the contribution is added to $R^{m,\text{mix}}_{\text{INTER}}(i, j, \phi)$ using term $C_{0m}$. In the conditional, indices $i$ and $j$ are used to check the material since they are constant over vector $\overline{d}$ and apply to all $d-1$ and $d$ in $\overline{d}$.

- *More than one stacking state*: the case where there are two or more stacking states in subsequence $[i, j]$ in a multiloop context. The $3'$-most stacking state starts at $e+1$ and ends in the interval $[e+2, j]$ (depicted by a straight dashed line between $e+1$ and $j$); the contribution of subsequence $[e+1, j]$ is incorporated by element $Q^{ms,\text{single}}_{e+1,j}$. There are one or more additional stacking states in the interval $[i, e]$ (the straight dashed line denotes that $i$ and $e$ may or may not be paired); the contribution of subsequence $[i, e]$ is incorporated by element $Q^{m,\text{single}}_{i,e}$. The resulting contribution $Q^{ma,\text{single}}_{i,e} \otimes Q^{ms,\text{single}}_{e+1,j}$ of term $C_{2s}$ produces a single-material multiloop only if the material matches for both elements, which is checked with the conditional $\mu_j = \mu_i$. If the conditional is satisfied, the contribution is added to $R^{m,\text{single}}_{\text{INTER}}(i, j, \phi)$ using term $C_{0s}$. Otherwise, if the conditional fails, the contribution is added to $R^{m,\text{mix}}_{\text{INTER}}(i, j, \phi)$ using term $C_{0m}$. In the conditional, indices $i$ and $j$ are used to check the material since they are constant over vector $\overline{e}$ and apply to all $e$ and $e+1$ in $\overline{e}$.

- $R^{m,\text{mix}}_{\text{INTER}}(i, j, \phi)$: This recursion operates over the $R^{m}_{\text{INTER}}(i, j, \phi)$ conditional ensemble on $[i, j]$ in a multiloop context where the terminal base pairs and unpaired nucleotides overall contain a mixture of materials. This recursion distinguishes two cases that are combined using $\oplus$ in the recursion equation:

  - *One stacking state*: the case where there is exactly one stacking state in subsequence $[i, j]$ in a multiloop context. This stacking state starts at $d$ and ends in the interval $[d+1, j]$ (depicted by a straight dashed line between $d$ and $j$); subsequence $[i, d-1]$ does not have any stacking states (depicted by a hollow dotted line between $i$ and $d-1$). For the edge case with $d = i$, the entire region $[i, j]$ is incorporated using a $Q^{ms,\text{mix}}_{i,j}$ element with term $C_{3m}$. No further material checks are needed since the $R^{ms,\text{mix}}_{\text{INTER}}(i, j, \phi)$ recursion only includes mixed material contributions. For the base case, then the unpaired region $[i, d-1]$ is incorporated using a $Q^{ma}_{i,d-1}$ element while the region $[d, j]$ is incorporated using a $Q^{ms}_{d,j}$ element. Because $Q^{ma}_{i,d-1}$ and $Q^{ms}_{d,j}$ are each split into single- and mixed-material recursions, there are 4 possible combinations when multiplying them. The three combinations containing at least one mixed element ($Q^{ma,\text{single}}_{i,d-1} \otimes Q^{ms,\text{mix}}_{d,j}$, $Q^{ma,\text{mix}}_{i,d-1} \otimes Q^{ms,\text{single}}_{d,j}$, $Q^{ma,\text{mix}}_{i,d-1} \otimes Q^{ms,\text{mix}}_{d,j}$) must represent mixed contributions, and are included in $R^{m,\text{mix}}_{\text{INTER}}(i, j, \phi)$ using term $C_{1m}$. As noted above, the remaining combination, $Q^{ma,\text{single}}_{i,d-1} \otimes Q^{ms,\text{single}}_{d,j}$, calculated in term $C_{1s}$ represents a mixed contribution only if $\mu_j \neq \mu_i$, and is incorporated in $R^{m,\text{mix}}_{\text{INTER}}(i, j, \phi)$ using term $C_{0m}$.
  - *More than one stacking state*: the case where there are two or more stacking states in subsequence $[i, j]$ in a multiloop context. The $3'$-most stacking state starts at $e+1$ and ends in the interval $[e+2, j]$ (depicted by a straight dashed line between $e+1$ and $j$); the contribution of subsequence $[e+1, j]$ is incorporated by element $Q^{ms}_{e+1,j}$. There are one or more additional stacking states in the interval $[i, e]$ (the straight dashed line denotes that $i$ and $e$ may or may not be paired); the contribution of subsequence $[i, e]$ is incorporated by element $Q^{m}_{i,e}$. Because $Q^{m}_{i,e}$ and $Q^{ms}_{e+1,j}$ are each split into single- and mixed-material recursions, there are 4 possible combinations when multiplying them. The three combinations containing at least one mixed element ($Q^{m,\text{single}}_{i,e} \otimes Q^{ms,\text{mix}}_{e+1,j}$, $Q^{m,\text{mix}}_{i,e} \otimes Q^{ms,\text{single}}_{e+1,j}$, $Q^{m,\text{mix}}_{i,e} \otimes Q^{ms,\text{mix}}_{e+1,j}$) must represent overall mixed contributions, and are included in $R^{m,\text{mix}}_{\text{INTER}}(i, j, \phi)$ using term $C_{2m}$. As noted above, the remaining combination, $Q^{m,\text{single}}_{i,e} \otimes Q^{ms,\text{single}}_{e+1,j}$, calculated in term $C_{2s}$ represents a mixed contribution only if $\mu_j \neq \mu_i$, and is incorporated in $R^{m,\text{mix}}_{\text{INTER}}(i, j, \phi)$ using term $C_{0m}$.

The shading does not represent any recursion energies as all multiloop contributions are handled by other recursions: 1) there are no terminal base pairs in a multiloop context explicitly defined in this case, 2) there are no unpaired bases in a multiloop context explicitly defined in this case. Nucleotide $d$ must always be on the first strand ($d \leq \text{FIRST}(\eta) - 1$) to ensure that there are no nicks in the subsequence $[i, d]$, which would lead to either a disconnected structure (which is not permitted in the complex ensemble) or an exterior loop state (which is not handled by this multiloop recursion). Also, $d \geq i+1$ in order to provide a minimum of one nucleotide for the $Q^{ma}_{i,d-1}$ recursion, so we must have $i \leq \text{FIRST}(\eta) - 2$ for terms $C_{1s}$ and $C_{1m}$. To exclude exterior loop states that are not treated by this multiloop recursion, the function VALID returns the set of valid vectorization ranges for which nucleotides $e$ and $e+1$ are on the same strand for terms $C_{2s}$ and $C_{2m}$ (i.e., such that $e$ and $e+1$ do not take on values that would place a nick between them).

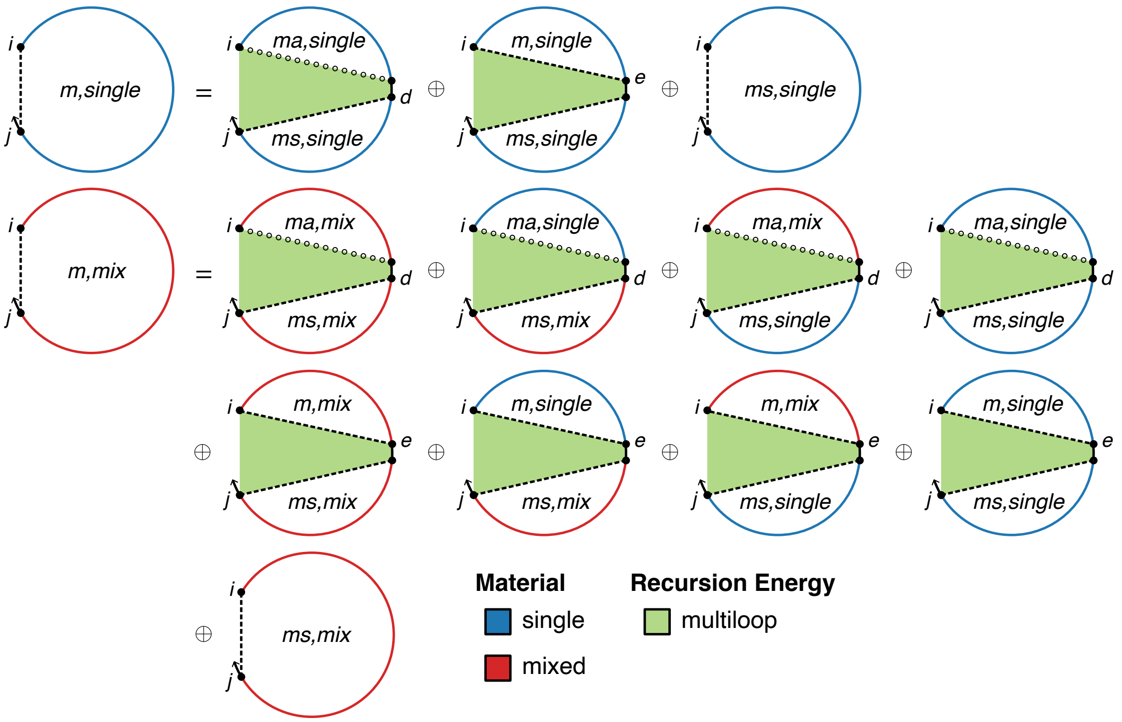

$$R_{\text{INTER}}^{m,\text{single}}(i,j,\phi) \equiv C_{0s} \oplus C_{3s}$$

$$C_{0s} \equiv \begin{cases} C_{1s} \oplus C_{2s}, & \mu_j = \mu_i \\ \mathbb{0}, & \text{otherwise} \end{cases}$$

$$C_{1s} \equiv \begin{cases} \text{DOT}\left(Q_{i,\bar{d}-1}^{ma,\text{single}}, Q_{\bar{d},j}^{ms,\text{single}}\right), & i < \text{FIRST}(\eta) - 1 \\ \mathbb{0}, & \text{otherwise} \end{cases}$$

$$C_{2s} \equiv \bigoplus_{\bar{e} \in \text{VALID}(i,j,\eta)} \text{DOT}\left(Q_{i,\bar{e}}^{m,\text{single}}, Q_{\bar{e}+1,j}^{ms,\text{single}}\right)$$

$$C_{3s} \equiv Q_{i,j}^{ms,\text{single}}$$

$$R_{\text{INTER}}^{m,\text{mix}}(i,j,\phi) \equiv C_{0m} \oplus C_{1m} \oplus C_{2m} \oplus C_{3m}$$

$$C_{0m} \equiv \begin{cases} \mathbb{0}, & \mu_j = \mu_i \\ C_{1s} \oplus C_{2s}, & \text{otherwise} \end{cases}$$

$$C_{1m} \equiv \begin{cases} \text{DOT}\left(Q_{i,\bar{d}-1}^{ma,\text{mix}}, Q_{\bar{d},j}^{ms,\text{mix}}\right) \oplus \text{DOT}\left(Q_{i,\bar{d}-1}^{ma,\text{single}}, Q_{\bar{d},j}^{ms,\text{mix}}\right) & \\ \oplus\, \text{DOT}\left(Q_{i,\bar{d}-1}^{ma,\text{mix}}, Q_{\bar{d},j}^{ms,\text{single}}\right), & i < \text{FIRST}(\eta) - 1 \\ \mathbb{0}, & \text{otherwise} \end{cases}$$

$$C_{2m} \equiv \bigoplus_{\bar{e} \in \text{VALID}(i,j,\eta)} \left[\text{DOT}\left(Q_{i,\bar{e}}^{m,\text{mix}}, Q_{\bar{e}+1,j}^{ms,\text{mix}}\right) \oplus \text{DOT}\left(Q_{i,\bar{e}}^{m,\text{single}}, Q_{\bar{e}+1,j}^{ms,\text{mix}}\right) \oplus \text{DOT}\left(Q_{i,\bar{e}}^{m,\text{mix}}, Q_{\bar{e}+1,j}^{ms,\text{single}}\right)\right]$$

$$C_{3m} \equiv Q_{i,j}^{ms,\text{mix}}$$

where $\bar{d} \equiv [i+1 : \text{FIRST}(\eta) - 1]$

Figure S42: $R_{\text{INTER}}^{m}$ recursion with coaxial and dangle stacking. Top: recursion diagram. Bottom: recursion equation.

#### S7 Additional studies

##### S7.1 Comparison of single- and mixed-material algorithm speeds for MFE determination and MFE proxy structure determination

Figure S43 compares the speed of single- and mixed-material algorithms for MFE determination (panels a and b) and MFE proxy structure determination (panels c and d). The MFE proxy structure is calculated by running the suboptimal structure generation algorithm with $\Delta G_{\text{gap}} = 0$. Like the partition function algorithm (Figure 10), these mixed-material algorithms are $\approx$2.0–3.5$\times$ slower than the single-material algorithms across a wide range of complex sizes (panels b and d).

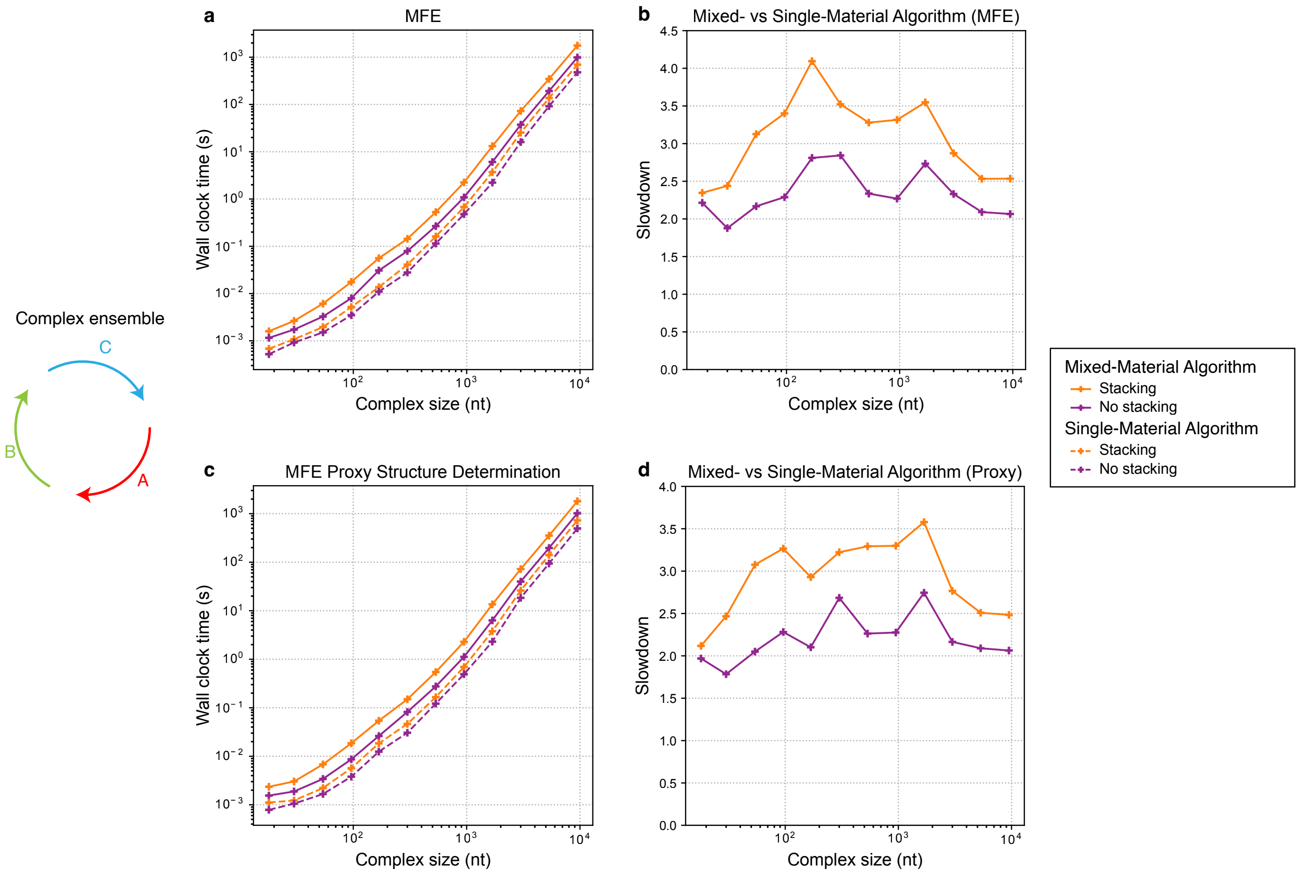

Figure S43: Relative speed of single- and mixed-material algorithms for MFE determination and MFE proxy structure determination. Calculations performed for a complex of 3 strands, each with a different random sequence of uniform length. For the mixed-material tests, each strand contained a random mix of RNA and DNA nucleotides. For single-material tests, the same strand sequences were used with only DNA nucleotides. Single- and mixed-material partition function algorithms with or without coaxial and dangle stacking. (a) Computational cost of MFE determination. (b) Computational slowdown for MFE determination (ratio of mean wall clock times). (c) Computational cost of MFE proxy structure determination. (b) Computational slowdown for MFE proxy structure determination (ratio of mean wall clock times). Mean wall clock time over 75 sets of random sequences per complex size for sizes less than 300 nucleotides, and 5 sets of random sequences per complex size for sizes greater than or equal to 300 nucleotides. Conditions: 37 °C, 1M $Na^{+}$.

#### S7.2 Melting temperature accuracy with coaxial and dangle stacking

Figures S44–S46 use the `stacking` ensemble as compared to Figures 7–9 which use the `nostacking` ensemble.

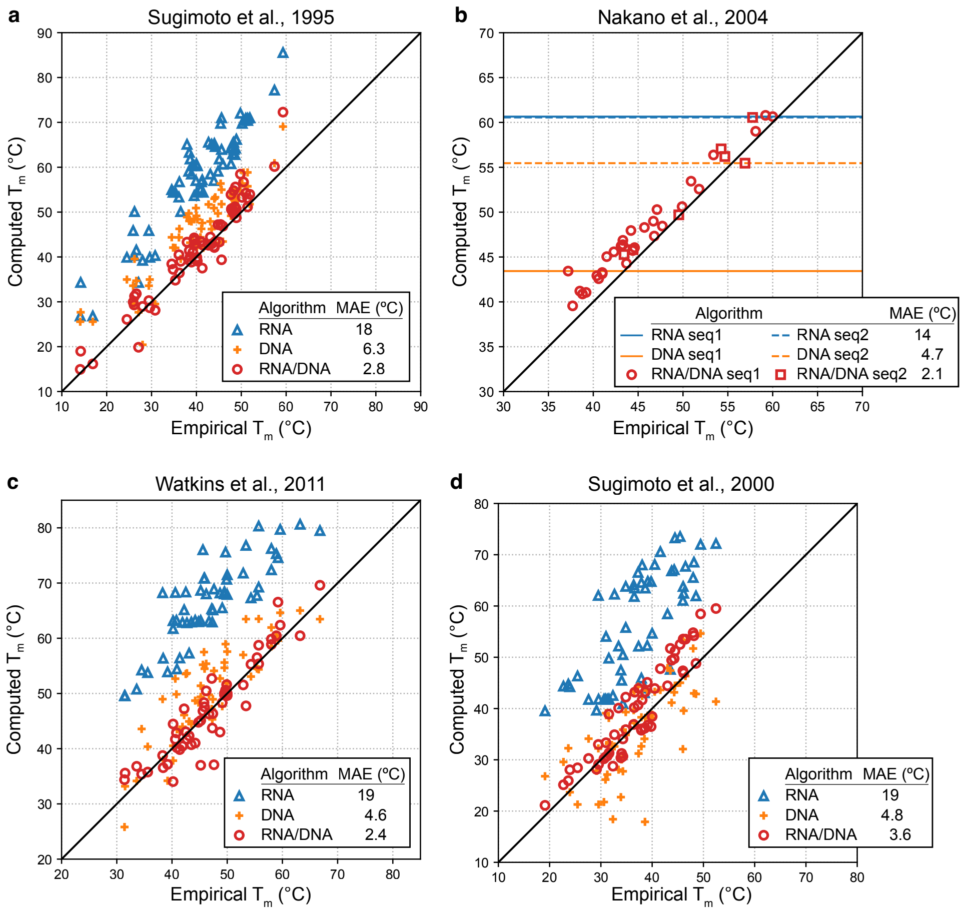

Figure S44: Prediction of duplex melting temperature with `stacking` ensemble using a mixed-material RNA/DNA algorithm compared to single-material RNA-only or DNA-only algorithms (cf. Figure 7 using `nostacking` ensemble). Computed vs empirical melting temperatures ($T_m$) for four different experimental studies. Mean absolute error (MAE). (a) Sugimoto et al, 1995.[32] 67 RNA/DNA hybrid duplexes with no mismatches; one duplex was omitted that did not melt between 0 °C and 100 °C using RNA parameters. (b) Nakano et al., 2004.[35] 37 chimeric duplexes with no mismatches (30 duplexes with the same non-self-complementary sequence (termed seq1) with varying RNA and DNA composition e.g., rGACUAdGGT/rACCUAGUC, dGACTrAGGU/dACCTAGTC, dGACTrAGGU/rACCUdAGTC; 7 duplexes with the same self-complementary sequence (termed seq2) with varying RNA and DNA composition e.g., rCGCGCdG/rCGCGCdG, rCGCdGCG/rCGCdGCG; the single-material predictions are the same for all duplexes with the same sequence (depicted as horizontal lines). (c) Watkins et al., 2011.[33] 54 RNA/DNA hybrid duplexes: 43 contain a single internal rA×dA, rC×dC, rG×dG, or rU×dT mismatch, 11 contain no mismatch; duplexes that appear in Sugimoto et al., 2000[34] were omitted. (d) Sugimoto et al., 2000.[34] 57 RNA/DNA hybrid duplexes: 39 contain a single mismatch (rA×dA, rA×dC, rA×dG, rC×dA, rC×dT, rG×dA, rG×dG,rU×dC, rU×dG, or rU×dT), 9 contain a wobble pair (rG·dT), 4 contain a chimeric mismatch (dT×dG), 5 contain no mismatch; eight sequences containing dU were omitted since there are no DNA parameters for U. Each strand: 50 μM. Salt conditions: 1.0 M $Na^+$.

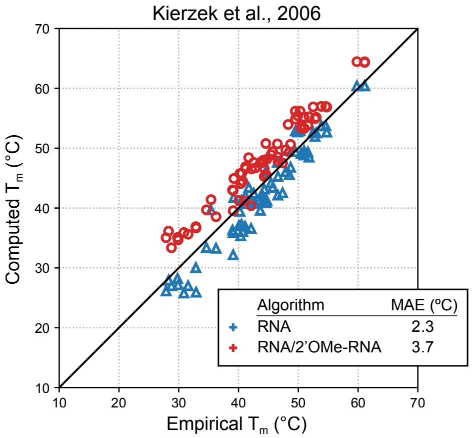

Figure S45: Prediction of duplex melting temperature with `stacking` ensemble using a mixed-material RNA/2′OMe-RNA algorithm compared to a single-material RNA-only algorithm (cf. Figure 8 using `nostacking` ensemble). Computed vs empirical melting temperature ($T_m$) for the experimental study of Kierzek et al., 2006.[30] Mean absolute error (MAE). 68 RNA/2′OMe-RNA hybrid duplexes without mismatches.[30] Each strand: 50 μM. Salt conditions: 0.12 M $Na^+$.

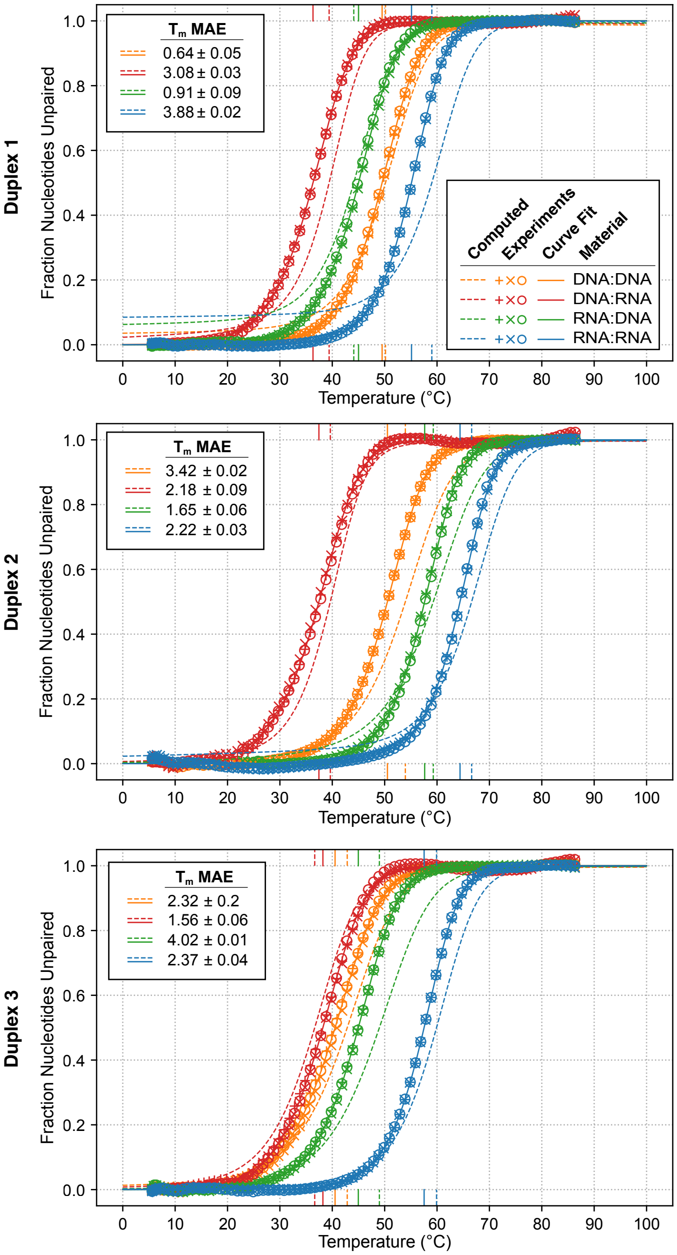

Figure S46: Comparison of melt curves computed with `stacking` ensemble vs experimental melt curves and melting temperatures for three duplex sequences each synthesized with four different strand compositions (DNA:DNA, DNA:RNA, RNA:DNA, and RNA:RNA) (cf. Figure 9 using `nostacking` ensemble). Computed melt: fraction of nucleotides unpaired, using `stacking` ensemble. Experimental melt: fraction of nucleotides unpaired based on curve fit with 2-state model. Symbols denote $N = 3$ replicate experiments; each experimental curve fit represents the mean of the curve fits for the replicates. Computed and experimental melting temperatures ($T_m$) depicted as vertical lines. Mean absolute error (MAE) $\pm$ SEM for $N = 3$ replicate experiments. Each strand: 5 µM. Salt conditions: 1.0 M $Na^+$.

### S8 Validation studies

Unit tests and regression tests were performed for all combinations of the following test scenarios:

- Number of strands per complex (1 scenario): 3-stranded complex.
- Number of nucleotides per strand (1 of 2 scenarios): 30 nt per strand for tests that involve comparisons between output from two different dynamic programs, 8 nt per strand for tests that involve comparison between output from a dynamic program and output from an exhaustive enumeration algorithm.

- Structural ensemble (2 scenarios): without coaxial and dangle stacking (`nostacking` ensemble) or with coaxial and dangle stacking (`stacking` ensemble).
- Single-material types (3 scenarios): RNA, DNA, or 2′OMe-RNA, with parameter sets `rna06`, `dna04.2`, or `merna06`, respectively.
- Mixed-material types (2 scenarios): RNA/DNA mixtures and RNA/2′OMe-RNA mixtures. For RNA/DNA mixtures, `rna06`, `dna04.2`, and `rna_dna06` parameters were used for RNA, DNA, and mixed RNA/DNA, respectively. For RNA/2′OMe-RNA mixtures, `rna06`, `merna06`, and `rna_merna06` parameters were used for RNA, 2′OMe-RNA, and mixed RNA/2′OMe-RNA, respectively.
- Mixed-material pattern (3 scenarios): two strands with one material and one strand with the other, alternating materials for each nucleotide, random material for each nucleotide.
- Number of sequences for each test scenario: 30 random sequences are generated for each test scenario.

#### S8.1 Unit tests

Over 250 C++ unit tests were run to validate the mixed-materials algorithm. Some unit tests compare the output from a dynamic program to output from an exhaustive enumeration algorithm (see Section S7.3 of Reference 1). This type of comparison can be done only for small complexes because the time complexity of exhaustive enumeration is exponential in the number of nucleotides. An exhaustive enumeration algorithm explicitly enumerates all possible stacking states in the complex ensemble (`stacking` ensemble) or all possible secondary structures in the complex ensemble (`nostacking` ensemble). For a given secondary structure, each loop is categorized as either single- or mixed-material, the appropriate parameters are applied within the loop, and the free energy of the secondary structure is calculated using equation (1).

- **Comparisons between dynamic programing and exhaustive enumeration for the following physical quantities:**
  - **Partition function:** exhaustive enumeration is used to calculate $\overline{Q}(\phi)$ using the sum of (S60) and post-processed to calculate $Q(\phi)$ (see Section S5 of Reference 1). Compare to dynamic programing post-processed output $Q(\phi)$ to the relevant floating point precision.
  - **Equilibrium base-pairing probability matrix:** exhaustive enumeration is used to calculate $\overline{P}(\phi)$ using the sum of (S62). Compare to dynamic programing output to the relevant floating point precision.
  - **MFE:** for the `stacking` ensemble, exhaustive enumeration is used to calculate the free energy of the MFE stacking state, $\overline{\Delta G}(\phi, s^{\shortparallel}_{\text{MFE}})$, using the minimization of (S63); for the `nostacking` ensemble, this reduces to calculating the free energy of the MFE secondary structure, $\overline{\Delta G}(\phi, s^{\text{MFE}})$. Compare to dynamic programing output to the relevant floating point precision.
  - **MFE proxy structure:** for the `stacking` ensemble, exhaustive enumeration is used to calculate the MFE proxy structure(s) $s_{\text{MFE}'}$ using equation (S64); for the `nostacking` ensemble, this reduces to calculating the MFE secondary structure(s), $s_{\text{MFE}}$. The dynamic program for MFE proxy structure determination is the the suboptimal secondary structure algorithm with $\Delta G_{\text{gap}} = 0$ kcal/mol. Compare to dynamic programing output.
  - **Suboptimal structures:** for the `stacking` ensemble, exhaustive enumeration is used to calculate the set of suboptimal secondary structures, $\overline{\Gamma}_{\text{subopt}}(\phi, \Delta G_{\text{gap}})$ within a specified $\Delta G_{\text{gap}} \geq 0$ of the MFE stacking state using equation (S65). Tests are performed using $\Delta G_{\text{gap}} = 0.4$ kcal/mol. Compare to dynamic programing output.

- **Comparison of equilibrium base-pairing probability and Boltzmann sampled structure dynamic programs.** Estimate equilibrium the base-pairing probability matrix from Boltzmann-sampled structures and check convergence to the exact base-pairing probability matrix as the number of sampled structures increases.

- **Comparison of MFE and free energy of MFE proxy structure.** Calculate the free energy of the MFE stacking state, $\overline{\Delta G}(\phi, s^{\shortparallel}_{\text{MFE}})$ using the forward dynamic program. Calculate the MFE proxy structure, $s_{\text{MFE}'}$, using the backtracking dynamic program for suboptimal structure generation with $\Delta G_{\text{gap}} = 0$ kcal/mol. Calculate the free energy of the MFE proxy structure using equation (1) and compare it to the value calculated using the forward algorithm to the relevant floating point precision.

- **Overflow-safe partition function evaluation algebra:** Check that overflow-safe partition function agrees with non-overflow-safe partition function (see Sections S3.1.4 and S6.5 of Reference 1). Compare 32 bit overflow-safe with 64 bit non-overflow-safe.

#### S8.2 Regression tests

Regression tests were performed by comparing output from mixed-material dynamic programs to output from single-material dynamic programs (NUPACK 4.1).[1]

- **Comparison of single- and mixed-material algorithms on single-material complexes.** For single-material complexes, single- and mixed-material algorithms should return identical calculations for any physical quantity. Comparisons are made for the following physical quantities: 1) partition function, 2) equilibrium base-pairing probability matrix, 3) MFE, 4) MFE proxy structure, 5) suboptimal structures.

- **Comparison of single- and mixed-material algorithms on mixed-material complexes using only mixed-material parameters.** For mixed-material complexes: the single- and mixed-material algorithms are used with only mixed-material parameters (either `rna_dna06` or `rna_merna06`) so that all loops are treated with mixed-material parameters even if a given loop is single-material. This scenario allows the single- and mixed-material algorithms to be compared for mixed-material complexes. Comparisons are made for the following physical quantities: 1) partition function, 2) equilibrium base-pairing probability matrix, 3) MFE, 4) MFE proxy structure, 5) suboptimal structures.
